# Cancer Metastases of 202 Kinds of Cancers via Cytocapsular Tubes

**DOI:** 10.1101/2021.05.13.443996

**Authors:** Tingfang Yi, Gerhard Wagner

## Abstract

**BACKGROUND:** Cancer metastasis is the primary source of solid cancer lethality, but the underlying mechanisms have essentially remained elusive. Recently, we found that cytocapsular tubes, a newly discovered organelle of mammalian cells, conduct cell translocation, suggesting an efficient pathway for cancer cell metastases.

**METHODS:** We performed immunohistochemistry staining and fluorescence microscope imaging analyses of 6 kinds of normal organs (n=14 patients), 38 subtypes of benign tumors (n= 126 patients), and 8,061 clinical solid cancer tissue samples (covering 202 types and subtypes of cancers, and including cancers, paracancer tissues and metastatic cancers) taken from 7,125 cancer patients. These solid cancers cover 30 types of organs. We characterized cytocapsular tubes (CTs), cell migration in CTs, and CT quantity, density, network, superstructures and lifecycle in clinical samples of normal tissues, benign tumors, cancers, paracancer tissues, and metastatic cancers.

**RESULTS:** There is no cytocapsular tubes (CTs) in normal organ tissues. Among the 126 benign tumor samples (covering 38 subtypes) from 126 patients, 86.6% do not have CTs and the other 13.4% show benign-to-malignancy transitions with CTs. Of the 8,061 solid cancer tissue and paracancer samples from 7,125 patients, 100% of cancers (including carcinomas *in situ*), 100% of paracancer tissues, and 100% of metastatic cancer tissues/organs, harbor large quantities (from thousands to hundreds of thousands) of CTs with many cancer cells in migration inside.

**CONCLUSION:** Cytocapsular tubes and networks provide membrane-enclosed physical freeway systems for clinical cancer cell metastasis to neighboring and far-distance tissues and organs.

## Introduction

Cancer is a leading cause of human mortality^1^. It has been estimated that approximately 19.3 million new cancer cases and approximately 10 million cancer deaths occurred worldwide in 2020 alone^1^. Cancer metastasis is the major source of cancer lethality^2–6^. The verifiable identification of cancer metastasis mechanisms is a prerequisite for the effective prognosis, diagnosis, pharmacotherapy and treatment for patients with suspicious, progressive, symptomatic, metastatic and recurrent cancers, for effective drug development, for minimizing side effects and sufferings, and for reliably determining the benefits of treatments. Prognosis, diagnosis and treatment responses are usually assessed with the application of radiographic/ MRI imaging at millimeter (mm) level, which fails to detect the cancer metastasis pathways at micrometer (μm) level^7–9^. The methods of precise identification of initial cancerous transformation at μm level in tissues are absent. Therefore, there is an urgent need for verifiable identification of cancer metastasis mechanisms and biomarkers that effectively detect cancer metastasis physical pathways with high sensitivity and specificity.

Cancer cell metastasis proceeding unprotected through tissues *in vivo* would encounter numerous heterogeneous neighbor cells and diverse extracellular matrices (ECM), all of which function as both migration clues and substantial environmental obstacles impeding the migration and long distance translocation of cancer cells. Although a number of hypotheses have been proposed^10–16^, consistent models are still missing that could explain the mechanisms underlying cancer metastasis to different sites.

Recently, we discovered that previously unknown membrane-enclosed organelles, dubbed cytocapsular tubes (CTs)^17^, can conduct efficient cell translocation (Fig. S1A; Movies S1-2). We developed anti-CM-01 antibodies that recognize a CT membrane protein marker *in-vitro* (Fig. S1A). We have now used the same membrane protein marker, to investigate CTs in 8,061 solid cancer and paracancer tissues from 7,125 cancer patients (covering 202 kinds of cancers, and including cancers, paracancer and metastatic cancer tissues). We found that the cancer cells in all examined 202 kinds of cancers employ a similar mechanism for near- and long-distance metastases based on CT networks.

## Methods

### Patients and Sample Collection

The sources of clinical samples are described in Fig. 1. The patients providing formalin-fixed paraffin-embedded (FFPE) tissue samples gave informed consent that indicated that they understood that the biopsies (needle biopsy or surgical biopsy) were performed for research purposes only. Comparative deidentified samples of normal tissues, benign tissues, carcinoma *in situ*, cancer, paracancer, metastatic tissues with their cancer stages identified according to the TNM system (cancer stages: 0, I, II, III, IV) were obtained from archival materials. The cancer, paracancer and metastatic cancer tissues, in which the original cancer niches were identified by indicated cancer specific molecular markers, were identified by hospital pathology laboratories and obtained from archival materials.

**Figure 1.**
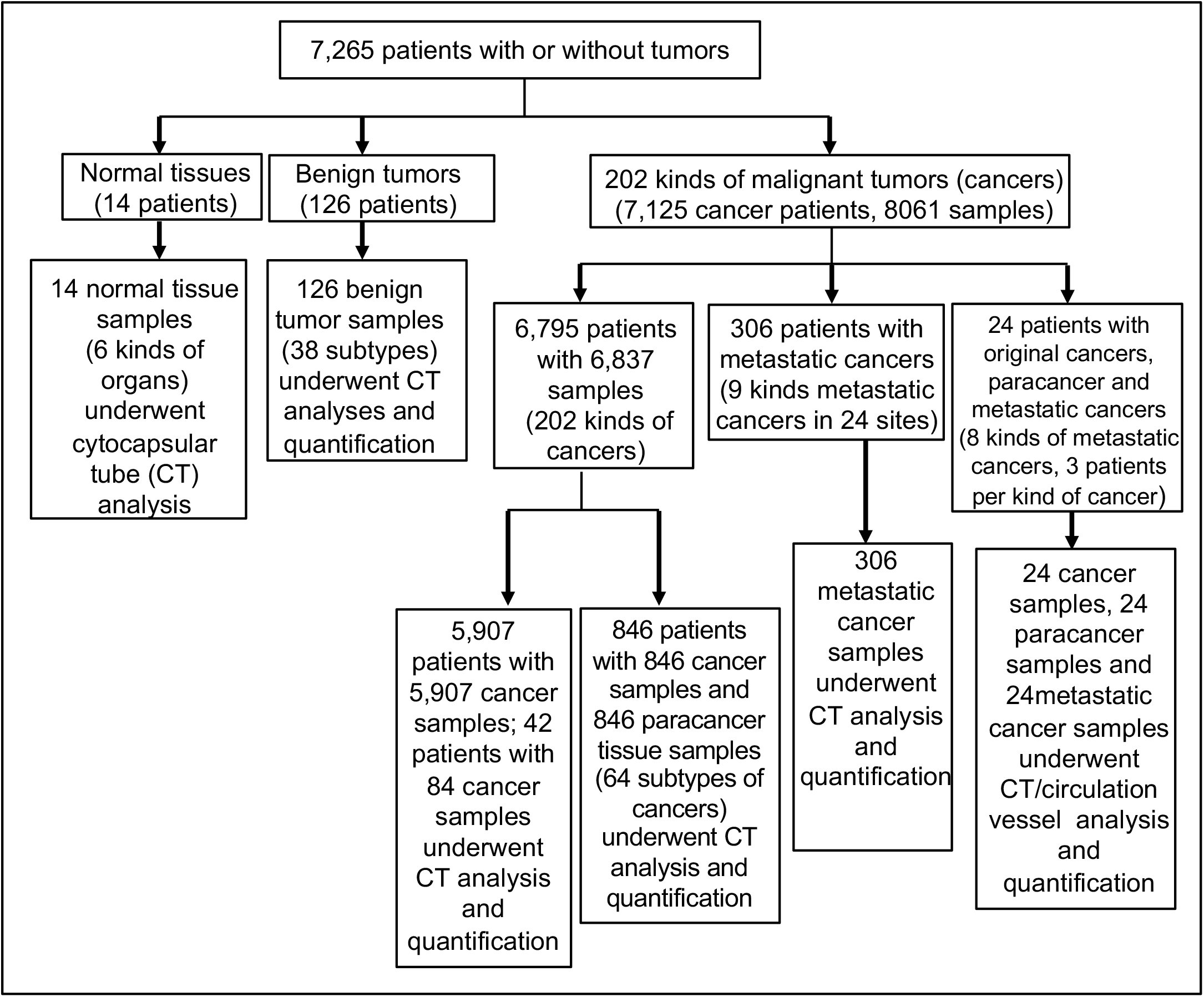
Origen and Enrollment of Clinical Samples from 7,253 Patients. In the 7,253 cancer patients, 846 cancer patients provided both cancer and paracancer tissues, 42 cancer patients provided 84 samples (2 tissue samples per patient), and 5,907 cancer patients provided one tissue sample each. In the 24 cancer patients, each provided with 1 original cancer sample, 1 paracancer sample and 1 metastatic cancer sample.

### Histology and Immunohistochemical testing

Cancer lesion samples were obtained from the index patients from clinical hospitals in USA (8,036 specimens from 7,088 patients, US Biomax; others samples directly from clinical hospitals). Samples were processed for histopathological evaluation with the use of hematoxylin and eosin (H&E) staining. Immunohistochemical (IHC) fluorescence tests were performed to stain CTs using rabbit anti-CM-01 monoclonal and polyclonal primary antibodies (self-made), mouse anti-gamma-actin monoclonal primary antibodies (Abcam, ab123034) (Fig. S1).

### Data collection

Cytocapsular tubes (not sectioned, longitudinally sectioned, and cross sectioned) without degradation (3∼6μm in measured diameter) were counted using a fluorescence microscope and ImageJ. The presence of CTs degrading into thick strands (1∼2μm in measured diameter), thin strands (0.2∼1μm in measured diameter), or the disintegration state were reported without quantification.

### Statistical analysis

For each specimen, the number of fully intact cytocapsular tubes was counted in 5 areas (0.35mm x 0.35mm, length x width) of the sample (top, bottom, left, right, and center), and the CT density (CT/mm^2^) was calculated and determined for each area. The average CT density across the 5 sites was treated as the specimen’s overall CT density.

## Results

### Cancer metastasis via cytocapsular tubes in 202 kinds of cancers

#### Normal tissues

In the 6 types of normal tissues analyzed immunohistochemically (breast, colon, liver, lung, prostate, and stomach; n=1 to 3 patients for each type, n=14 patients in total), no cytocapsular tubes were found (CT, 0%) (Fig. 2A **and** Fig. S2A, 2D,).

**Figure 2.**
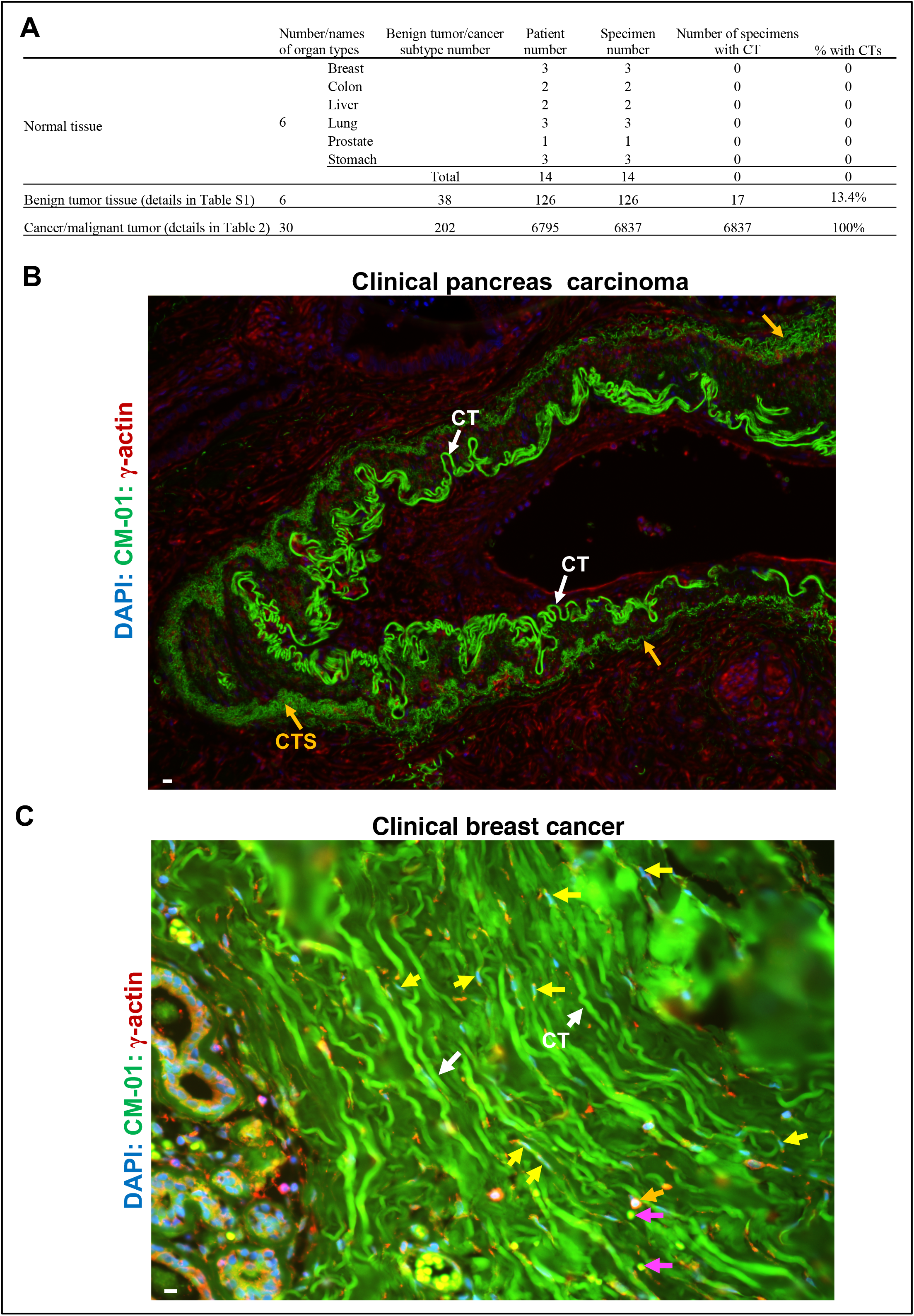
Cytocapsular tubes that are ubiquitously present in clinical solid cancer tissues (but not in normal tissues or benign tumors) are membrane-enclosed, physical freeways of cancer cell metastasis. **A:** Comprehensive analyses of 6 kinds of normal human tissues, 38 kinds of benign tumors in 6 kinds of tissues, and 202 types/subtypes of solid cancers in 30 kinds of tissues evidence that CTs, which are ubiquitously distributed in solid cancers but not in normal tissues or non-transformed benign tumors, are physical freeways of cancer cell metastasis and dissemination. **B:** Representative fluorescence microscope image of a clinical pancreas carcinoma stained with DAPI, anti-CM-01 and anti-γ-actin antibodies in immunohistochemistry (IHC) assays. A long and highly curved, membrane-enclosed cytocapsular tube (CT) with many hairpins, turns and coils (white arrows, measured length is approximately 5400μm in the sectioned specimen analyzed by ImageJ), forming a super-large structure in an irregular morphology. Outer side that well-formed structure, there are many CT strands (CTS) degraded from another layer of super-large CT structure(s). **C:** Representative fluorescence microscope image of a clinical breast cancer with many aligned CTs (white arrows) and CT bunches. Breast cancer cells (yellow and orange arrows) migrate in the long and membrane enclosed freeways of CTs. Indicated are: longitudinally sectioned CT fragments with longitudinally sectioned cells migrating in CTs (yellow arrows, in a thin and long morphology shaped by CT membranes), cross sectioned CTs with cross sectioned cells in migration in CTs (orange arrows), and cross sectioned CT surfaces without cells (purple arrows). Scale bar, 10μm.

#### Benign tumors

Of the 6 types (breast, lung, prostate, stomach, liver, colon; covering 38 subtypes) of clinical benign tumor samples (n= 1 to 15 patients of each subtype, n=126 patients in total), approximately 86.6% don’t present CTs; however, approximately 13.4% harbor CTs, displaying the transition of a benign tumor to a malignant tumor (Fig. 2A**, and** Fig. S2B, 2E **and** Table S1). CTs are invisible in conventional hematoxylin & eosin staining assay or in IHC staining assays with multiple popular cancer protein marker antibodies (such as anti-ER, anti-PR and anti-HER2 antibodies for breast cancer; anti-MSH-2 antibody for colon cancer) (Figs. S2-3), but visible in IHC staining assays with anti-CM-01 antibodies in multiple types of cancers, including cancers in breast, colon, oral and thyroid (Figs. S2-S5).

#### Carcinoma in situ

This is the early stage before spreading. In breast carcinoma in situ tissues (n=12), 100% of them have plenty of CTs and the average CT density is high up to 106 CT/mm^2^ (**Excel S1**).

#### 202 subtypes of cancer

Next, we performed CT analyses in 202 subtypes of cancers. In the 6, 837 cancer tissue samples from 6,795 cancer patients (covering 202 subtypes of cancer tissues), all of them (100%) harbor CTs with various CT densities, and cancer cells were found in migration inside the CTs (Figs. 2-3, S2**, Tables 1-2**, **Excel S1-2**). These 202 subtypes of cancers cover up to 30 kinds of organs: adrenal gland, bladder, bone, brain, breast, cervix uteri, colon, esophagus, gallbladder, head/neck, intestine, kidney, liver, lung, lymph node, oesophagus, oral cavity, ovary, pancreas, penis, prostate, rectum, skin, spleen, stomach, testis, thymus, thyroid, uterus, and vulva (n=3∼2774 patients per type of organ). The CT densities in cancers of these 30 kinds of organs vary in a range from 6 to 111 CT/mm^2^ (Figs. 2-3**, Excel S1-2**).

**Figure 3.**
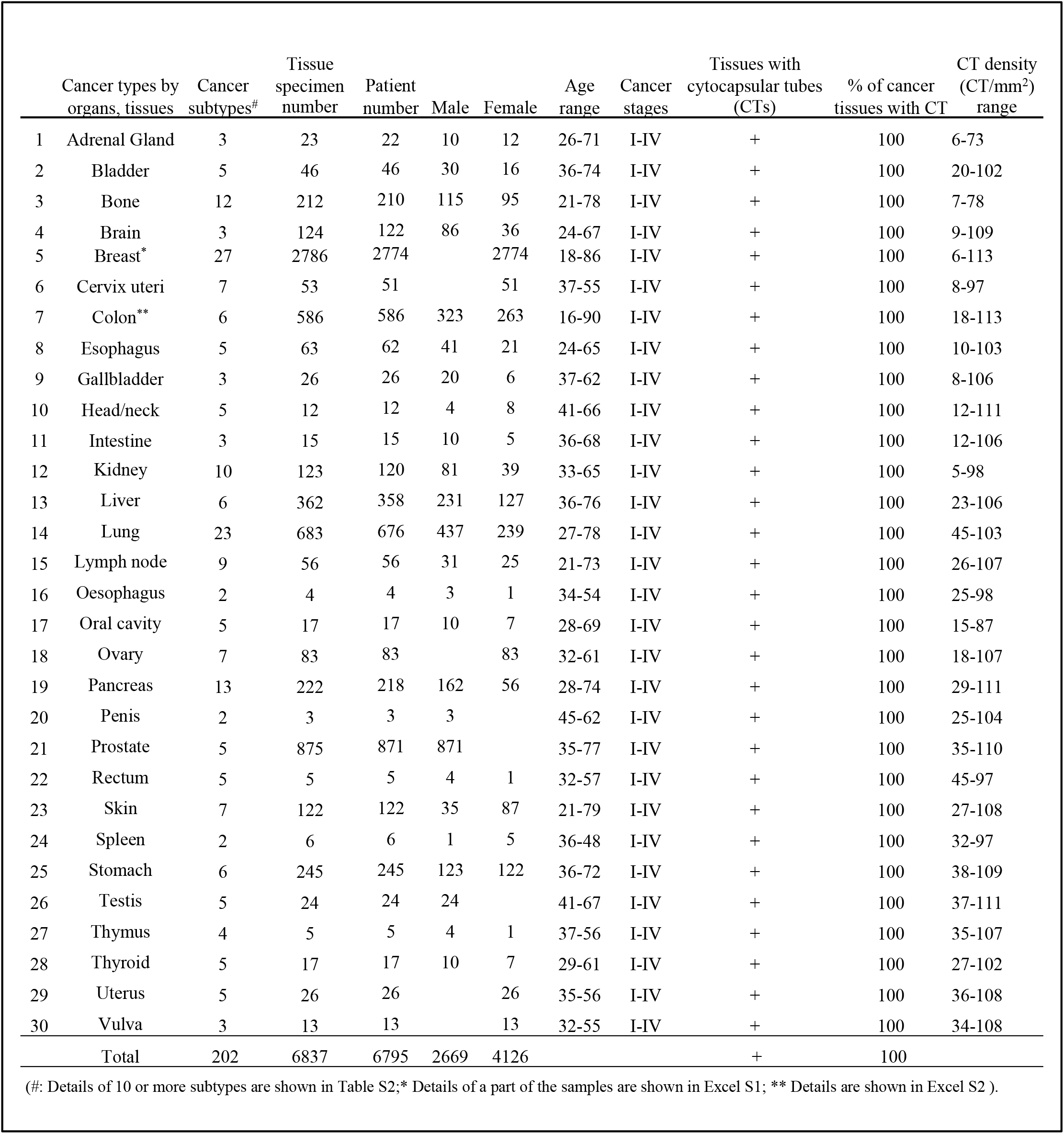
Characterization of cytocapsular tubes in 202 kinds of clinical cancers in 30 types of human organs. The cancer patient and tissue sample collection underwent randomization. The cytocapsular tube presence, percentages and densities in all the 6,837 examined cancer tissue samples from 6,795 cancer patients (covering 202 kinds of solid cancers and 30 types of organs).

#### Paracancer

This is a state tissues must pass towards solid cancer metastasis. We performed IHC staining analyses with clinical paracancer tissues (namely “normal adjacent tissue, NAT”) of breast and prostate cancers, and found that these paracancer tissues contain plenty of CTs and CT networks with many cancer cells migrating in CTs (Fig.2). Next, we performed IHC staining analyses with paracancer tissues of 12 types of cancers (covering 64 subtypes of cancers) in bladder, breast, esophagus, colon, intestine, kidney, liver, lung, ovary, pancreas, stomach and uterus (Fig. 4). In these paracancer tissues of 64 subtypes of cancers (n=3∼365 samples per subtype), 100% paracancer tissues harbor plenty of CTs. The CT densities of these paracancer tissues range from 67 to 114CT/mm^2^ (Fig. 4). There are many cancer cells migrating in all the CTs in paracancer tissues of all the 64 cancer subtypes (Fig. 4).

**Figure 4.**
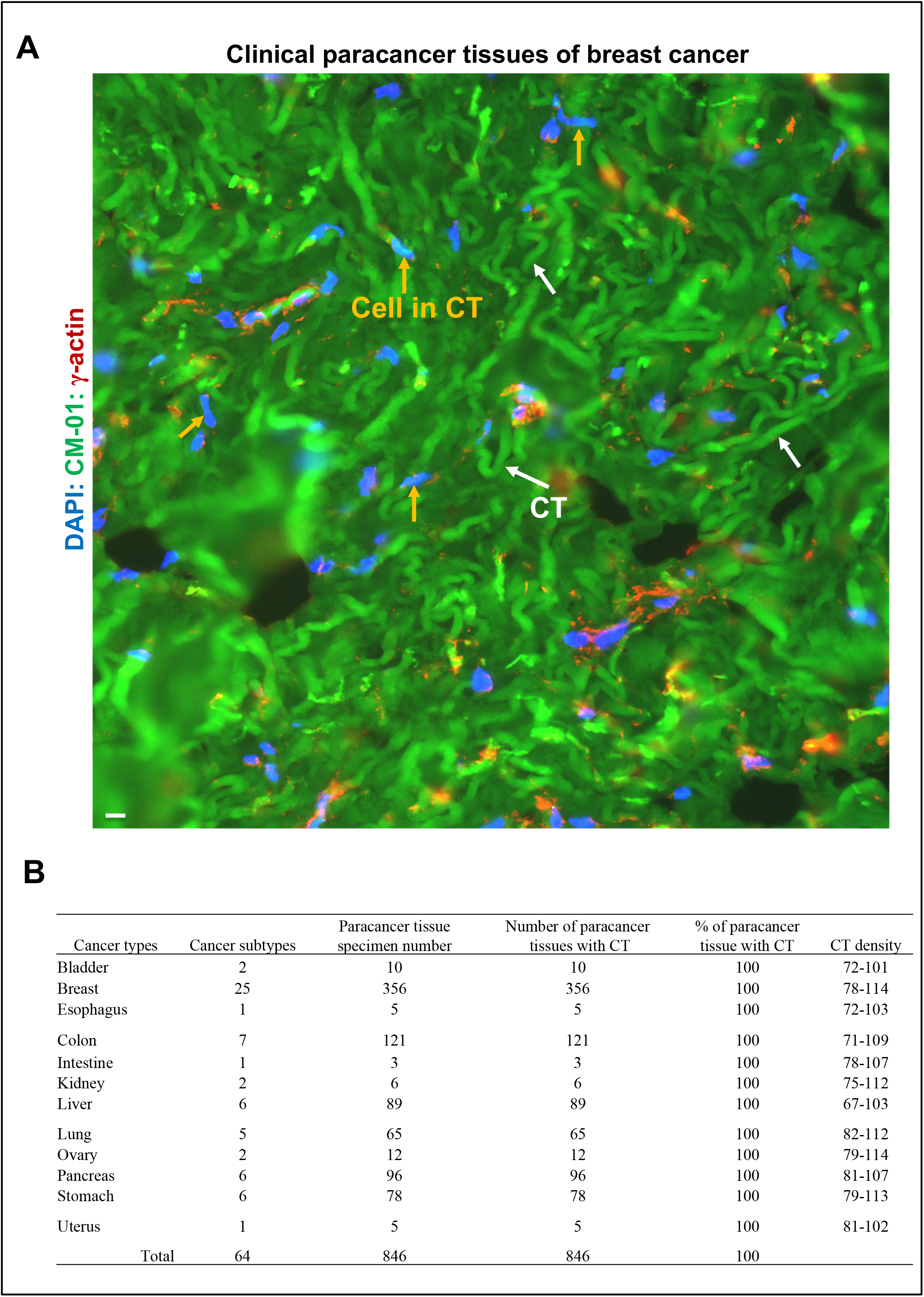
Cytocapsular tubes are massively distributed in paracancer tissues. **A:** Representative fluorescence microscope image of paracancer or “normal adjacent tissues (NAT)” of a breast cancer with massive long, curved and entangled CTs (green color, white arrows) conducting cancer cell metastasis. There are many breast cancer cells in migration (orange arrows, the nuclei are in blue color) in the thousands of thousands of massive CTs. These images show that previously thought “normal adjacent tissues” are not real normal tissues, but harbor thousands or millions of membrane-enclosed highways of CTs conducting massive cancer cell dissemination. **B**: Table of characterization of CTs in the examined clinical paracancer tissues of 64 subtypes of cancers in 12 kinds of organs.

#### Metastatic cancers

Subsequently, we investigated CTs in clinical metastatic mucinous adenocarcinoma from colon in liver, metastatic adenocarcinoma from ovary in epiploon, metastatic breast adenocarcinoma in lymph node, and metastatic mucinous adenocarcinoma from unknown sites in epiploon, 100% of them harbor numerous CTs with many cancer cells migrating in CTs (Fig. S6). We performed CT analyses in 9 types of metastatic cancer tissues in multiple metastatic sites, and found that 100% of these clinical metastatic cancers harbor a lot of CTs with many cancer cells in migration in CTs (Fig. 5A). We performed CT analyses in more metastatic cancers with successive cancer-paracancer-metastatic sites in breast, colon, pancreas and ovary, 100% of these samples along the cancer-paracancer-metastatic site axis harbor plenty of CTs with increased CT density in paracancer tissues comparing to cancer original niches and metastatic cancers (Fig. 5B).

**Figure 5.**
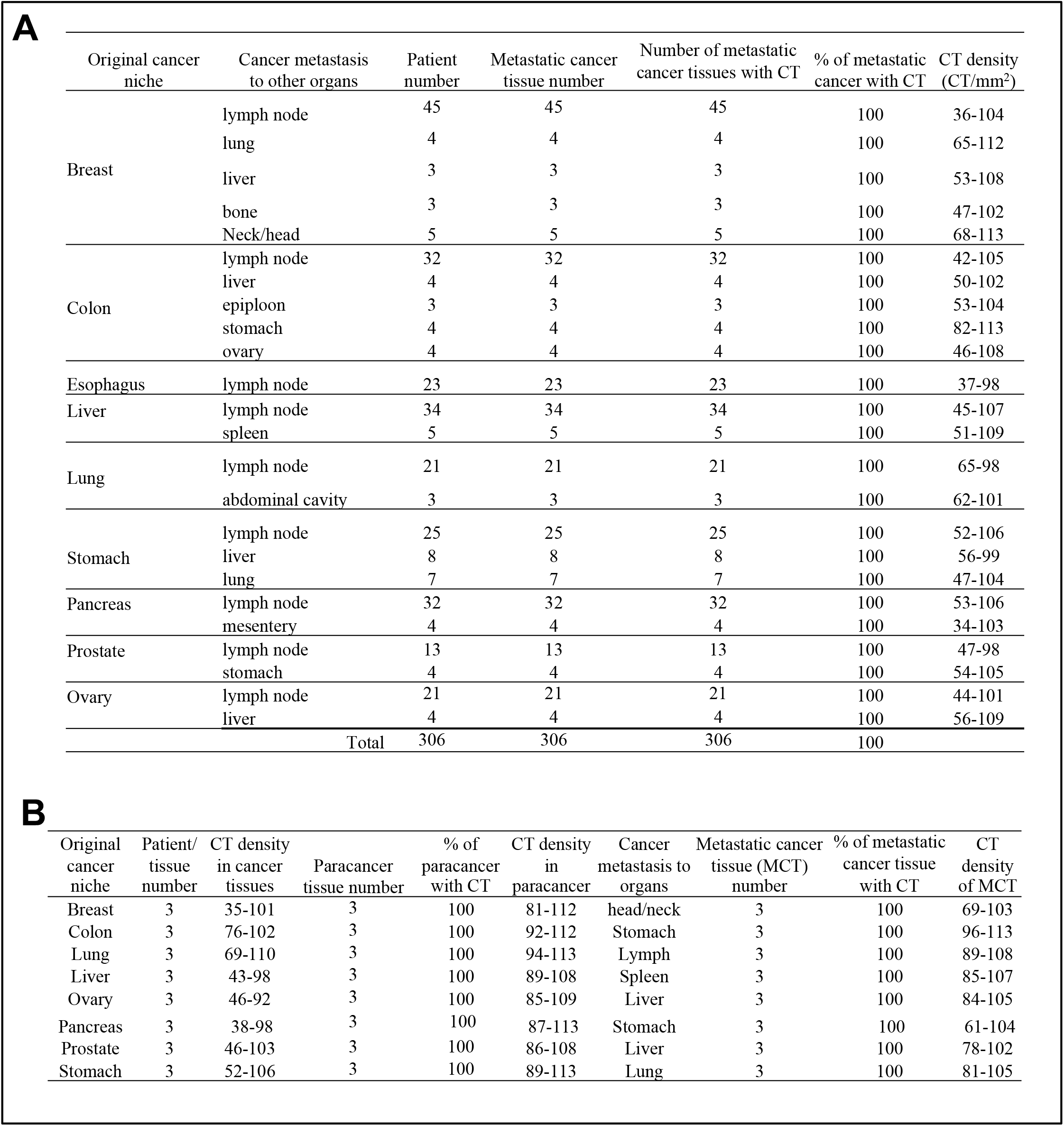
Cytocapsular tubes (CTs) conduct crossing-organ cancer metastasis in solid cancers. **A:** Characterization of CTs in crossing-organ cancer metastasis from original cancer niches (in 9 kinds of organs) to multiple far-distance and crossing-organ dissemination. **B**: Characterization of CTs in 8 kinds of cancers with crossing-organ metastasis from examining CTs in original cancer niches, paracancer tissues, and metastatic cancer tissues.

### Lifecycle of cytocapsular tubes in clinical tissues

We analyzed the acquired >60,000 frames of IHC fluorescence microscope images of the examined 8,061 cancer/paracancer/metastatic samples taken from 7,125 patients covering 202 kinds of cancers, and found variations in CT quantity, density, morphologies, distribution, CT networks, stages in CT life cycle, and superstructure textures, displaying a huge diversity of CTs and networks in all kinds of clinical cancers. Thus, based on the comprehensive analyses of these images, we can identify a lifecycle of cytocapsular tubes in clinical cancers. It can be categorized into 5 successive stages: 1) CT initiation; 2) CT elongation, network formation and cancer cell migration inside; 3) CTs and networks invasion into and pass through paracancer tissues; 4) CTs and CT networks cross tissues and organs and reach neighboring and far-distance destinations; 5) CT degradation and decomposition.

In the initiation phase, individual cancer cells generate cytocapsulas that envelope the cancer cells. Cancer cells migrate inside and generate elongated CTs. The rear part of the CTs display shrunk and thin-tailed morphologies as they are not well anchored in the ECM (Fig. S7A). CTs continue to elongate and form long CTs with consistent diameter/width (2∼6μm in diameter). Individual elongated CTs aggressively invade into neighboring loose and compact tissues, even invade through the tightly-connected and thickness-enhanced epithelial layer of terminal ducts in breast cancer (Fig. S7B). Membrane-enclosed CTs interconnect to each other and form superlarge 3D CT networks and provide membrane-enclosed 3D freeway systems that allow multiple migration directions for CT-conducted cancer cell dissemination in tissues (Fig. S8). Elongated CTs generate curved and entangled CT masses that occupy spaces and create CT mass cavities (CMCs). Locally, still-free cancer cells invade into CTs and migrate away via CTs, which creates free (liquid-filled) spaces and cavities that grow in size (Figs. S9A-B). After CT masses degrade and decompose, the empty but liquid filled CT mass cavities (CMCs) are left in the compact cancer tissues (Figs. S9C). Numerous CTs align together and form CT bundles and invade into compact tissues (Fig. S10A). Nearby cancer cells invade into CTs and migrate away, CT masses occupy the spaces while few/no cancer cells remain outside of CTs (Fig. S10B). Cancer cells generate highly curved and coiled CTs, and invade into compact tissues in multiple directions, providing more freeways for cancer cell metastasis (Fig. S11). Free cancer cells invade into CTs generated by other cancer cells, make more CTs and CT networks, and escape to neighboring and far-distance destinations, leaving no free cancer cells beyond CT masses (Figs. S12-13). Cancer cells can generate highly curved CT masses and big curved CT masses, which provide more closed CT freeways for cancer cell invasion into CTs for metastasis (Fig. S14). Cancer cells generate multiple layered, highly curved, coiled superstructures invading into compact tissues, providing large CT contacting surfaces for cancer cell invasion into CT networks for cancer cell metastasis (Fig. S15). During CT conducted cancer metastasis, cancer cells engender many kinds of different complex superstructures for invading into and pass through diverse compact tissues (Fig. S16), huge, dozens-layers complex superstructures invading into tissues with diverse densities (Fig. S17), highly compact cone-shaped superstructure (Fig. S18A), or multiple layered complex superstructures (Fig. S18B). Massive CT clusters can powerfully invade into all kinds of loose and compact tissues (Figs. S19-20). Straight, curved and coiled CTs interconnect together and form large CT networks for cancer metastasis. Even though some parts of CT networks decompose due to CT degradation, the left CT networks remain capable of providing CT efficient freeway systems for cancer cell metastasis progression. Thus, CT networks display potent damage resistance capacities and provide efficient CT freeway systems for cancer metastasis (Fig. S21). Subsequent CT degradation employs a multiple-step format: 1) numerous dot membranes on CT surfaces collapse, 2) CTs form a thin CT strand-interconnected network in tube shapes; 3) thin CT strands decompose, disconnect and form individual CT strands; 4) disconnected thin CT strands, decompose and disappear in a complex manner (Figs. S22-24).

The numbers of CTs are always much larger than those of circulating system vessels (blood/lymph vessels) (Fig. S25A). In most instances, plenty of CTs pass through, surround and embed circulating system vessels (blood vessels and lymph vessels) without invading into blood vessels and lymph vessels (Fig. S25A). In the examined 8 kinds of organs, the average number ratio of CTs versus blood and lymph vessels in cancers, paracancer tissues and metastatic cancers is around 1000 (Fig. S25B). In the late (but not early) stage cancers (stage III or IV), CTs invade into the tightly connected endothelial layers of approximately 1 in 10^4^ blood/lymph vessels and release cancer cells into circulation system vessels (Fig. S26A). Approximately 1% of CT invaded blood/lymph vessels present leak/disassembly followed by local blood cell spreading (Figs. S26B-C). CT analyses with needle biopsy samples provide a linear CT scan in multiple-layer tissues (Fig. S27).

### Cytocapsular tube based cancer metastasis grading

Based on the above observations and data of cancer CT numbers, density, degradation, and metastasis situations of the 8,061 solid cancer and paracancer tissue samples from 7,125 cancer patients (covering 202 kinds of cancers), we suggest a five stage CT-based cancer metastasis (CM) grading reference: CM0, CM1, CM2, CM3, and CM4 (Table S3). On the basis of the comprehensive analyses of the cancer stages and CT based CM grade of 7251 benign tumor and cancer patients, we found relationships between cancer stages and CT based CM grades listed in Table S4.

## Discussion

It is vital to understand the long-term elusive mechanics underlying cancer metastasis^18–25^. Here, we investigated the metastasis mechanics of 202 kinds of clinical cancers, and evidenced that cancer cells generate substantial membrane-enclosed freeway systems formed by cytocapsular tubes and CT networks that conduct cancer-cell metastases to neighboring and far-distant tissues and organs; this was found to be the case in all 202 kinds of solid cancers tested.

Our data indicate that cancerous cells upon becoming malignant decide to generate extra-cellular cytocapsular tubes initiated by a not-yet-known mechanism. CTs are used by the originator cells and other cancer cells. From there on, cancer cells can migrate in large quantity CTs and protected from outside attacks. CTs can be very long consistent with far-distance metastasis. While most of our data shown here are obtained from snapshots of clinical cancers samples received from cancer tissue banks, the emerging picture is confirmed by clinal data from live cancer patients with cancers in 9 kinds of organs: breast, colon, lung, oral, pancreas, prostate, skin, testis, and thyroid.

It was unforeseen that cytocapsular tubes would be ubiquitously used as physical freeways for cancer metastasis in all 202 kinds of examined solid cancers. Membrane-enclosed homogeneous CTs shield cancer cells from external heterogeneous obstacles of neighbor cells and ECM, and significantly facilitate efficient cancer cell dissemination. 3D CT networks provide substantial physical freeways leading in multiple and expanding directions for cancer cell metastases to far-distance tissues and organs. Collective and self-organization activities of long CTs, large networks and superstructures augment collective cancer metastasis through dense, compacted and diverse tissues, cross organs and reach far-distance destinations. The number of cancer cell-filled CTs is approximately 1000-fold larger than that of blood and lymph vessels, and only approximately one 1 of 10,000 blood/lymph vessels present CT-vessel invasion in late stage cancers where cancer cells are released into the circulation systems. This is consistent with clinical observations that there are rare circulating cancer cells (CTCs) in the circulation system^26–28^. These above observations evidenced that the ubiquitous and large quantities of CT networks provide the major physical pathways for neighboring and far-distance cancer metastases in cancer patients.

It is essential to precisely evaluate the cancer metastasis status in prognosis, before and after treatments, and during cancer management to achieve better outcomes for patients. So far, a quantitative and precise cancer metastasis evaluation system is absent^2–3, 29^. The cytocapsular tube-based cancer metastasis (CM) grading described here may facilitate the accurate evaluation of cancer cell status and metastasis, thereby improving therapy outcomes. To take advantage of the possibilities for the precise and qualitative evaluation of CT-based cancer metastasis, more clinical analyses are required. In the future, we expect that CT-based drug and technology development will significantly improve cancer pharmacotherapy and management.

## Acknowledgements

Supported by a grant (to Dr. Yi) from Cytocapsula Research Institute Fund for Cytocapsular Tube Conducted Cancer Metastasis Research, and a grant (to Dr. Yi) from Cellmig Biolabs Inc Fund for Cytocapsular Tube Cancer Metastasis Research. We greatly acknowledge Dr. Ed Harlow, Harvard Medical School (USA) and Cancer Institute of University of Cambridge (UK), Dr. Jon Aster, Harvard Medical School, Dr. Jelle Wesseling, Leiden University (Netherlands), Dr. David Golan of Harvard Medical School (USA), and Steven Ganshirt, MD of Northwestern University Medical School for their help and meaningful discussion in the study.

## Supplementary Methods

### Proteome characterization of cancer cell cytocapsulas and cytocapsular tubes and cytocapsular tube protein marker identification

Multiple stable dual SILAC labeled human cancer cells (100% SILAC labeling with ^13^C_6_, ^15^N_2_-Lys and ^13^C_6_, ^15^N_4_-Arg) of pancreas cancer cell Bxpc3, non-small cell lung cancer cell A-427, breast cancer cell MCF-7, and colon cancer cell SK-CO-1 cells were cultured on 3D-Matrigel matrix to grow cytocapsulas and cytocapsular tubes^1^. With cytocapsulae and cytocapsular tube ecellularization and ecellularized cytocapsulae and cytocapsular tube collection methods, SILAC labeled, ecellularized, and cell-free cytocapsulas and cytocapsular tubes were collected, followed by SDS-gel purification and LC-MS/MS proteome analyses. We identified 412 cytocapsula and cytocapsular tube proteins. Protein marker candidate screenings were performed in cancer cell cytocapsulae and cytocapsular tubes *in vitro* with antibodies and IHC-fluorescence microscope analyses. An identified cytocapsular tube membrane protein marker named as CM-01 (Cancer Metastasis-01, an established membrane protein) was used for cytocapsular tube analyses in clinical tissues *in vivo*.

## Supplementary Figure Legends

**Figure S1.**
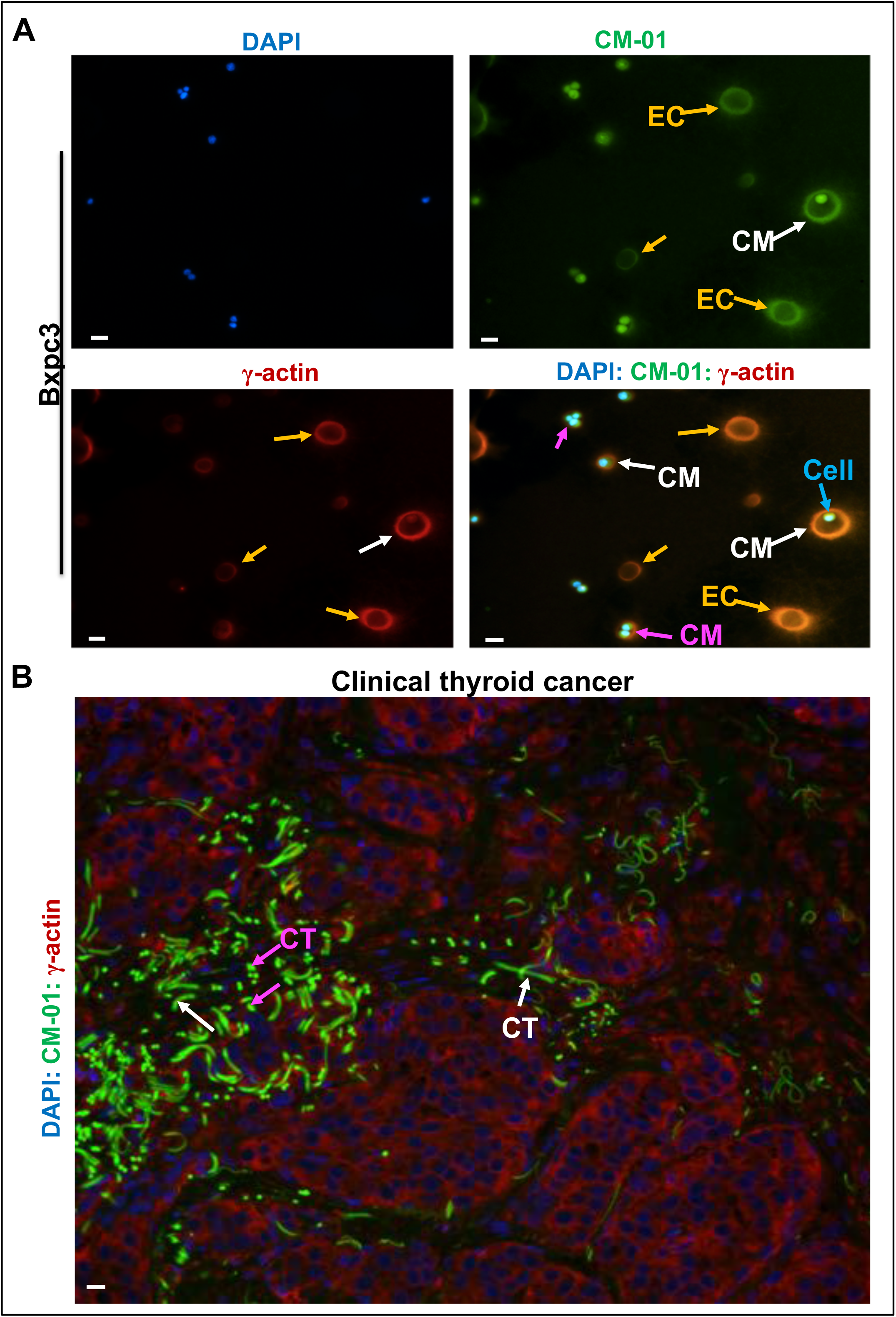
Verification of a cytocapsular tube protein marker in cancer cell cytocapsular membranes *in vitro* and *in vivo*. **A**: Representative fluorescence microscope images of pancreas cancer cell Bxpc3 cytocapsulae with anti-CM-01 and anti-γ-actin antibodies in immunohistochemistry (IHC) assays. Bxpc3 cells were implanted onto and cultured on a 3D-Matrigel matrix. Bxpc3 pancreas cancer cells grow cytocapsulas. After ecellulation, Bxpc3 cells existed and left ecellulated cytocapsulas (EC, orange arrows). Cytocapsular membranes (CM, white arrows) of single Bxpc3 cells, Cytocapsular membranes enclosing two or more Bxpc3 cells (CM, purple arrows), and cells outside CMs (cyan arrows) are shown. Panels of DAPI staining, anti-CM-01 antibody staining, anti-γ-actin antibody staining, and merge are shown. **B**: Representative image of collective thyroid cancer cytocapsular tubes (CTs) invading into compact thyroid cancer tissues. The bright green color specific for cytocapsular tubes, and the very low green background color in non-cytocapsular tube thyroid cancer tissues document the relative enhanced abundance of the protein marker CM-01 recognized by anti-CM-01 antibody and the specific binding ability of anti-CM-01 antibodies. Longitudinally-sectioned CTs (white arrows) and cross-sectioned CTs (purple arrows) are shown. Scale bar, 10μm.

**Figure S2.**
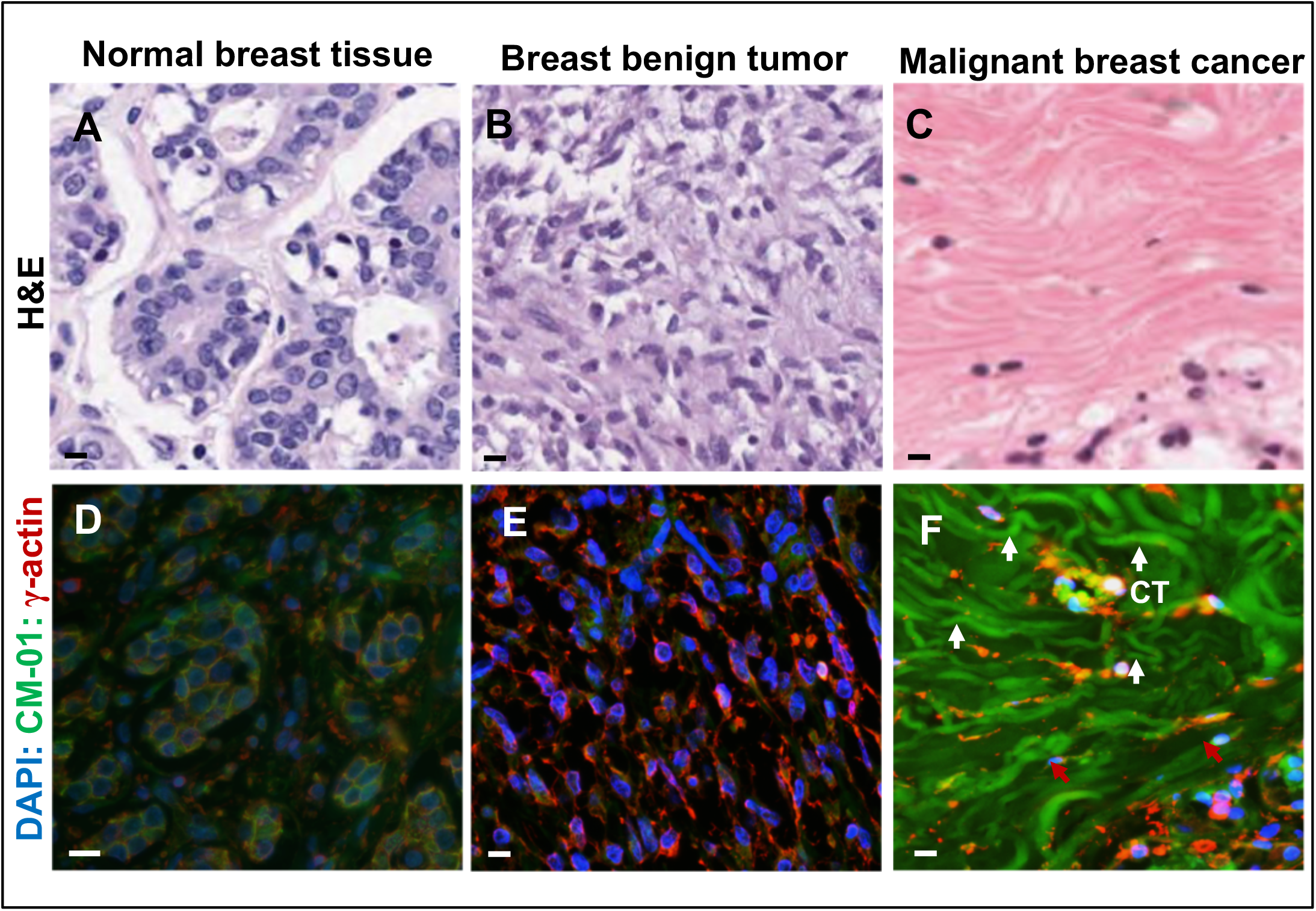
Cytocapsular tubes are present in clinical cancer tissues, but not in normal tissues or benign tumor tissues. **A-C:** Representative images of H&E stained samples of normal clinical breast tissue, clinical breast benign tumor and clinical malignant breast cancer. **D-F**: Representative images of IHC fluorescence images of anti-CM-01 (green), anti-γ-actin (red) and DAPI (blue) stained samples of clinical normal tissue, breast benign tumor and malignant breast cancer, respectively. Conventional H&E staining method does not clearly show cytocapsular tubes (CTs) in malignant breast cancer tissues, while immunohistochemistry (IHC) staining with anti-CM-01 antibodies shows numerous CTs (white arrows) in malignant breast cancer tissues. Scale bar, 10μm.

**Figure S3.**
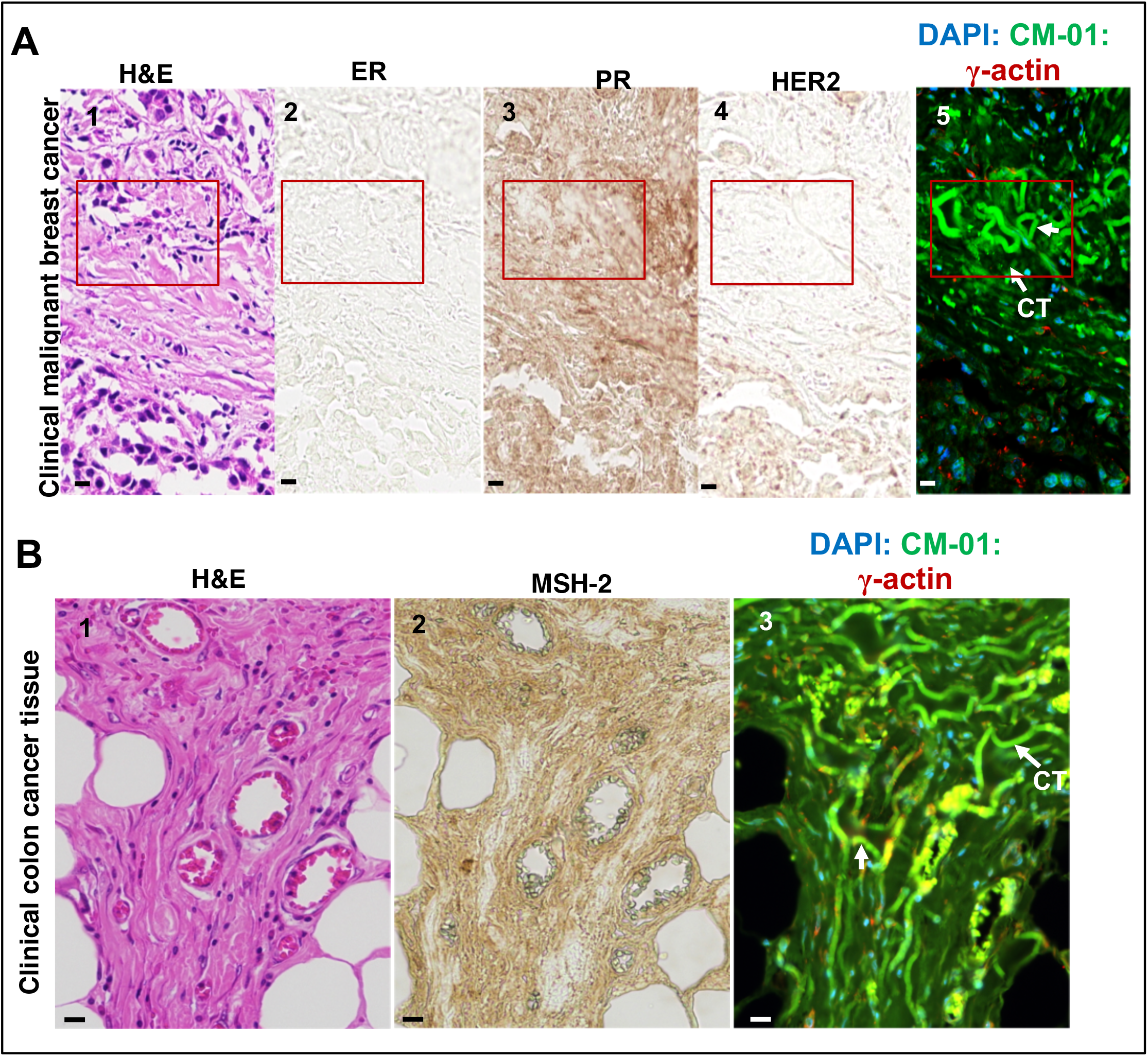
Anti-CM-01 antibodies, but not conventional popular cancer cell marker antibodies or H&E staining, show cytocapsular tubes in clinical cancer tissues. **A:** Representative images of H&E staining (panel 1) and immunohistochemistry (IHC) staining with antibodies recognize ER (estrogen receptor, panel 2), PR (Progesterone receptor, panel 3) and HER2 (human epidermal growth factor receptor 2, panel 4), and anti-CM-01 antibody (panel 5) with clinical breast cancer tissues. Only IHC staining with anti-CM-01 antibody show that there are many cytocapsular tubes (CTs, white arrow, in curved tube-shaped morphologies with 3-6 µm in diameter/width, in red-framed areas) in clinical breast cancer tissues, which are invisible in the panels 1-4. These 5 specimen analyzed here are 5 continuously-sectioned specimens (5μm in thickness, panels 1-5) from the same site of the same clinical breast tumor sample. **B:** Representative images of H&E staining (panel 1) and IHC staining with antibodies recognizing DNA mismatch repair protein MSH-2 (panel 2), and anti-CM-01 antibody (panel 3) in clinical colon cancer tissues. The specimen in the above analyses are three continuously-sectioned specimens (5μm in thickness, panels 1-3) from the same site of the same clinical colon cancer sample. There are many cytocapsular tubes (CT, white arrows) in clinical breast and colon cancer tissues, which are invisible in conventional methods of H&E staining and IHC staining assay with conventional cancer markers (ER, PR, HER2 for breast cancer, andMSH-2 for colon cancer), but can be clearly detected by anti-CM-01 antibodies in IHC the fluorescence staining assay. There are many colon cancer cells (in cyan color) in migration in CTs in the colon cancer tissue. Scale bar:10μm.

**Figure S4.**
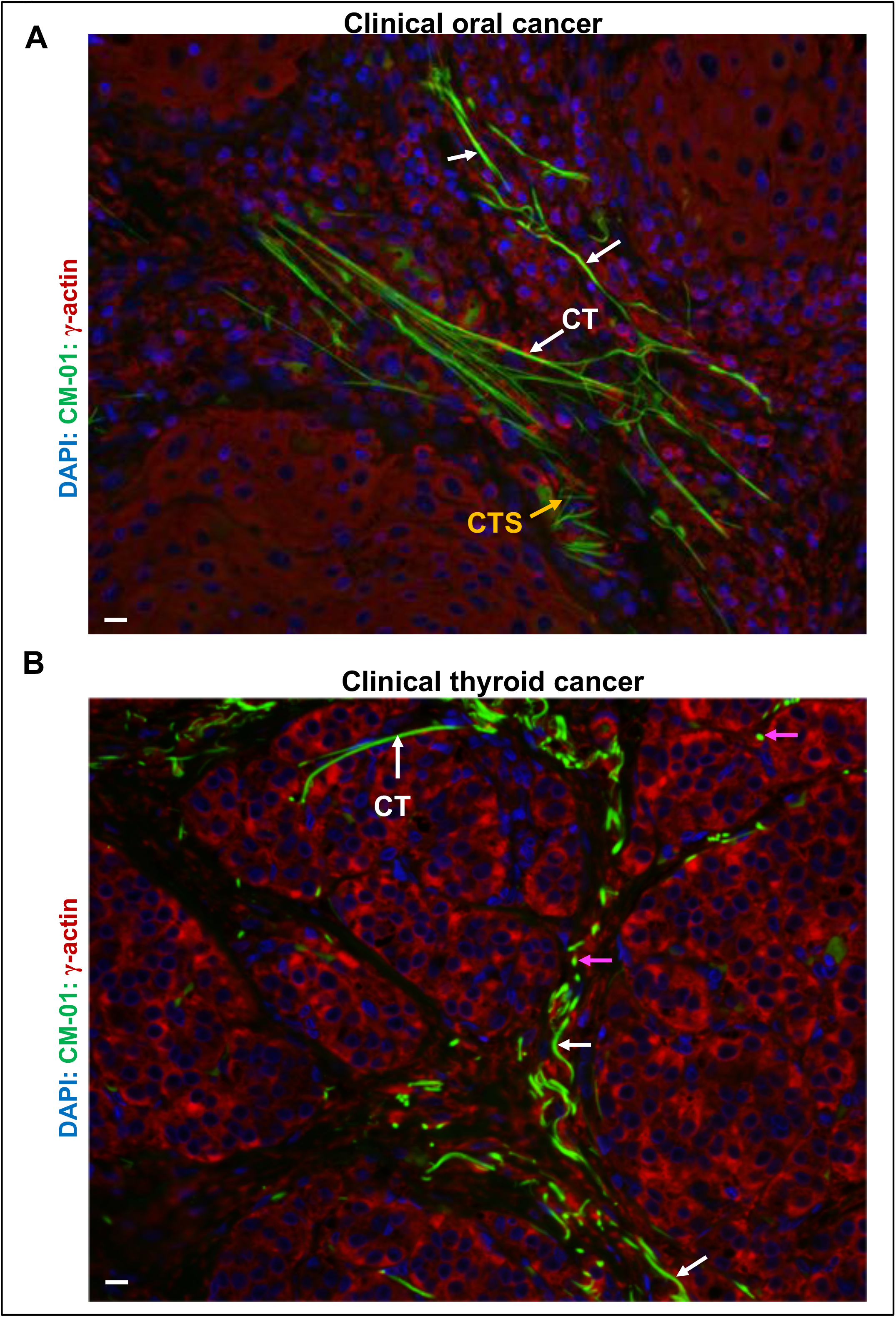
Individual cytocapsular tubes (CTs) potently invade into and pass through compact tissues. **A:** Representative image of compact clinical oral cancer tissue. Individual oral cancer cells are propertied with potent capacities to generate long and straight CT freeways crossing condensed and highly compacted tissues in CT conducting cancer cell metastasis. Longitudinal sectioned CT (white arrows) and CT strands (CTS, orange arrows) are shown. **B**: Representative image of thyroid cancer. Individual thyroid cancer cells exhibit potent abilities to generate CT freeways invading into dense and compacted tissues for cancer cell metastasis. Longitudinal-sectioned or not-sectioned curved CT (white arrows) and cross-sectioned CT (purple arrows) are shown. The specific binding ability of anti-CM-01 antibodies to the CT protein is shown. Scale bar:10μm.

**Figure S5.**
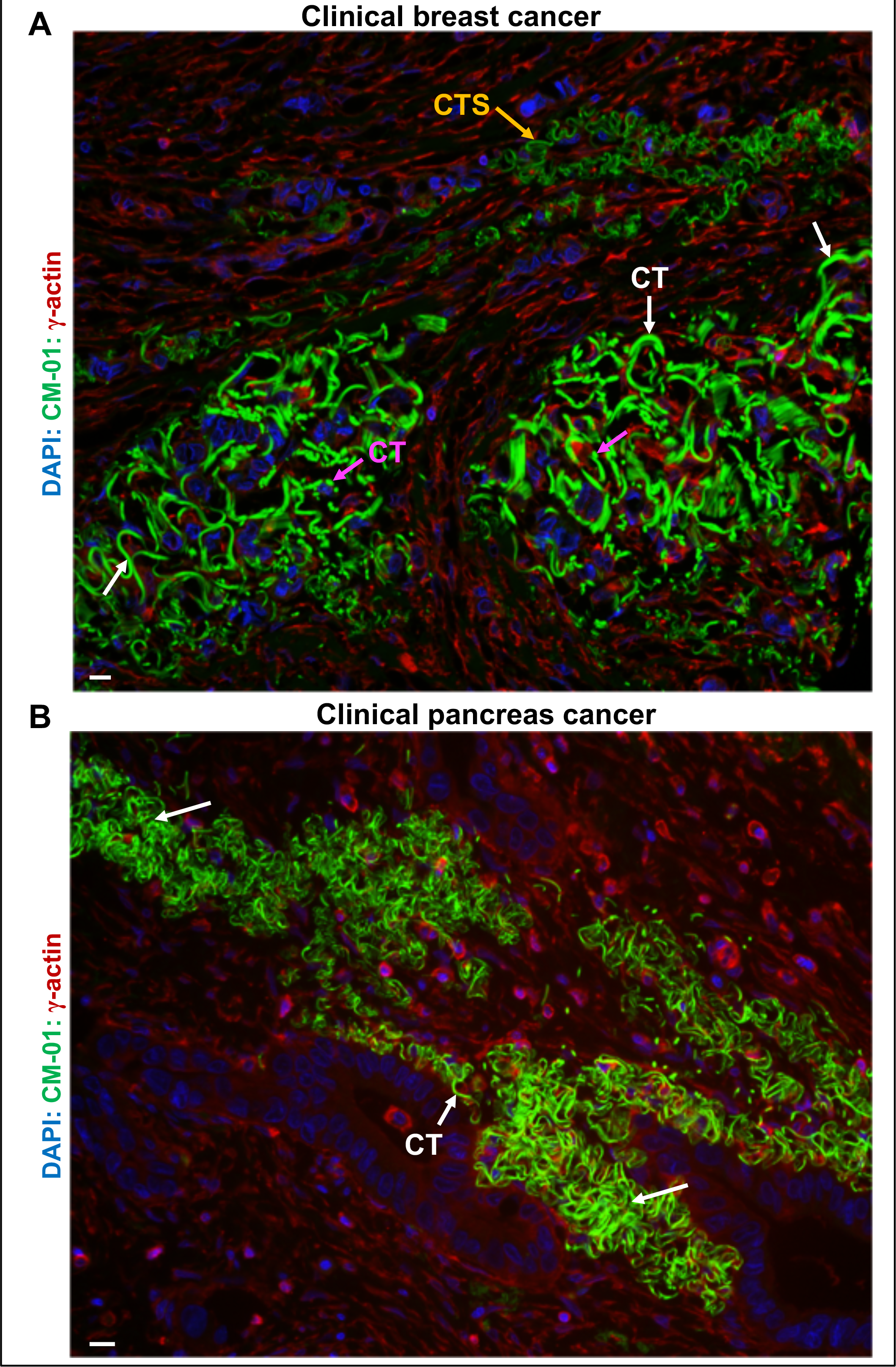
Collective cancer cell cytocapsular tubes (CTs) powerfully invade into compact tissues. **A:** Representative image of collective and curved breast cancer CTs invading into compacted tissues. Breast cancer cells present powerful abilities in engendering collective curved CTs for massive cancer metastasis in compacted tissues. Curved, longitudinal-sectioned or not-sectioned CT (white arrows), cross-sectioned CT (purple arrows), and CT strands (CTS, orange arrows) are shown. **B:** Representative image of cytocapsular tubes (CTs, white arrows) in clinical pancreas cancer tissues. Massive curved CTs invade into compact tissues. Scale bar:10μm.

**Figure S6.**
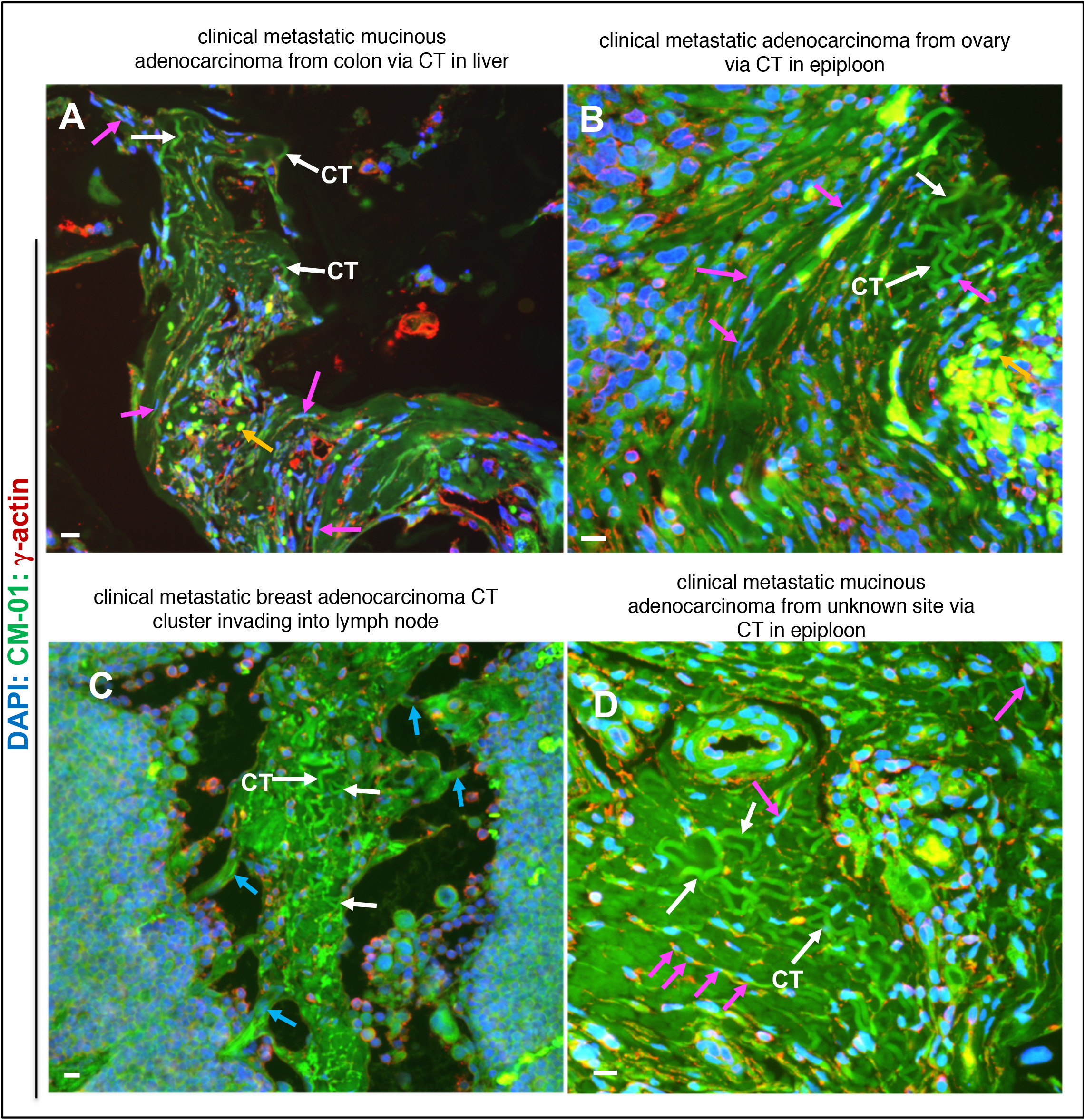
Long distance cancer metastasis crossing organ/tissue via cytocapsular tubes (CTs) and CT networks. **A:** Representative image of metastatic mucinous adenocarcinoma from colon via colon cancer cell CT in liver. Many CTs align and entangle together and form compact CT bundles with multiple sharp tips (top of the panel) invading into the liver with many cancer cells in migration inside colon cancer-cell CTs. Large quantities of liver cells surrounding the colon cancer cell CT cluster are apoptotic and disappear, leaving a big area absent of liver tissues. Longitudinally sectioned colon CTs (white arrows) with longitudinally-sectioned cell in migration in colon CTs (purple arrows), and cross-sectioned colon CTs (orange arrows) in liver are shown. **B:** Representative image of metastatic adenocarcinoma from ovary via ovary cancer cell CTs in epiploon. There are many straight and curved ovary cancer cell CTs entangled together, invading into compact epiploon tissues, with many ovary cancer cells migrating in CTs. Longitudinally sectioned ovary CTs (white arrows) with longitudinally sectioned cell in migration in CTs (purple arrows), and cross-sectioned ovary CTs (orange arrows) in epiploon are shown. **C:** Representative image of metastatic breast adenocarcinoma from breast via breast cancer cell CTs (and CT networks) in lymph node. Many curved breast cancer cell CTs entangled together and interconnected to form a condensed CT cluster powerfully invading into compact lymph node. Several small CT bunches (cyan arrows) invade into lymph node at the right and left sides. **D:** Representative image of mucinous adenocarcinoma from unknown site via CT in epiploon. Longitudinally sectioned ovary CTs (white arrows) with longitudinally sectioned cells migrating in CTs (purple arrows) are shown in epiploon. Scale bar:10μm.

**Figure S7.**
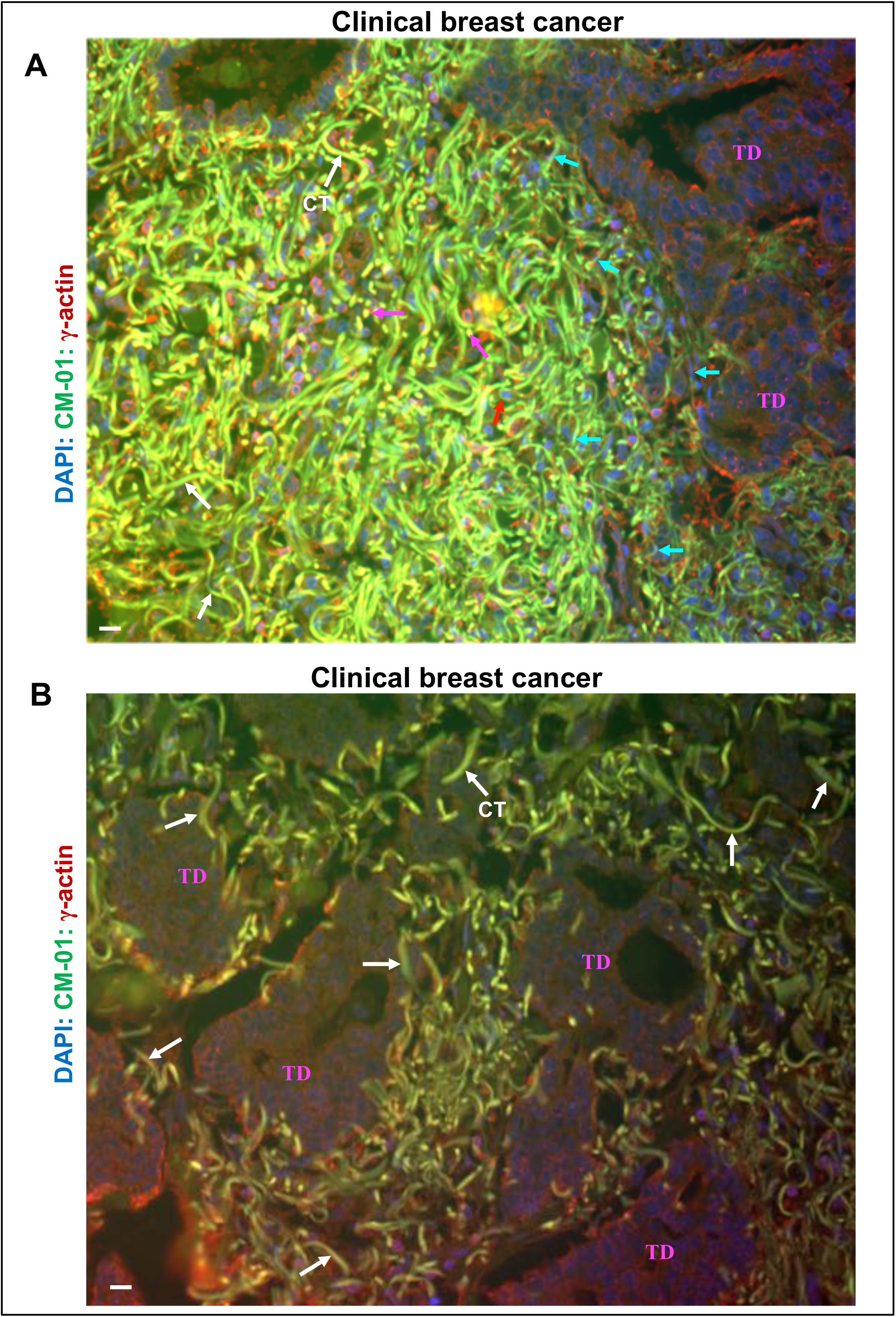
Initiation and early elongation of cytocapsular tubes in clinical breast carcinoma in situ. **A:** Initiation and early elongation of cytocapsular tube in clinical breast ductal carcinoma in situ. There are plenty of single breast cancer cell cytocapsular tube (CT) in curved morphologies showing shrunk thin tails without CT degradation. The longitudinally sectioned (or not sectioned) CT fragments (white arrows), longitudinal sectioned CT fragments carrying cancer cells (in cyan or blue color and in long and thin morphology) are shown in migration in CTs (cyan arrows), cross sectioned cytocapsula with cell inside (red arrow), and cross sectioned CTs (purple arrows). **B:** Elongation of CTs in clinical breast ductal carcinoma in situ. Breast cancer cell CTs (white arrows) elongate and aggressively invade into terminal ducts and neighboring tissues. The elongated CTs show consistent diameter/width without the shrunk thin tails. Terminal ducts (TDs) are shown. Scale bar:10μm.

**Figure S8.**
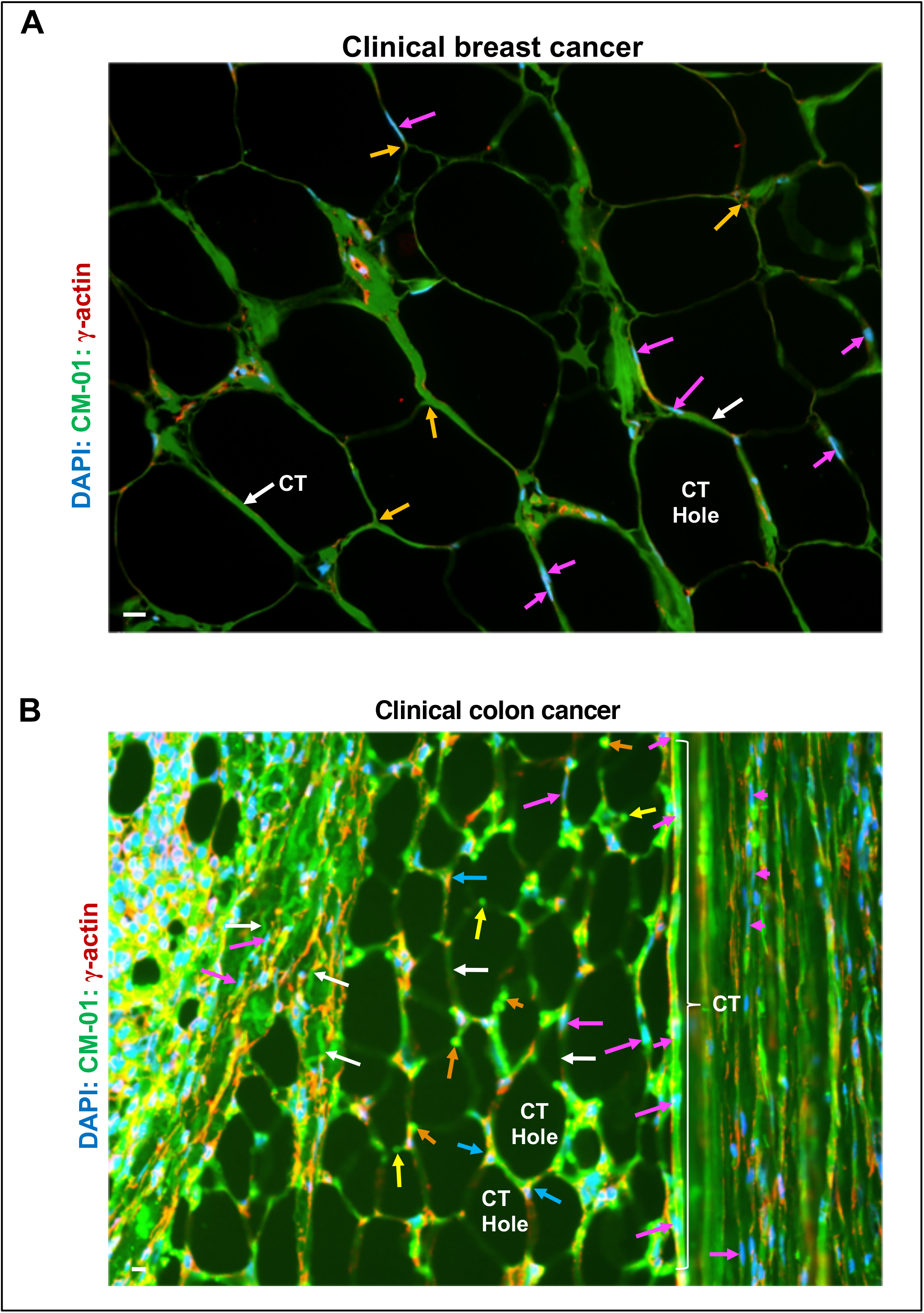
Cytocapsular tubes (CTs) interconnect and form large 3D CT networks for cancer cell dissemination. **A:** Representative image of a part of sectioned network of large and 3D breast cancer cell CT networks with breast cancer cells migrating in CT networks. Cells migrating in CTs can generate new CTs and CT branches in diverse directions. CTs (white arrows) interconnect to each other and form large, 3D and self-organized networks. There are many breast cancer cells (purple arrows) freely migrating in the CT networks, in which most cells are in long and thin morphologies squeezed and shaped by the CT membranes. CT connection node (orange arrows), and longitudinal sectioned CT fragment with longitudinal sectioned cells in CTs (purple arrows) are shown. **B**: Representative image of a part of sectioned 3D colon cancer cell CT networks with colon cancer cells migrating in CT networks. CT (white arrows), CT connection node (cyan arrows), longitudinal sectioned CT fragment with longitudinal sectioned cells in CTs (purple arrows), cross sectioned CT surfaces with exposed round CT membranes with the tiny CT lumen in a dimmed/darkness in the center (yellow arrows), cross sectioned CT surfaces with shrunk CT membrane and without dimmed lumens (orange arrows) are shown. Multiple (3 or more) CTs form a CT hole/grid. In the right part, there is a compact, dense and wide CT network bundle with many long and straight CTs (white brace) align together and interconnect to each other. In a long and straight CT fragment (white brace), there are 5 colon cancer cells (purple arrows) that migrate inside. In the left part, there is a compact and thin CT network bundle with many curved CTs (white thin arrows) entangled together, with many colon cancer cells (purple arrows) in migration in CTs. There is no CT degradation. The grids or holes among CTs in the 3D CT networks are formed during CT interconnection to generate 3D CT networks. Scale bar, 10μm.

**Figure S9.**
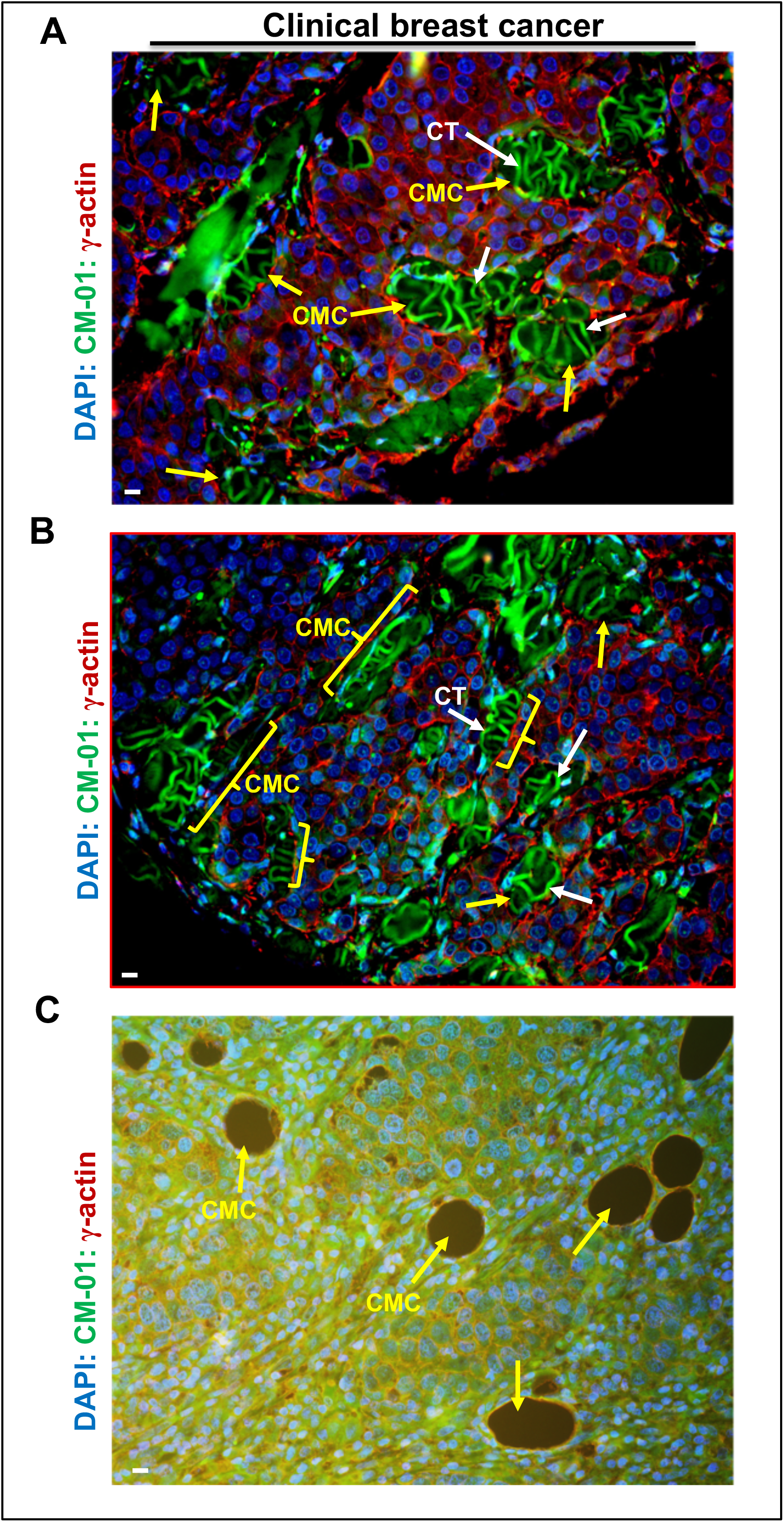
Development of cytocapsular tube (CT) mass cavities. **A-B:** Highly curved/coiled breast cancer cell CTs (white arrows) entangle together and engender CT masses. CT masses occupy spaces in compact breast cancer tissues and form CT mass cavities (CMCs, yellow arrows). Representative image of cross-sectioned CMCs (**A**) and longitudinally-sectioned CMCs (**B**, yellow braces) in compact breast cancer tissues are shown. These sectioned CMCs interconnect and form CMC clusters or networks distributed in tissues. Cancer cells surrounding the CT mass cavity can invade into the CTs and leave, and the cavity space size will increase, and the cavities grow into bigger ones. **C**: Representative image of empty CMCs after CT degradation and decomposition. After CT degradation, CTs disappear, leaving empty CMCs in the compacted cancer tissues. Scale bar:10μm.

**Figure S10.**
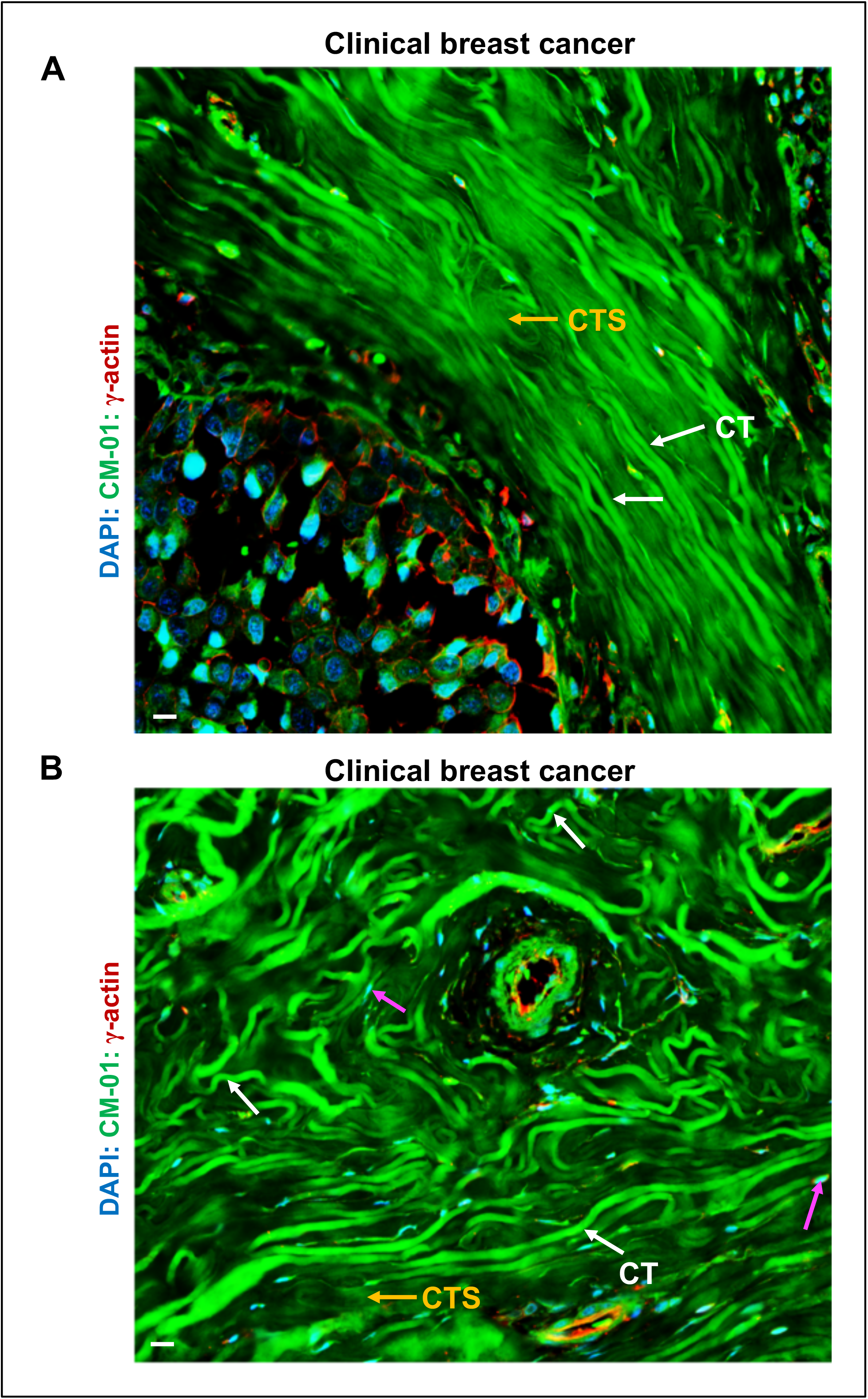
Collective cytocapsular tubes (CTs) align together and form CT bundles for massive cancer cell dissemination. **A:** Representative image of breast cancer presents many long and straight breast cancer cell CTs aligning together and form compact CT bundles that conduct cancer metastasis to far distance tissues and organs. **B:** Representative image of breast cancer displays that massive highly curved breast CTs entangled together occupying the space. The cancer cells in that place have invaded into these CTs and relocate to other sites via the CTs, leaving no breast cancer cell here. CT (white arrows), and cell in migration in CTs (purple arrows) are shown. Scale bar:10μm.

**Figure S11.**
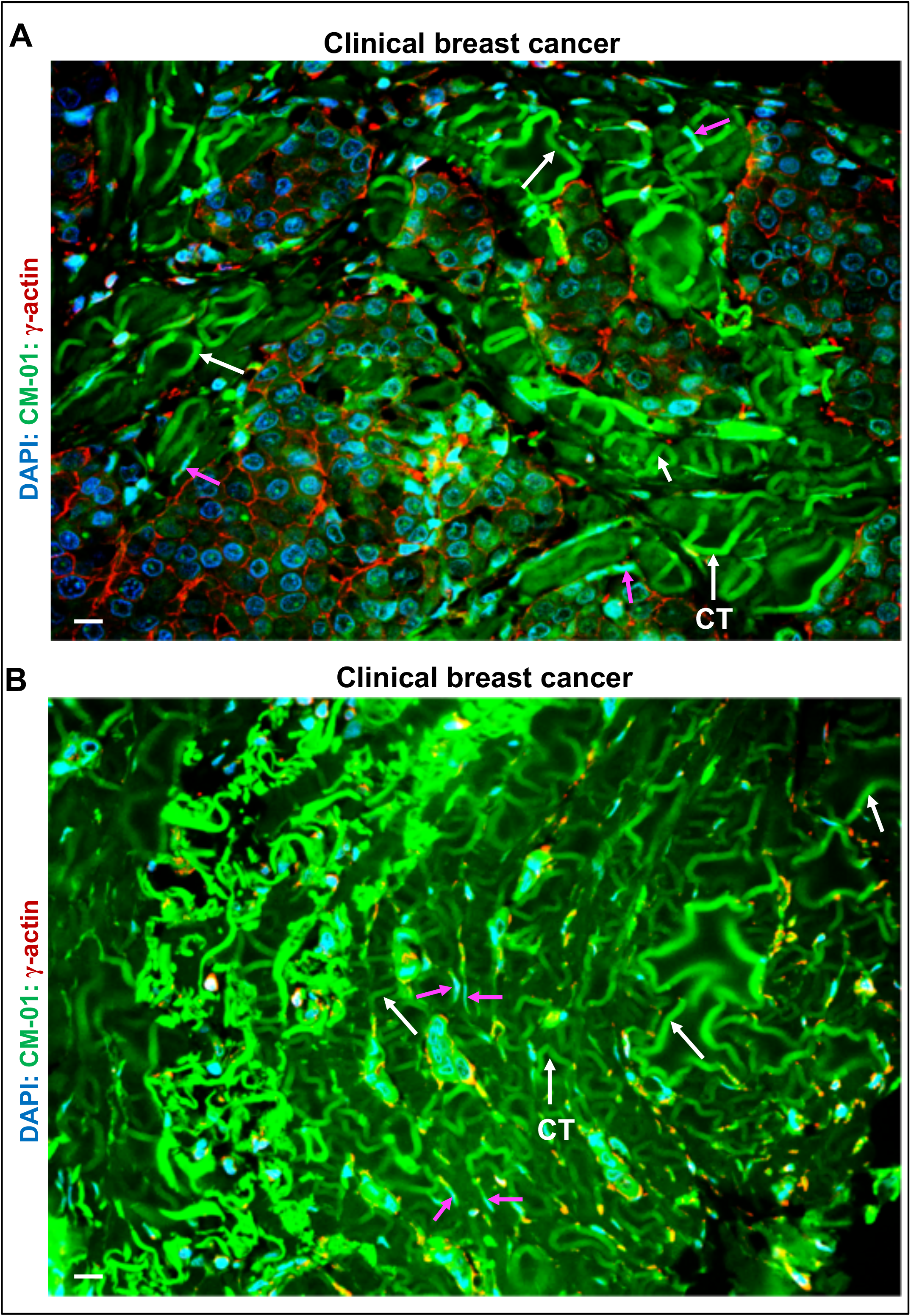
Curved cytocapsular tubes (CTs) and CT clusters invade into compact tissues in diverse pathways conducting cancer metastasis in multiple directions. **A:** Representative image of breast cancer tissues. Highly curved and coiled breast cancer cell CTs potently invade into and pass through compacted tissues in multiple and diverse pathways, and conduct cancer metastasis in multiple directions. The cancer cells outside of and close to the CTs can invade into the CTs and relocate to other sites via CTs. **B:** A representative image presents that numerous highly curved/coiled breast CT masses conquer the place. The cancer cells in the place have invaded into these curved CTs and relocate to other sites via the CTs, leaving no breast cancer cells. Even with the highly curved morphology, these CTs (without degradation) keep the similar and consistent sizes in diameter/width. CT (white arrows), and cell in migration in CTs (purple arrows) are shown. Scale bar:10μm.

**Figure S12.**
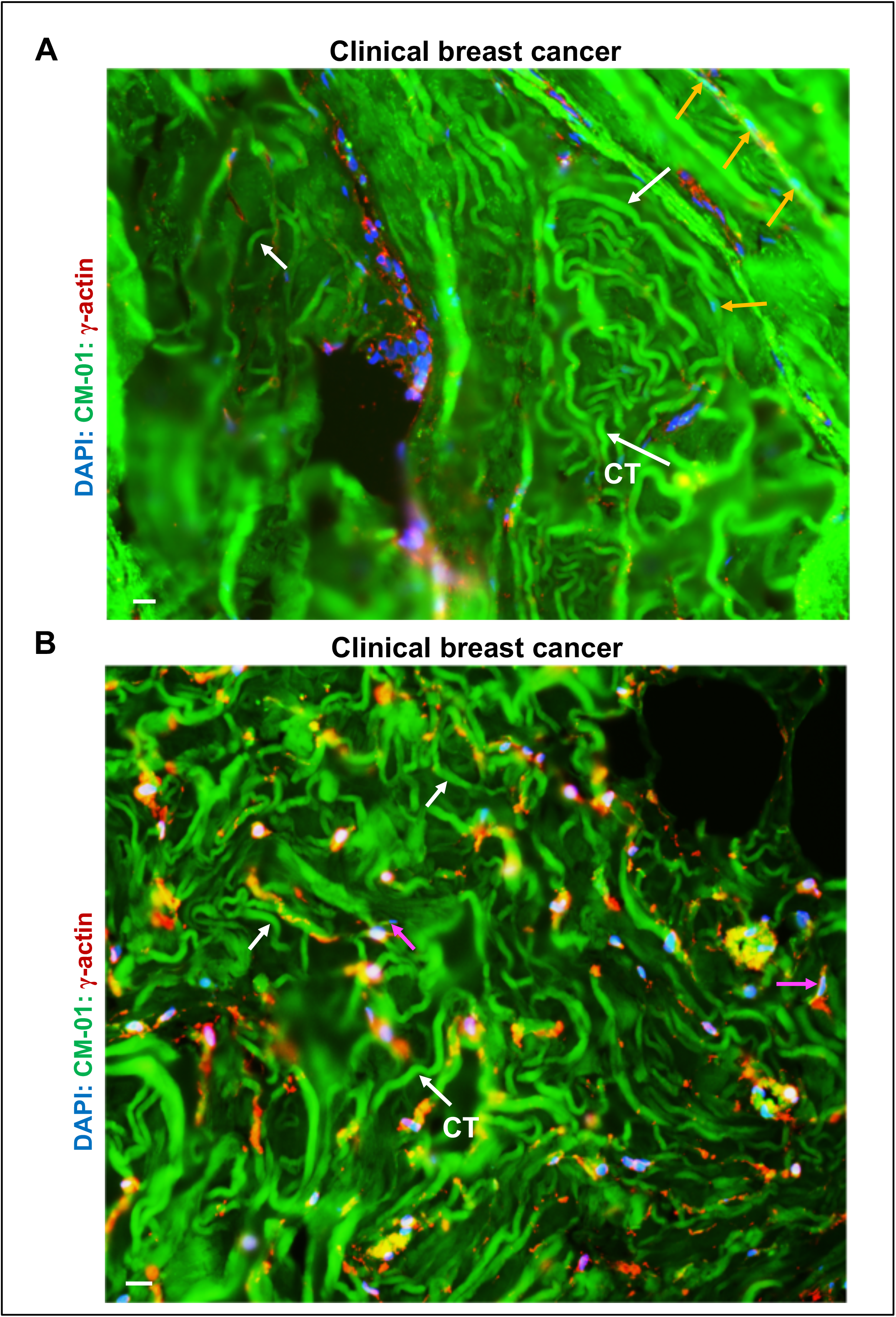
Massive breast cancer cell cytocapsular tubes (CTs) conduct cancer metastasis. **A**: Representative image of clinical breast cancer tissues with CT masses. There are many highly curved CTs that pile up and form compact CT masses. The major breast cancer cells in place invade into the CTs and relocate away, leaving very few cells in the center. **B**: Representative image of breast cancer tissues with CT masses. A large quantity of curved CTs pile up and form large CT masses occupying the spaces. The breast cancer cells in this place invade into these CTs and relocate away leaving no cancer cells outside of CTs. At the top right corner, there are two big cavities caused by the CT degradation and decomposition. CT (white arrows), and cell in migration in CTs (purple arrows) are shown. Scale bar:10μm.

**Figure S13.**
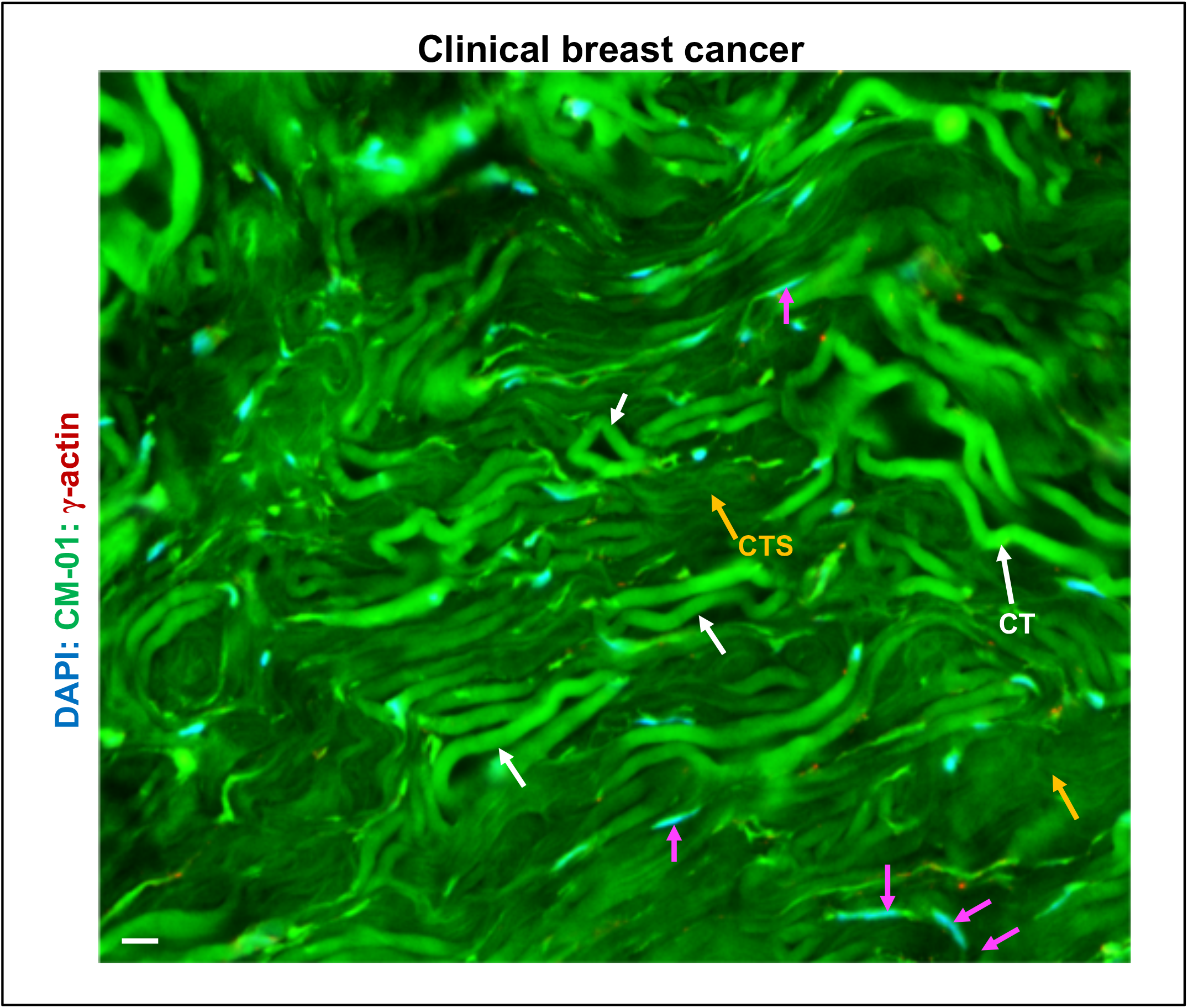
Massive breast cancer cell CTs conduct massive cancer metastasis. There are plenty of hairpin-shaped and highly curved CTs (white arrows) in clinical breast carcinoma with cancer cells (purple arrows) migrating in CTs. Massive cancer cells invaded the CTs leaving no cancer cell outside. Some CTs degrade into CT strands (CTS, orange arrows). Scale bar, 10μm.

**Figure S14.**
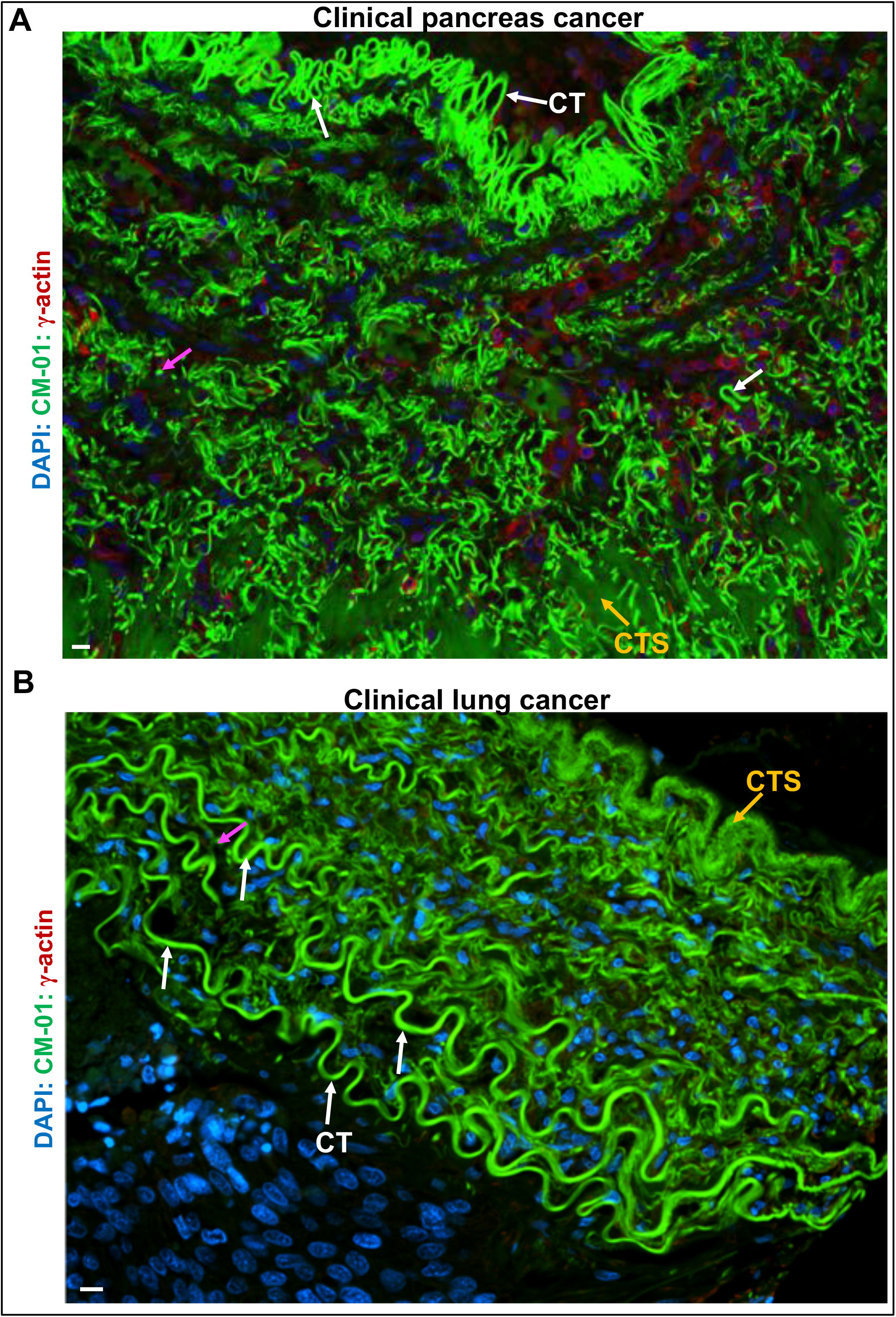
Highly curved pancreas and lung cancer cell CTs induce cancer metastasis. **A:** Representative image of pancreas cancer with plenty of highly curved pancreas cancer cell CTs randomly distributed in clinical pancreas cancer tissues. At the top, there is a long and highly curved pancreas cancer cell CT with many condensed hairpin-shaped turns and a continuously maintained diameter of 4 μm. **B**: Representative image of lung cancer with multi-layered, highly curved lung cancer cell CTs. At the bottom, there are multiple layers of CT individuals with regular and irregular turns and curves. The highly curved CT repeats significantly increase the area for contacting more cancer cells, providing more close freeways for cancer cell metastasis in an efficient format. Scale bar, 10μm.

**Figure S15.**
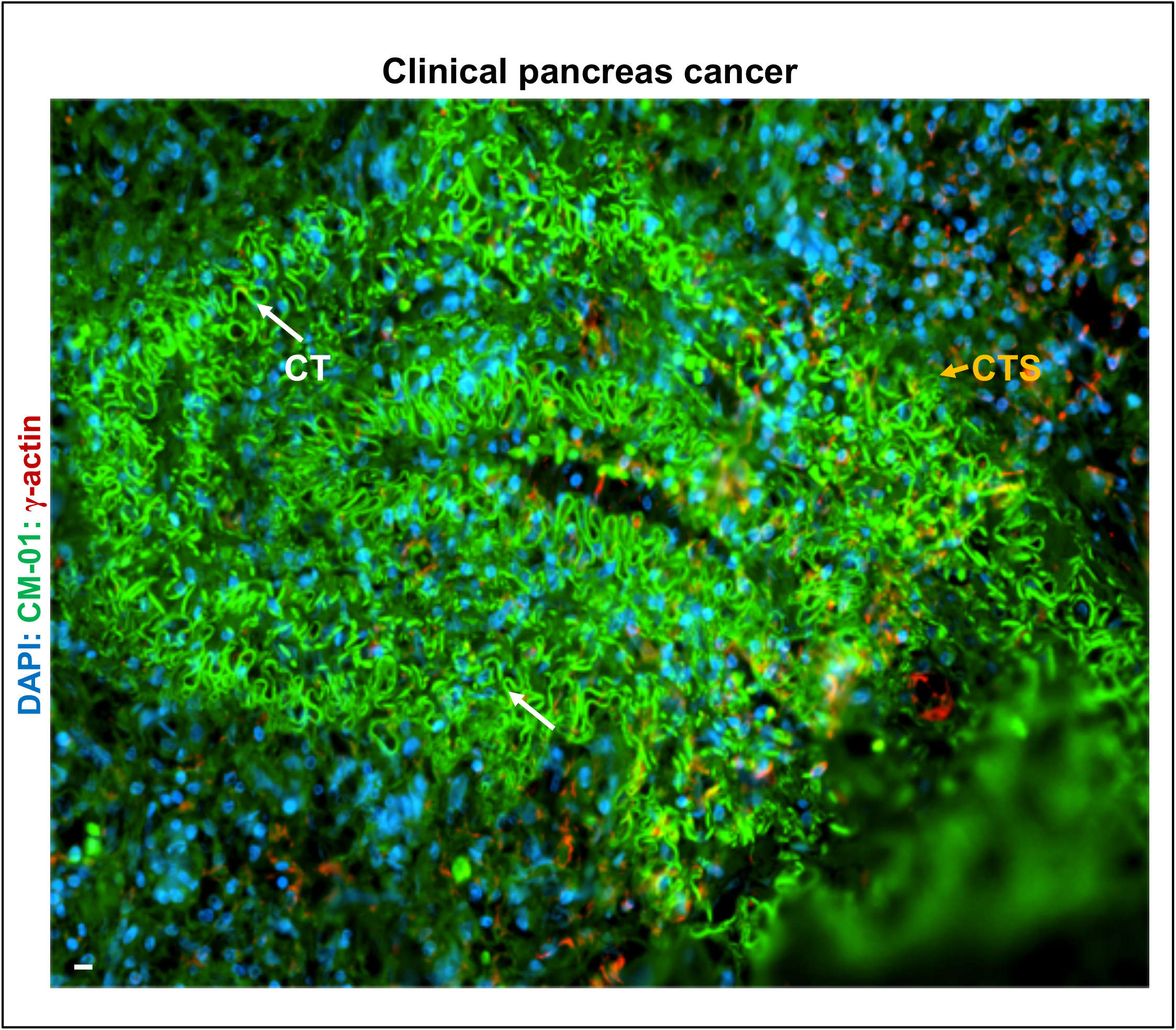
Cross-section view of highly curved, coiled, multi-layered pancreatic-cancer CT super-structures. The highly curved and coiled pancreatic cancer cell CTs form multi-layered CT superstructures in pancreas cancer, displaying the powerful capacities of CTs in generating complex super-structures when invading into tissues during the CT conducted cancer metastasis. CT (white arrows), and CT strands (CTS, orange arrows) are shown. Scale bar:10μm.

**Figure S16.**
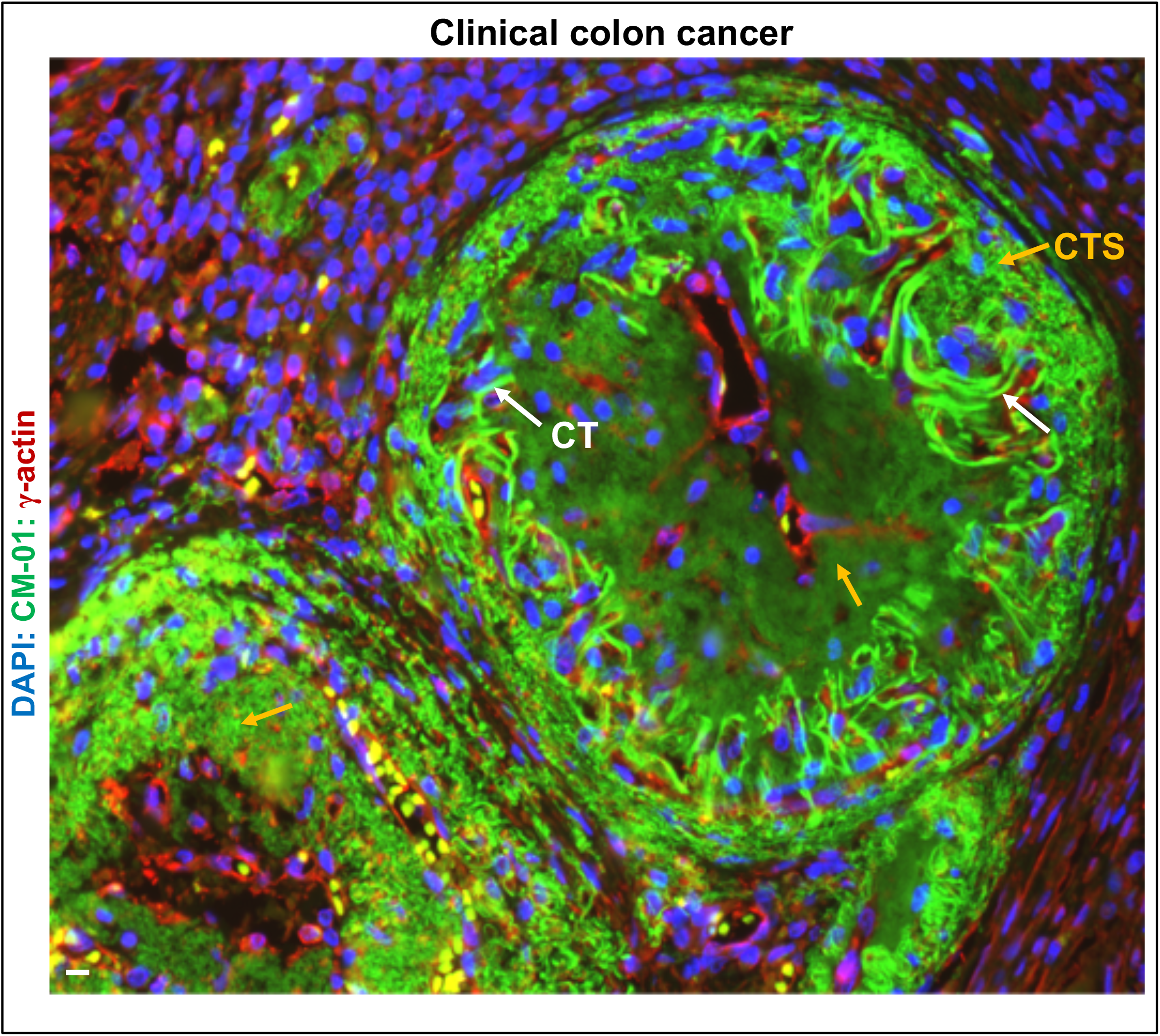
Cross-section view of two big, round, multi-layered, multi-staged, complex colon cancer CT superstructures. A representative cross-section view of two big and complex CT super-structures of round morphology. The bigger one displays multiple layers: outer side thin layer, in severe degradation into thin CT strands; middle thick layer, many highly curved CTs distributed in a ring circulating the center; and inner side layer, CTs with severe degradation into cloud-like status. At the bottom left corner, most CTs degrade into CT strands (CTS) in diverse thickness in this big complex CT super-structure. These two complex CT super-structures of round morphology and exhibiting multiple layers of multi-staged CTs display the vigorous abilities of collective CTs in engendering diverse and effective superstructures for efficient invasion during conducting cancer metastasis. CT (white arrows), and CT strands (CTS, orange arrows) are shown. Scale bar:10μm.

**Figure S17.**
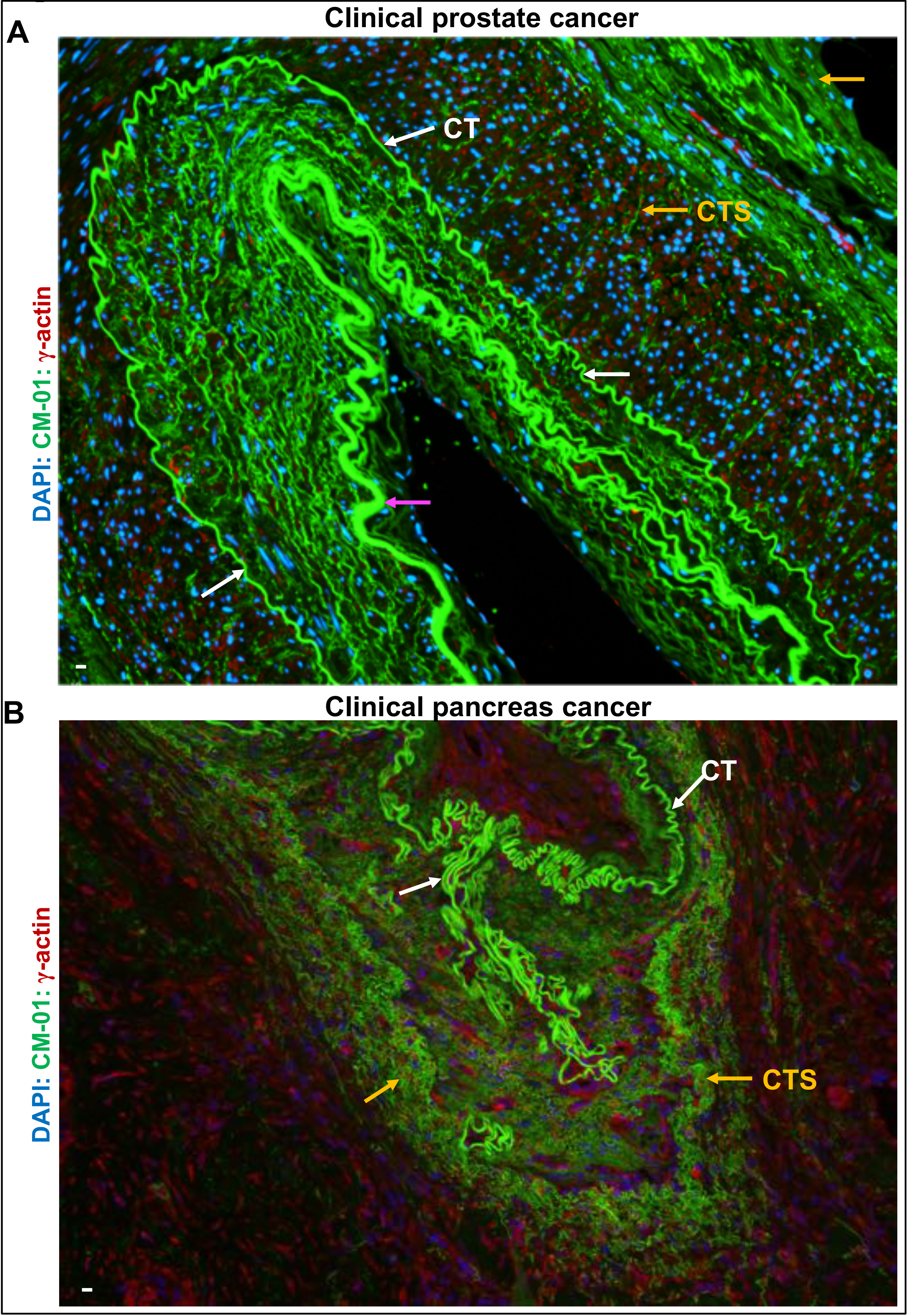
Cross-section view of huge, flattened and oval shaped, multi-layered, complex prostate cancer CT superstructures. **A**: Representative image of cross-section view of a part of a huge, flattened and oval shaped, multi-layered, complex prostate CT superstructure. There are many circulated CTs that form a complex CT superstructure. The purple arrow labeled CT bundle is formed by aligned 4 long and curved CTs with a big hair-pin shaped curve. This huge, multi-layered, complex CT superstructure presents the potent capacities of collective CT invasion in cancer metastasis. Multiple CTs tightly align together and form a CT bundle with enhanced invasion capabilities during conducting cancer cell metastasis in tissues. **B**: Representative image of a part of cross-section view of a huge, multi-layered, irregular-shaped morphology, multi-CT stages, and complex pancreas cancer cell CT superstructures. In the middle, there is a part of a single, long, highly curved pancreas cancer cell CT with many hairpins, turns, coils and irregular curves. In the outside, there are multiple layered CTs degrade into strands in various sizes in width. CT (white arrows), and CT strands (CTS, orange arrows) are shown. Scale bar:10μm.

**Figure S18.**
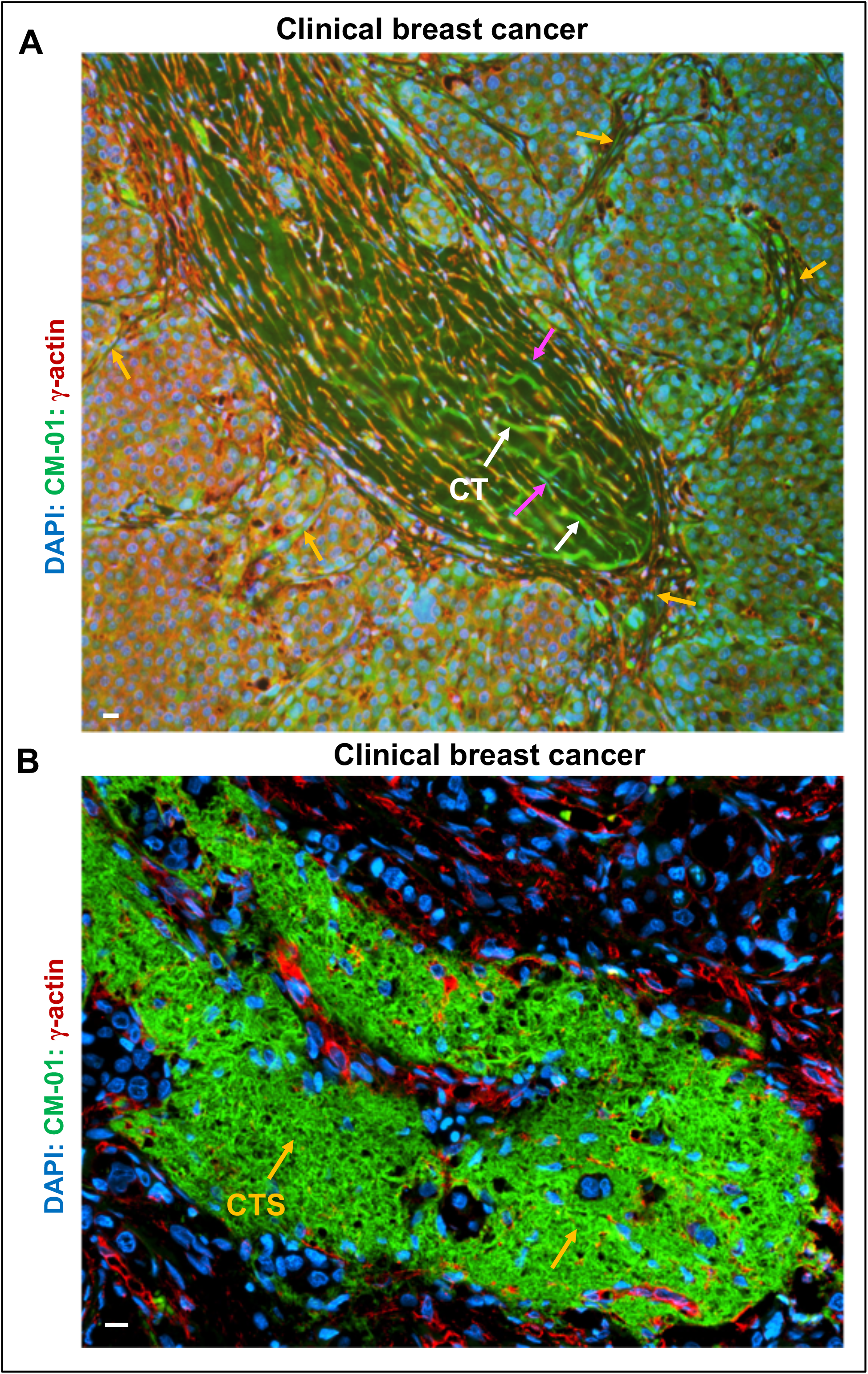
Longitudinal section view of long and big breast cancer CT superstructures in conducting cancer metastasis in compact tissues. **A:** Longitudinal section view of a long, big and needle-shaped breast cancer CT superstructure during cancer metastasis. This image shows that many long, curved or straight breast CTs entangle together, and form a long, big, self-organized and needle-shaped CT superstructure, massively and powerfully spur into the highly condensed and compacted tissues, conducting cancer metastasis. There are multiple CTs (orange arrows) extend into the compacted tissue surrounding the CT super-structure including the front. **B:** Longitudinal view of a long, big, sandwiched, irregular CT superstructure in conducting cancer metastasis. Most CTs are degraded into CT strands (CTS, orange arrows). CT (white arrows), and cells in migration in CTs (purple arrows) are shown. Scale bar:10μm.

**Figure S19.**
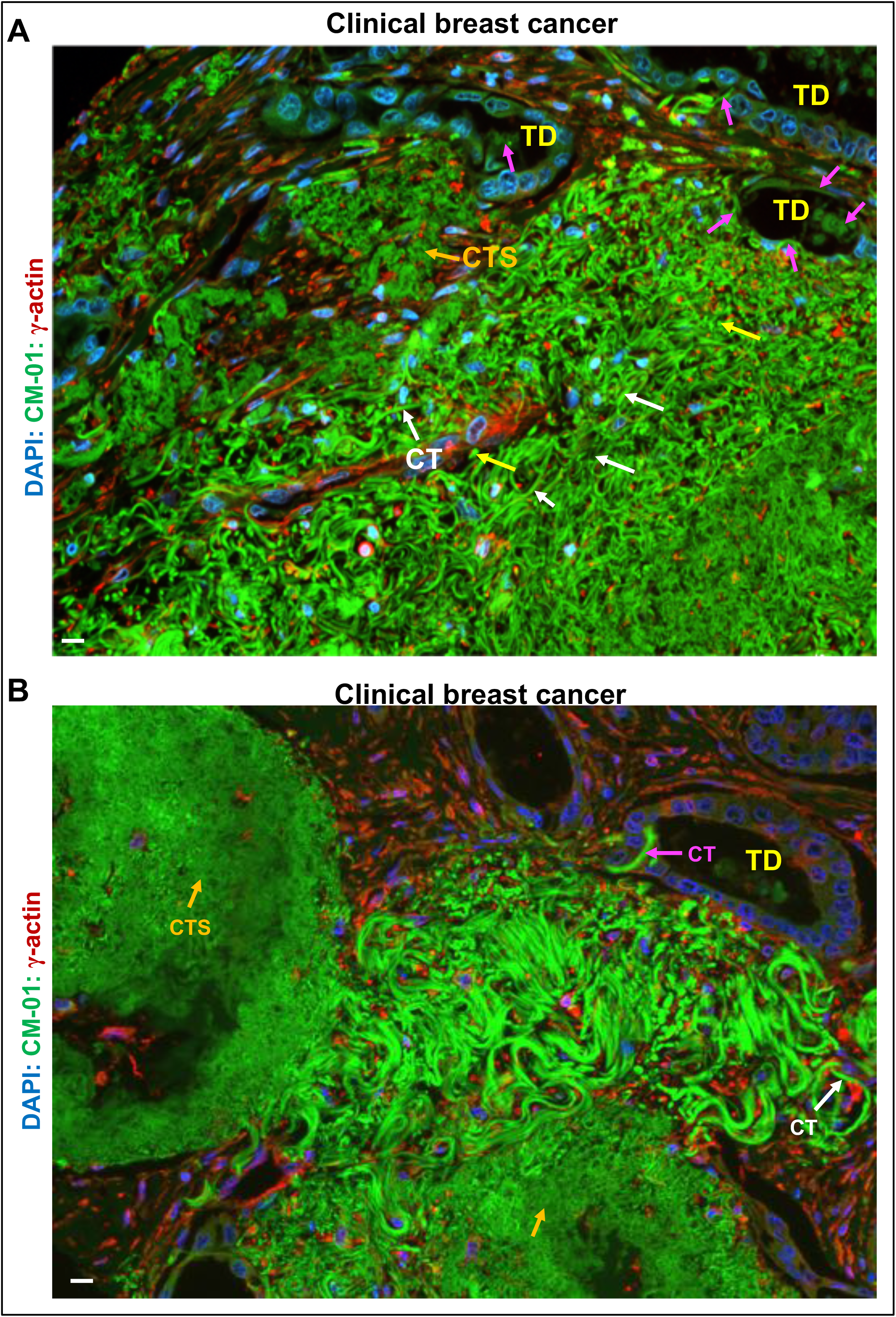
Massive invasion of breast cancer CTs into diverse types of tissues during cancer metastasis. **A:** A representative image of breast cancer tissue shows that a large quantity of breast cancer CTs massively invade into the neighboring tissue among terminal ducts (TD), and some CTs (purple arrows) even crossed through the tight contacted epithelial cell layers of TD and invade into the TD lumens. **B:** A representative image presents that massive CTs in three different CT lifecycle stages invade into diverse tissues during breast cancer metastasis. The right CT mass in a big round morphology is in a late CT stage with CT degradation into cloud-like status. In the middle CT mass, most CTs are not in degradation, and some CTs cross the tight TD epithelial layer and invade into the TD lumens (purple arrow). In the bottom CT mass, most CTs degrade into thick and thin strands. Longitudinal sectioned CT (white arrows), cross-sectioned CTs (yellow arrows), and CT strands (CTS, orange arrows) are shown. Scale bar:10μm.

**Figure S20.**
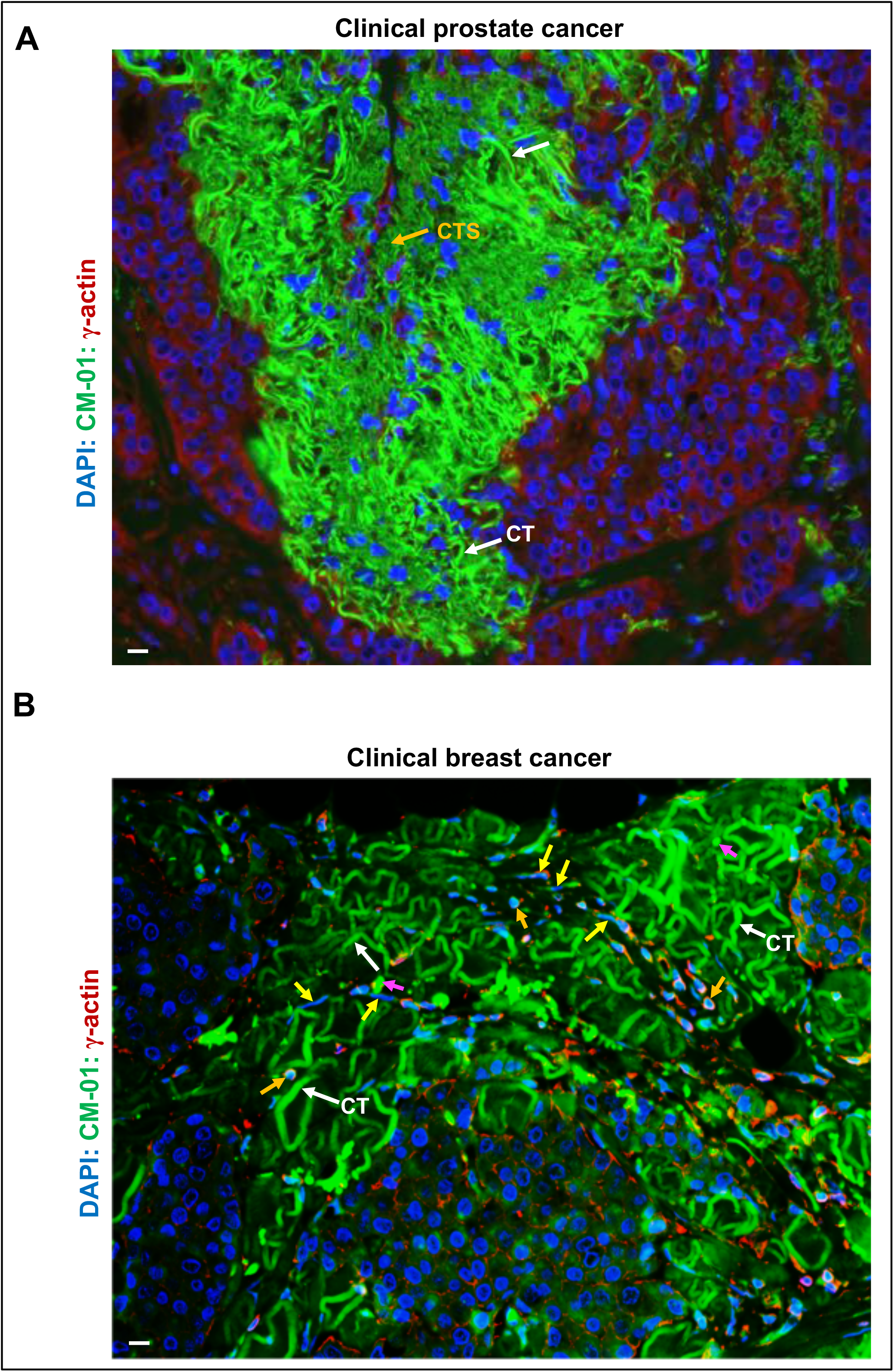
Collective CT clusters invade through compact tissues conducting long distance, CT-directed cancer metastasis. **A.** Representative fluorescence microscope image shows that many curved and entangled prostate cancer-cell CTs formed a curved cone-shaped CT network cluster that invades through a compact tissue layer during cancer metastasis. **B.** Representative image displays numerous curved, coiled and entangled breast cancer cell CT (white arrows) formed collective CT masses that invade into compact tissues in multiple directions, wrapping compact tissues and separating them into several islands. These images document that collective CT clusters can aggressively invade through compact tissues and reach far distance tissues/organs, conducting cancer metastasis in multiple CT-steered directions. Scale bar, 10μm.

**Figure S21.**
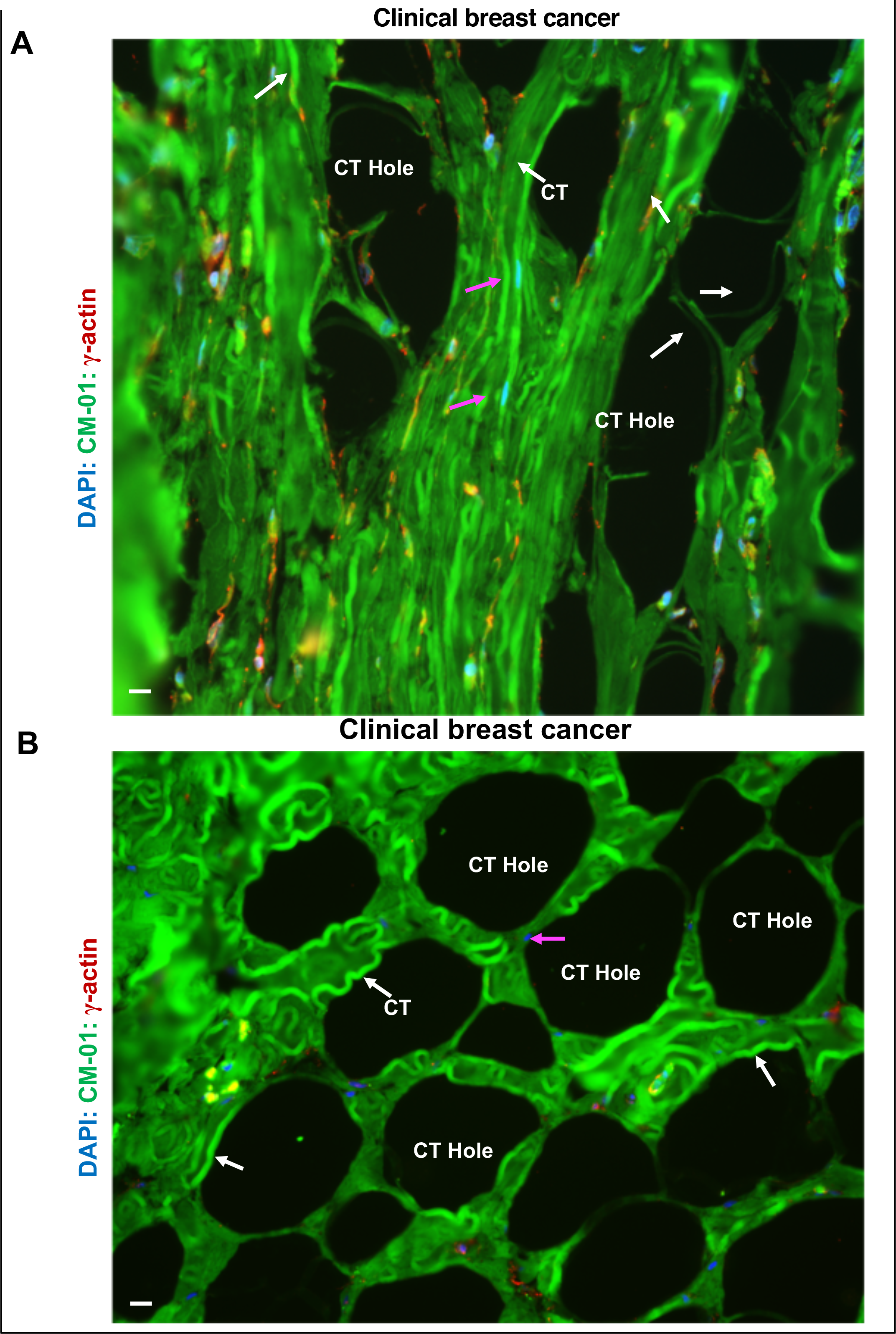
Cytocapsular tube networks possess robust capacities to direct cancer cell metastasis-pathway and are network damage resistant. **A:** Representative image of compacted straight breast cancer CT network bunches with CT degradation. There are several big holes in the compact CT network bundle caused by CT degradation and partly CT network decomposition, but there are still some CTs that connect between the small bundles, which enables the 3D CT network bundles to keep interconnections, thus allowing cancer cell metastasis among the whole CT networks to continue and provide CT network damage resistance. **B:** Representative image of compacted curved CT network masses with severe CT degradation. In the compact and condensed breast cancer CT network masses formed by highly curved, coiled and entangled CTs, part of CTs degrade and networks decompose, leaving many big holes in the CT network masses. The remaining 3D CT networks keep interconnections via curved or coiled CTs, which maintains the powerful cancer cell metastasis pathway capacities with CT network damage resistance. Scale bar:10μm.

**Figure S22.**
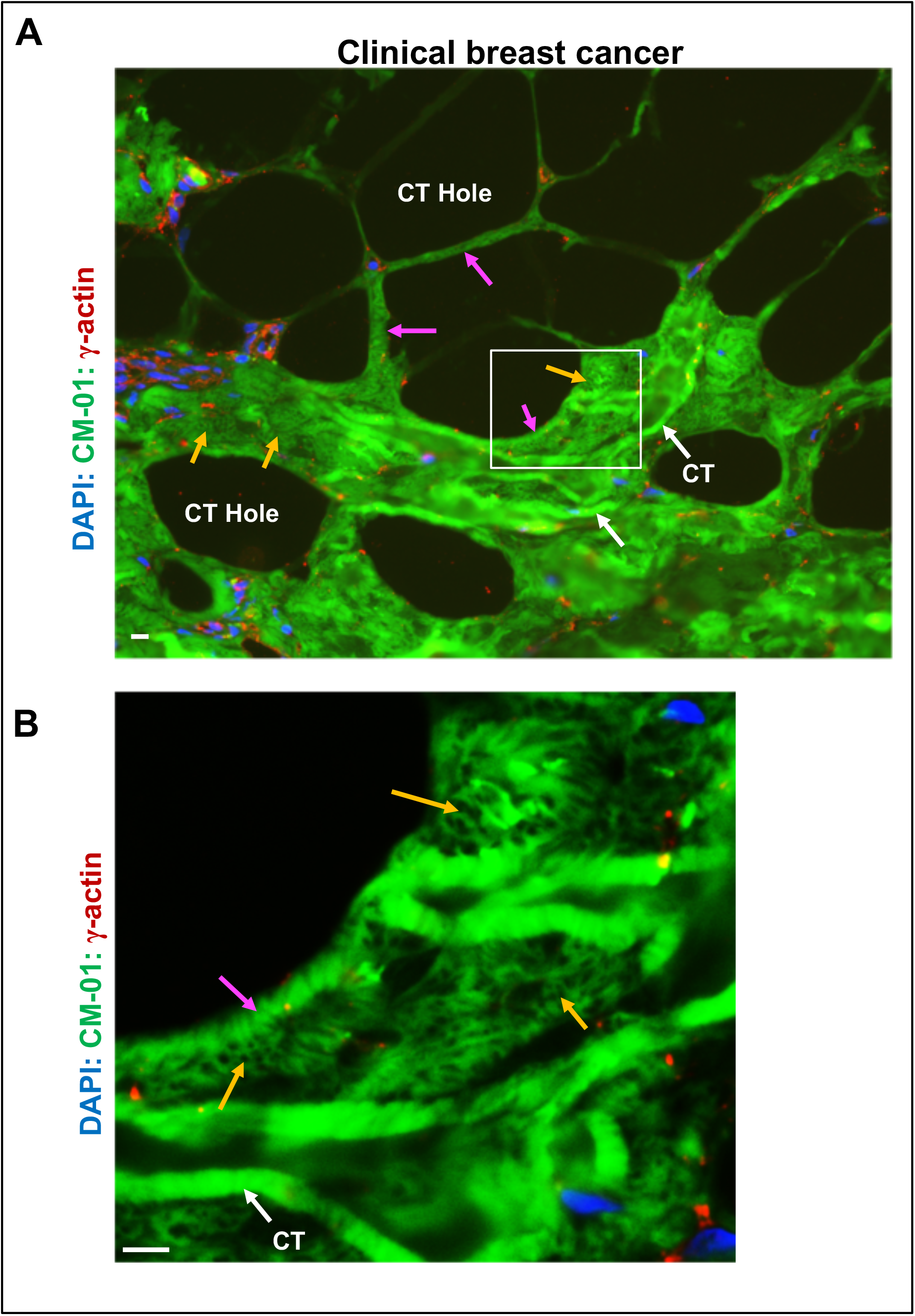
Cytocapsular tube degradation stage I: point array-membrane collapse and decomposition, and CT strand network-tube formation. **A-B:** Representative breast cancer CT degradation from a breast cancer specimen with fresh breast cancer tissues and instant FFPE (formalin-fixed, paraffin-embedded) treatment. There are many tiny points (point array, <0.1μm in width in measured sizes) in the CT membranes collapse and decompose, leaving plenty of tiny holes (0.05-0.2 μm in width in measured sizes) in CTs. The remaining CT strands interconnect and form CT strand-network in the CT tubes: the whole CTs transiently remain the outline of the original CT, while the CT surfaces are composed of thin or very thin CT strand networks with many point-shaped tiny holes (point array) inside. Subsequently, the membrane degradation continues, connected CT strand networks in the CT tubes decompose into disconnected thin strands until they completely disappear. The white-line framed area is enlarged and shown in panel B. This enlargement shows not-yet-degraded CTs (white arrows), micro CT strand network tubes (purple arrow), and tiny point CT membrane degradation-generated holes (orange arrows). Scale bar:10μm.

**Figure S23:**
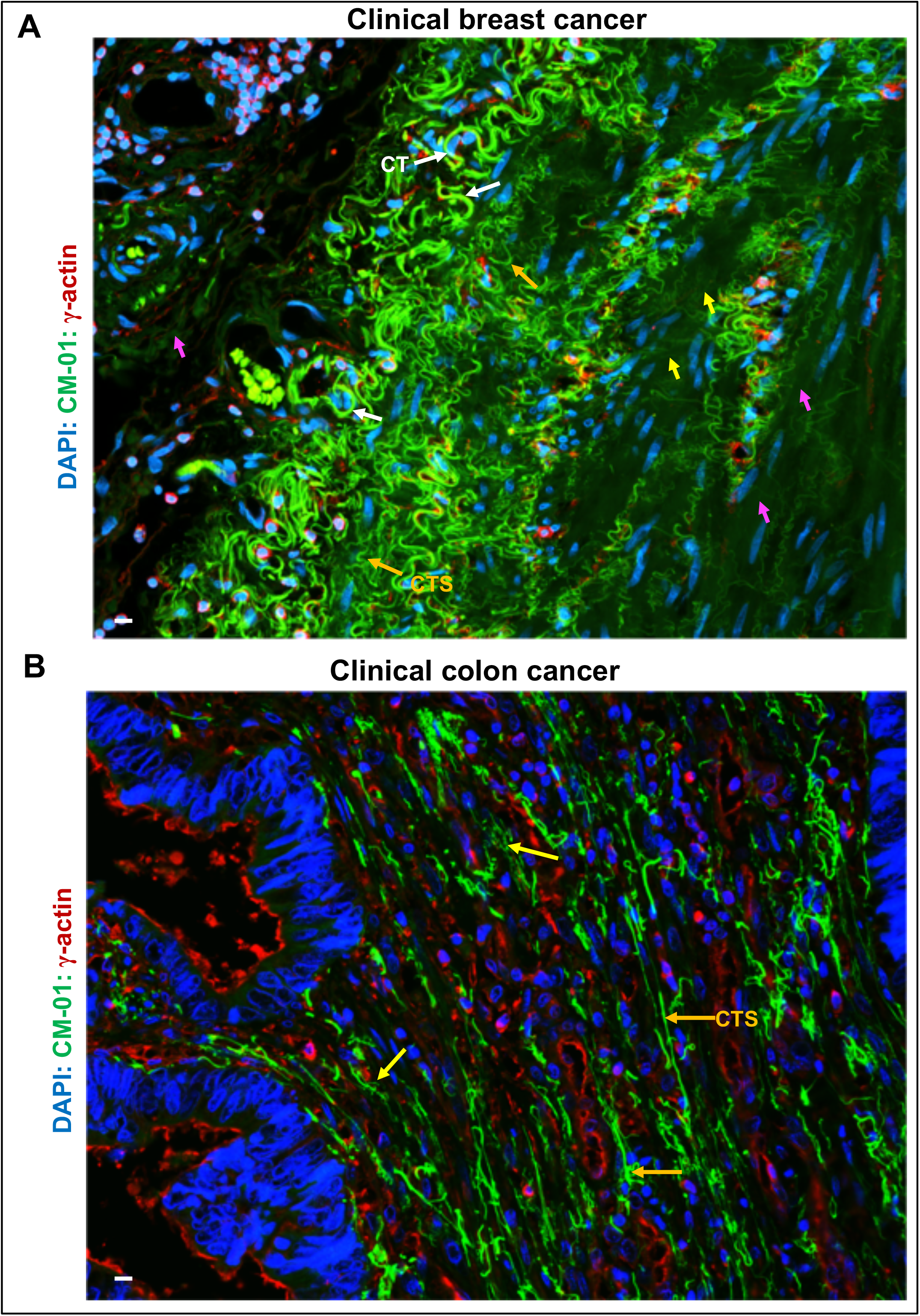
Cytocapsular tube degradation stage II: Disconnected CT strands, thin CT strands. **A:** Representative image of breast cancer showing degradation of micro CT strand network tubes into CT strands, thin CT strands, and cloud like status. Labeled are: CTs without degradation (white arrows), CT strands (orange arrows), thin strands (yellow arrows), and cloud-like status (purple arrows). **B**: Representative image of breast cancer cell CT degradation into CT strands and thin CT strands. Linear colon cancer cell CT strands (orange arrows) and thin strands (yellow arrows) sporadically distributed in colon tissues are shown. Scale bar:10μm.

**Figure S24.**
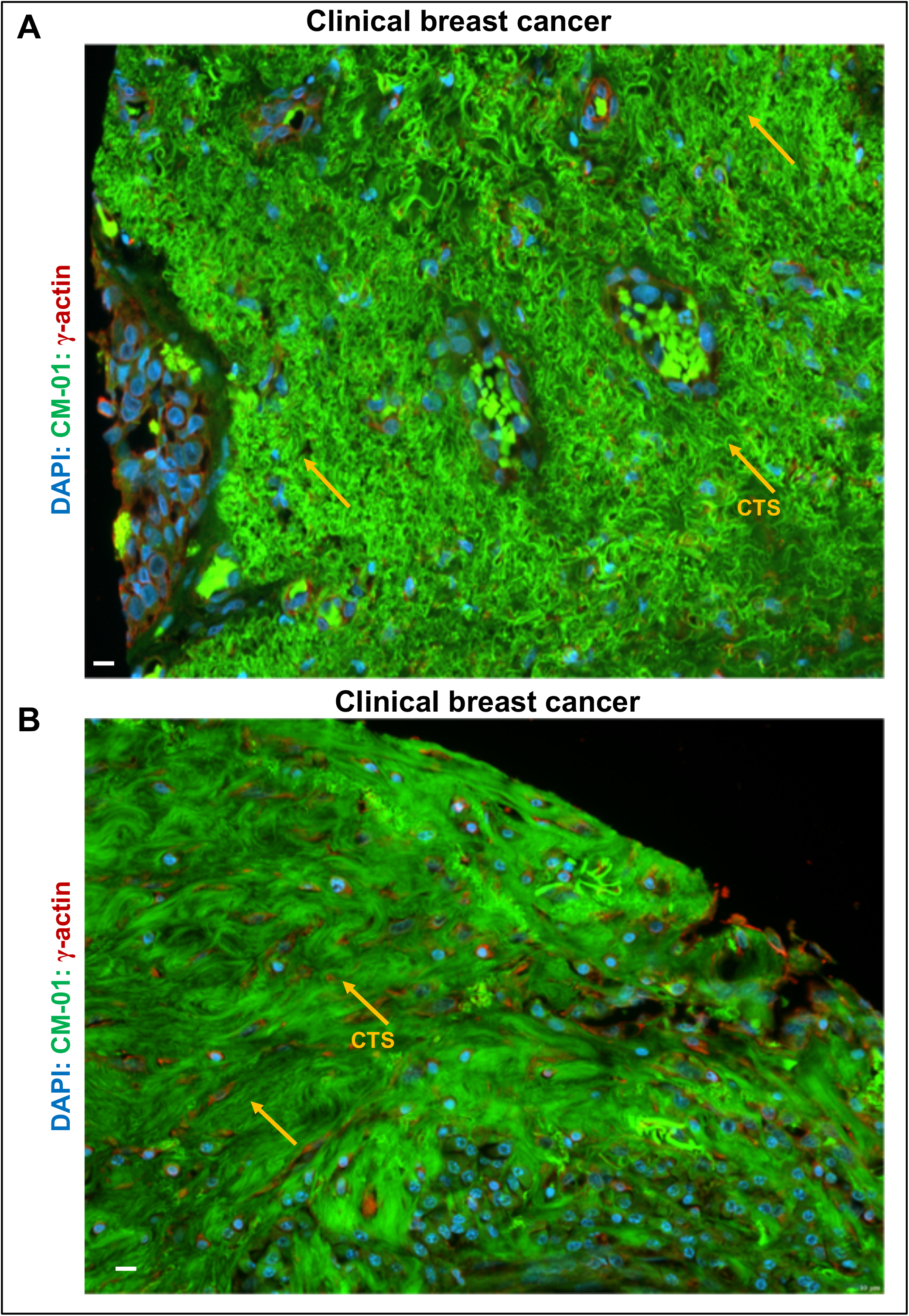
Massive cytocapsular tube degradation. **A**: After cancer cells invade into CTs and migrate away, massive CTs degrade into massive thick and thin strands. There are countless CT strands. **B**: CT strands continue to degrade into massive very thin strands, followed by decomposition and disappearance. After CT strands disappear, these sites will leave cavities filled with intercellular fluids. Scale bar:10μm.

**Figure S25.**
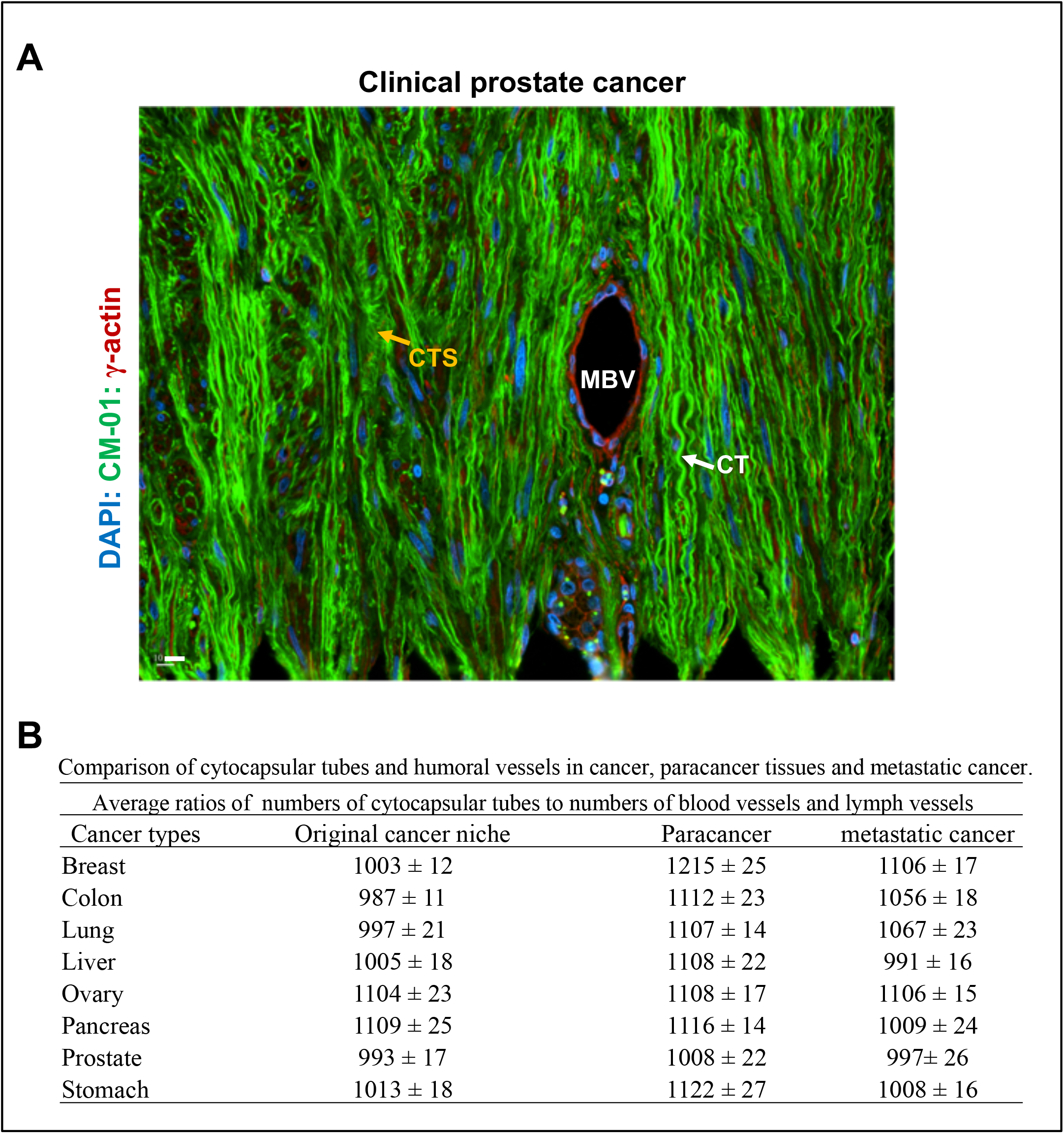
Cytocapsular tubes and networks dominate cancer metastasis pathways. **A:** Representative fluorescence microscope image of prostate cancer tissue specimen shows that there are several hundreds of CTs (white arrows) and only one micro blood vessel (MBV) in this imaged area. Some CTs degrade into CT strands (CTS, orange arrows). **B:** Comprehensive analyses of ratios of numbers of CTs and circulatory system vessels (blood vessels and lymph vessels) examined in cancer, paracancer and metastatic cancer for 8 types of cancers in breast, colon, lung, liver, ovary, pancreas, prostate and stomach. Scale bar, 10μm.

**Figure S26.**
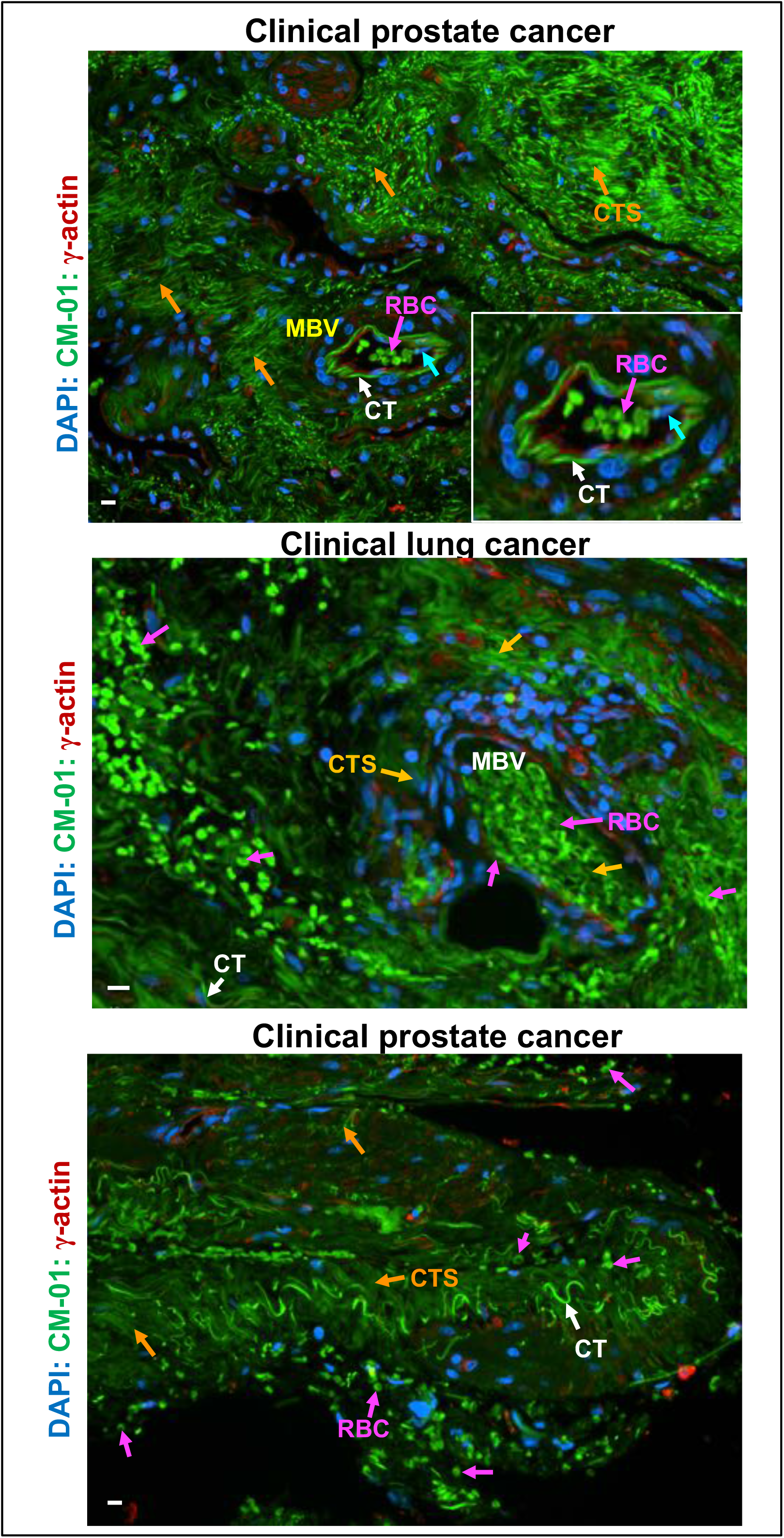
Cytocapsular tubes invade into micro blood vessels and release cancer cells into circulation systems at late cancer stages. **A.** Representative image of rare CT invasion into micro blood vessels (MBV) in stage III prostate cancer. Approximately 1 in 10^4^ CTs (white arrows) wrap around micro blood vessels, aggressively invade into and pass through the tightly connected endothelial layer of micro blood vessel. The cyan arrow labels a CT that protrudes into the micro blood vessel lumen and cancer cells migrate inside. This indicates that cancer cells can be released from the CTs that invade into blood and lymph vessels and subsequently allowing cancer cells release into the circulation systems as circulating tumor cells. There are numerous CTs (white arrow), degraded CT strands (CTS, orange arrows) outside the circulation system vessels. Red blood cells (RBC, purple arrows) are also shown. **B.** Representative image of a micro blood vessel that is leaky due to CT invasion in stage IV lung cancer. The micro blood vessel with leak caused by CT invasion is full of red blood cells (RBC, purple arrows), which spread and distribute around the local neighboring tissues. **C.** Representative image of blood vessels with leaks caused by severe CT invasion. This triggered blood vessel disassembly and decomposition in stage IV prostate cancer. There are many CT strands and fragments (CTS, orange arrows) and red blood cells (RBC, purple arrows). The local prostate cancer cells invaded into CTs (white arrows) and migrate away, leaving spaces without tissue cells. Scale bar:10μm.

**Figure S27.**
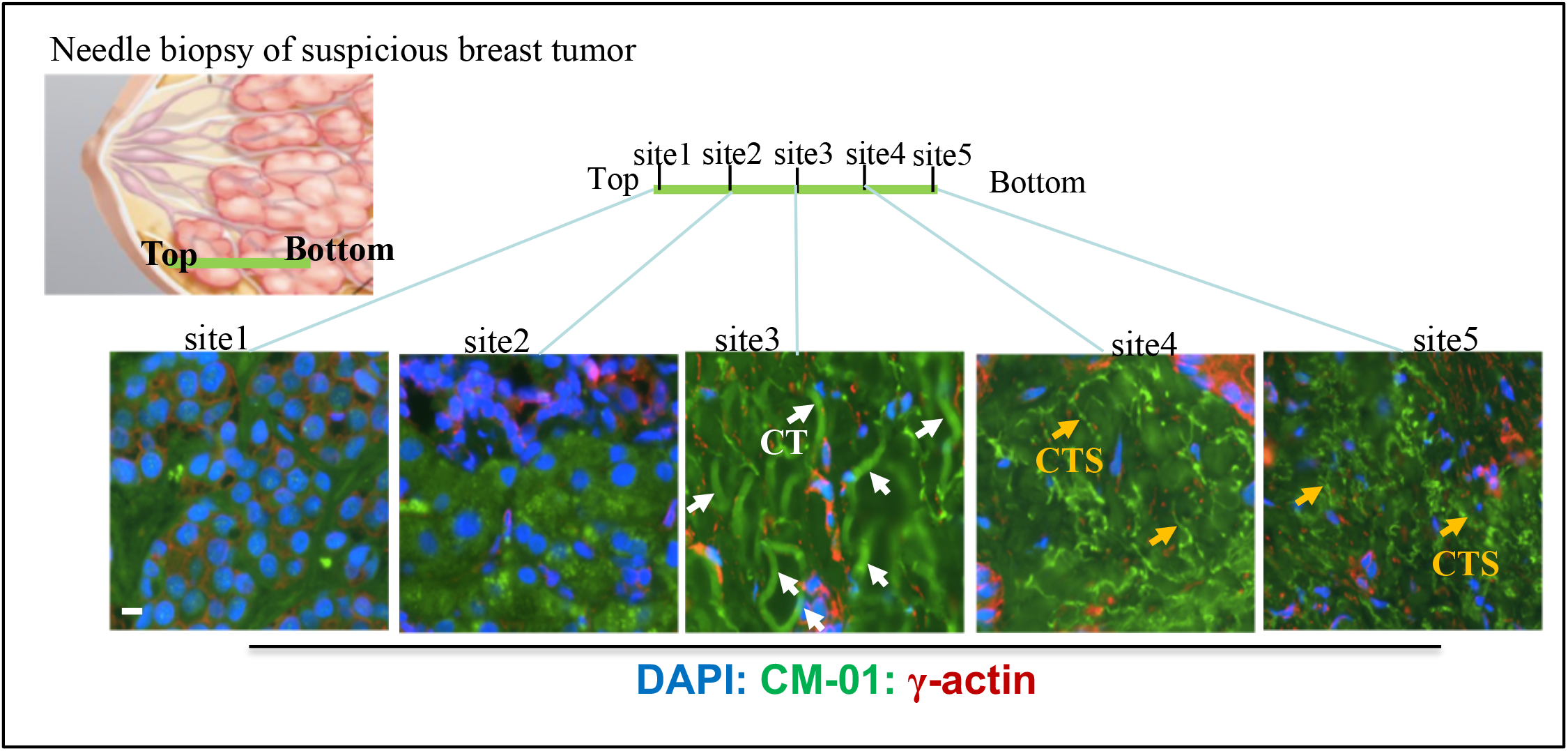
Continuous multi-layered tissue scanning of cancer cell cytocapsular tube in needle biopsy for cancer prognosis and diagnosis. Representative images of multiple sites in needle biopsy with suspicious breast cancer tissues by CT analyses for breast cancer diagnosis. In site 3, there are many breast cancer cell CTs without degradation. In sites 4 and 5, there are numerous breast cancer cell CT strands in degradation. Scale bar:10μm.

**Movie S1:**
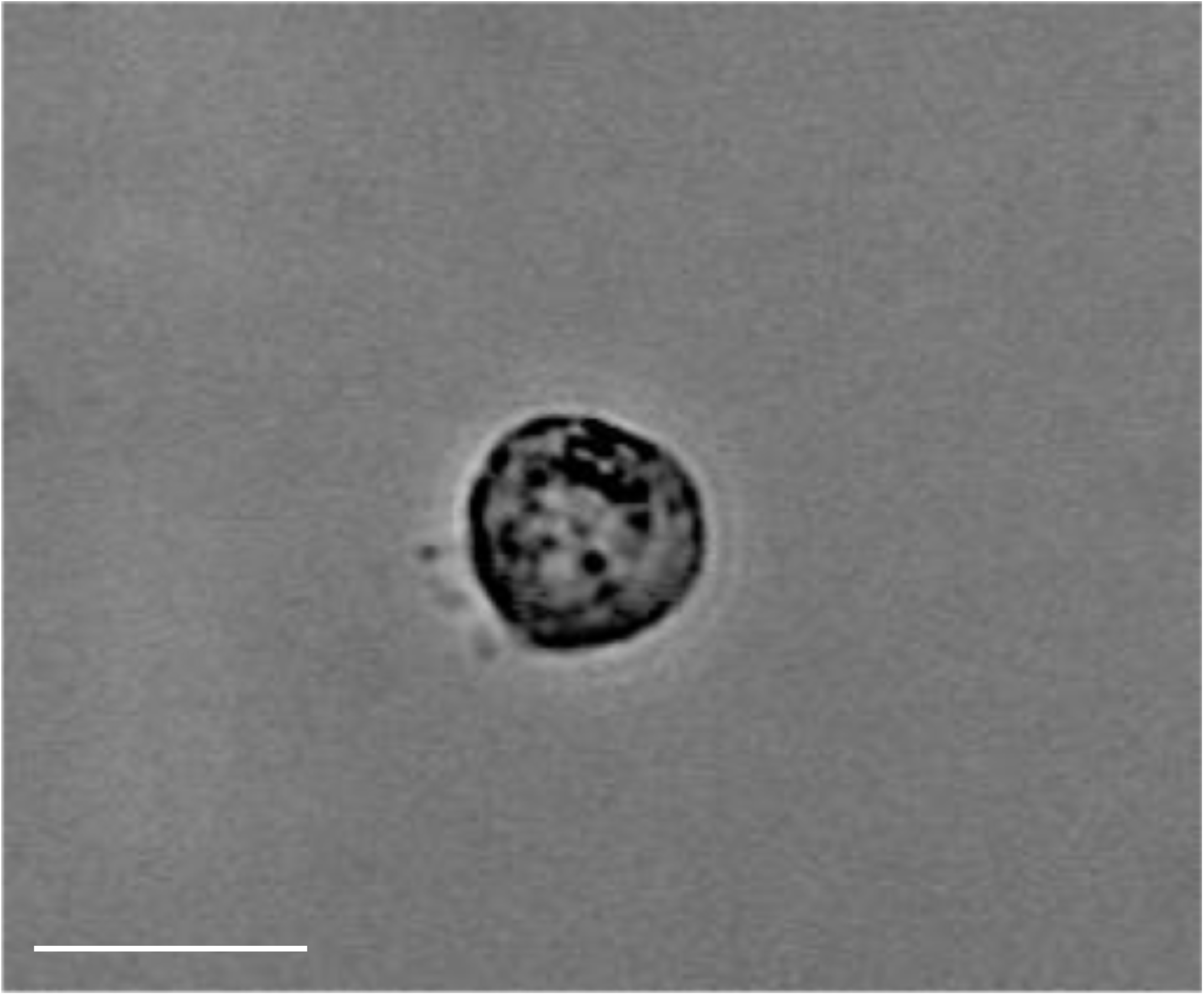
Movie of cancer stem cell (CSC) cytocapsular-tube initiation, generation, and bi-directionally migration in the CSC CT it generated. A single breast cancer stem cell from the HMLER(CD44^high^/CD24^low^)^FA^ subpopulation^1^ generates a short cytocapsular tube in the 3D Matrigel matrix. Subsequently, it migrates bi-directionally in the CT it engendered. Initially, the cell generates a cytocapsula outside of and disconnected from the plasma membrane. Then the cell moves to the left to elongate the cytocapsular membrane to engender a short cytocapsular tube (CT) in its wake. Then, the cell migrates in the CT to the right. There are several transient spike-like features of still unknown structure dynamically connecting the rear of the cell with the CT membranes. Scale bar:10μm.

**Movie S2:**
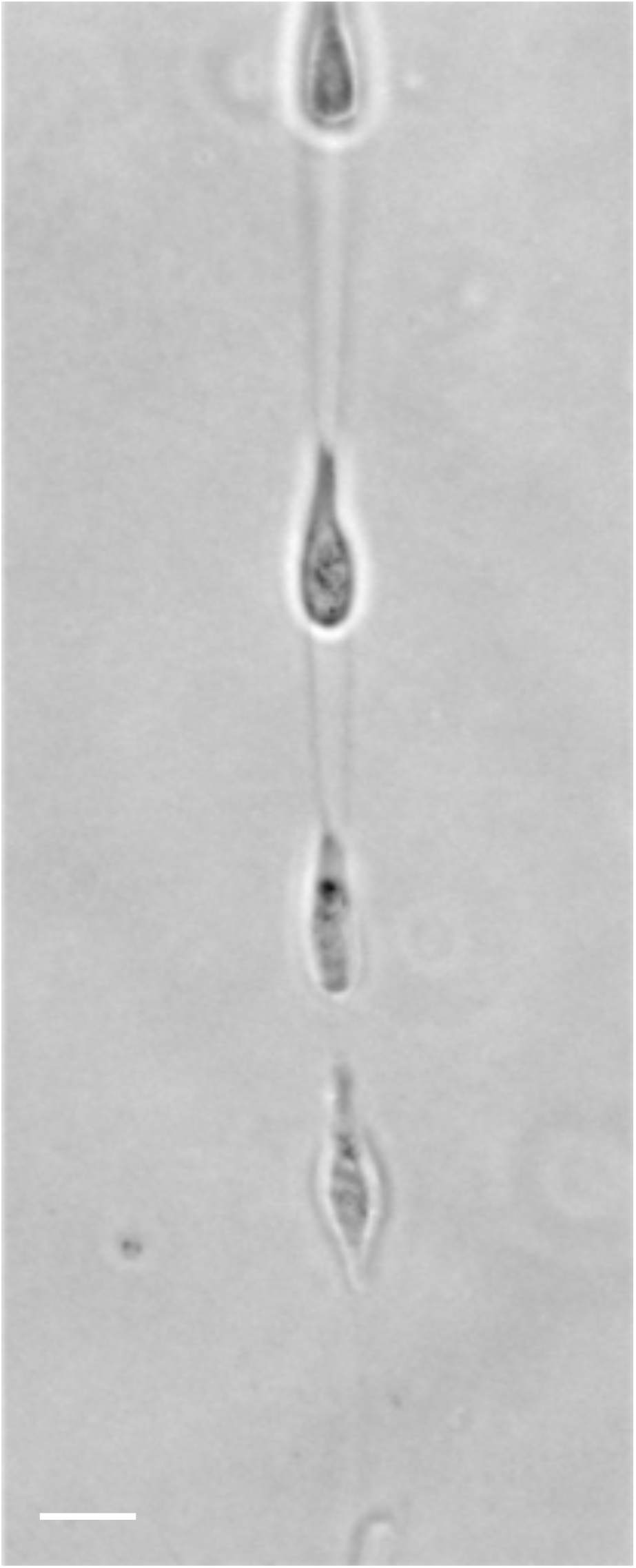
Multiple cancer stem cells bi-directionally migrate in a single long CSC cytocapsular tube. There are 4 breast cancer stem cell HMLER(CD44^high^/CD24^low^)^FA^ subpopulation cells migrate in a long cytocapsular tube (CT) generated by another breast CSC. The top CSC bi-directionally migrates inside the breast CSC CT. Scale bar:10μm.

---

**Table S1.** Clinical characteristics of cytocapsular tubes (CTs) in benign tumor tissues.

| Benign tumor types | Subtypes | Patients/specimens | with CT | % with CT |
| --- | --- | --- | --- | --- |
| Breast | Fibrosis and Simple Cysts | 11 |  |  |
|  | Ductal or Lobular Hyperplasia | 13 | 3 |  |
|  | Lobular Carcinoma in Situ | 2 |  |  |
|  | Adenosi | 1 |  |  |
|  | Fibroadenomas | 1 |  |  |
|  | Phyllodes Tumors | 1 | 1 |  |
|  | Intraductal Papillomas | 8 | 1 |  |
|  | Granular Cell Tumors | 3 | 1 |  |
|  | Fat Necrosis and Oil Cysts | 1 |  |  |
|  | Mastitis | 2 |  |  |
|  | Duct Ectasia | 2 |  |  |
| Lung | Hamartomas | 2 |  |  |
|  | Bronchial adenomas | 3 |  |  |
|  | Papillomas | 3 |  |  |
| Prostate | Benign prostatic hyperplasia | 4 | 1 |  |
| Stomach | fundic gland polyp | 12 | 4 |  |
|  | hyperplastic polyp | 1 |  |  |
|  | inflammatory fibroid polyps | 2 |  |  |
|  | xanthoma | 1 |  |  |
|  | hamartomatous polyps | 1 |  |  |
|  | leiomyoma | 1 |  |  |
|  | lipoma | 1 |  |  |
|  | hemangioma | 2 |  |  |
|  | glomus tumour | 2 | 1 |  |
|  | cystic tumour | 1 |  |  |
| Liver | Hemangioma | 1 |  |  |
|  | Focal nodular hyperplasia | 1 |  |  |
|  | Hepatocellular adenoma | 2 |  |  |
| Colon | Hyperplastic polyps | 15 | 3 |  |
|  | Hamartomas | 11 | 1 |  |
|  | Juvenile polyps | 4 |  |  |
|  | Inflammatory polyps | 2 | 1 |  |
|  | Lymphoid polyps | 4 |  |  |
|  | Tubular adenomas | 1 |  |  |
|  | Villous adenomas | 2 |  |  |
|  | Tubulovillous adenomas | 2 |  |  |
|  | Total | 126 | 17 | 13.40% |

**Table S2.**
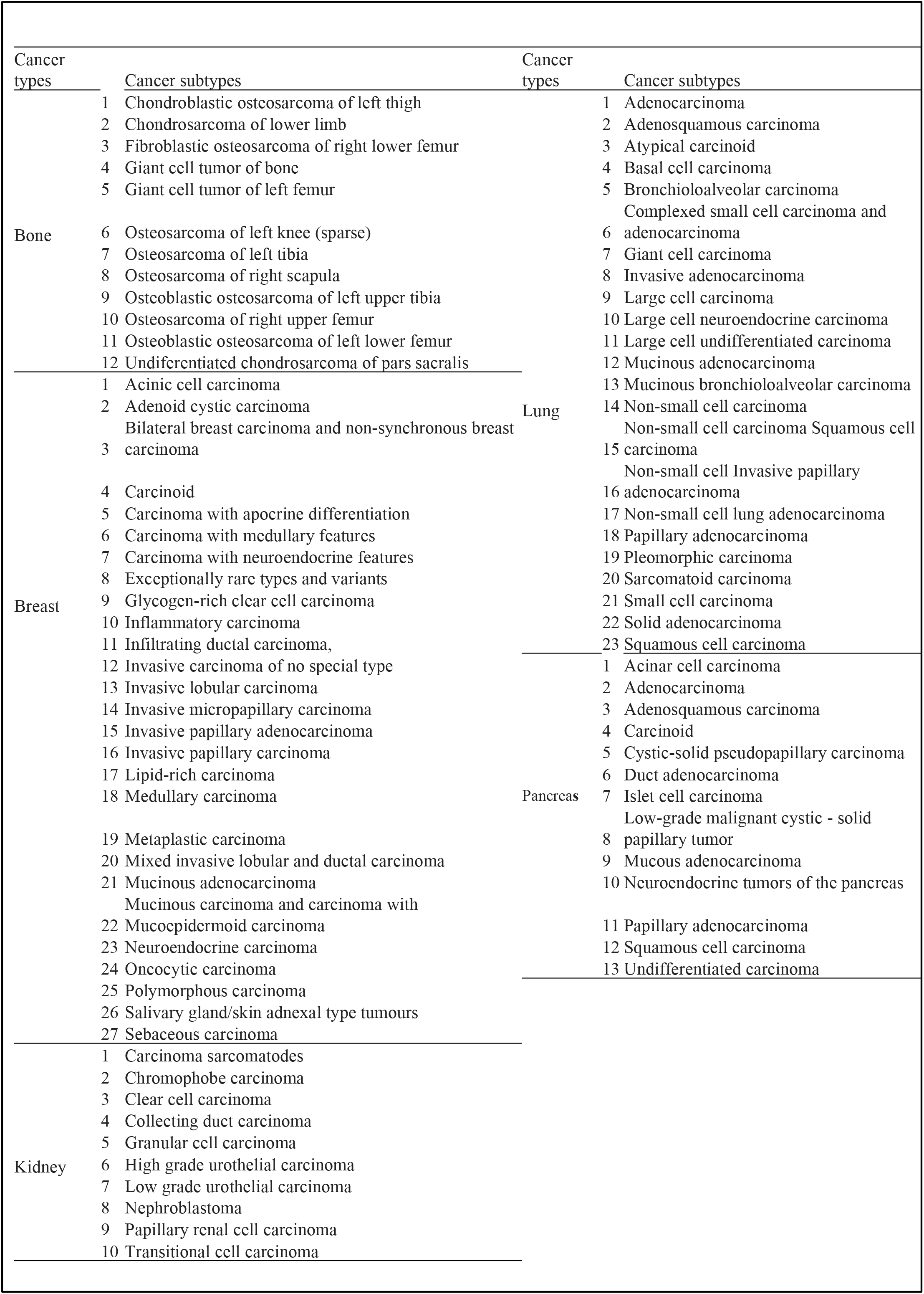
Subtypes of cancers with 10 (or more) kinds of investigated subtypes in Table 2.

**Table S3:**
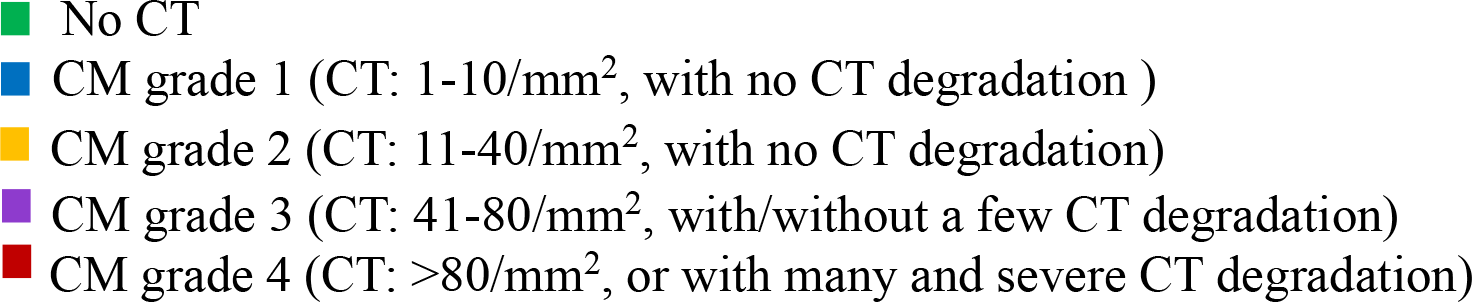
Cytocapsular tube (CT) density and suggested cytocapsular tube-based cancer metastasis (CM) grades:

**Table S4:** Relationship between conventional cancer stages and cancer metastasis (CM) grades based on the results in involved in Table S1 and Table 2:

| Cancer stage | Cancer Metastasis (CM) grade |
| --- | --- |
| 0 | 0, 1, 2, 3, 4 |
| I | 1, 2, 3, 4 |
| II | 2, 3, 4 |
| III | 3, 4 |
| IV | 4 |

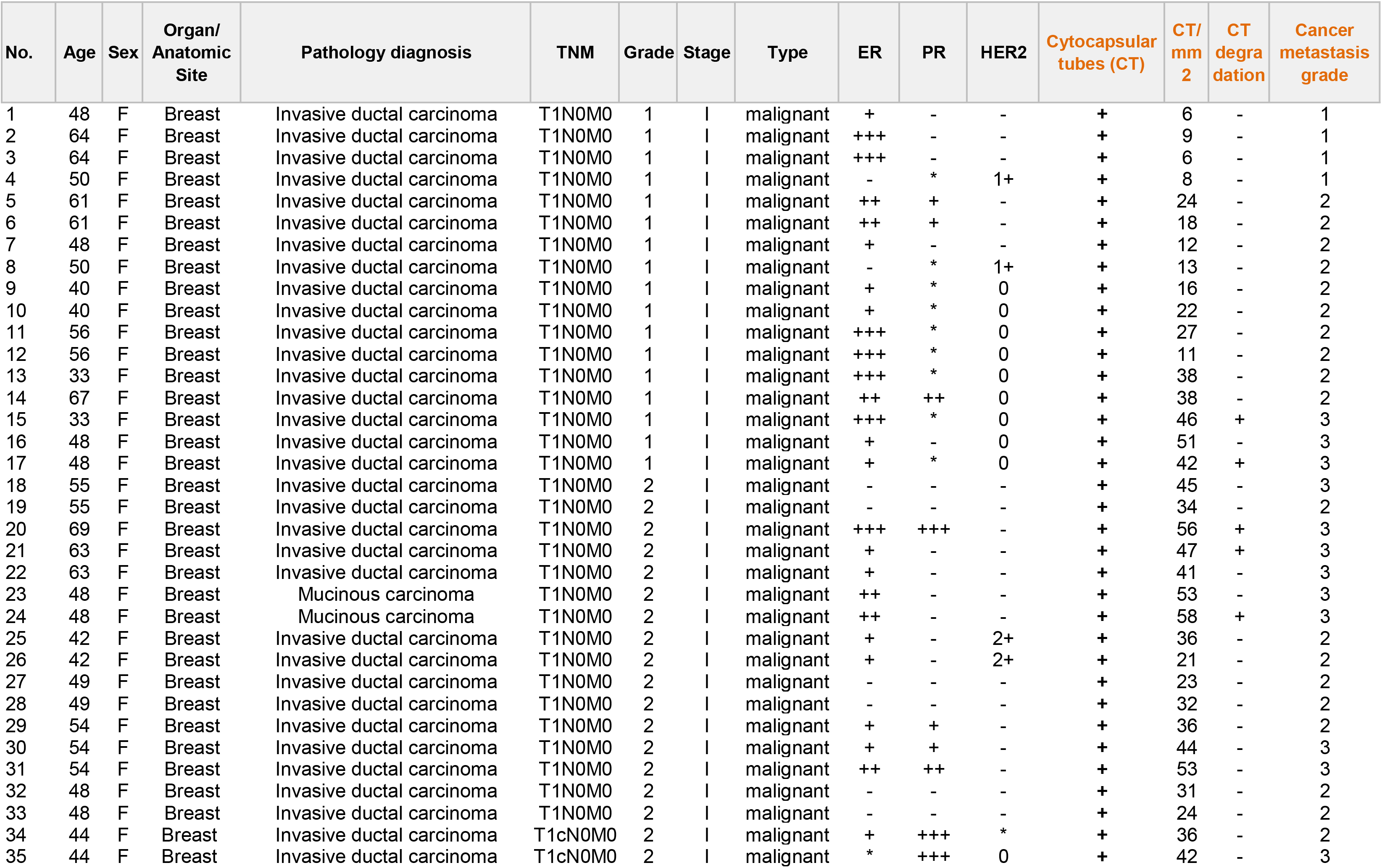
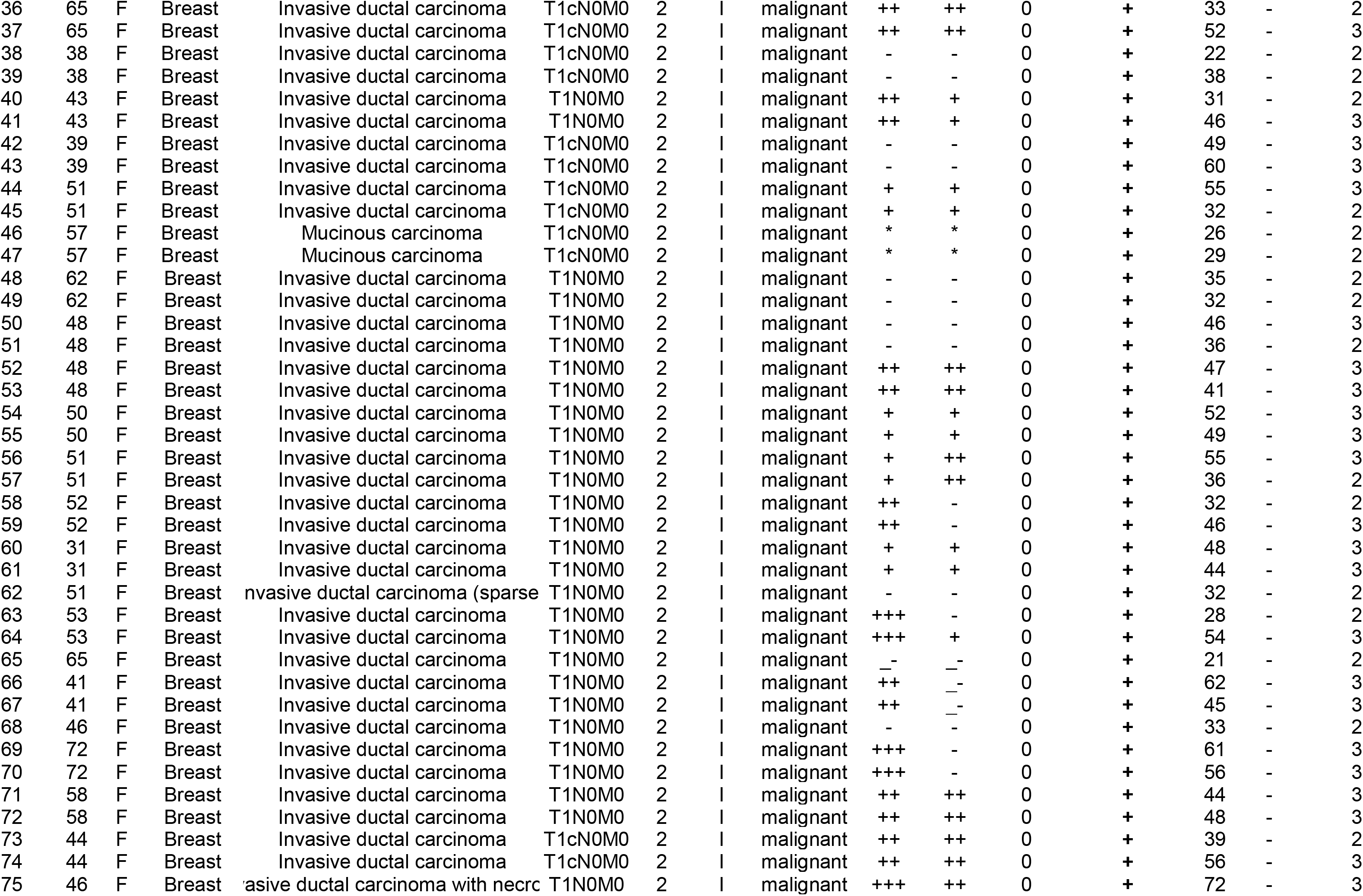
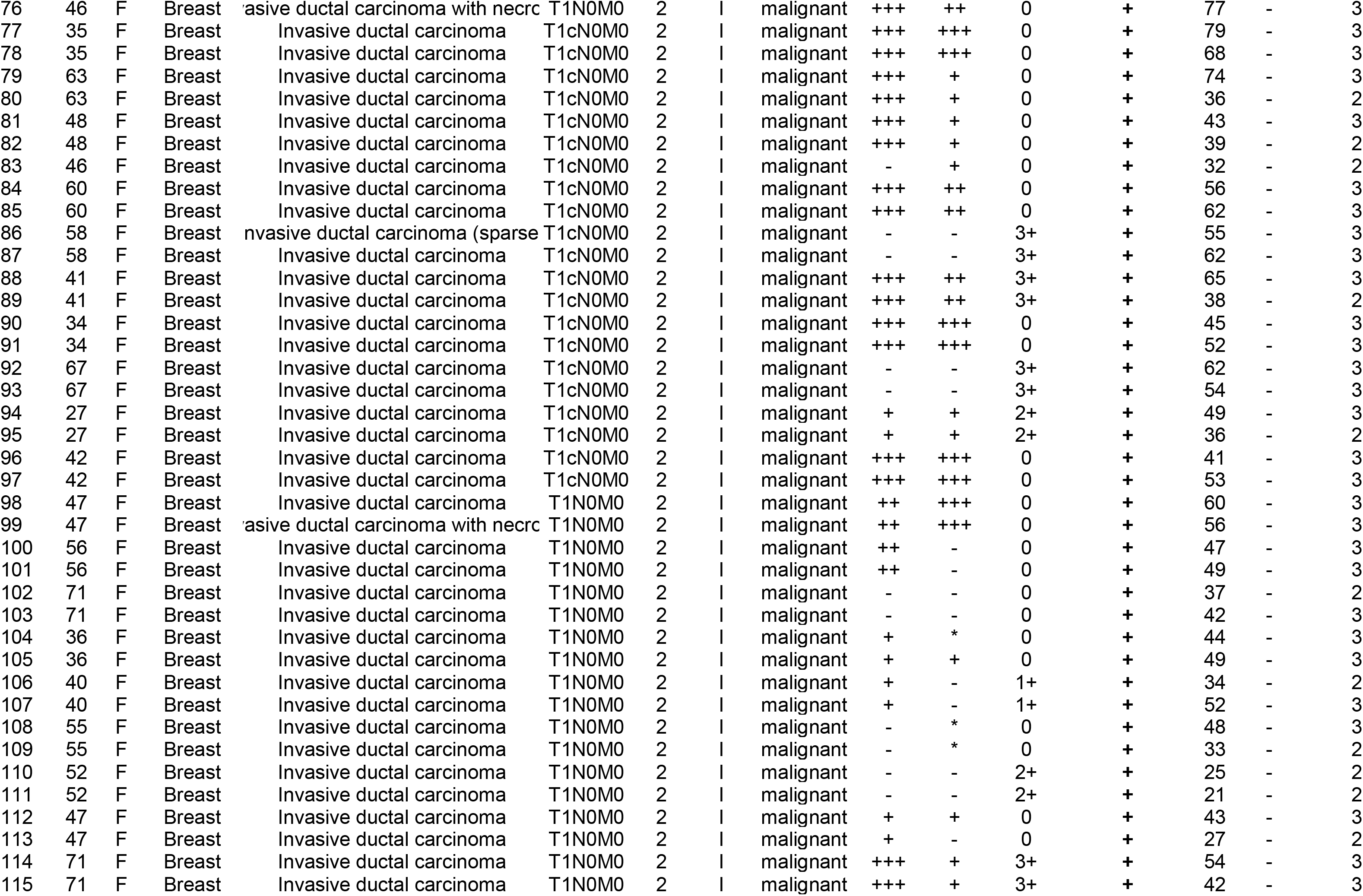
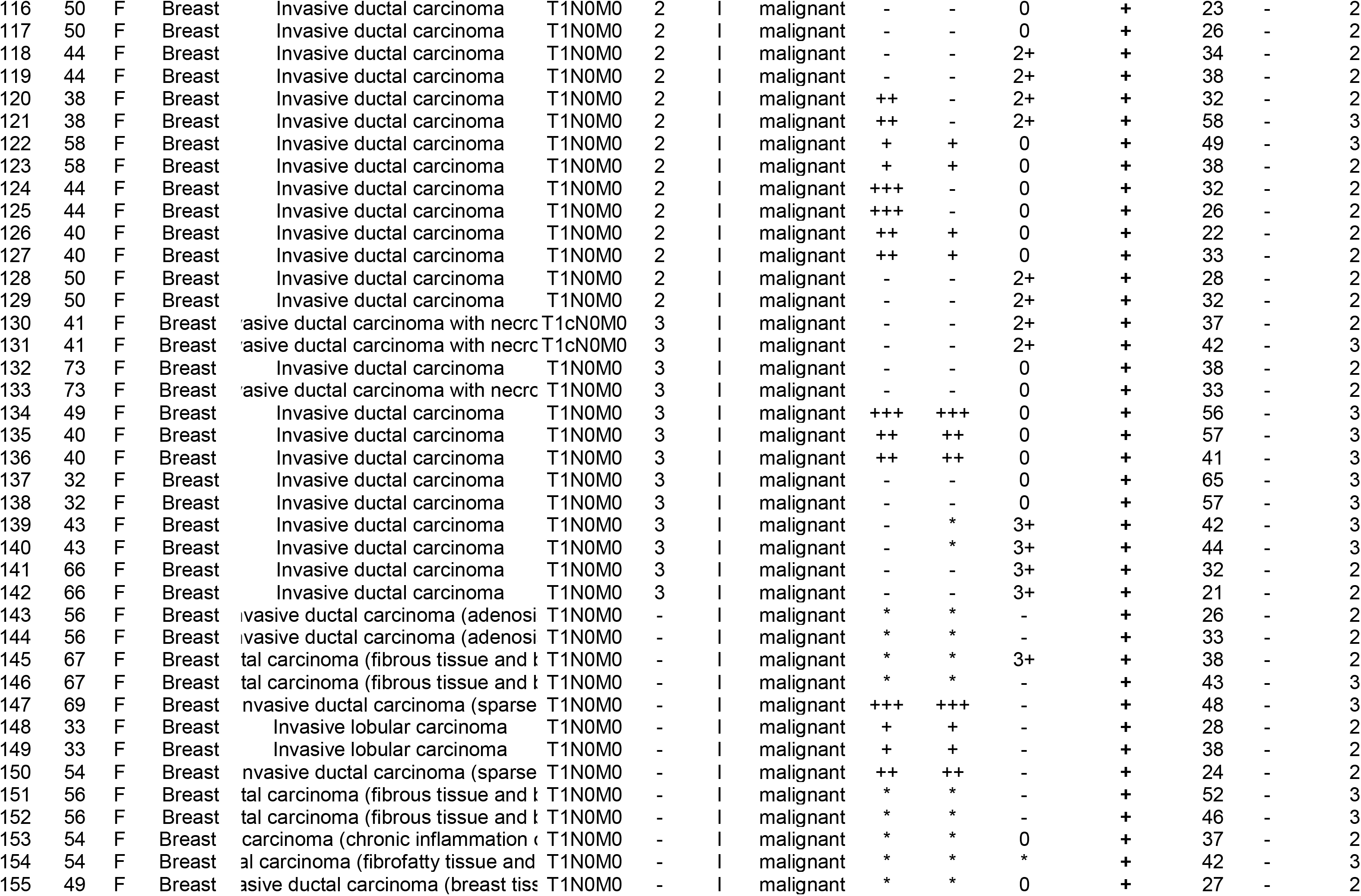
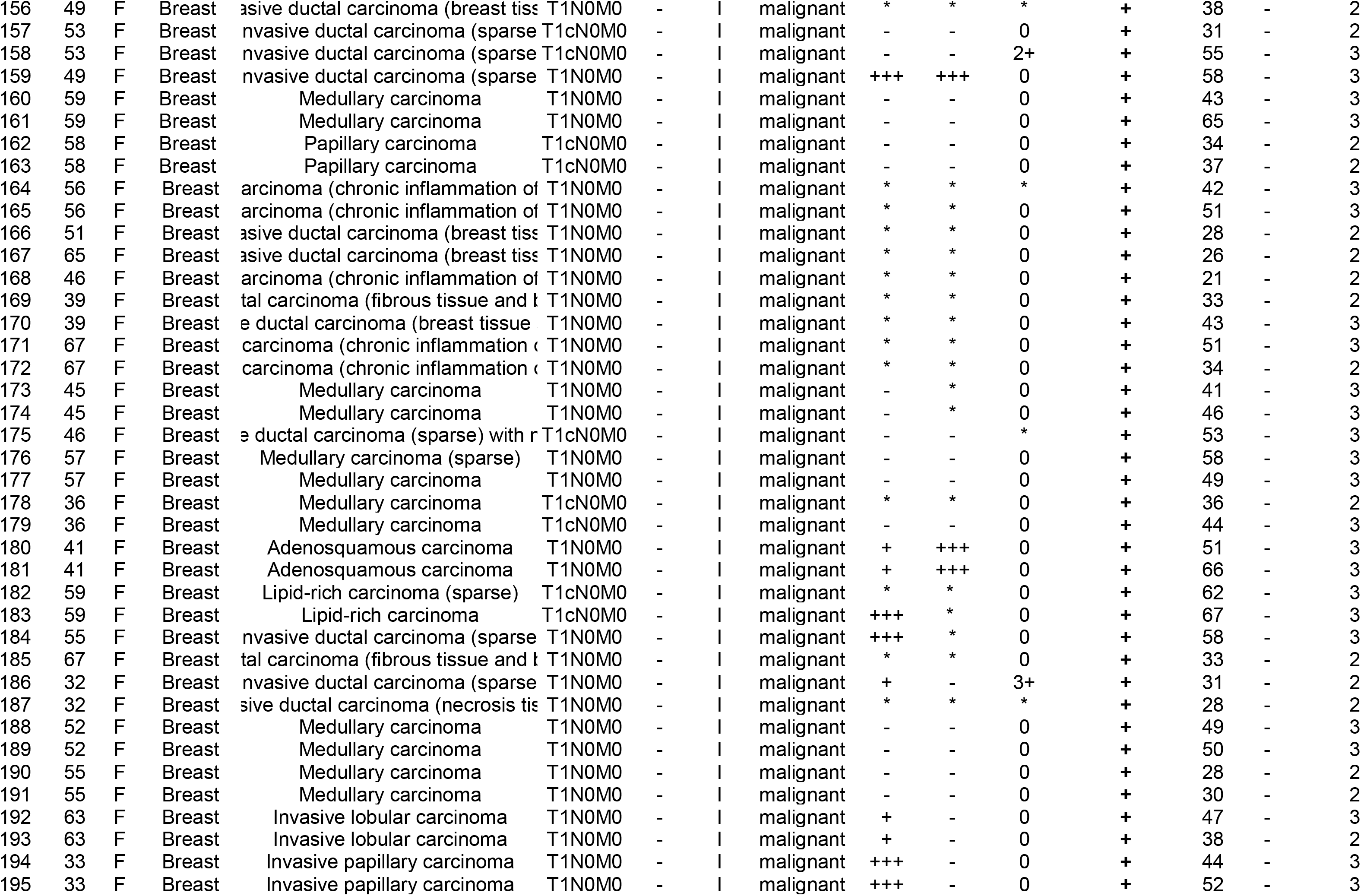
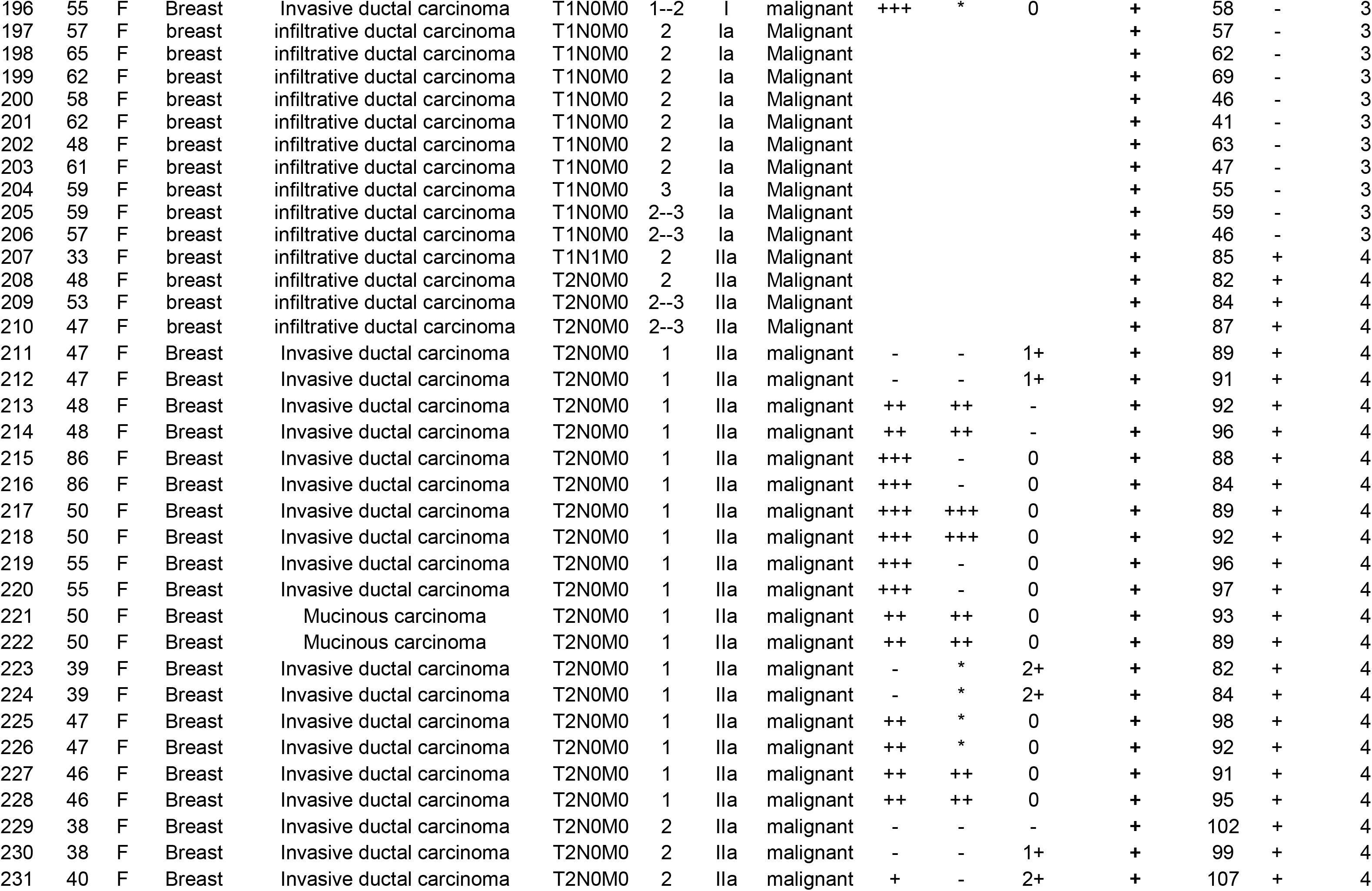
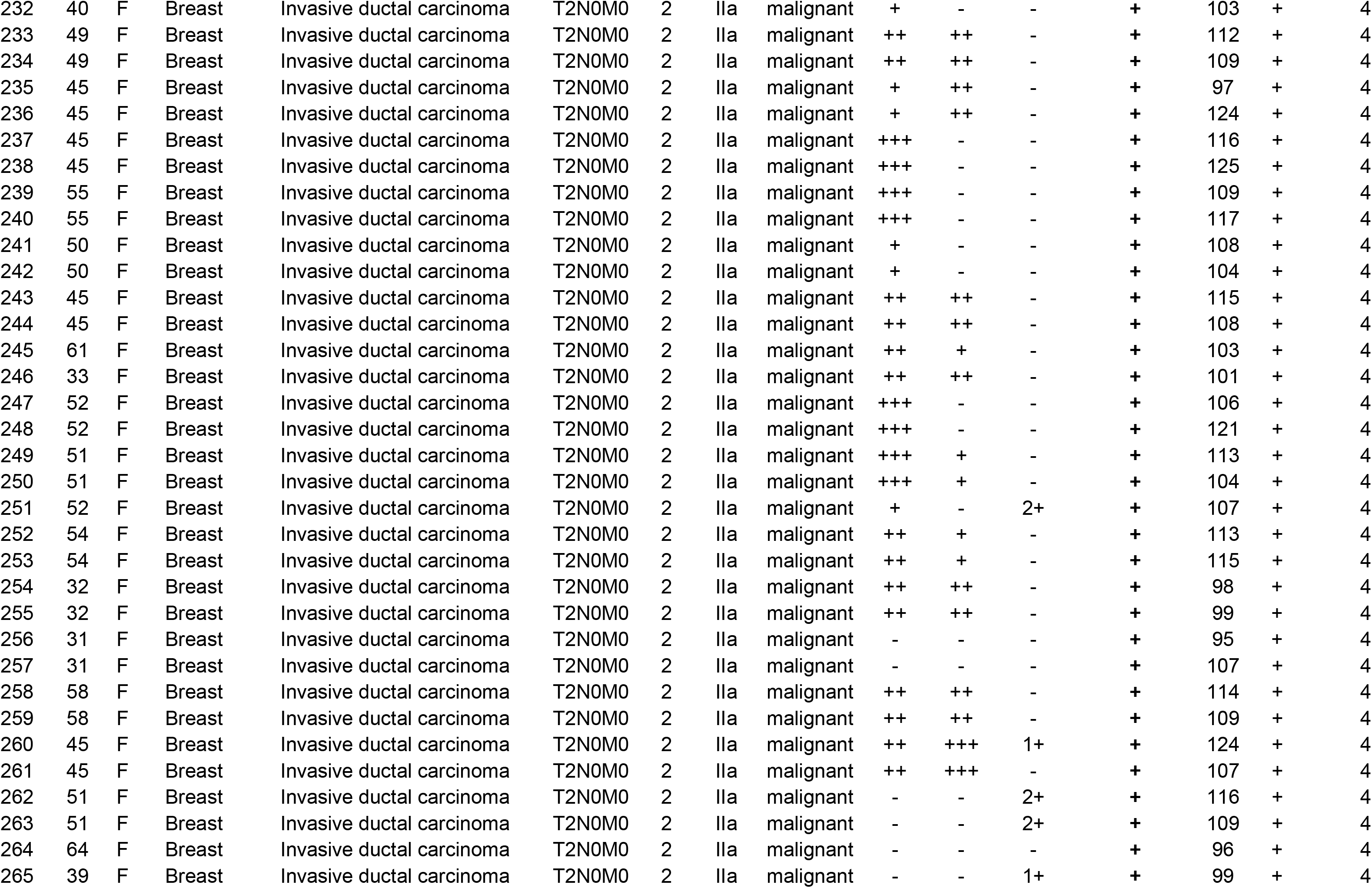
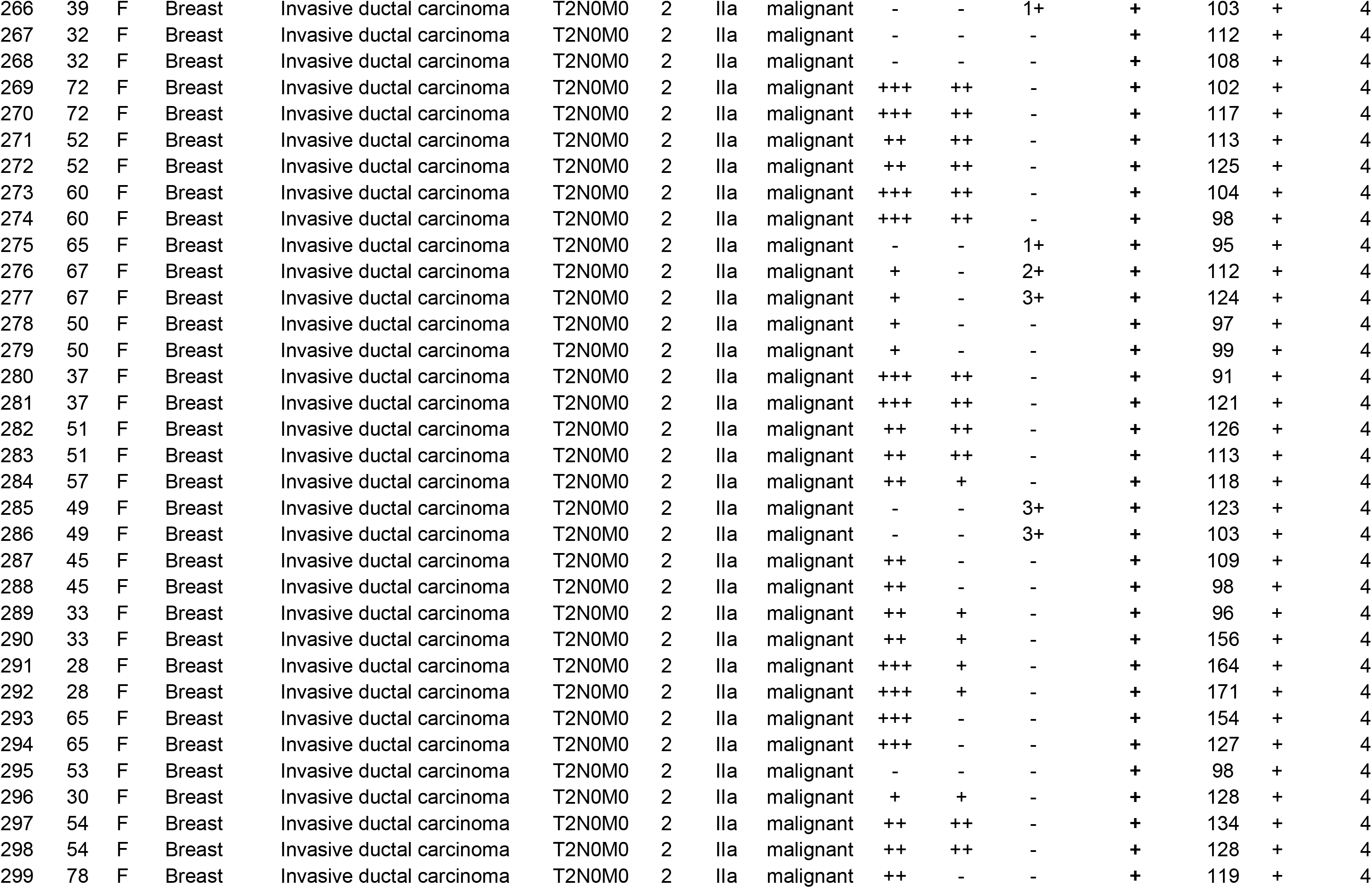
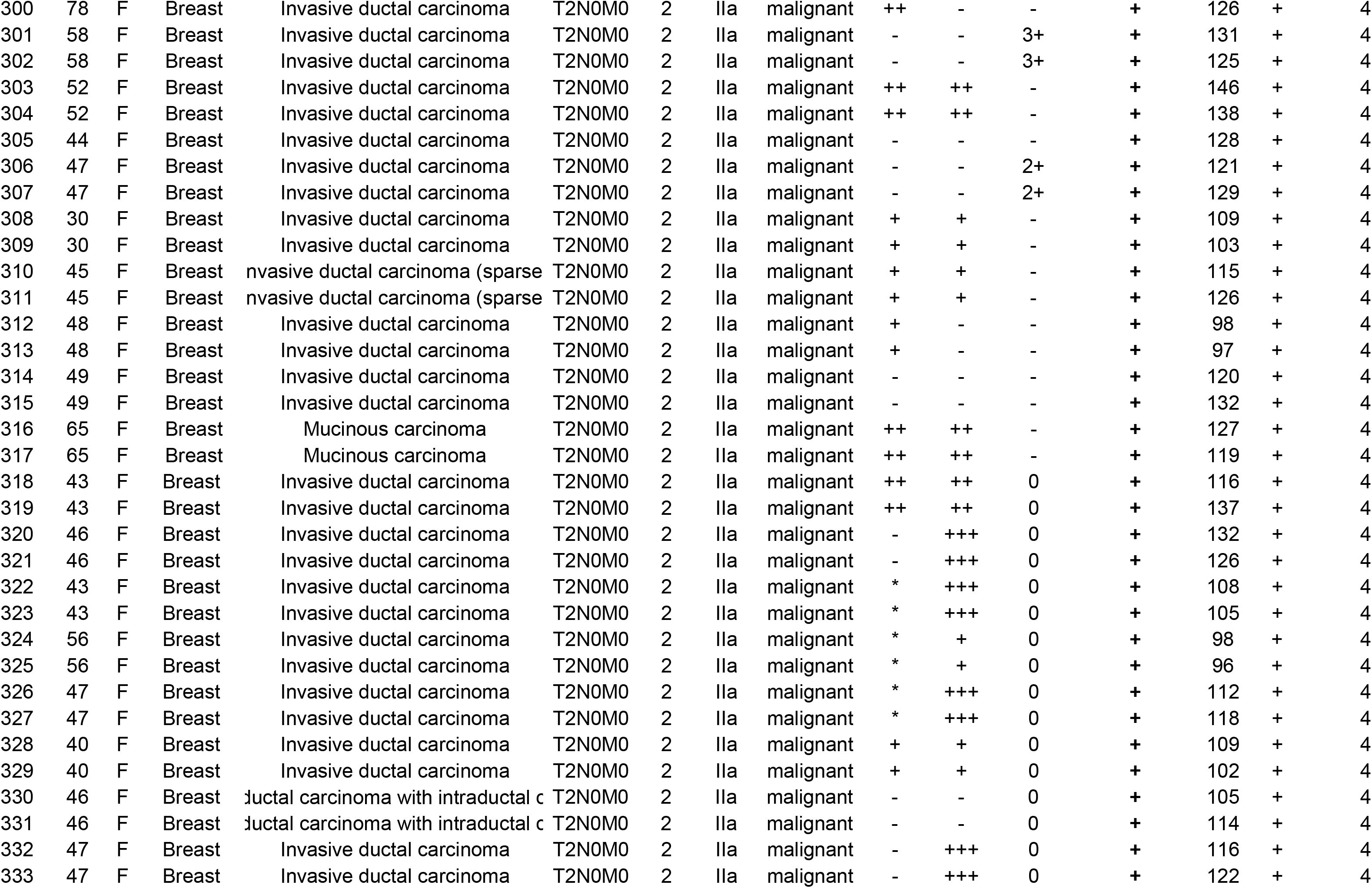
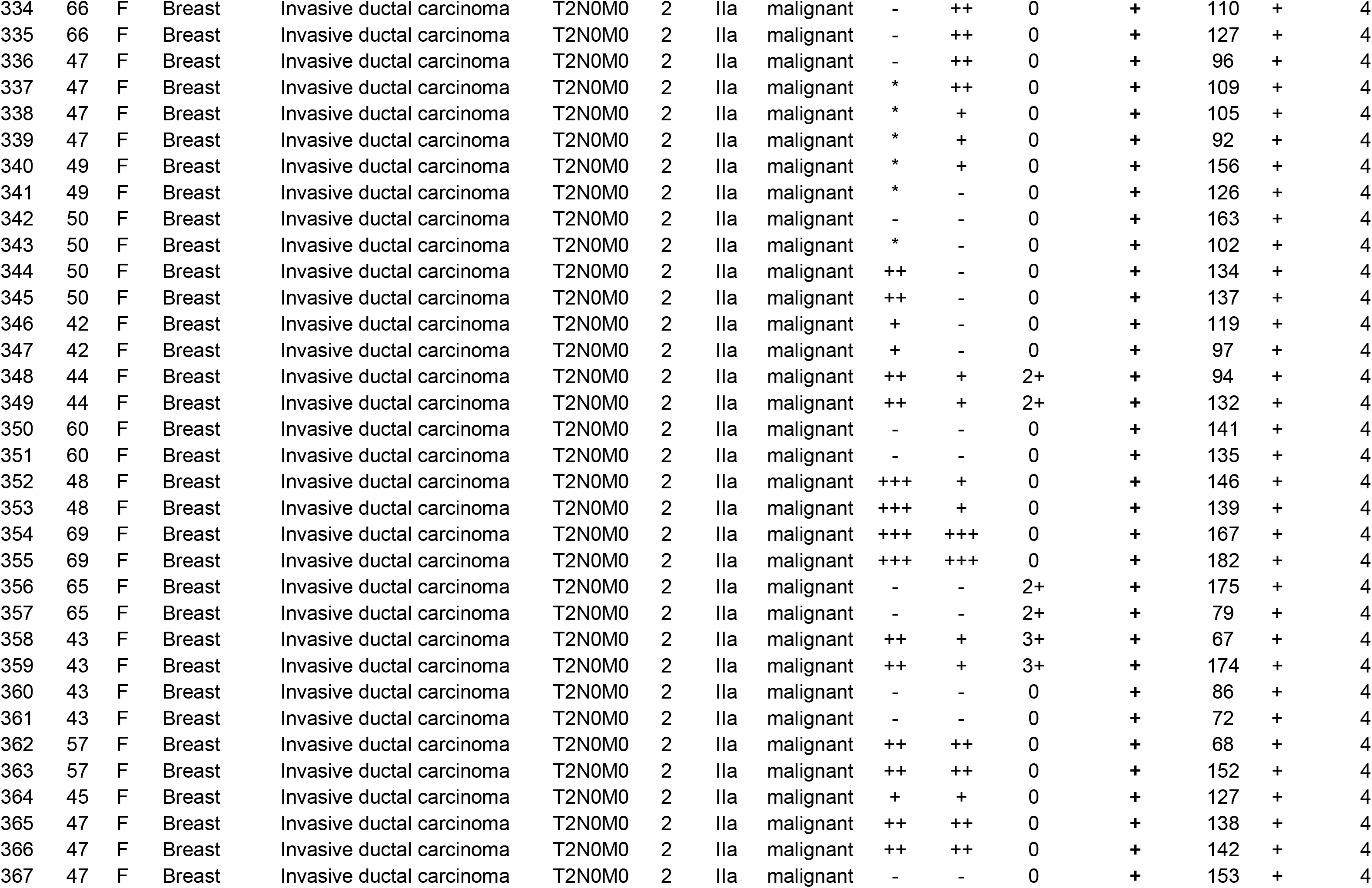
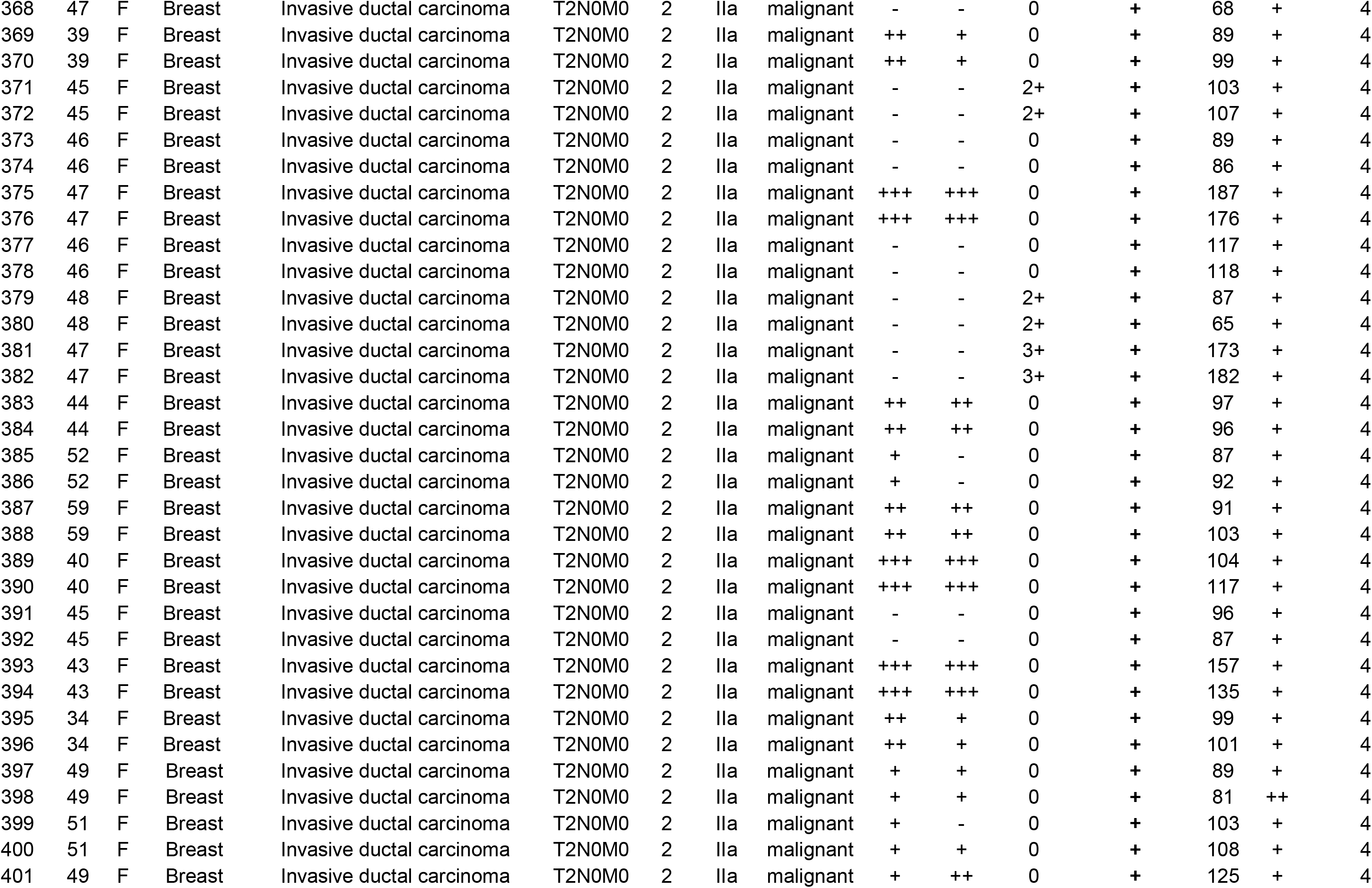
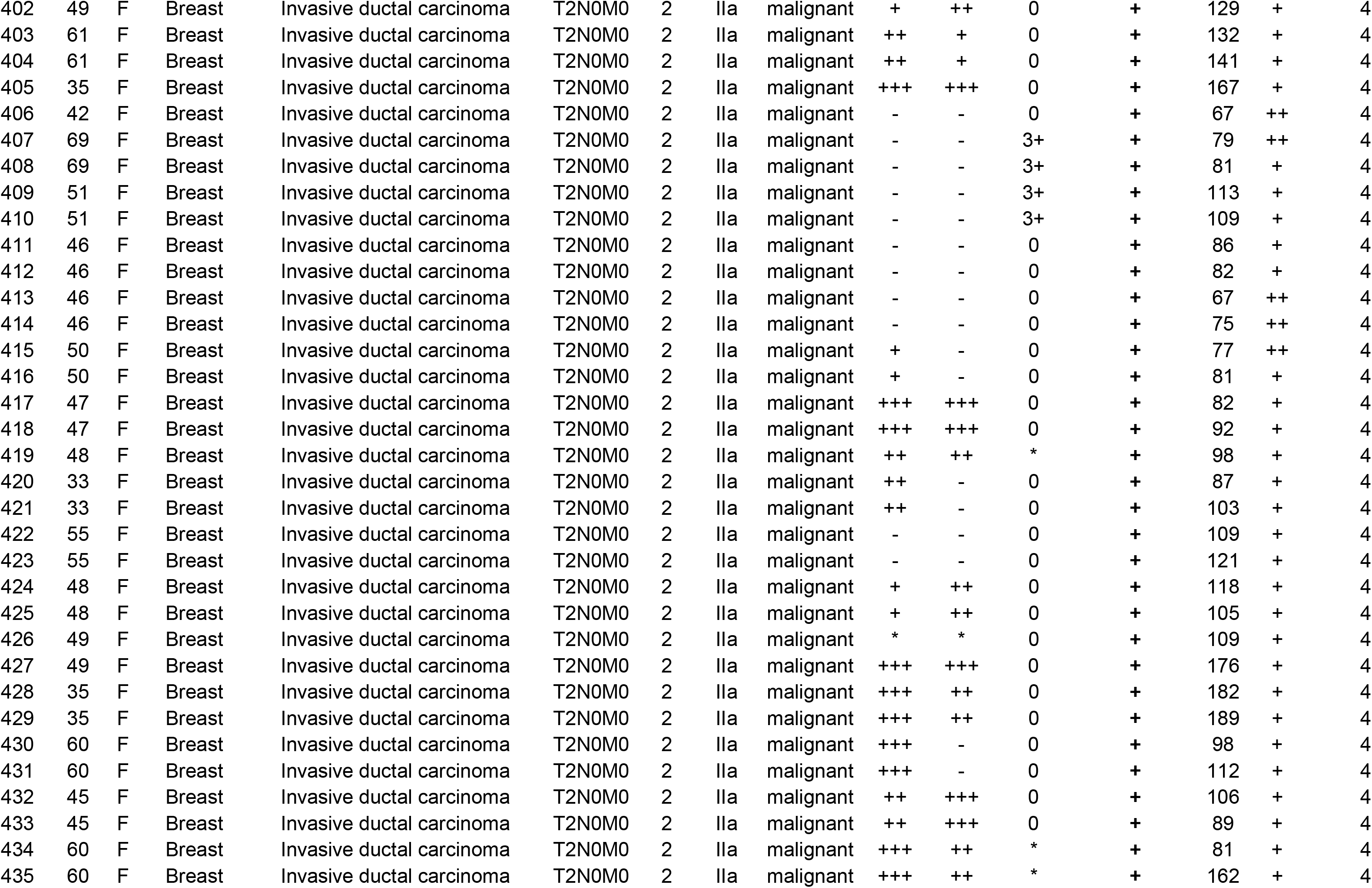
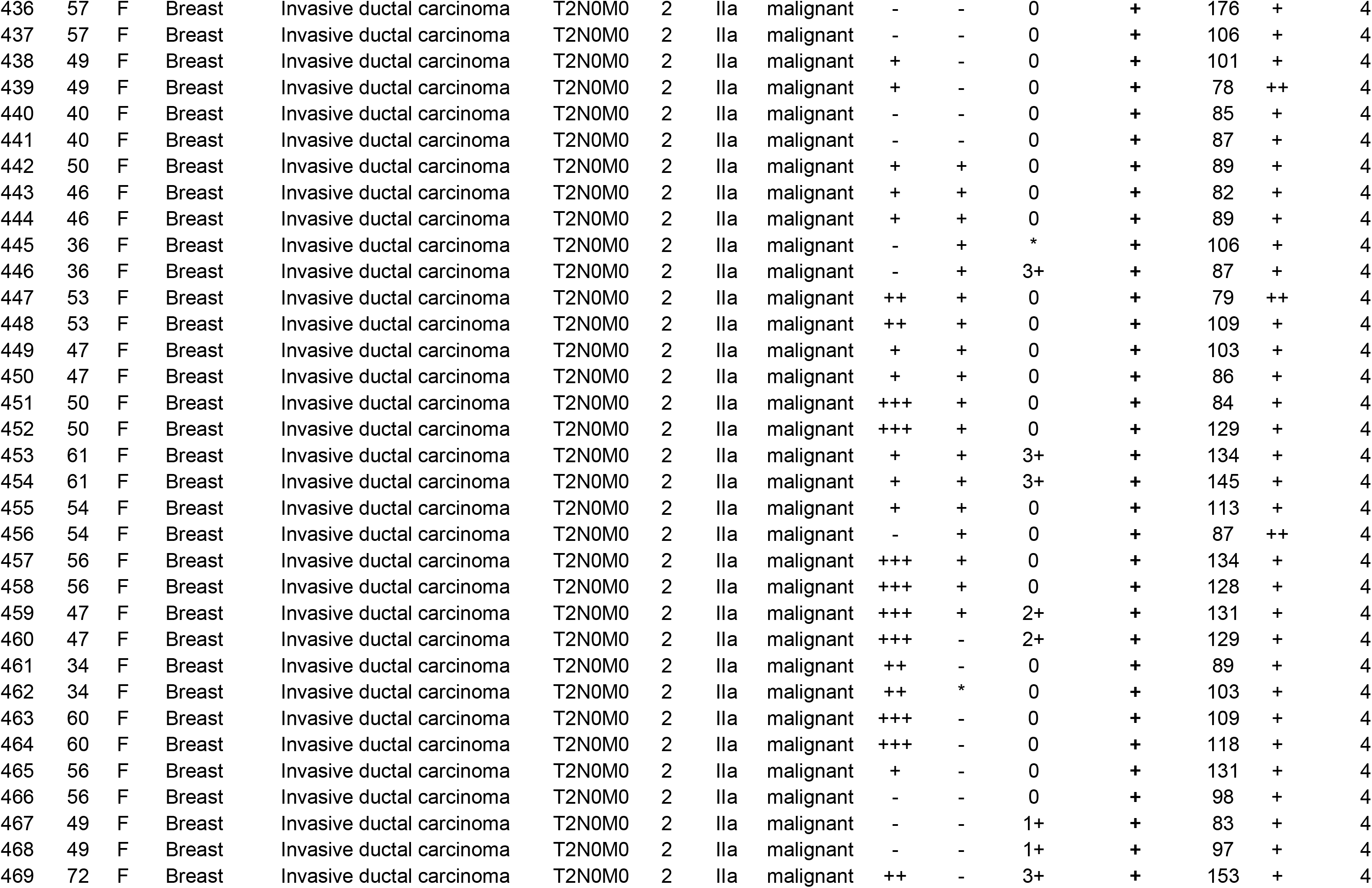
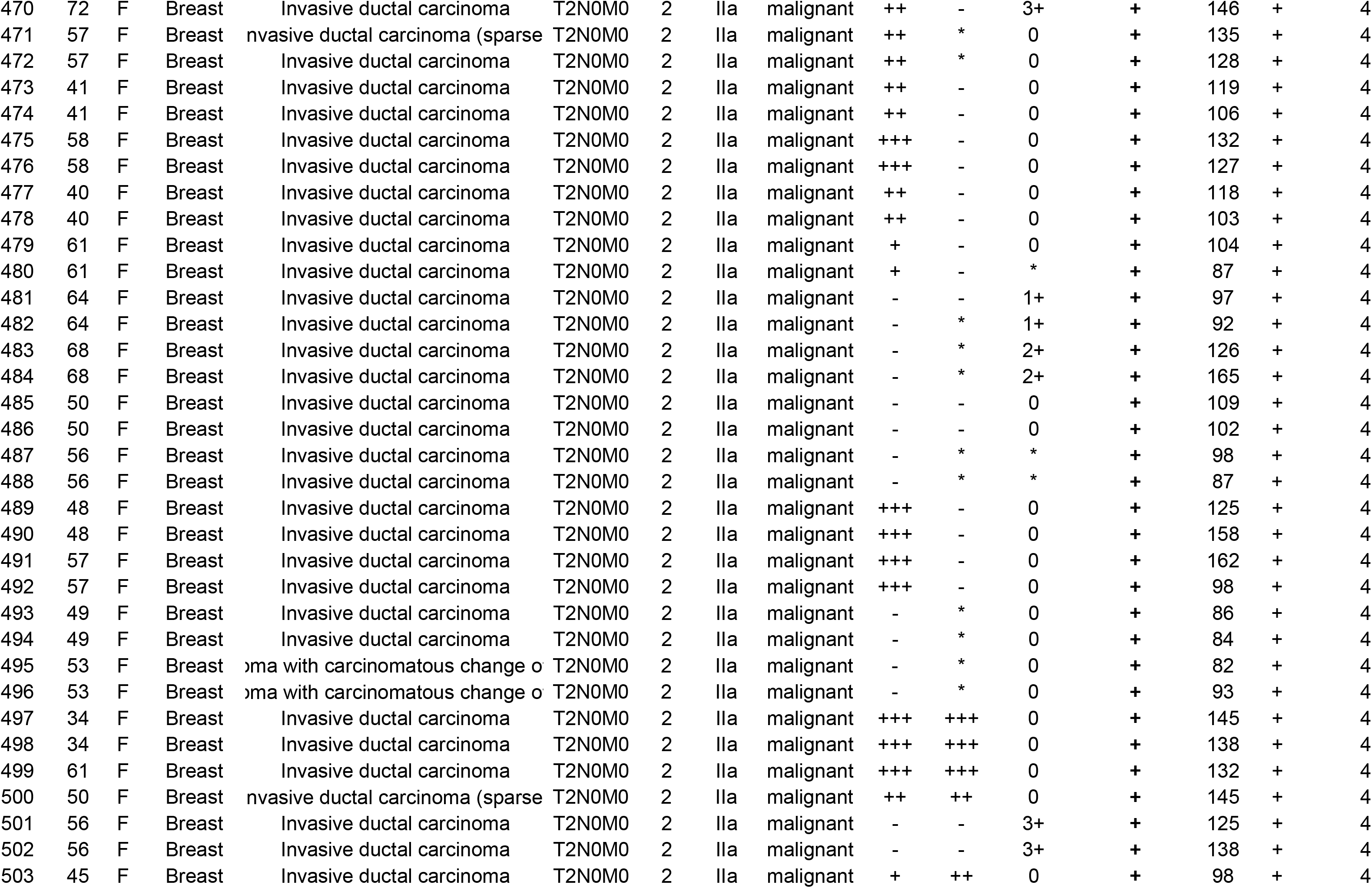
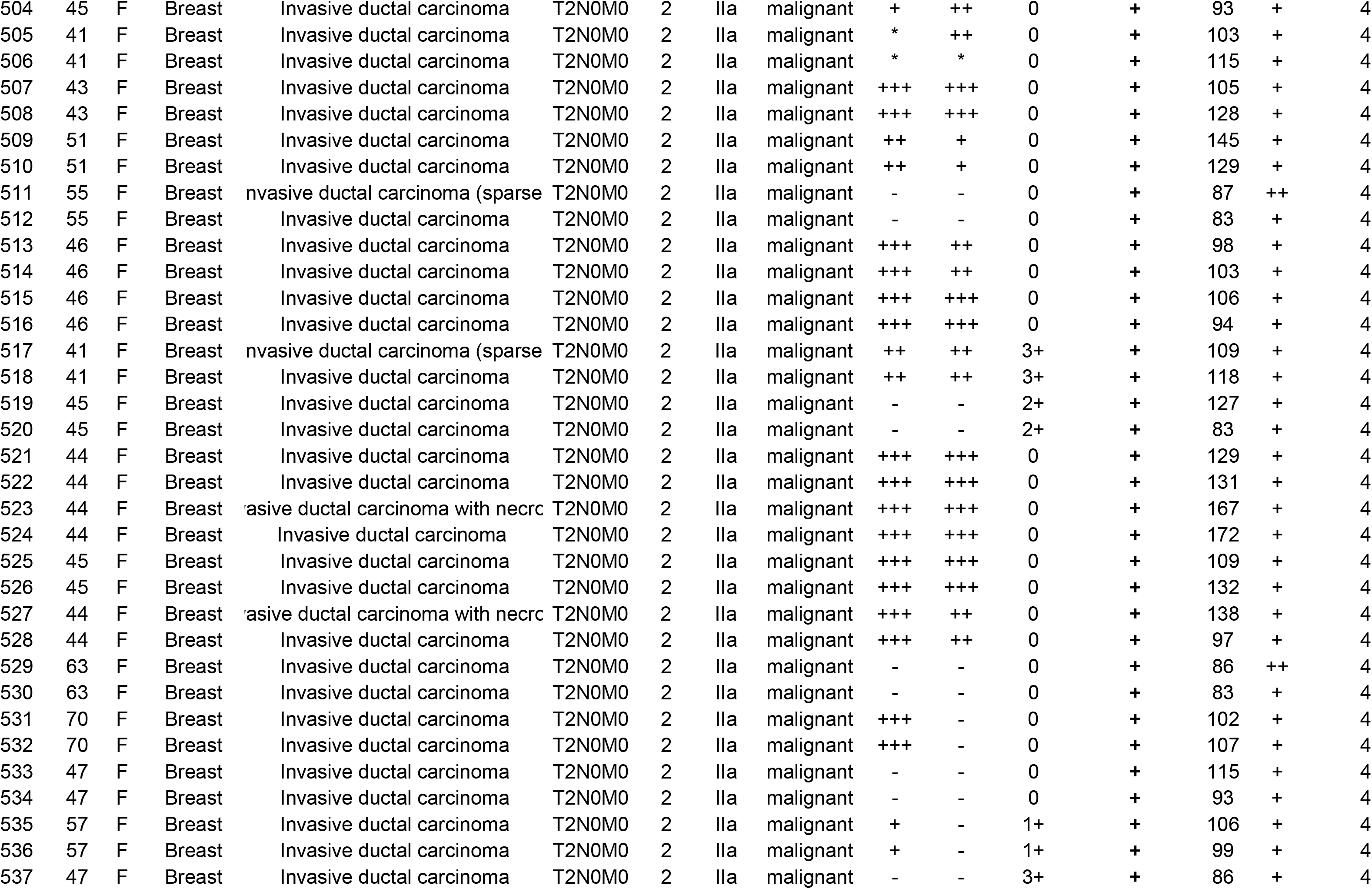
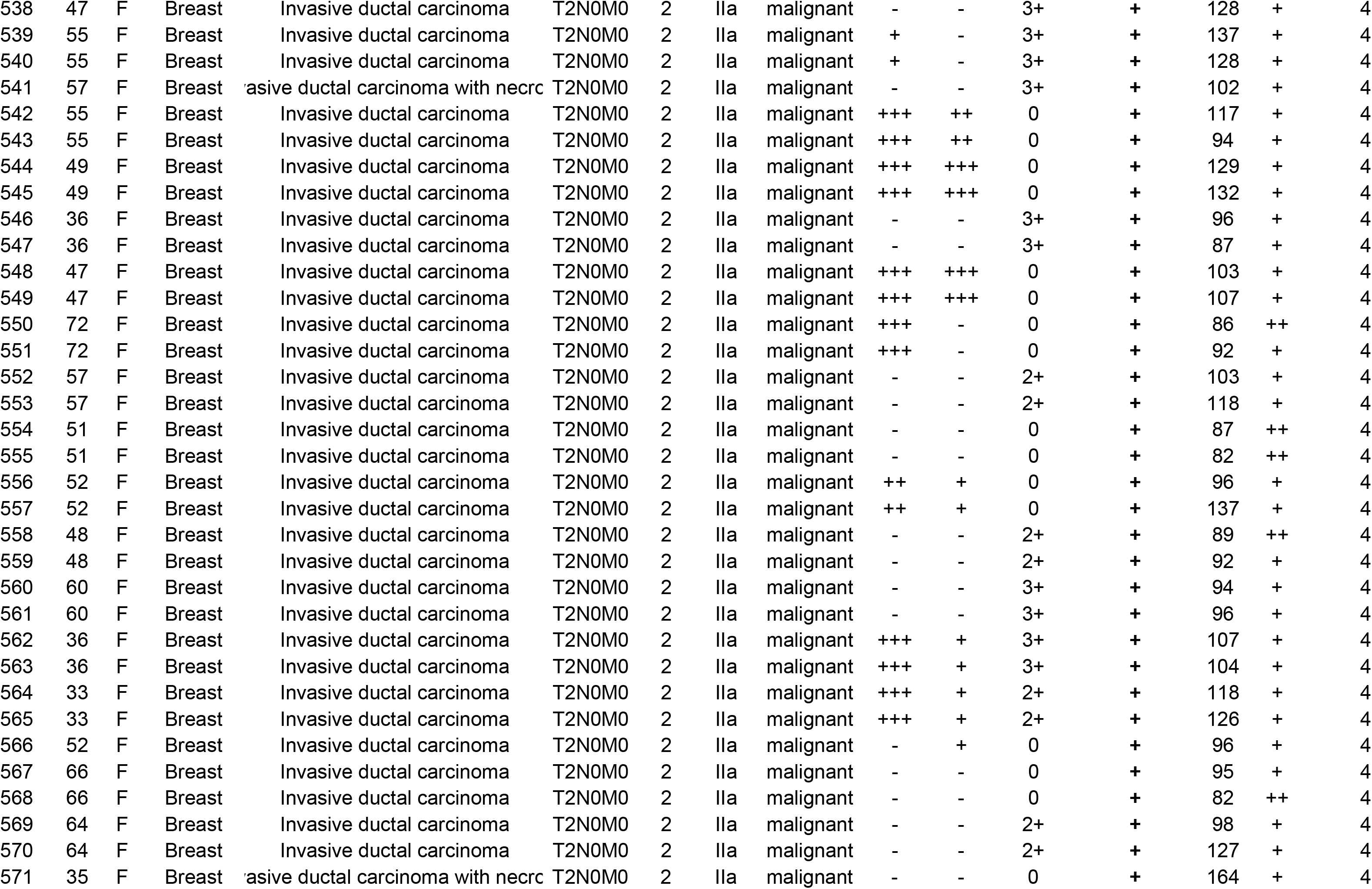
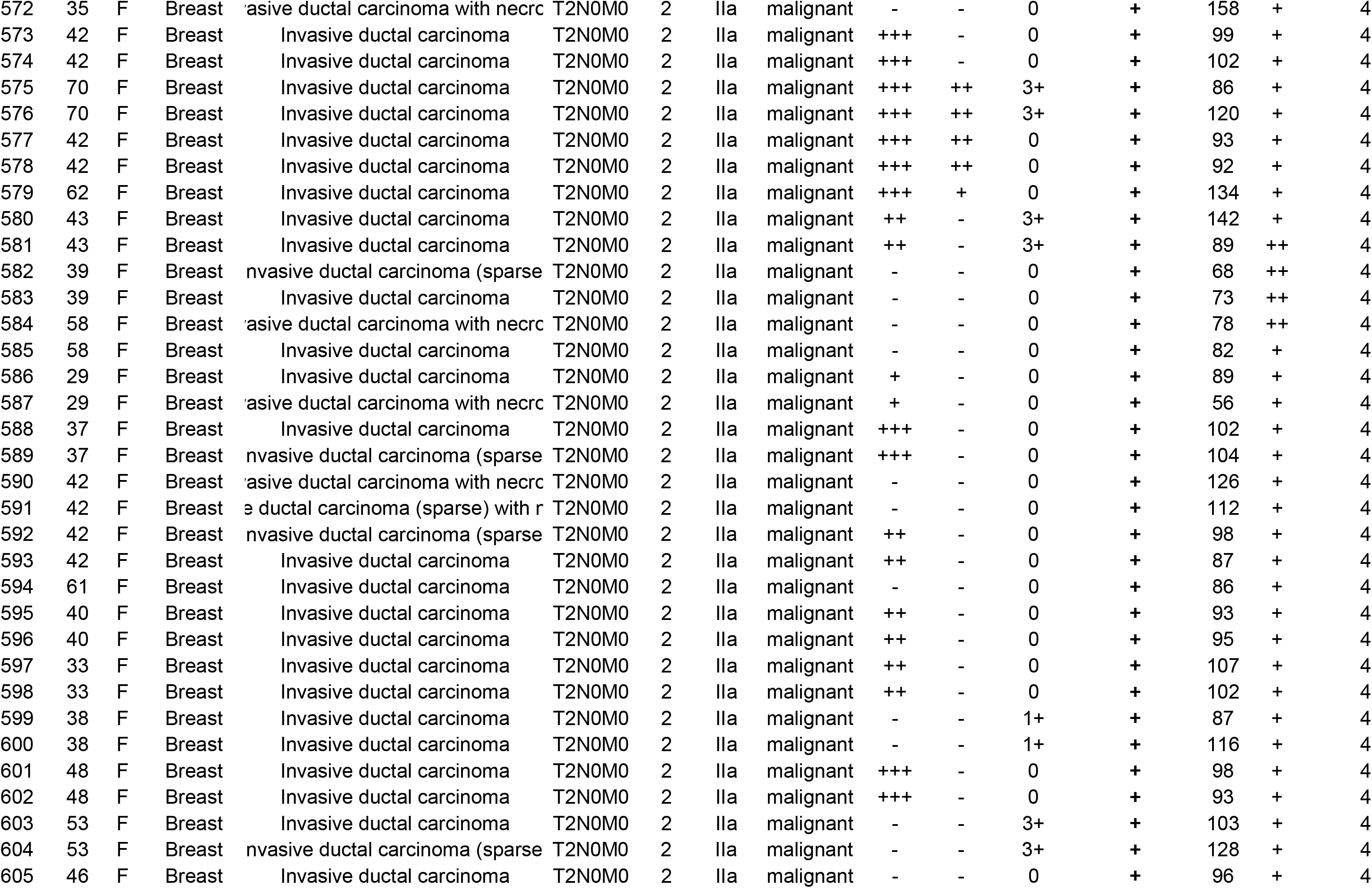
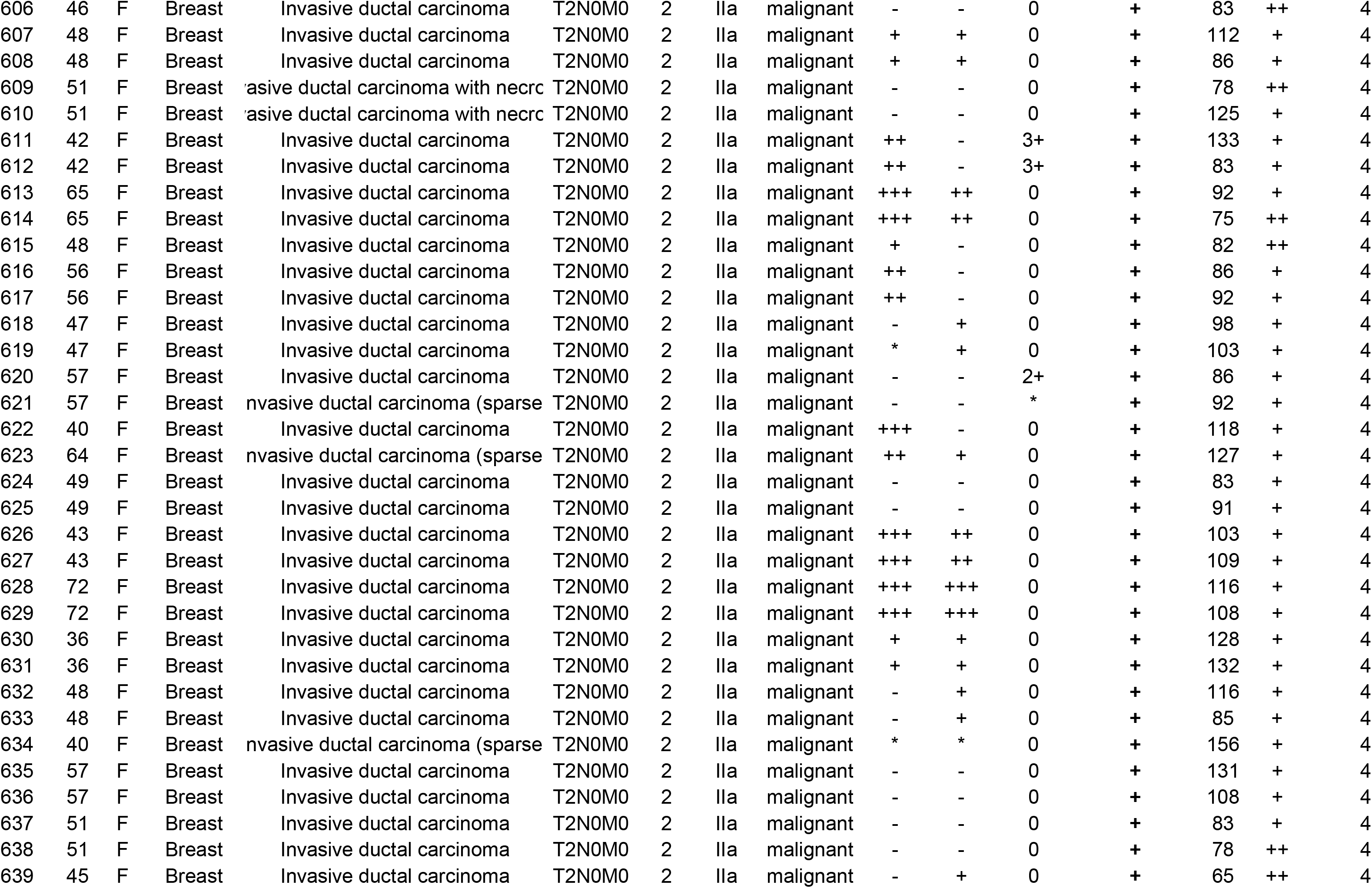
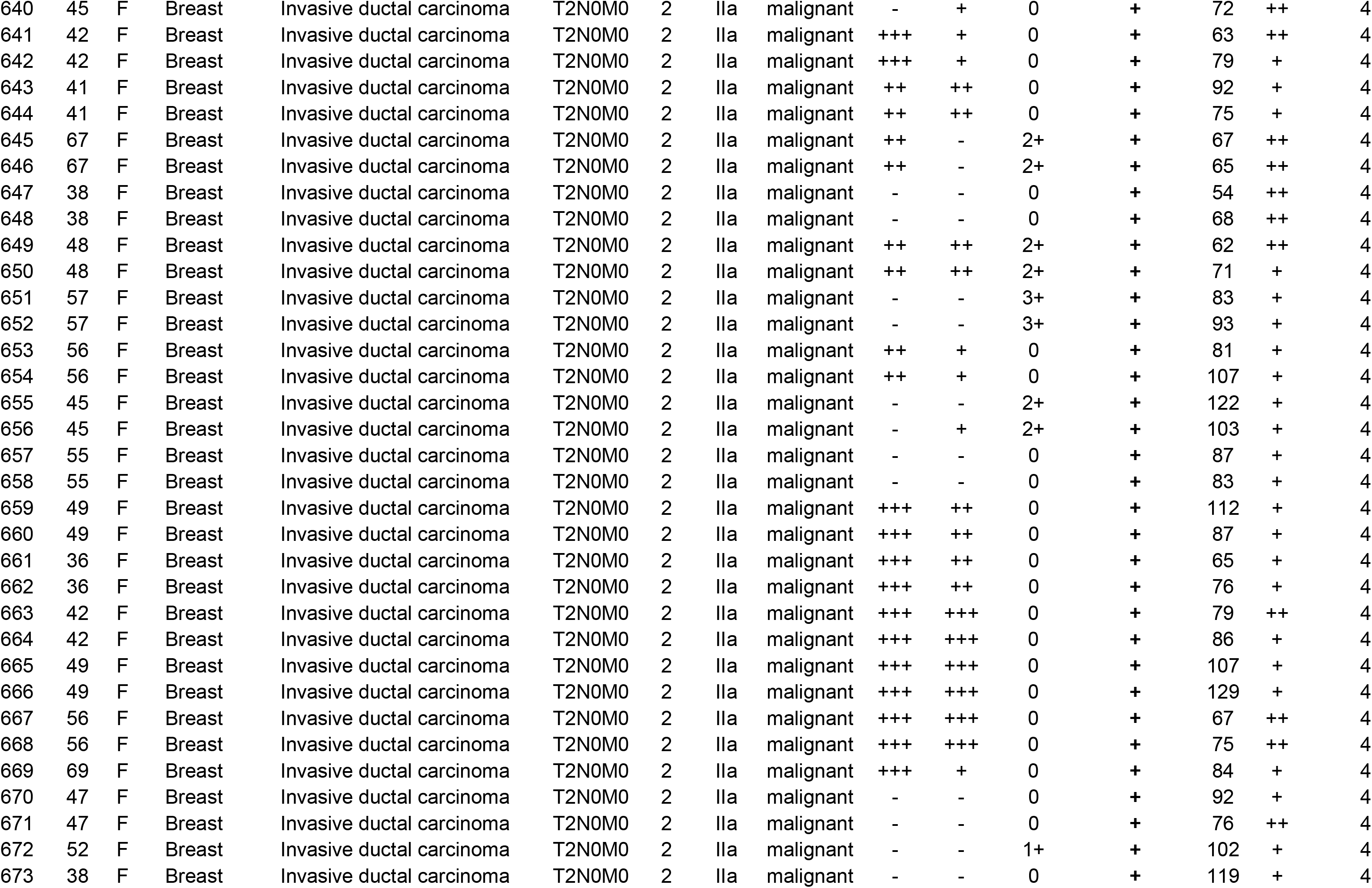
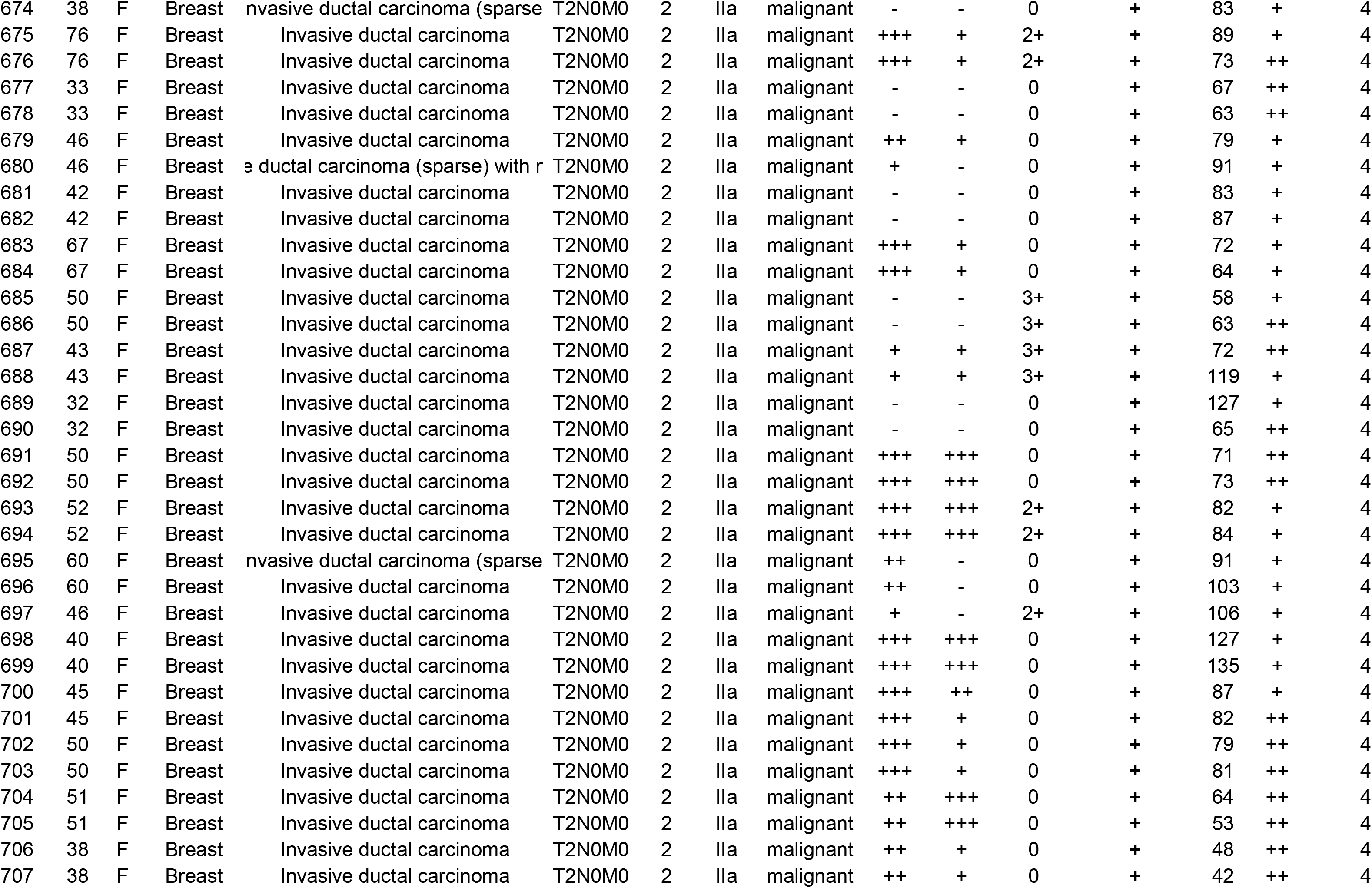
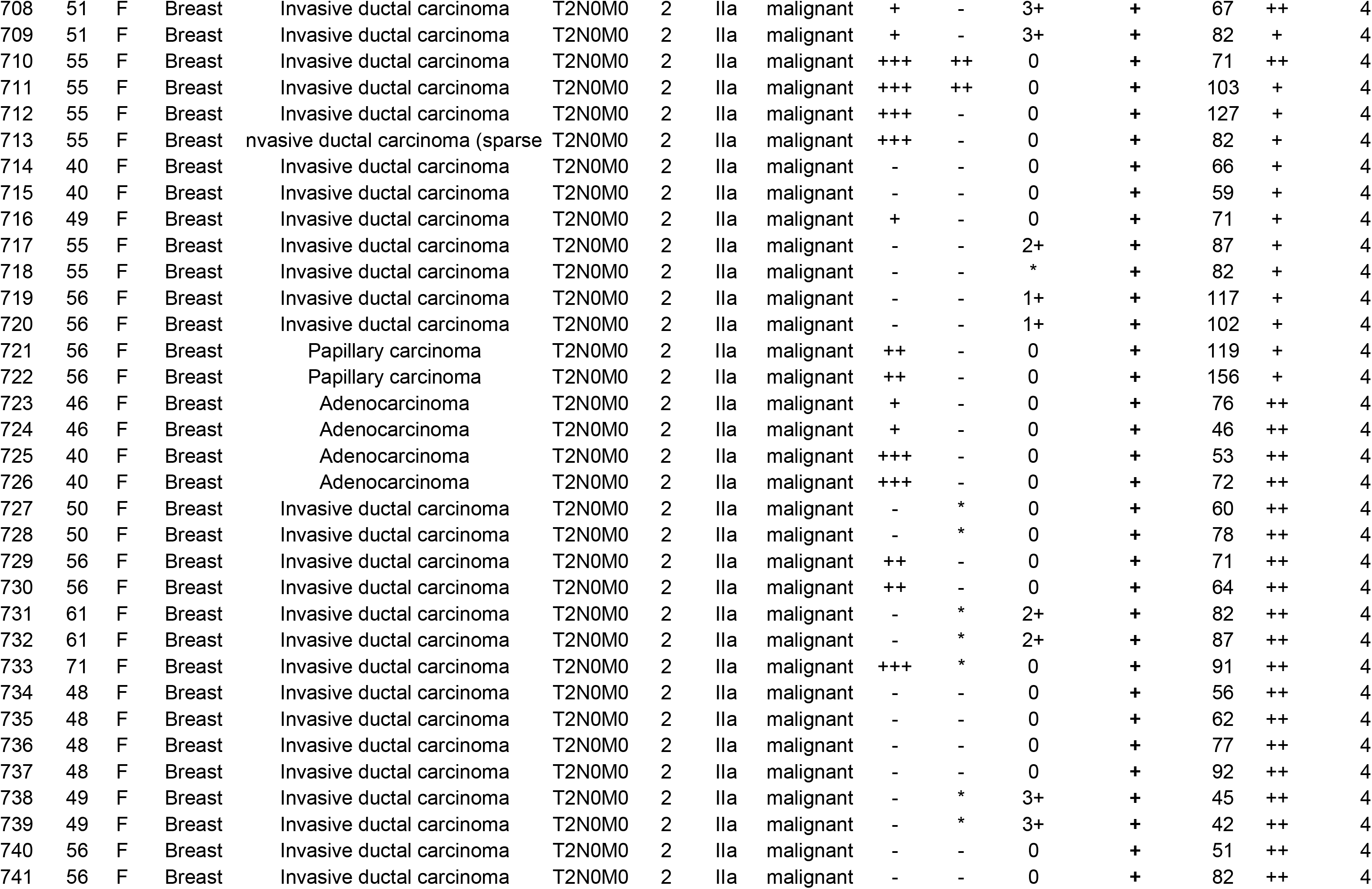
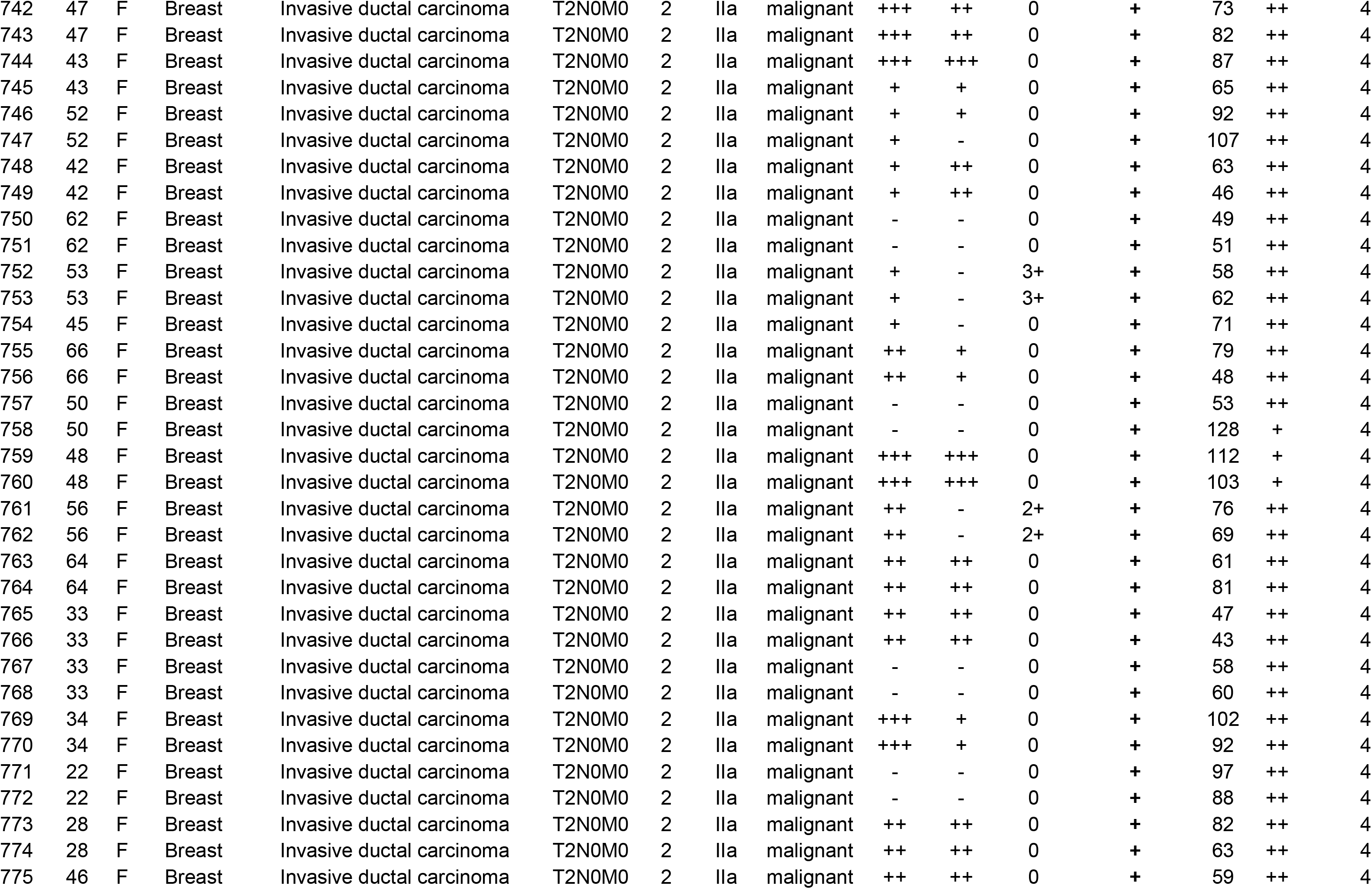
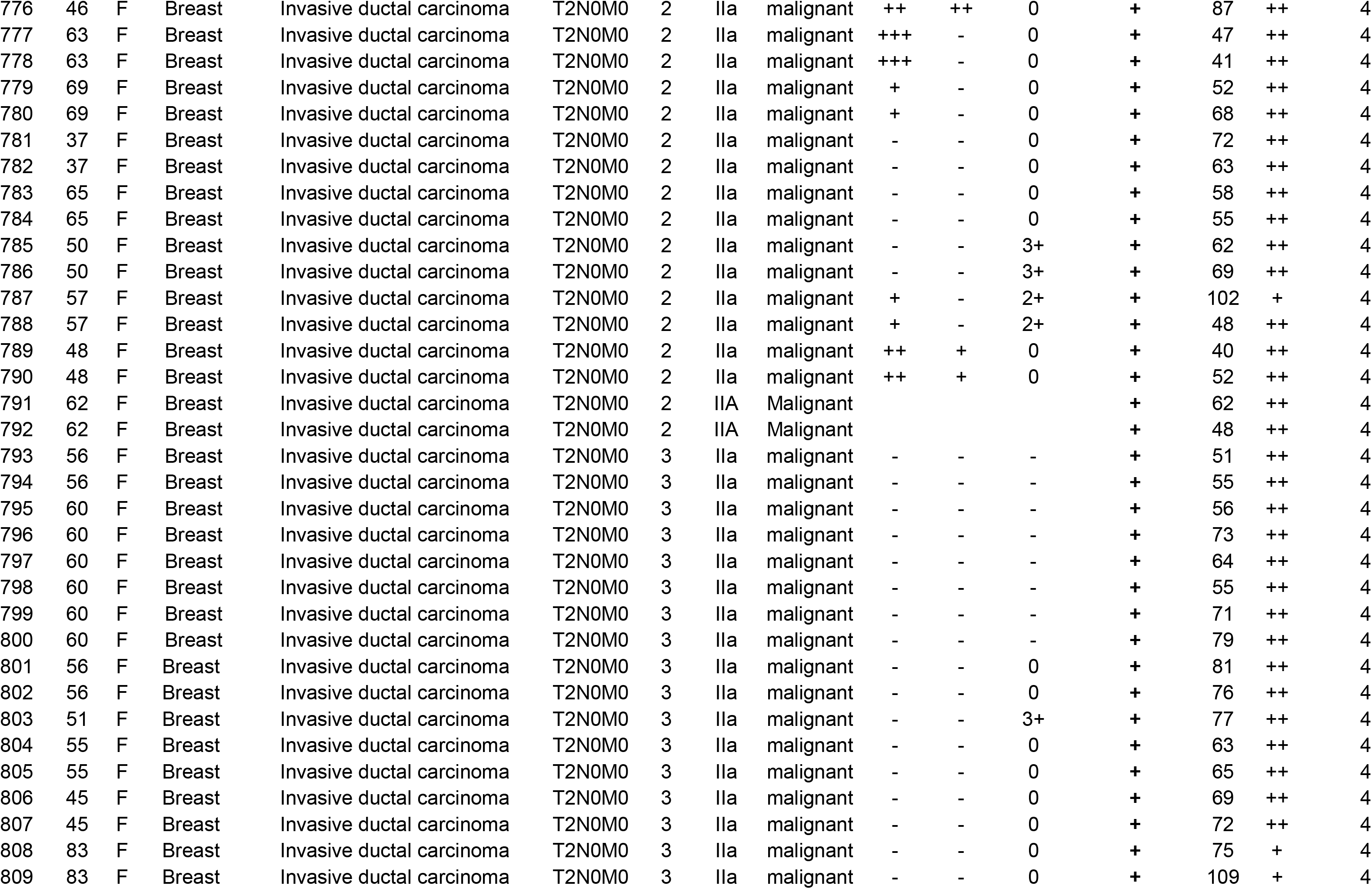
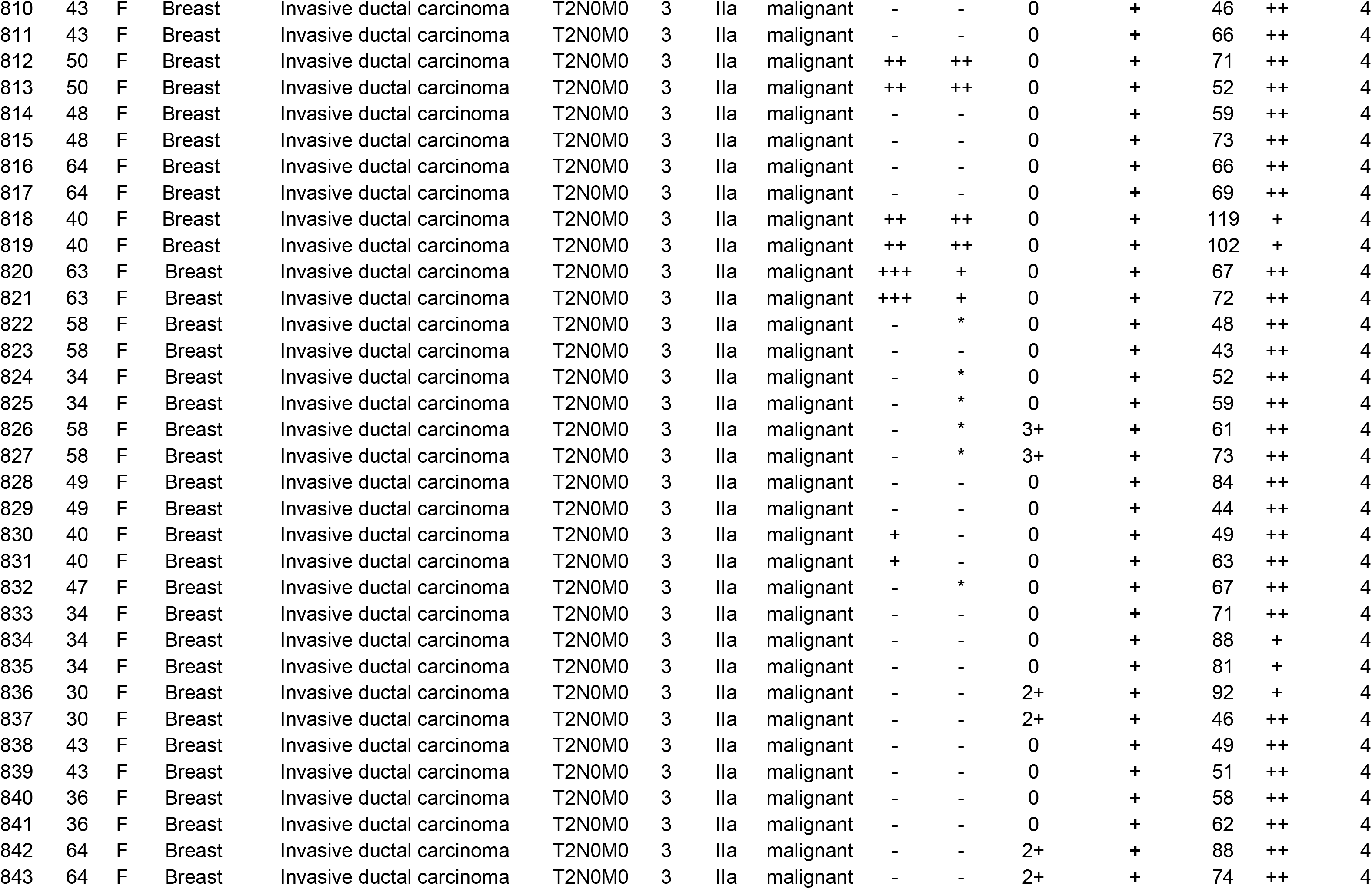
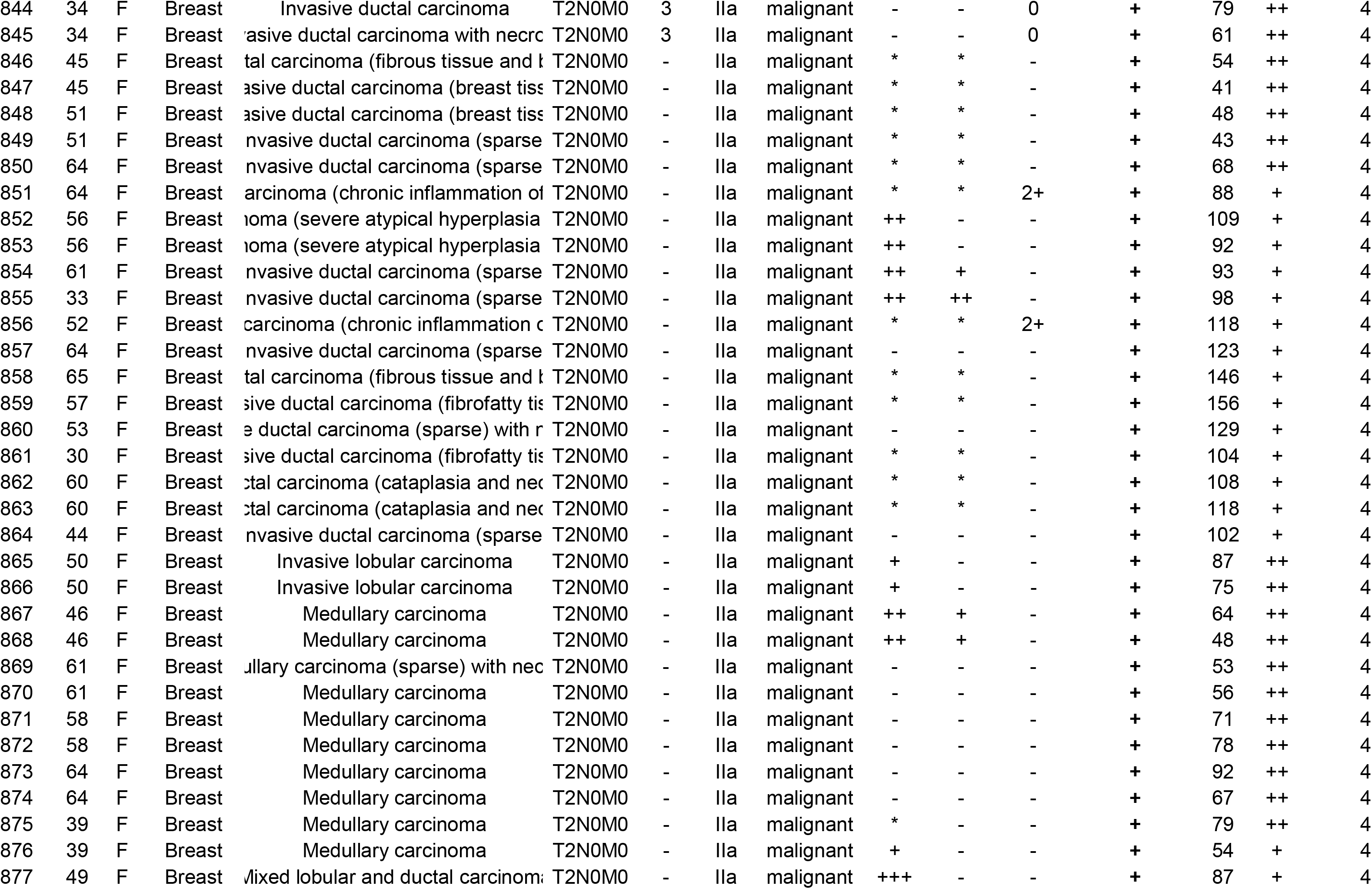
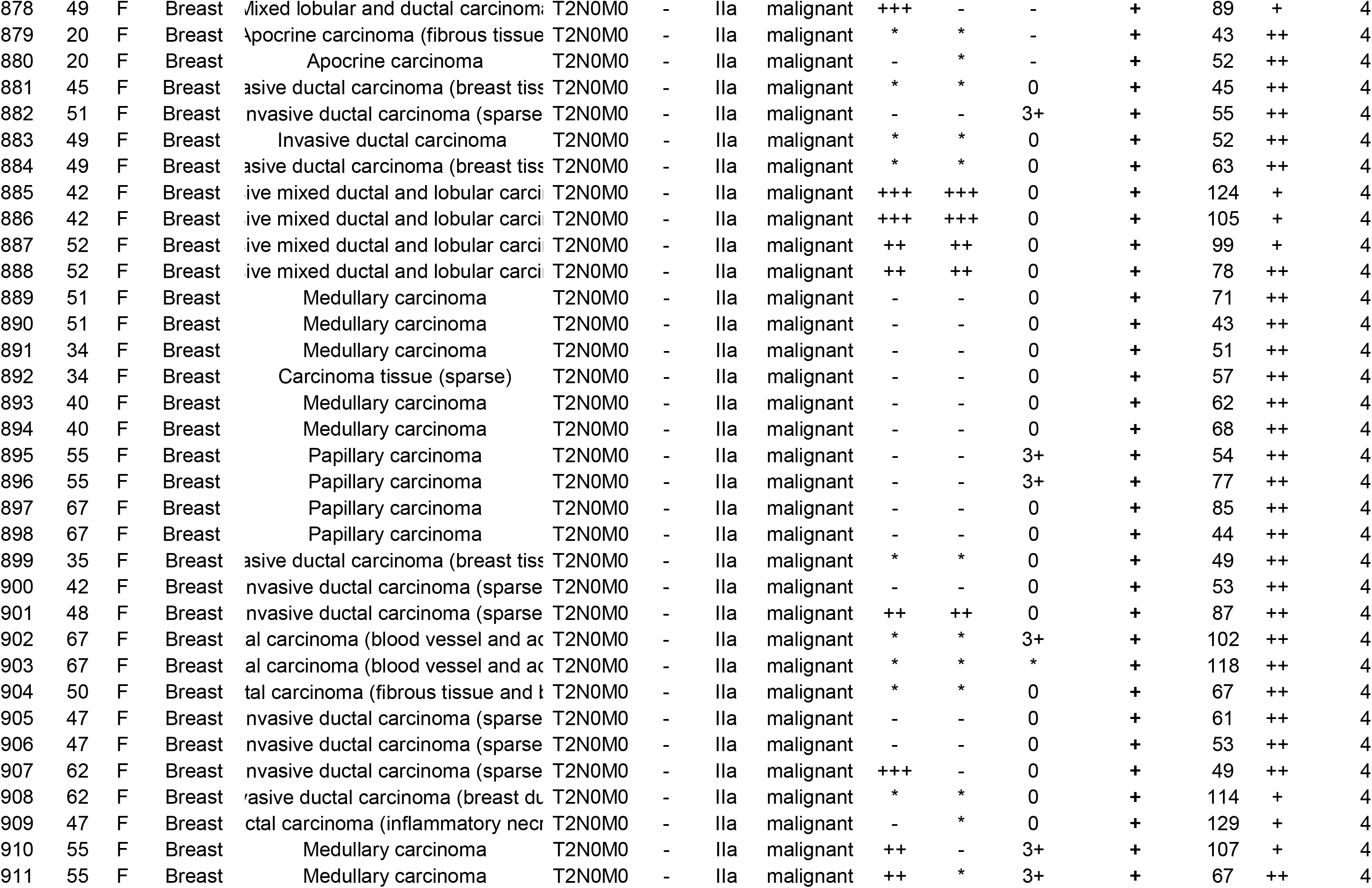
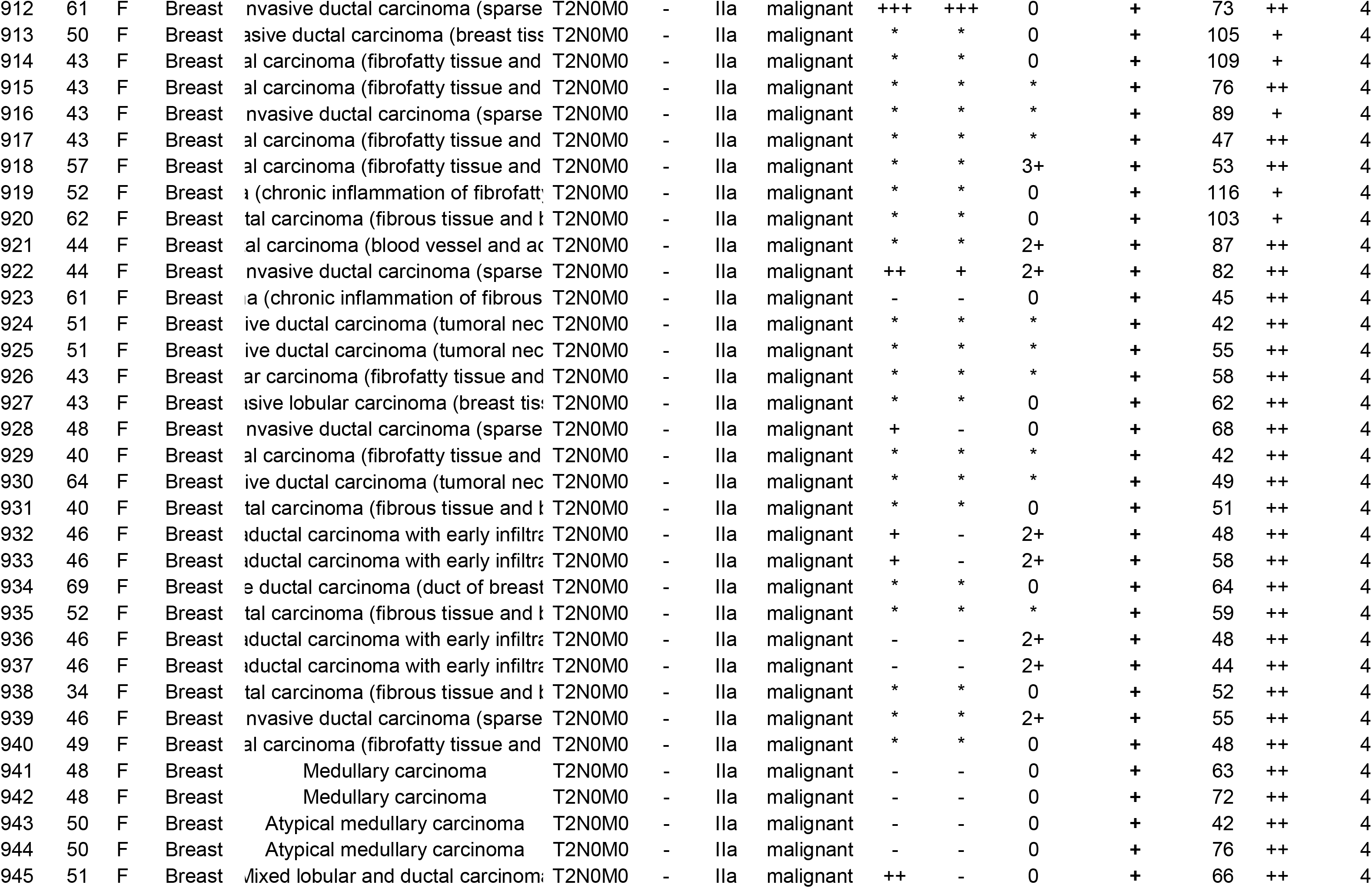
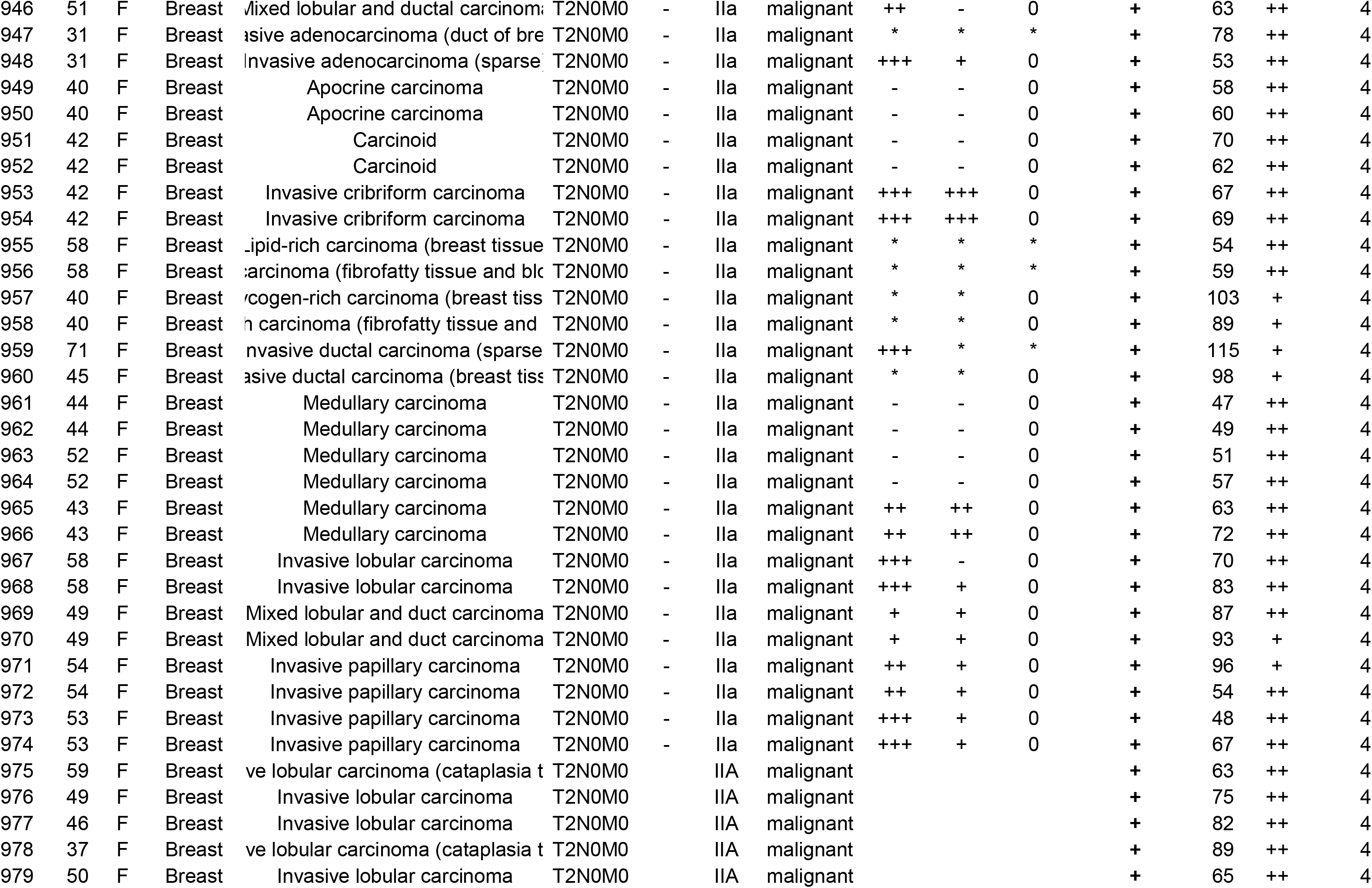
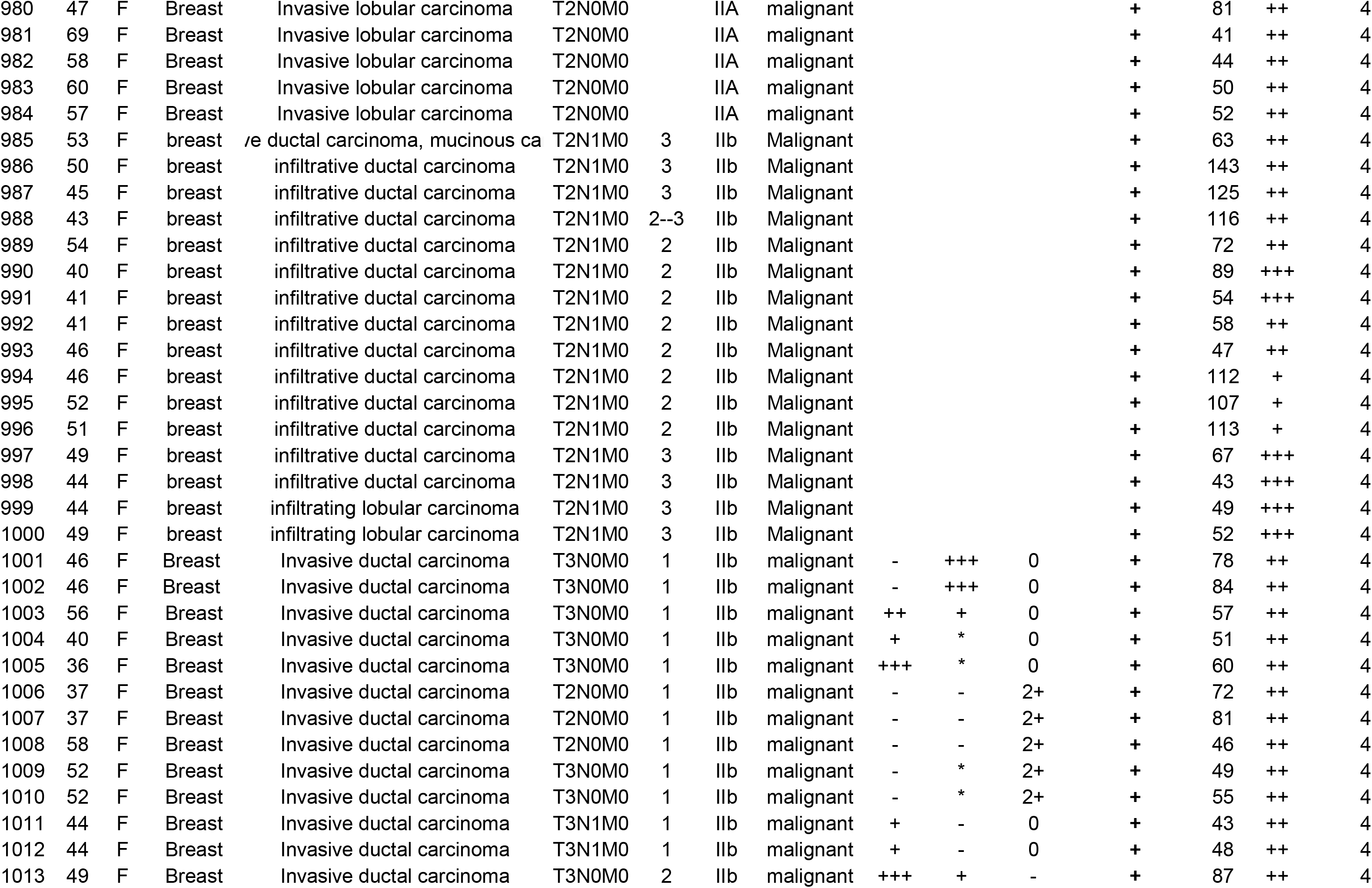
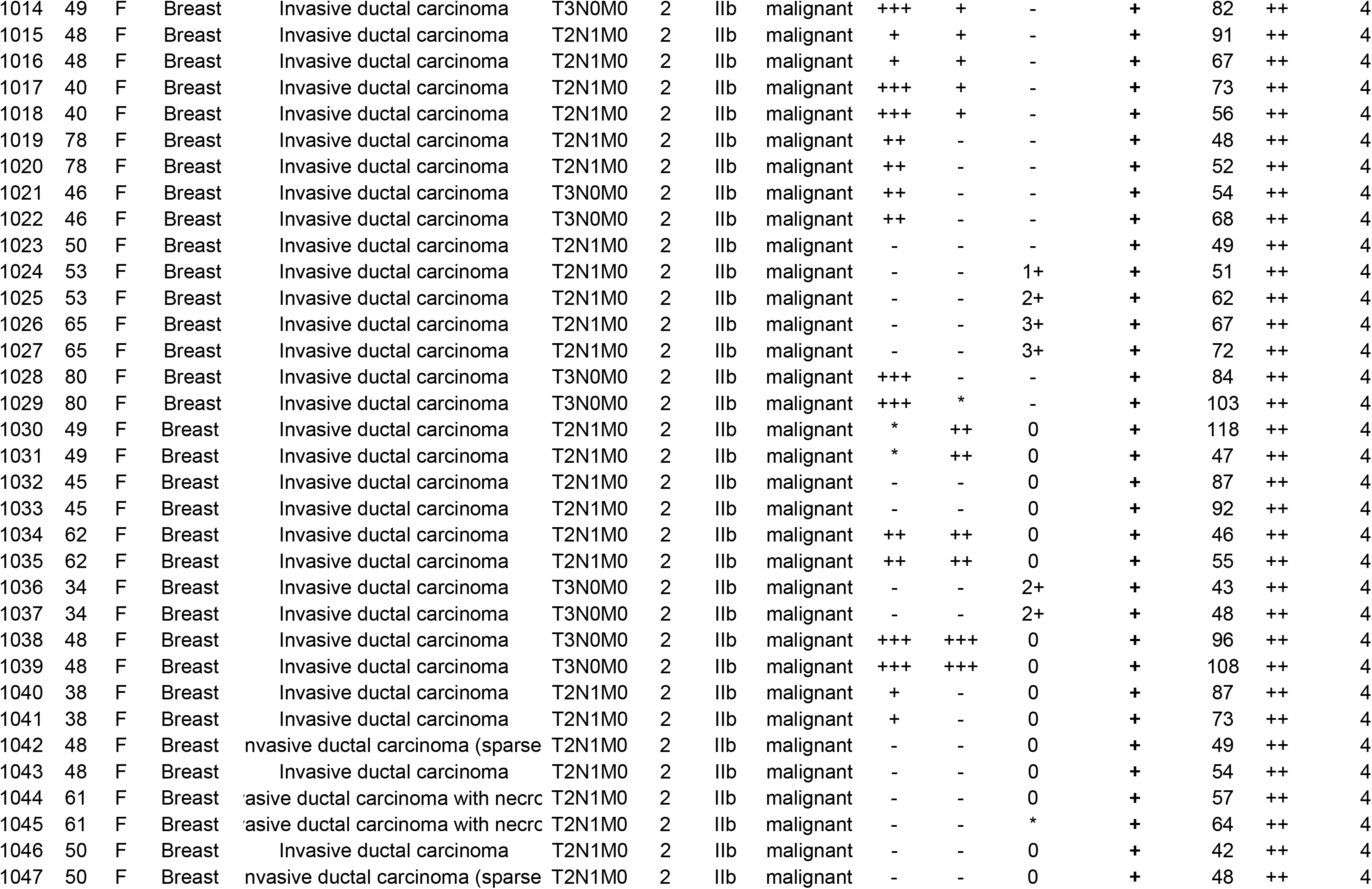
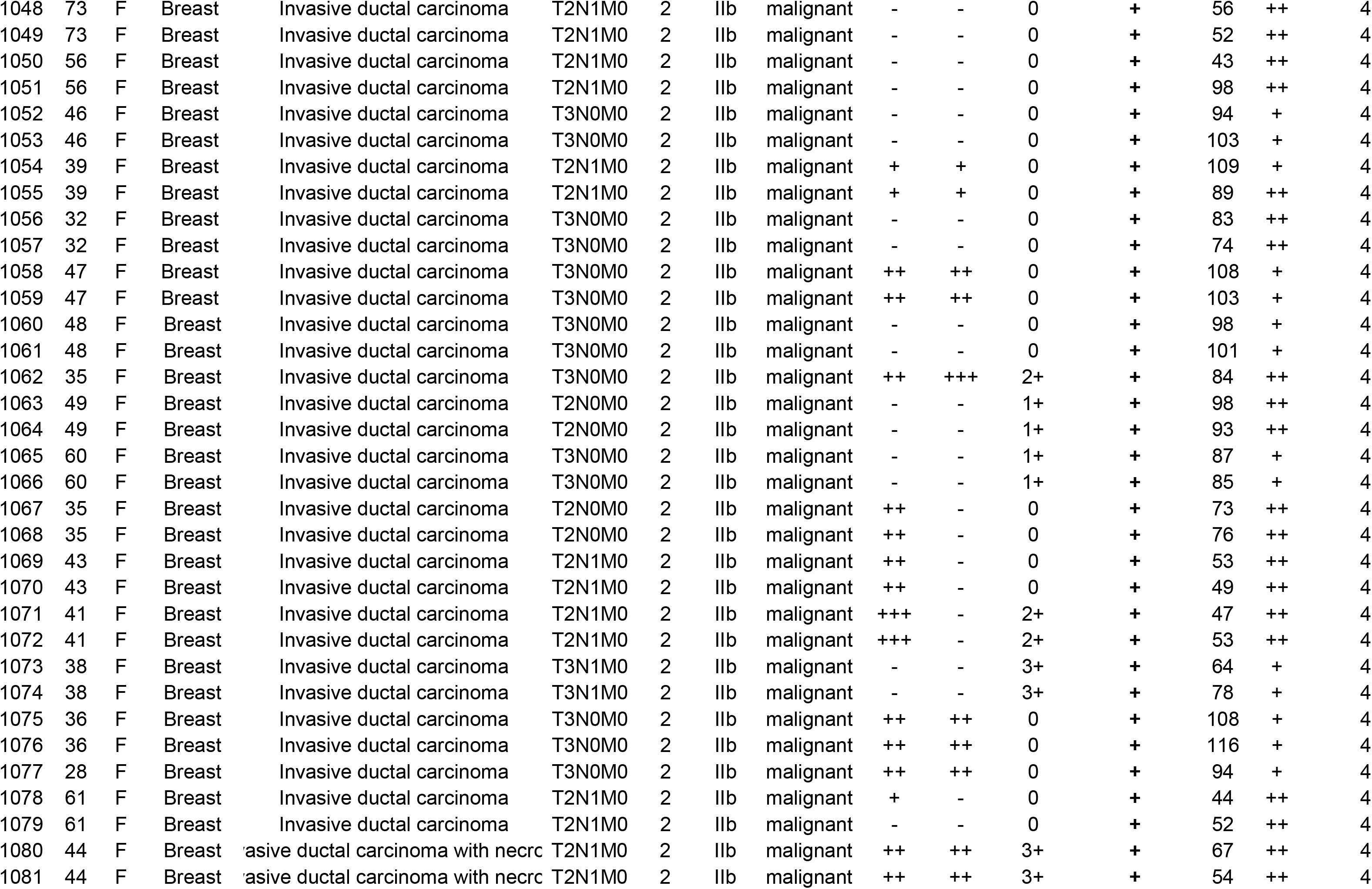
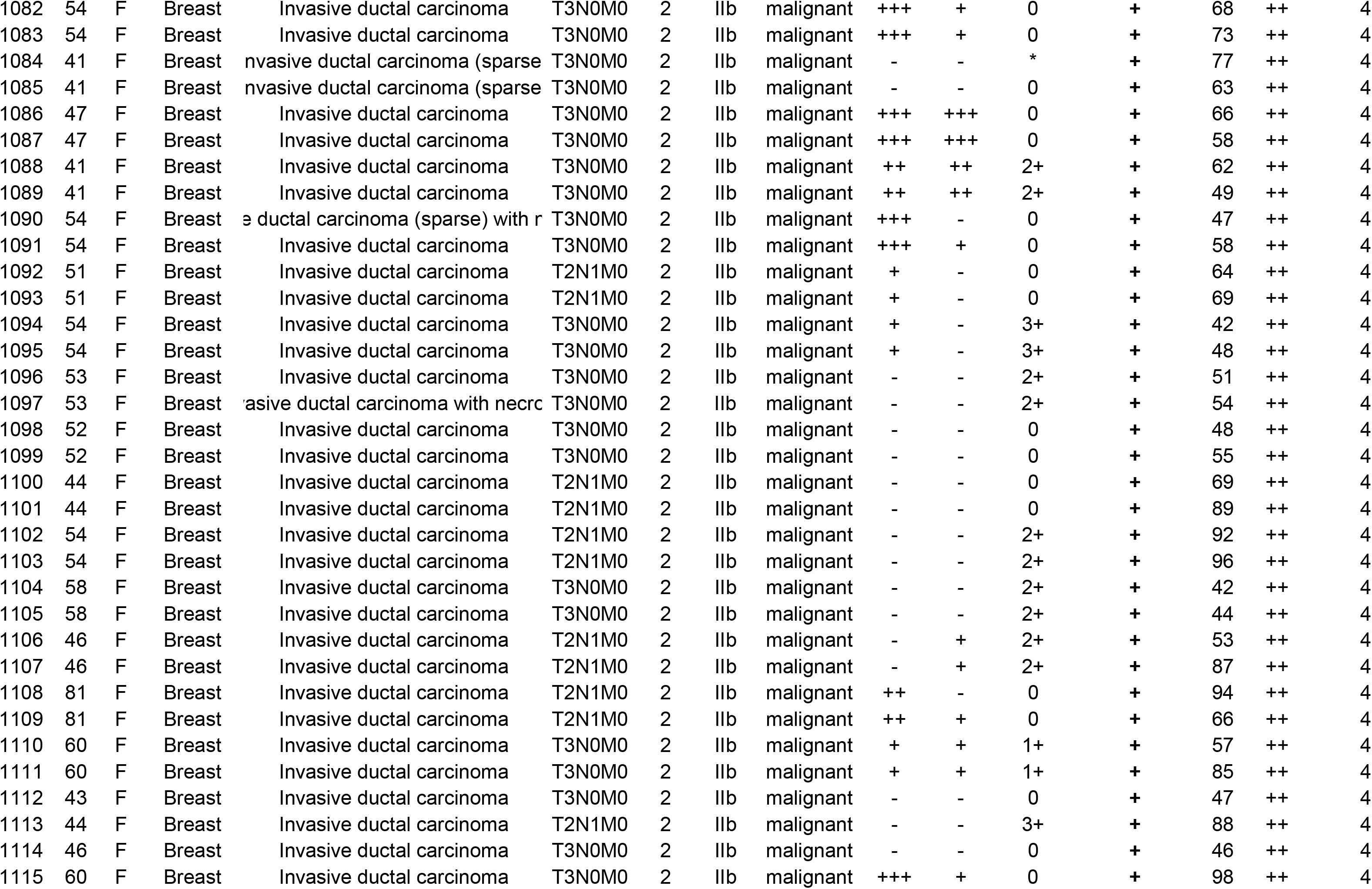
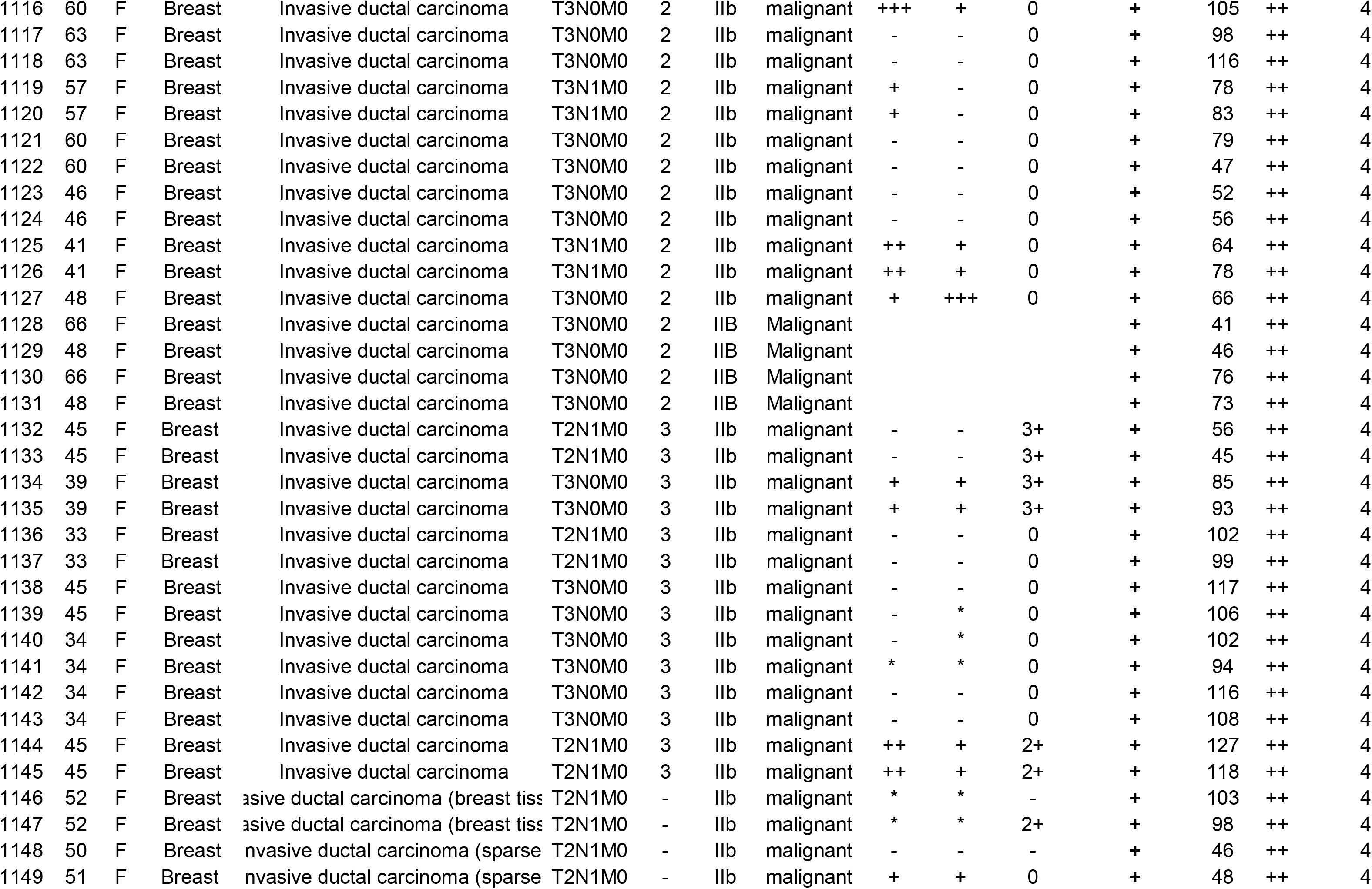
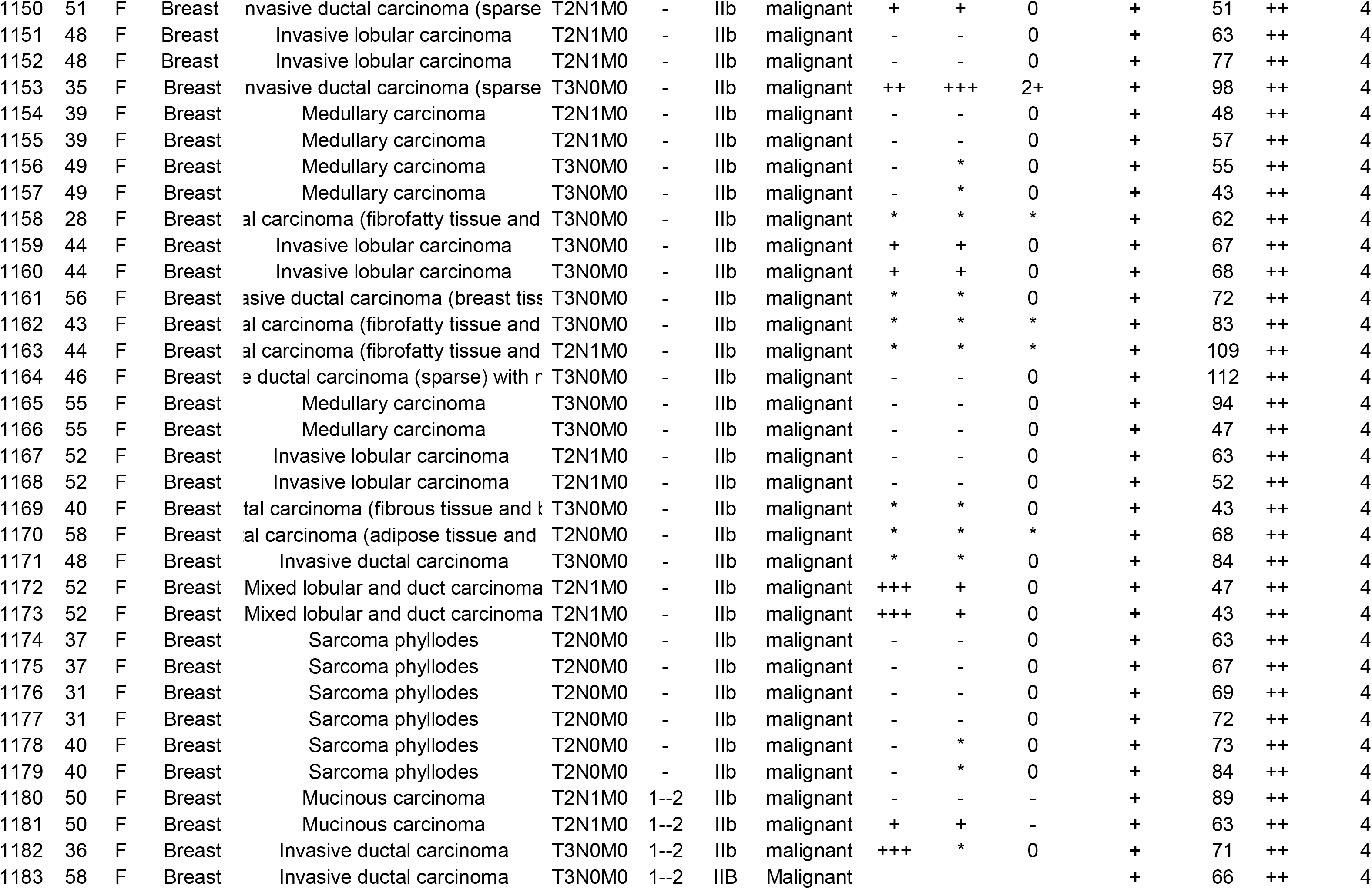
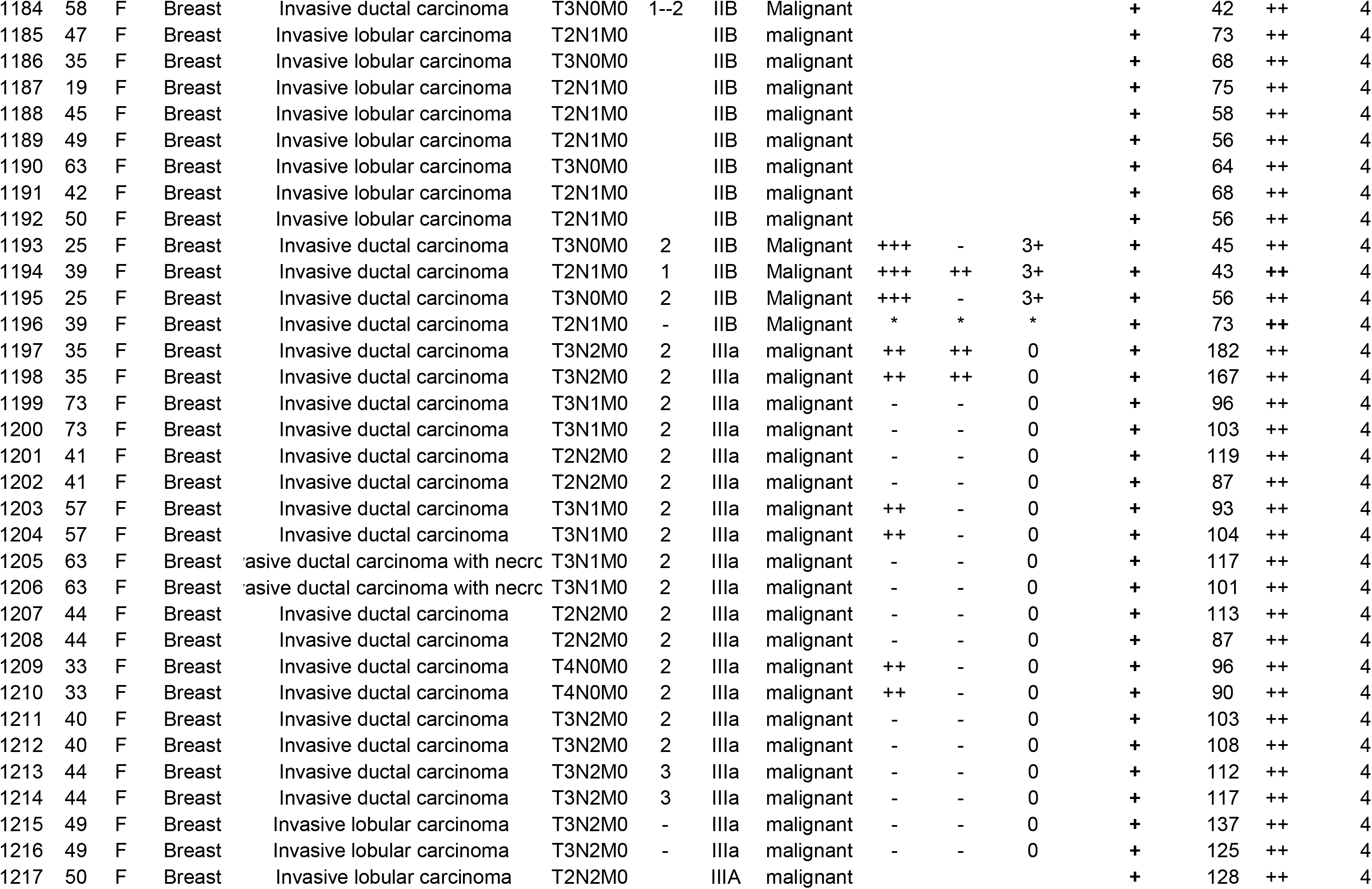
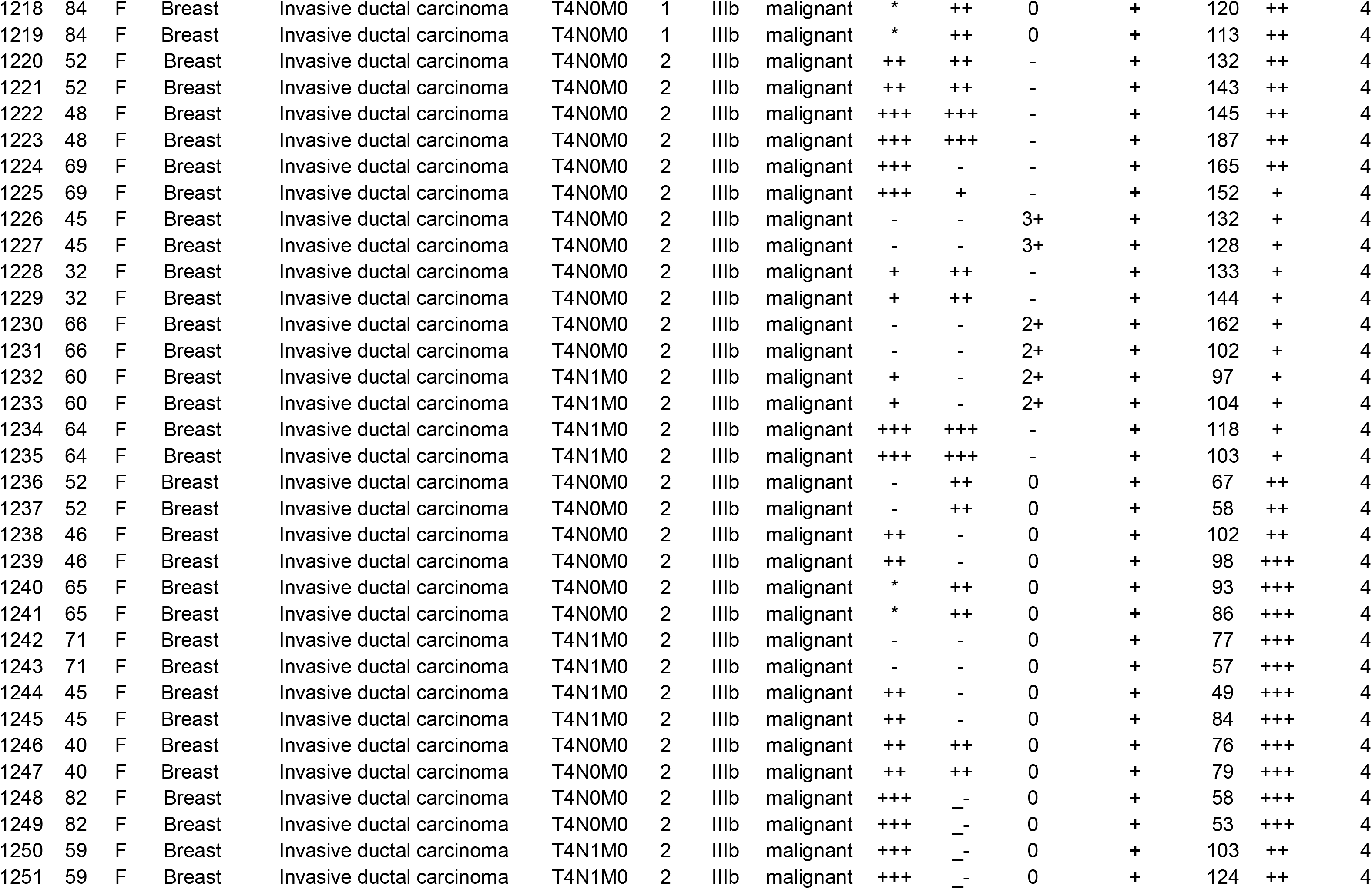
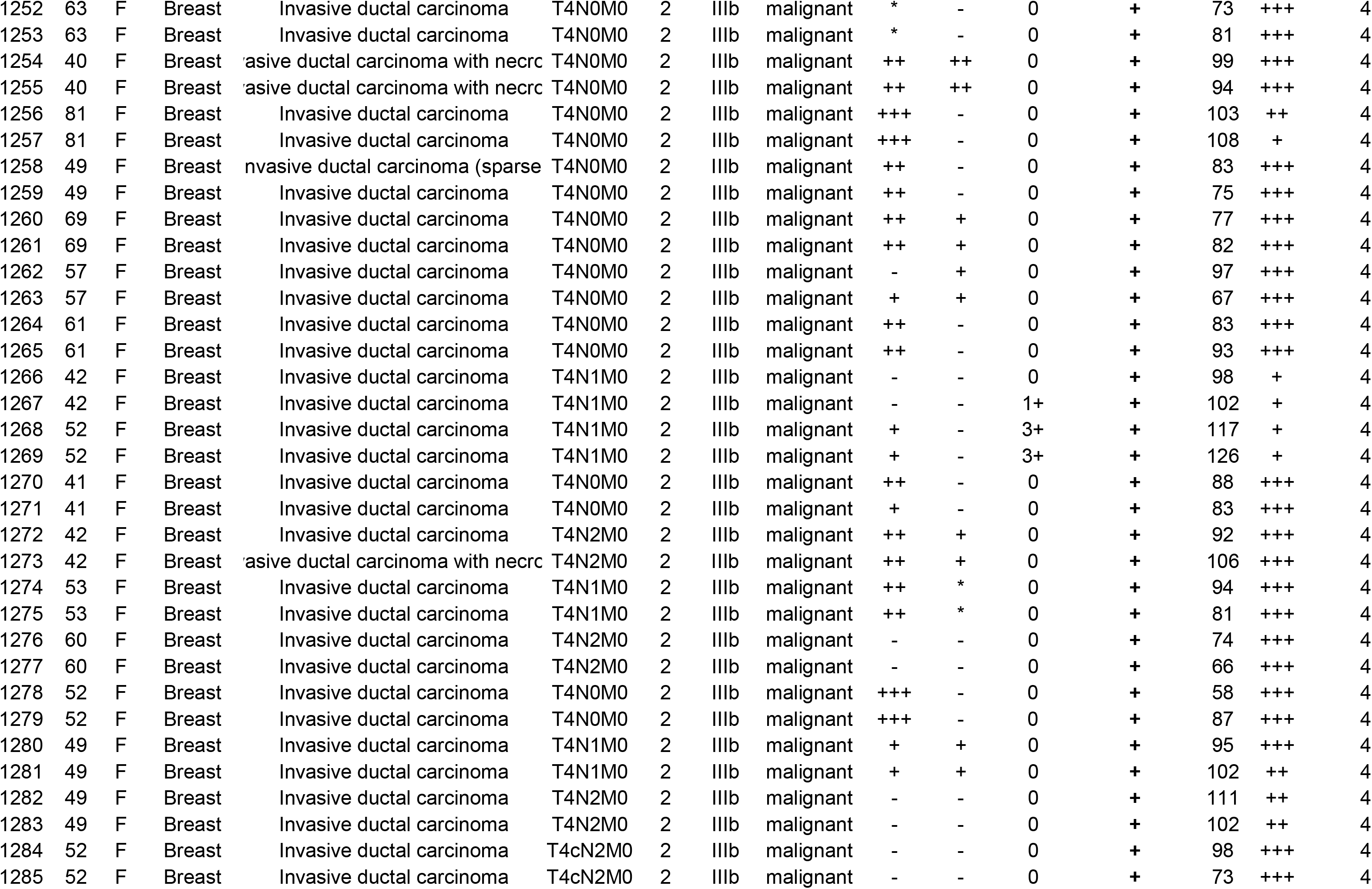
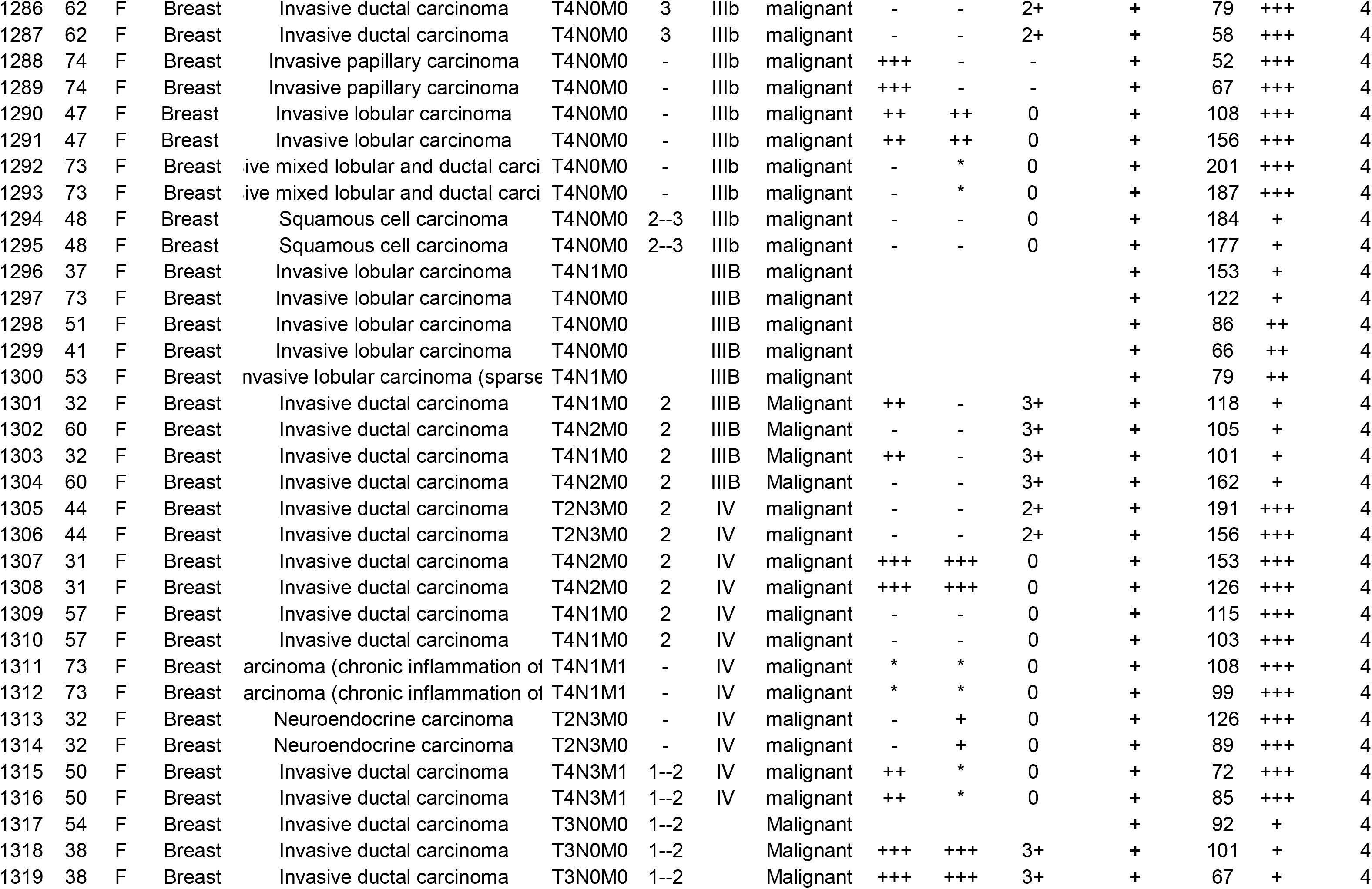
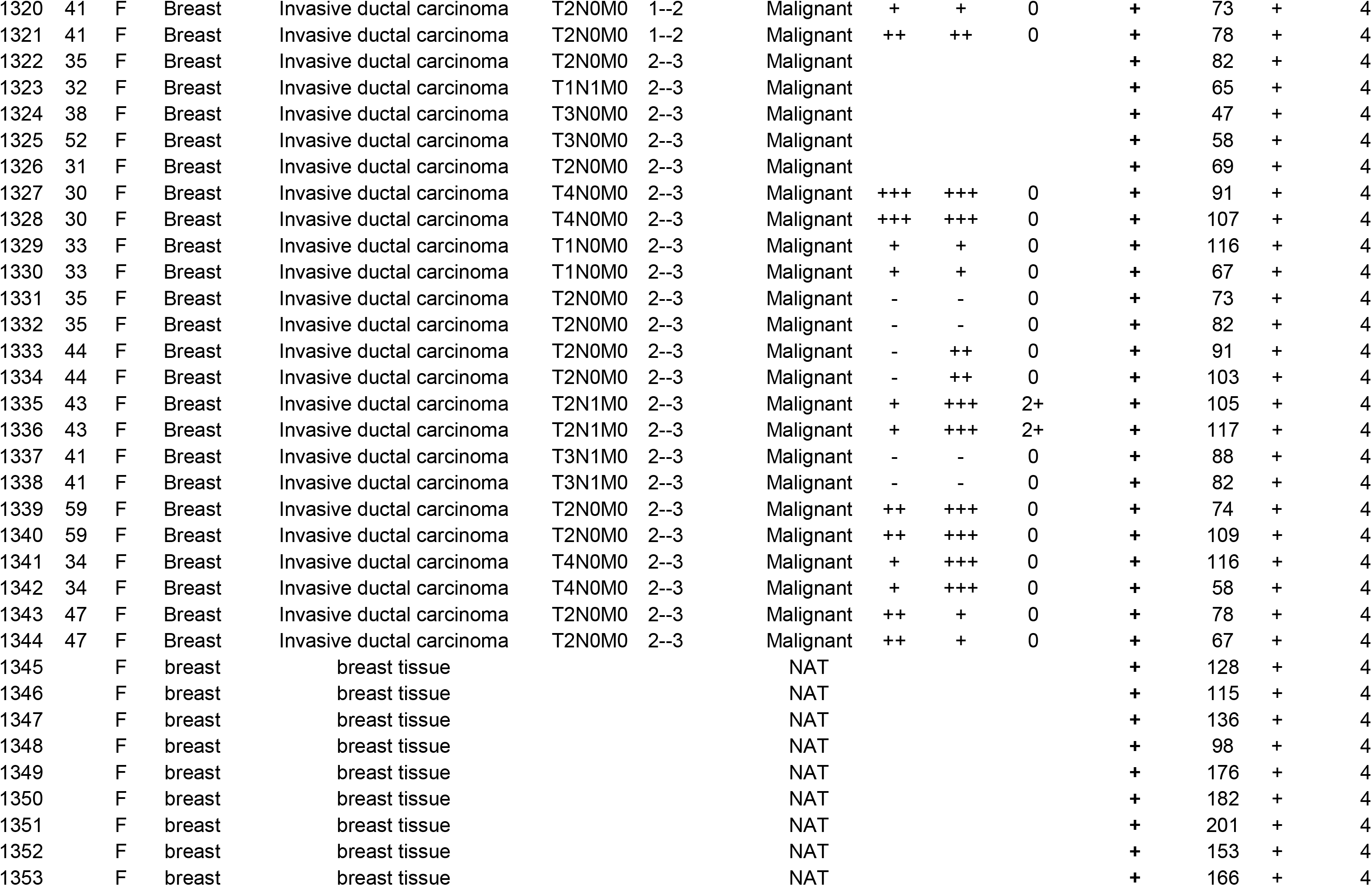
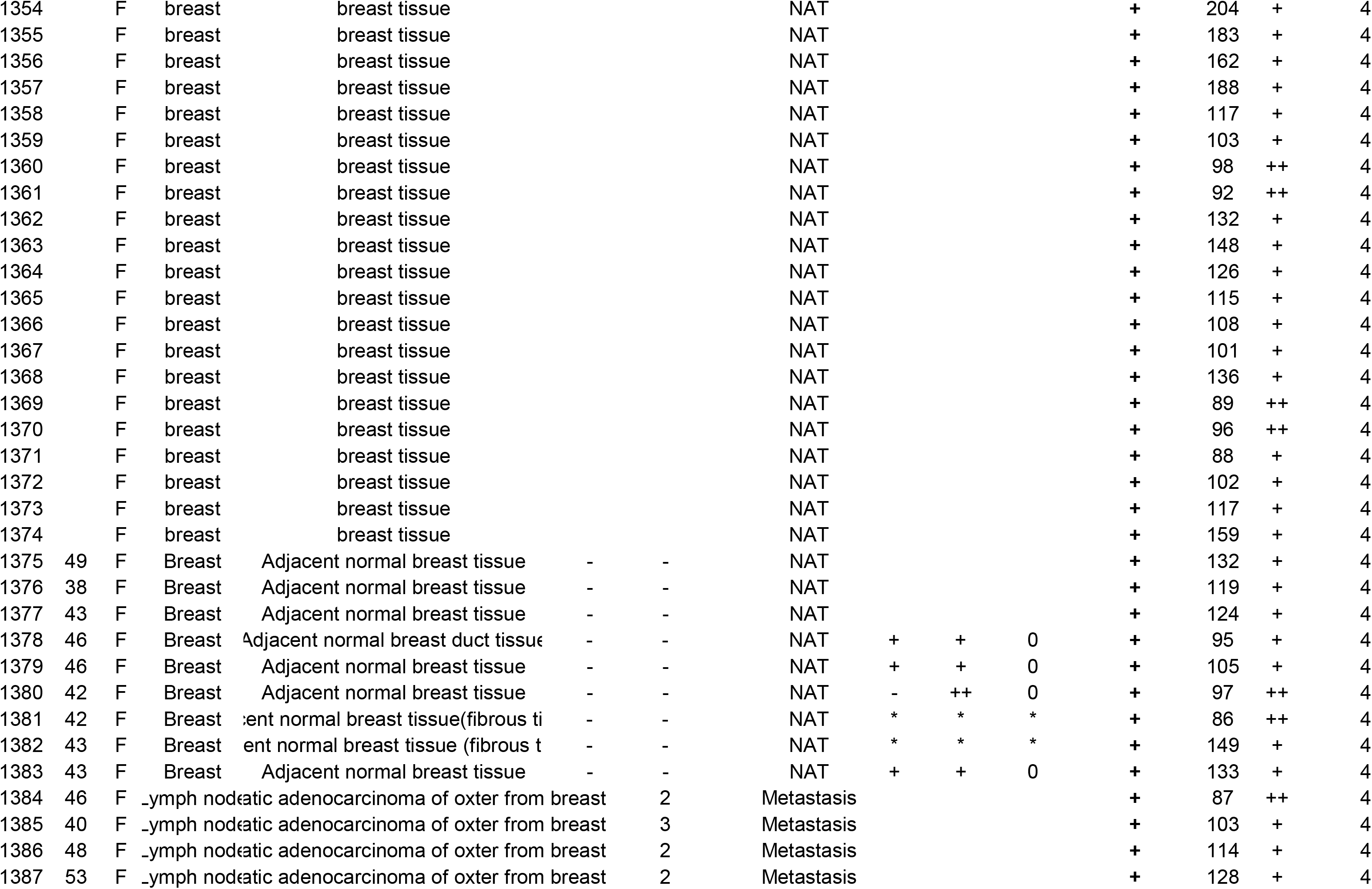
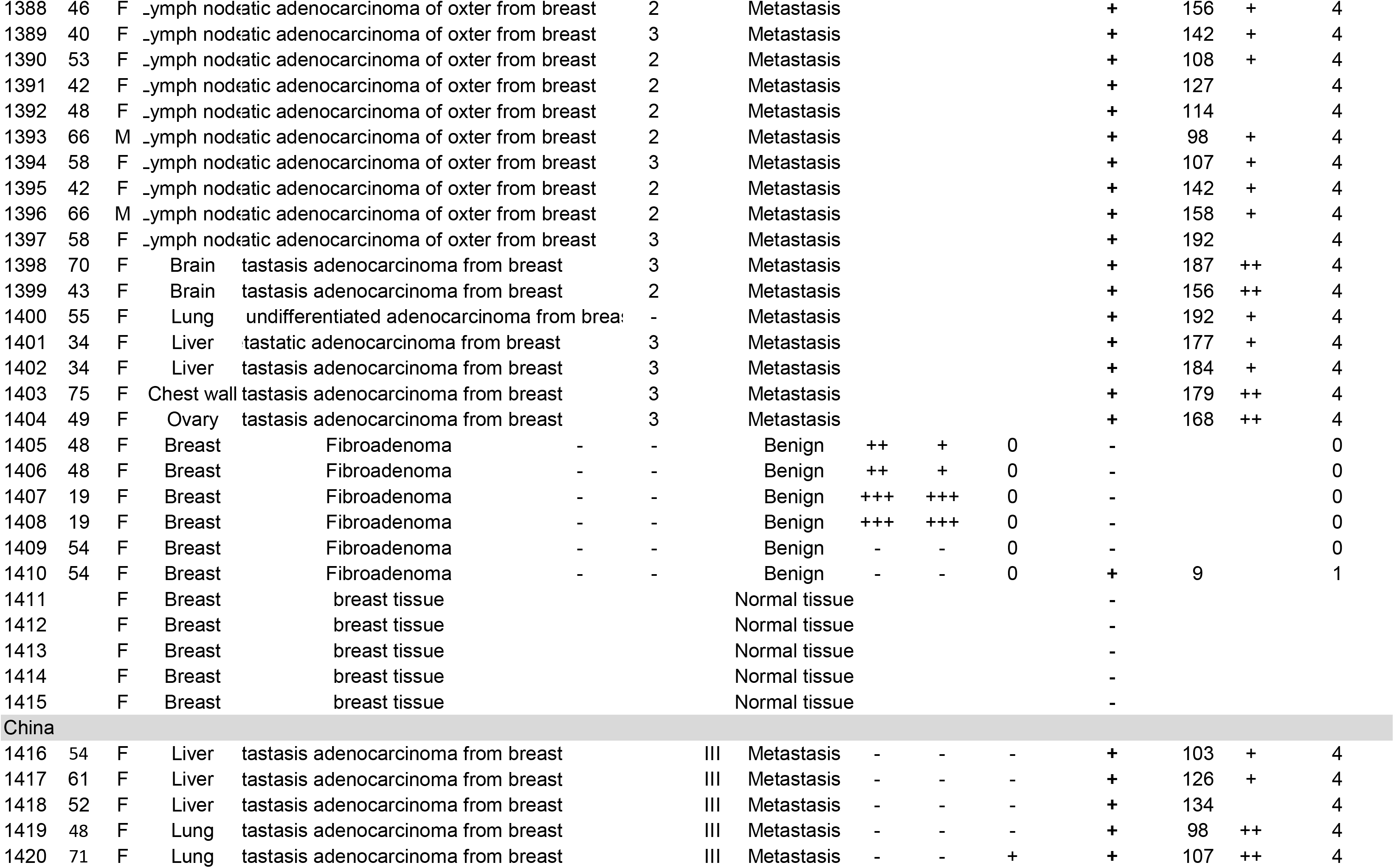
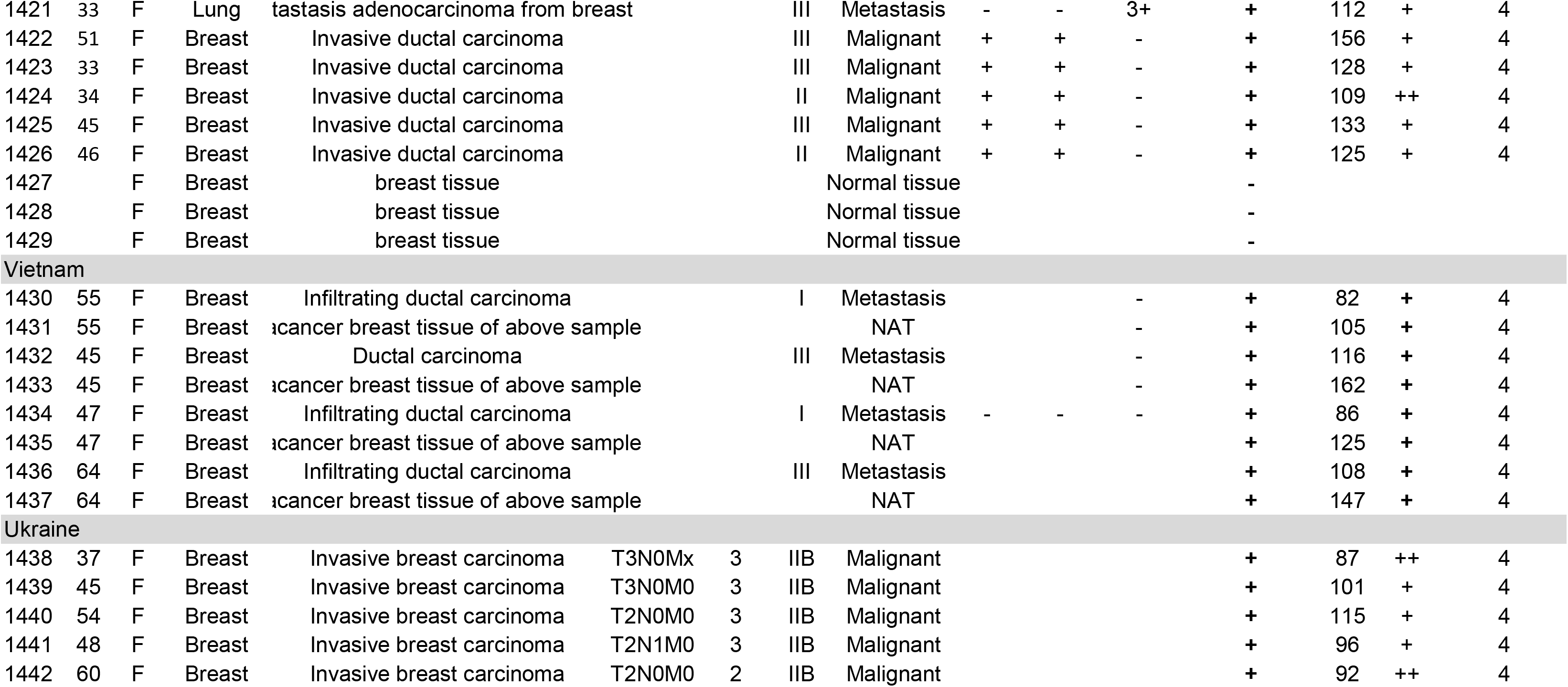

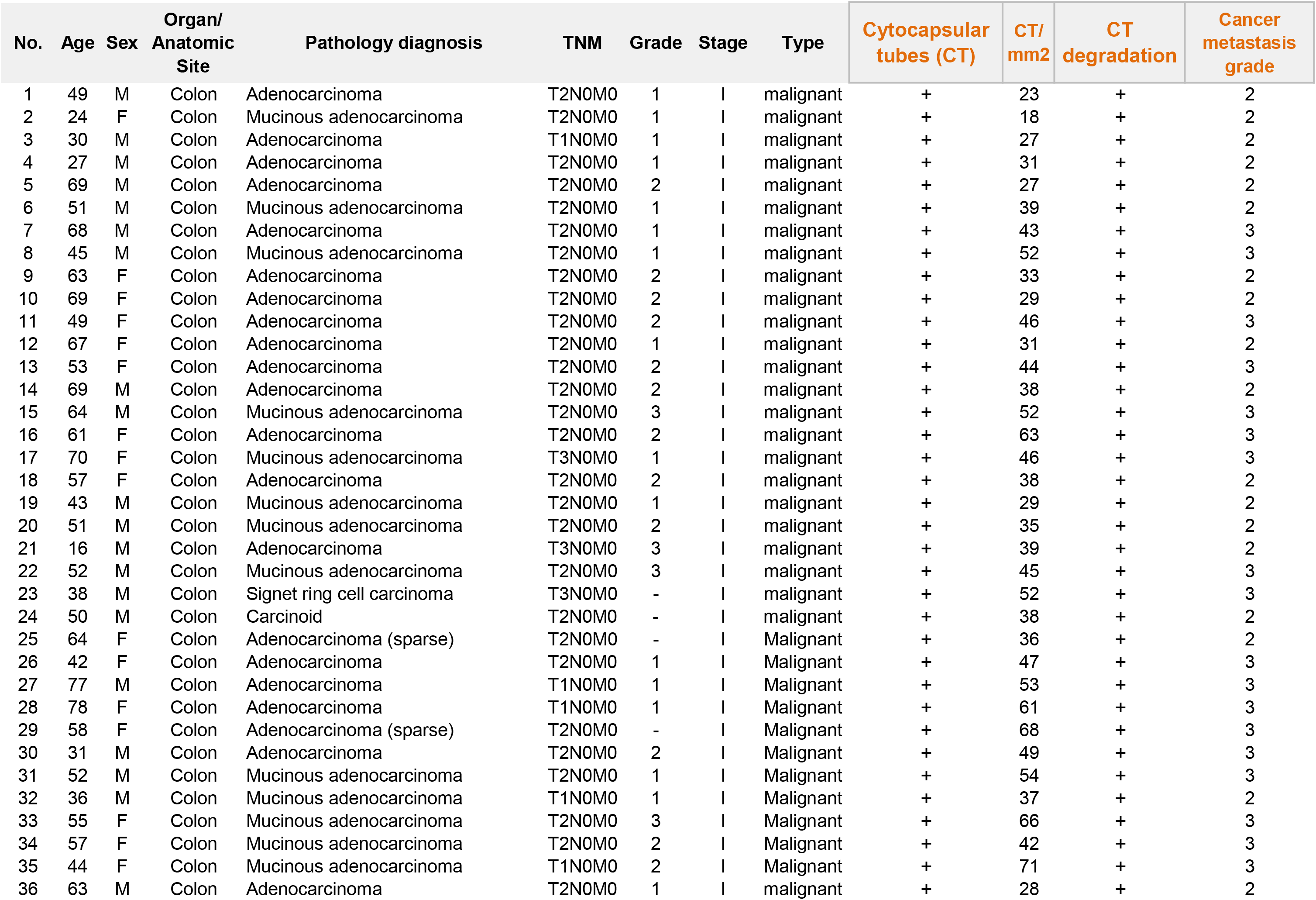
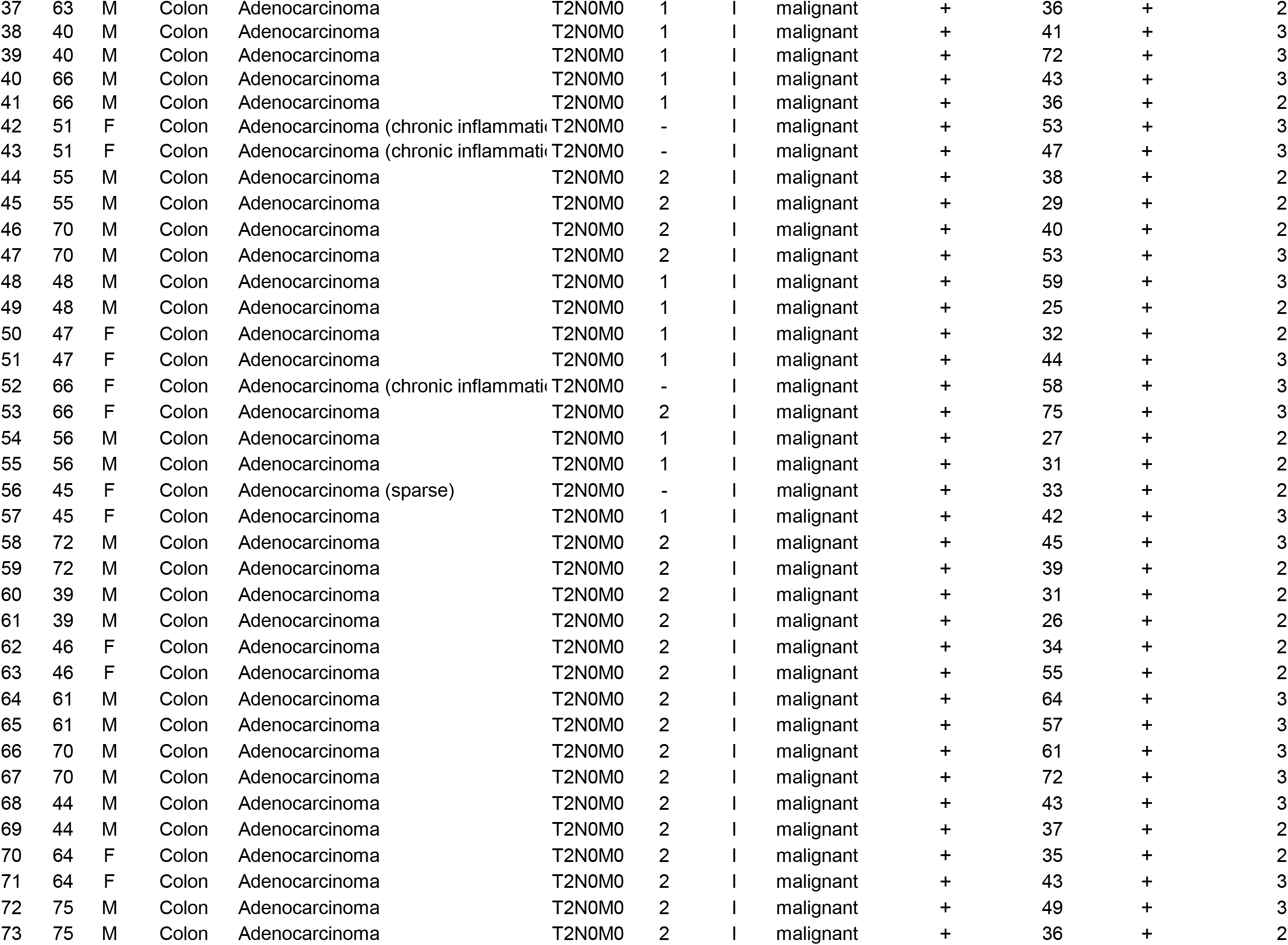
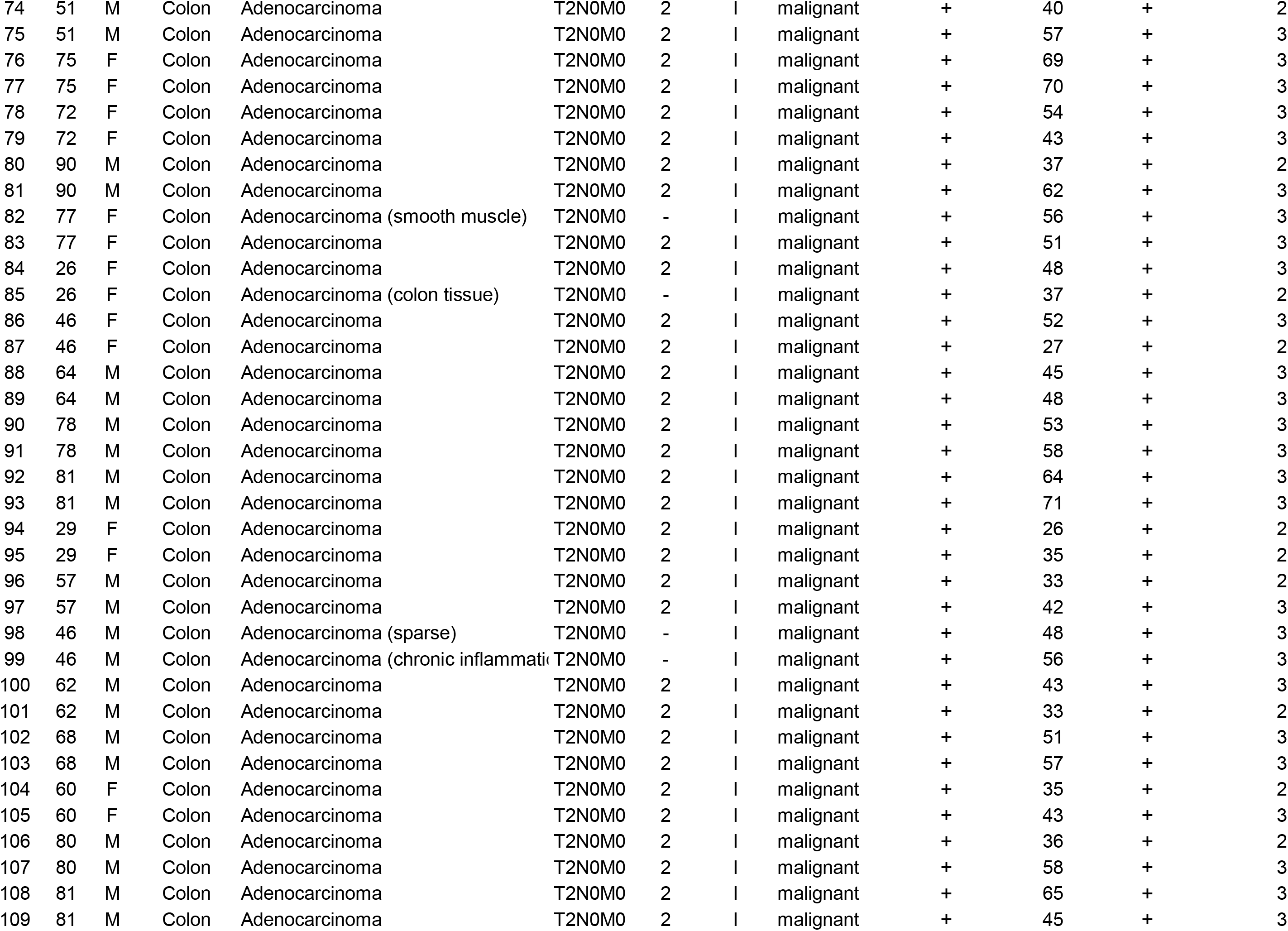
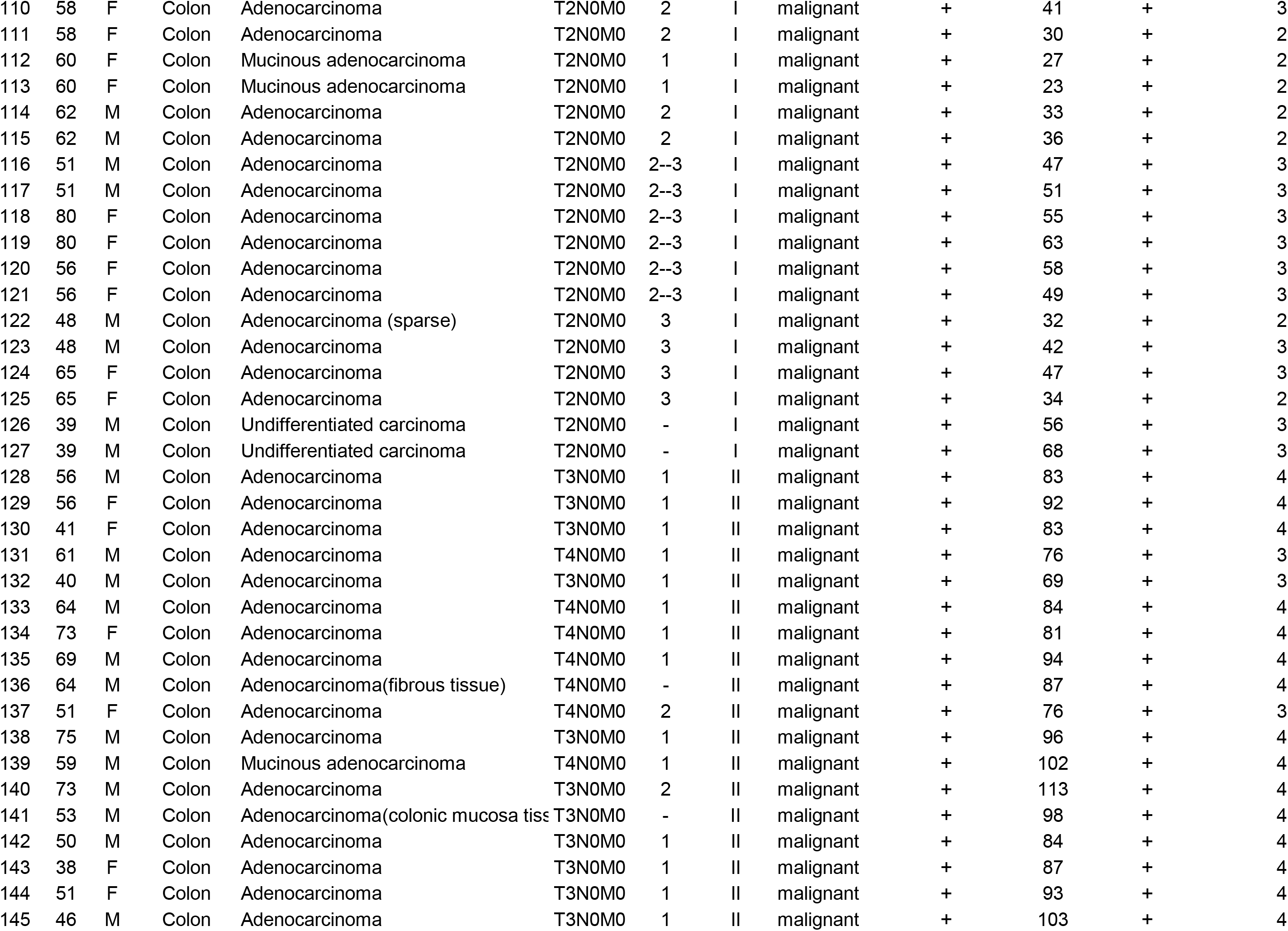
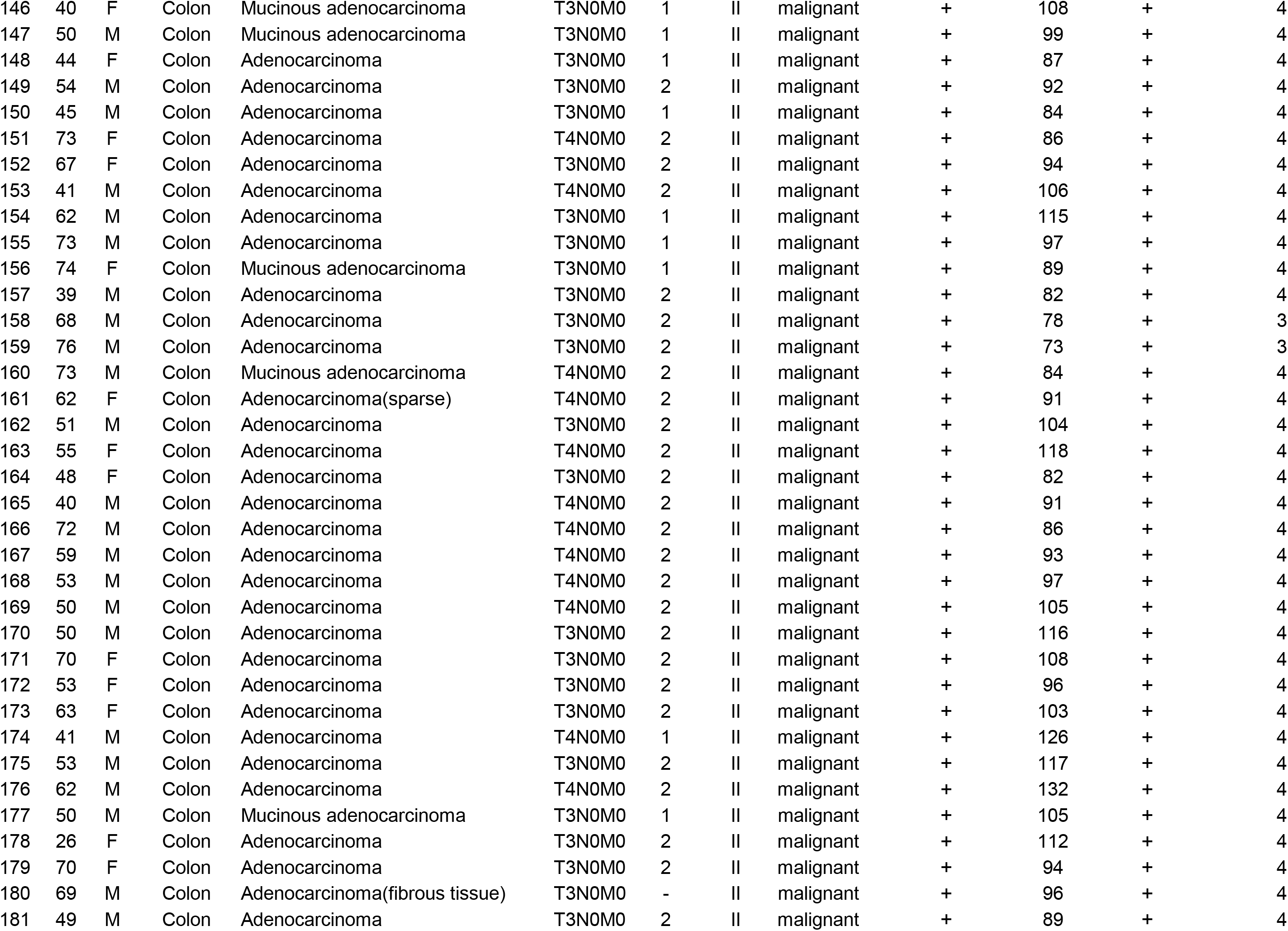
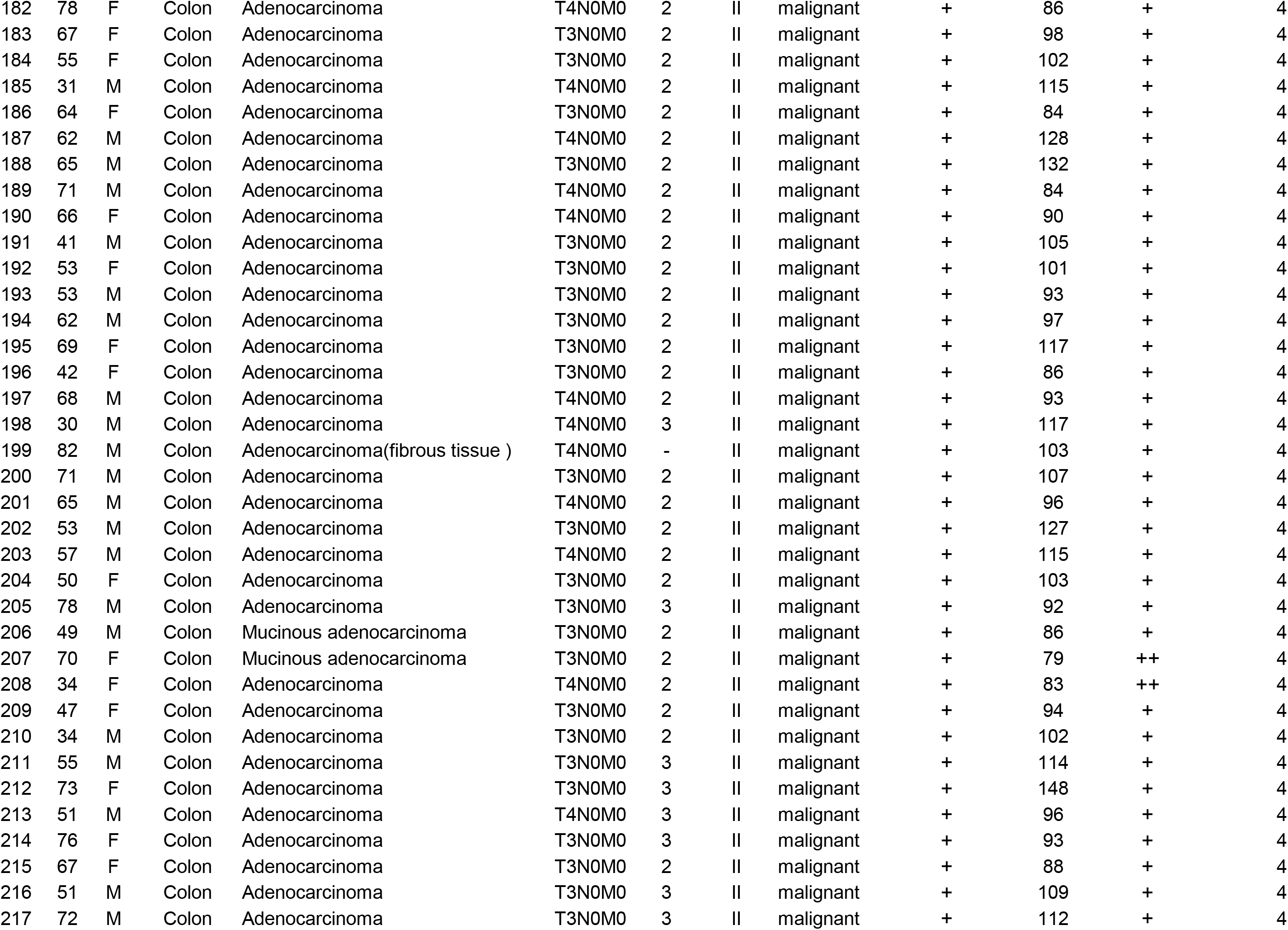
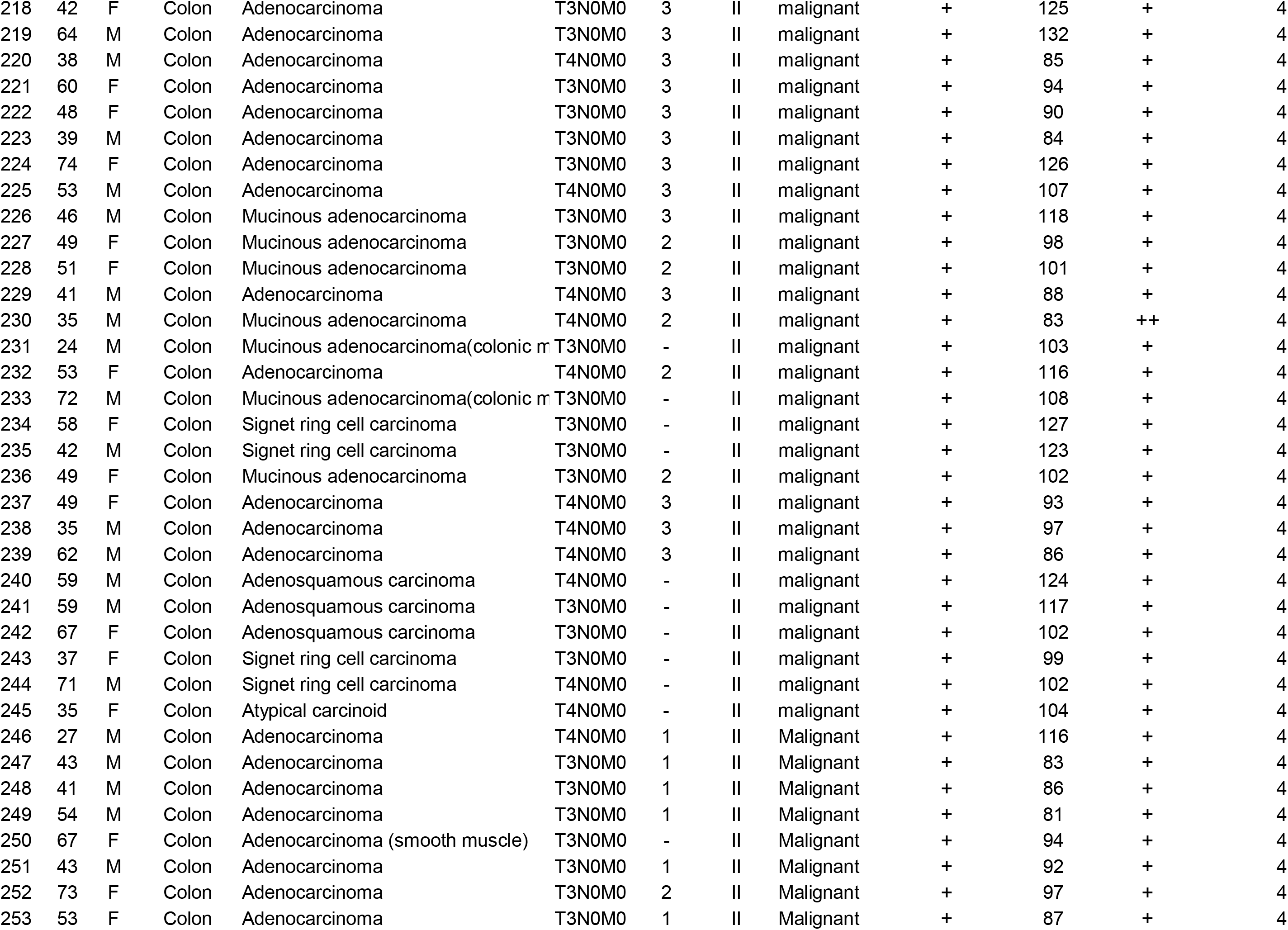
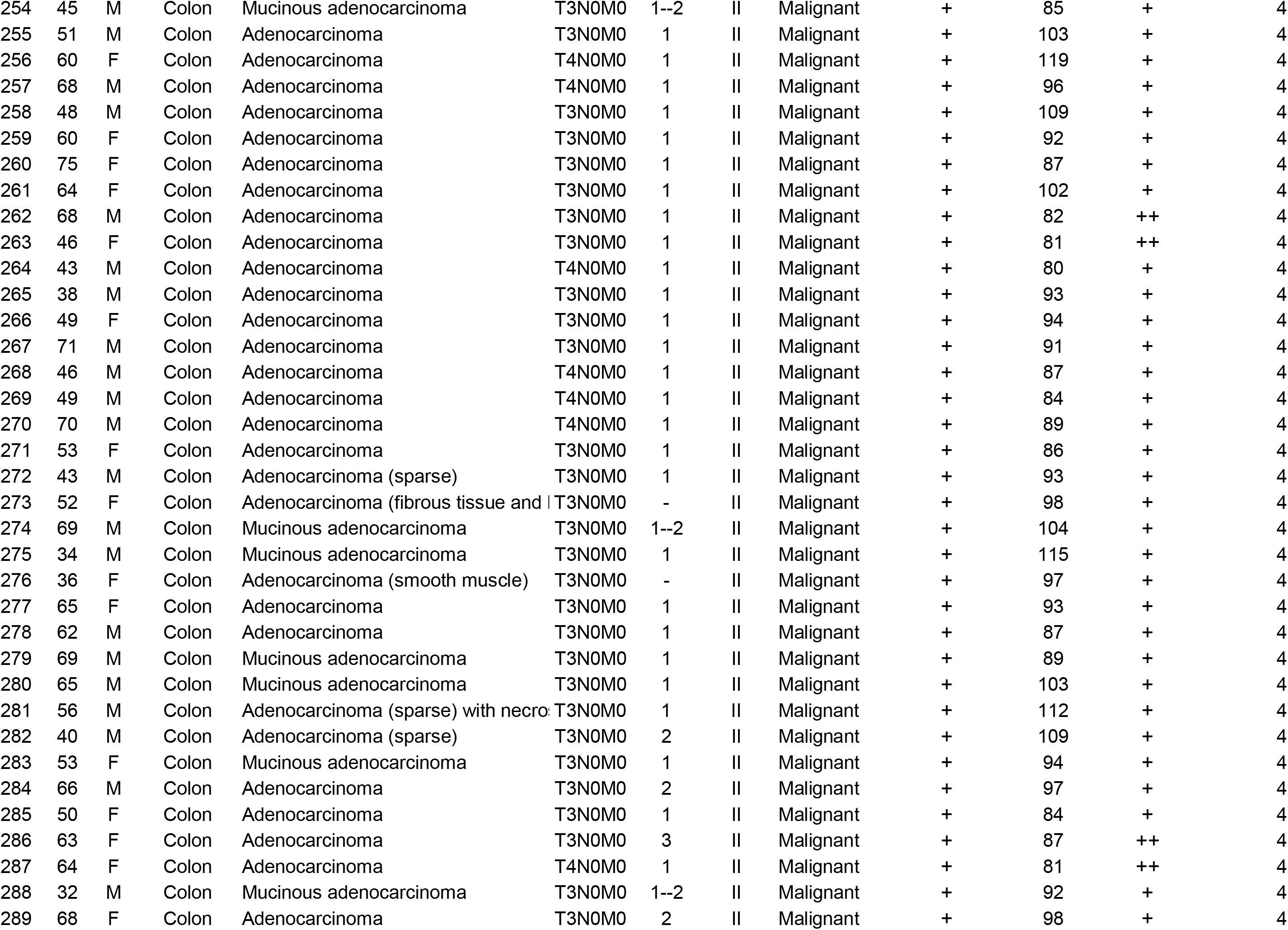
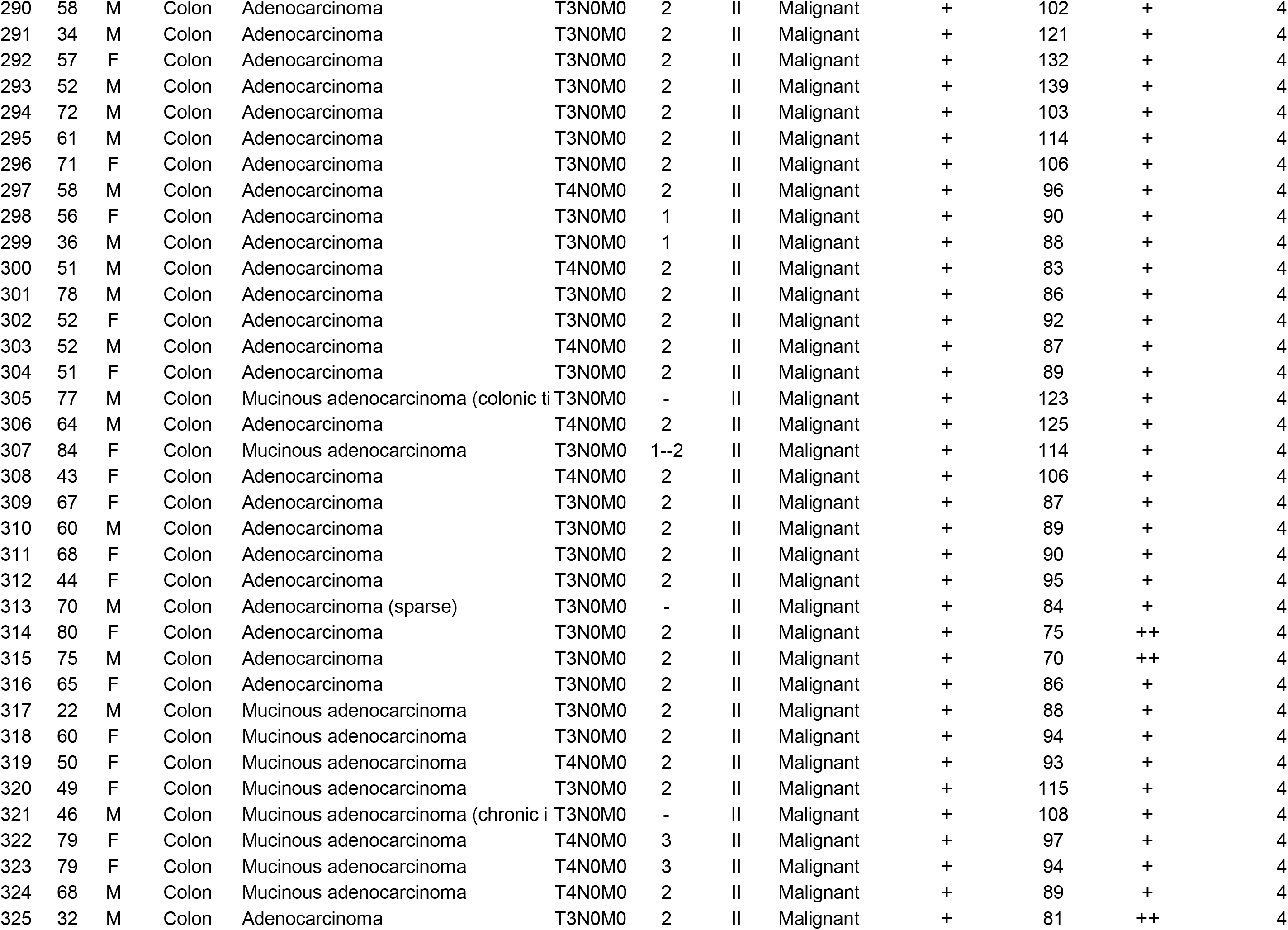
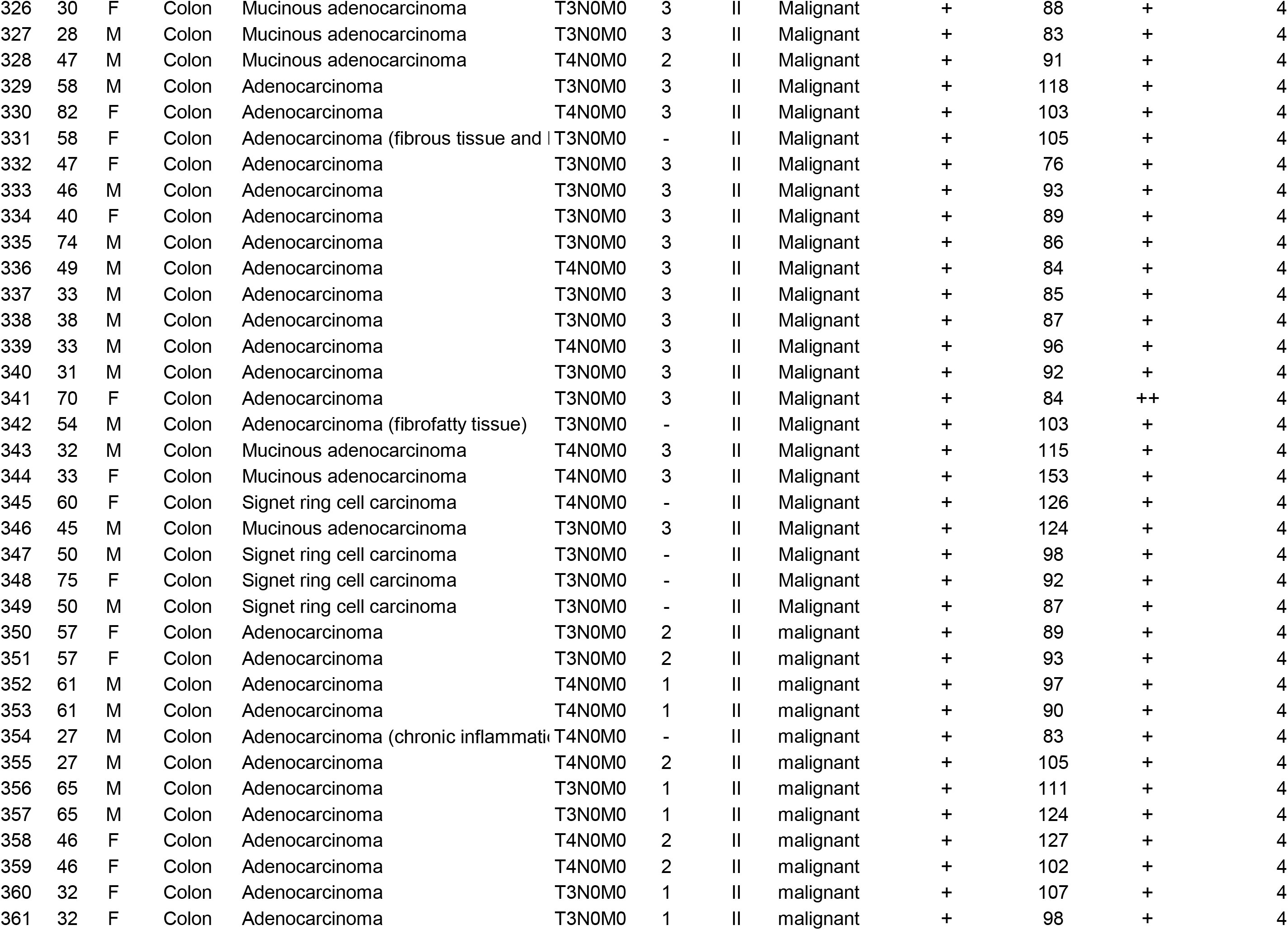
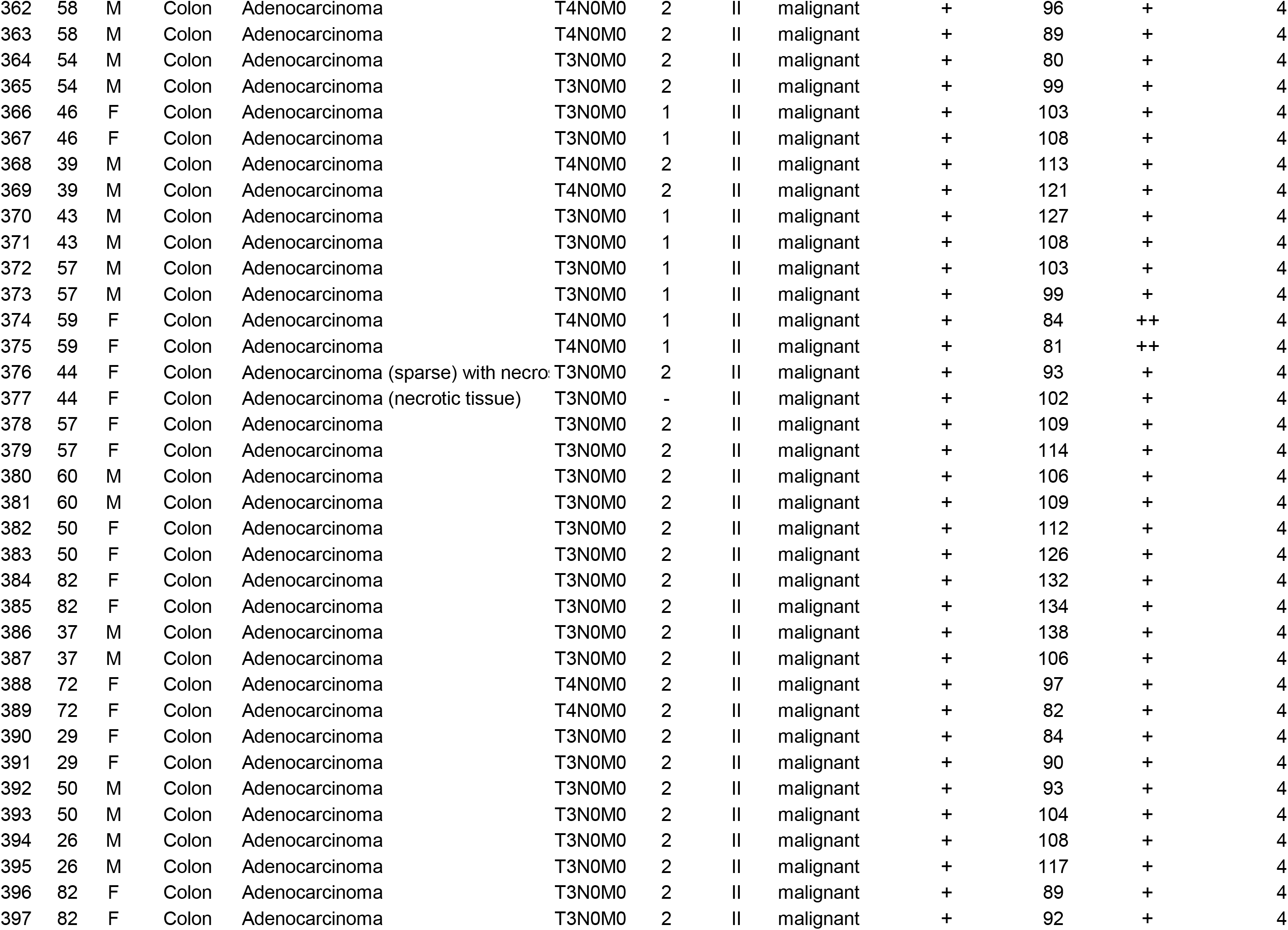
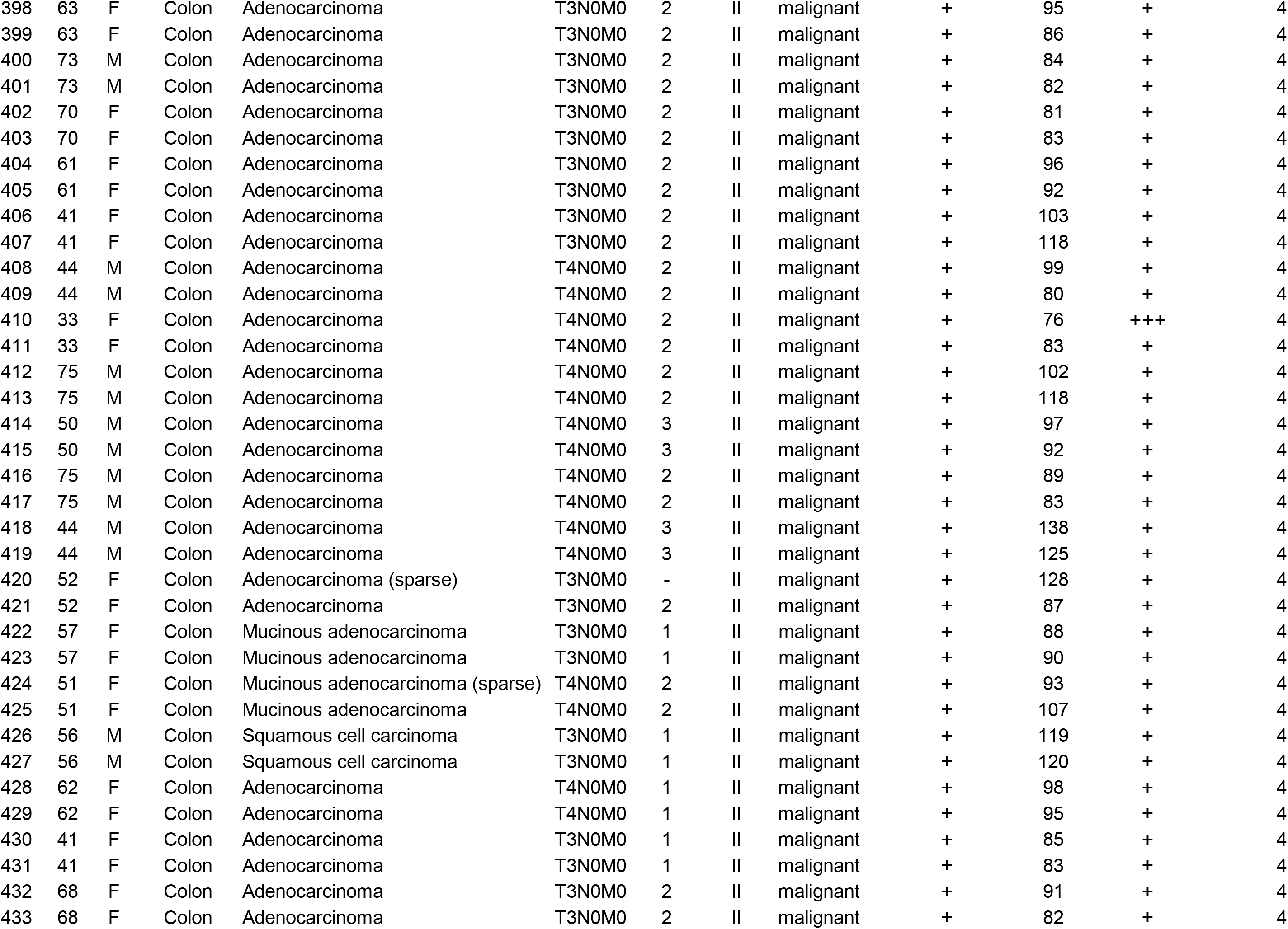
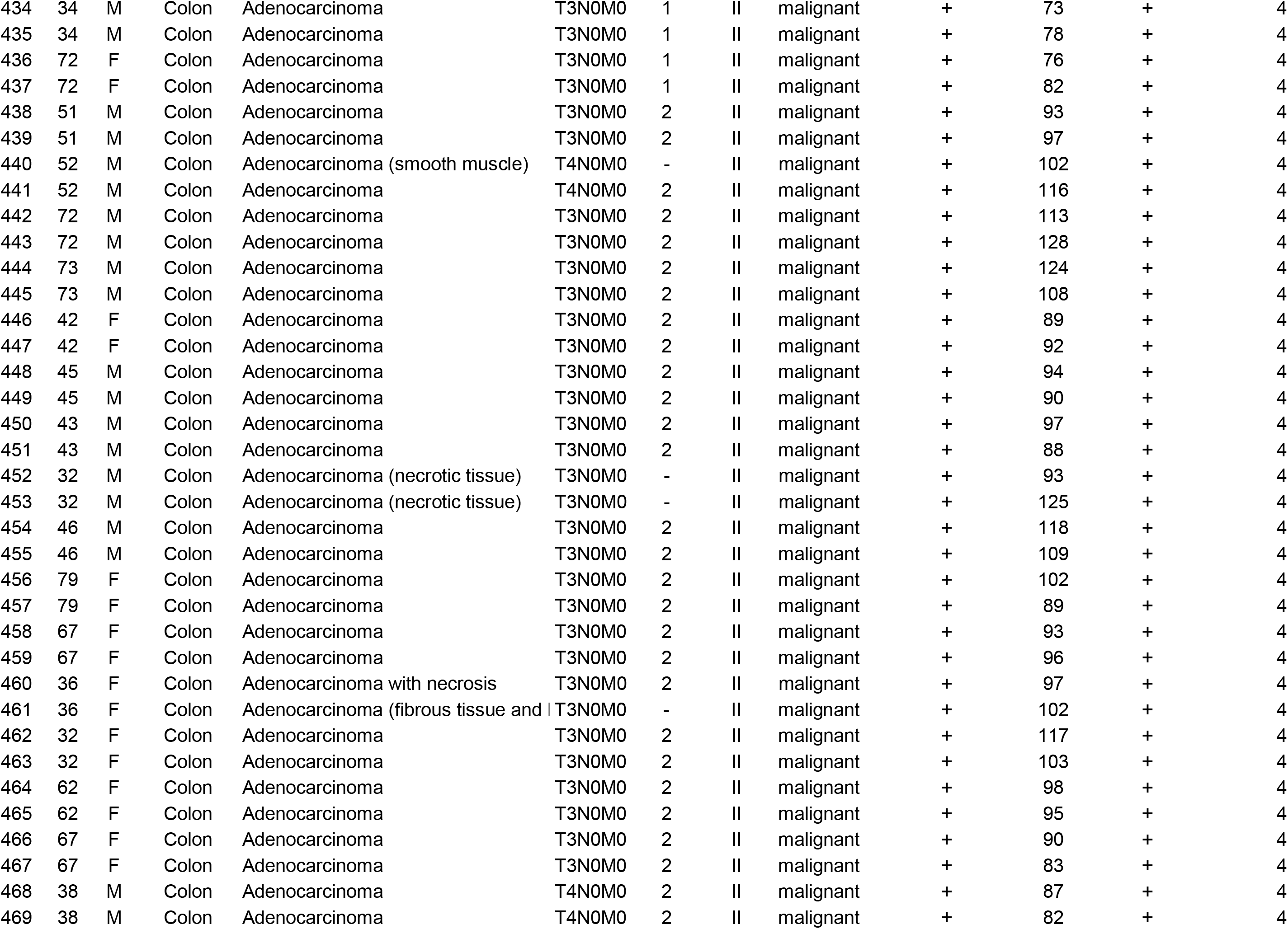
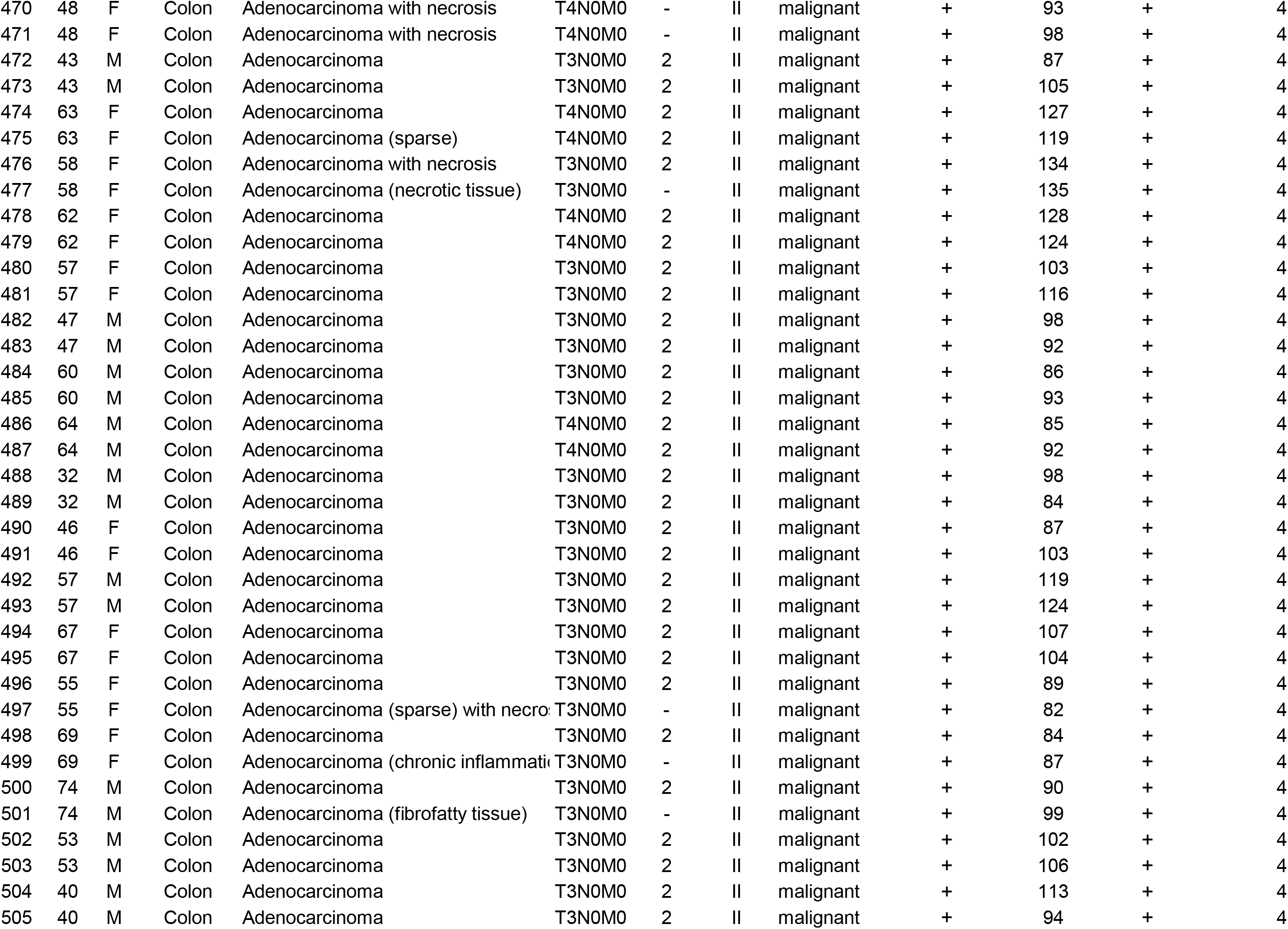
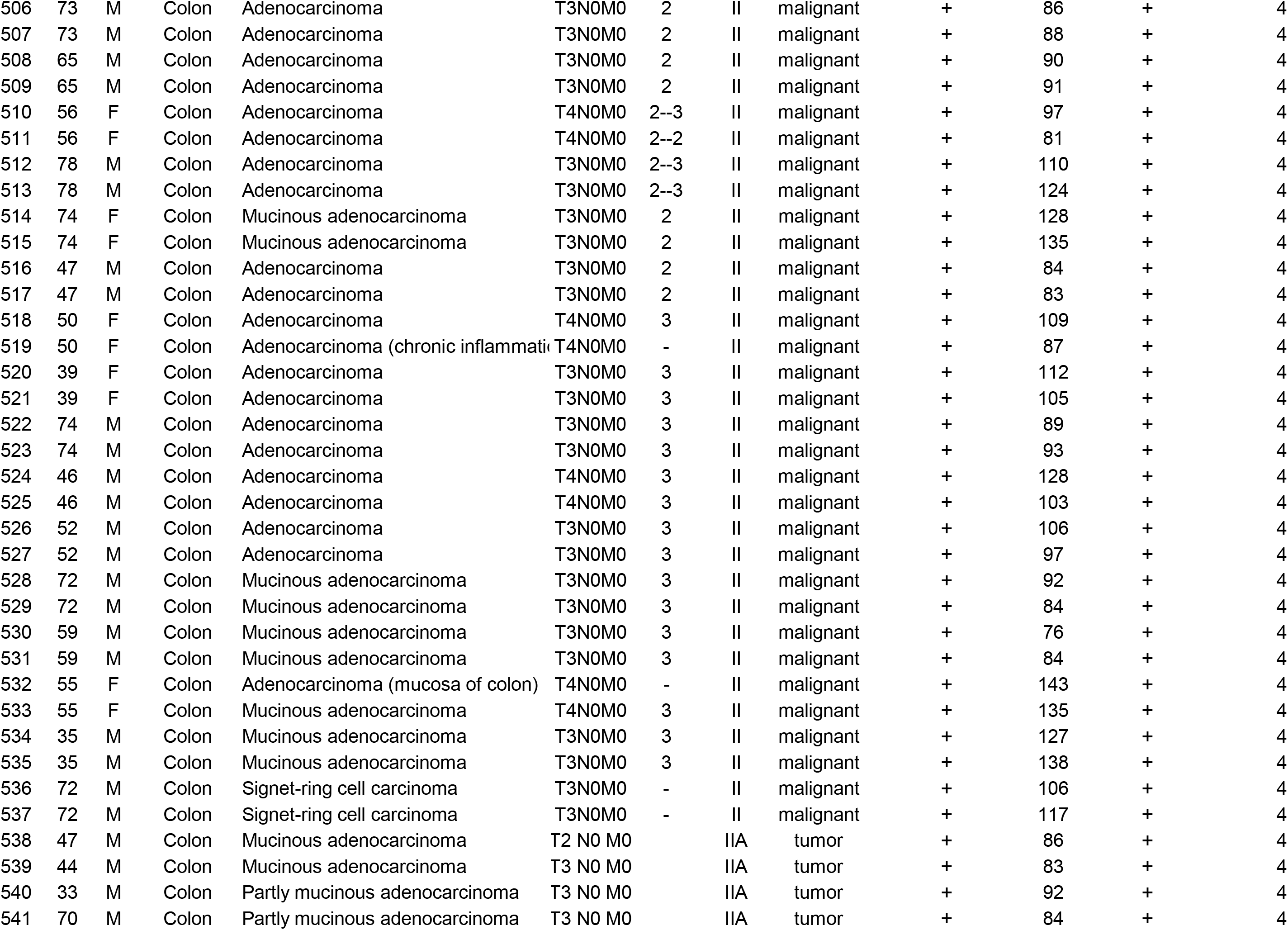
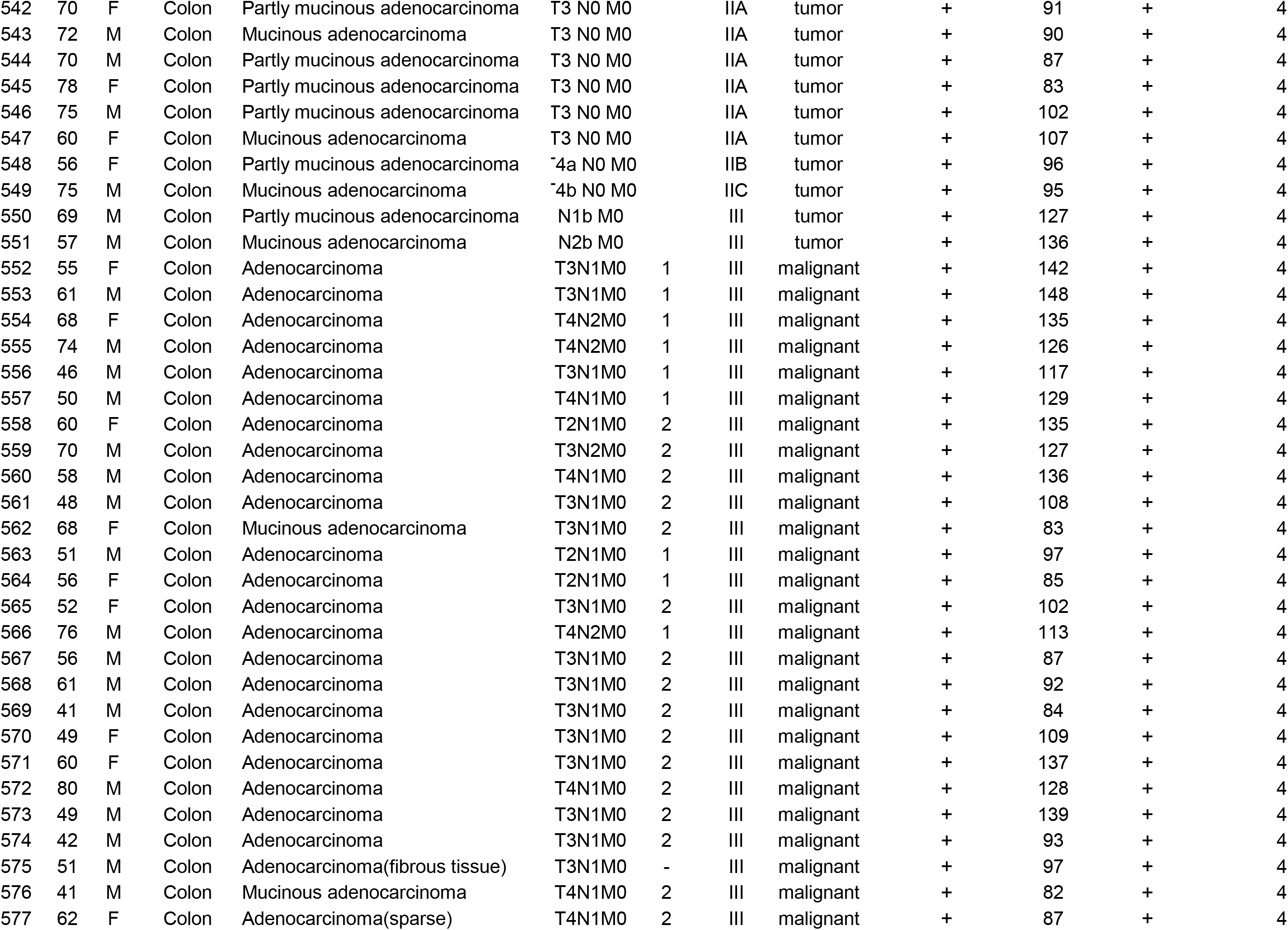
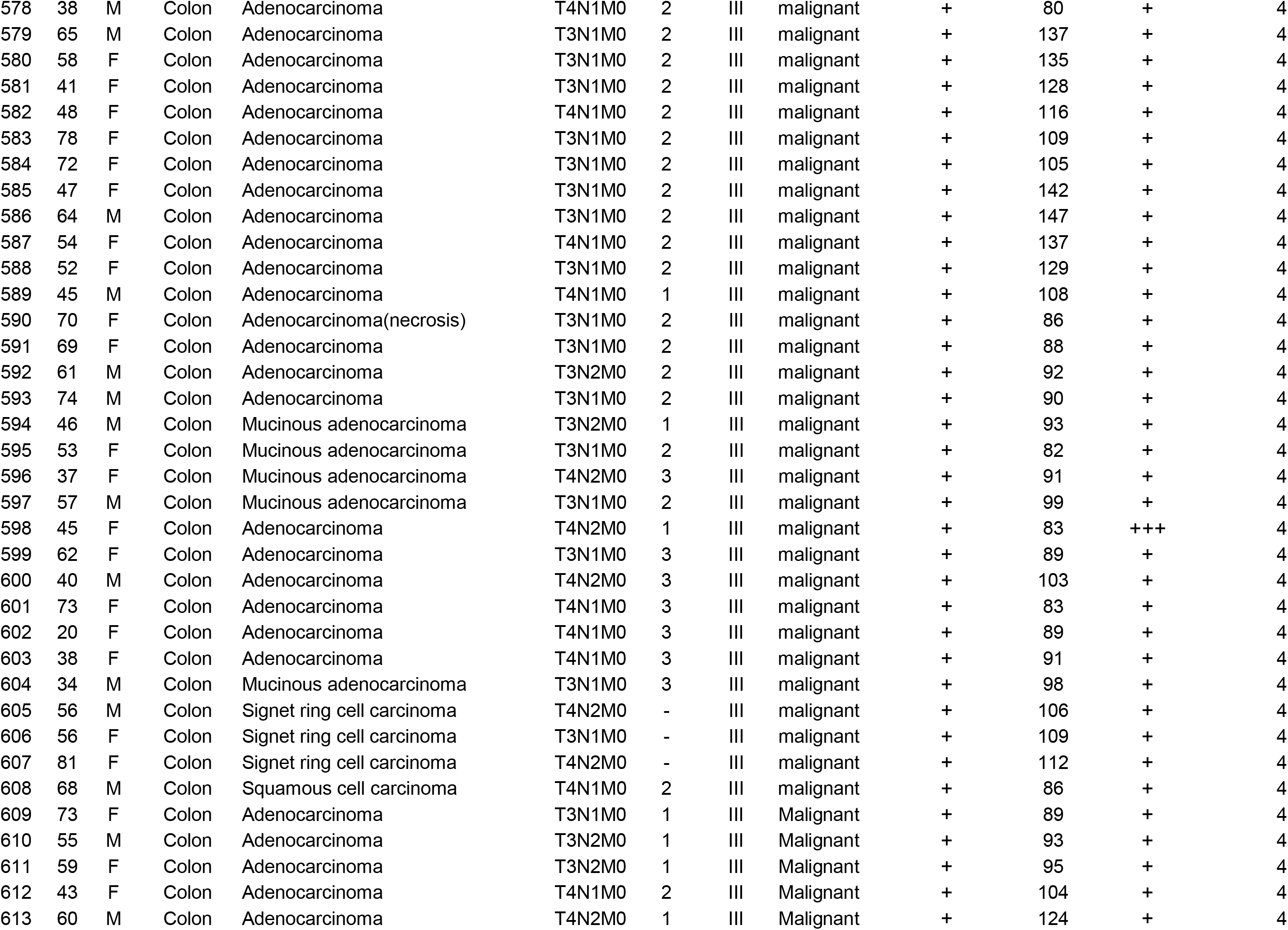
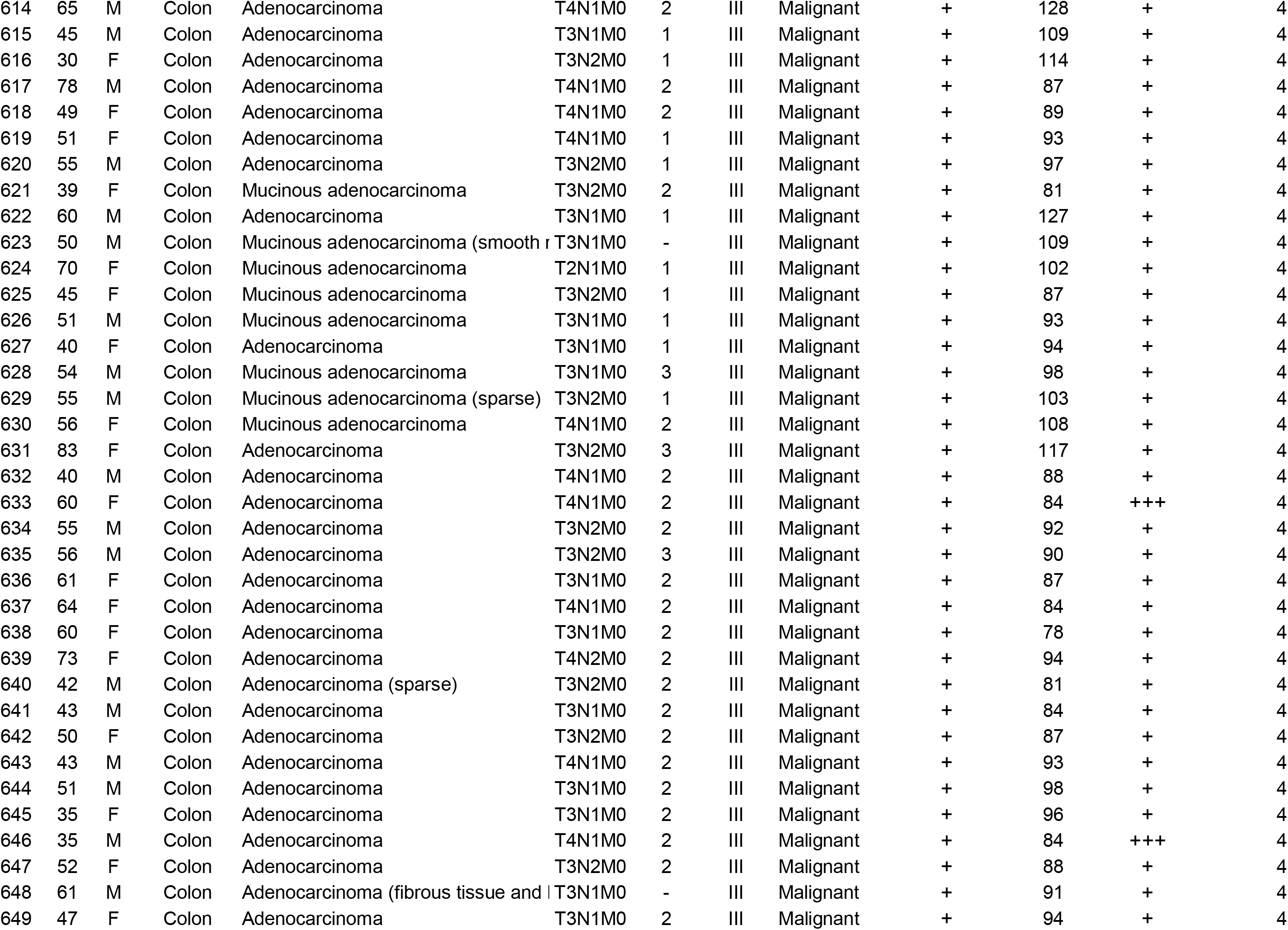
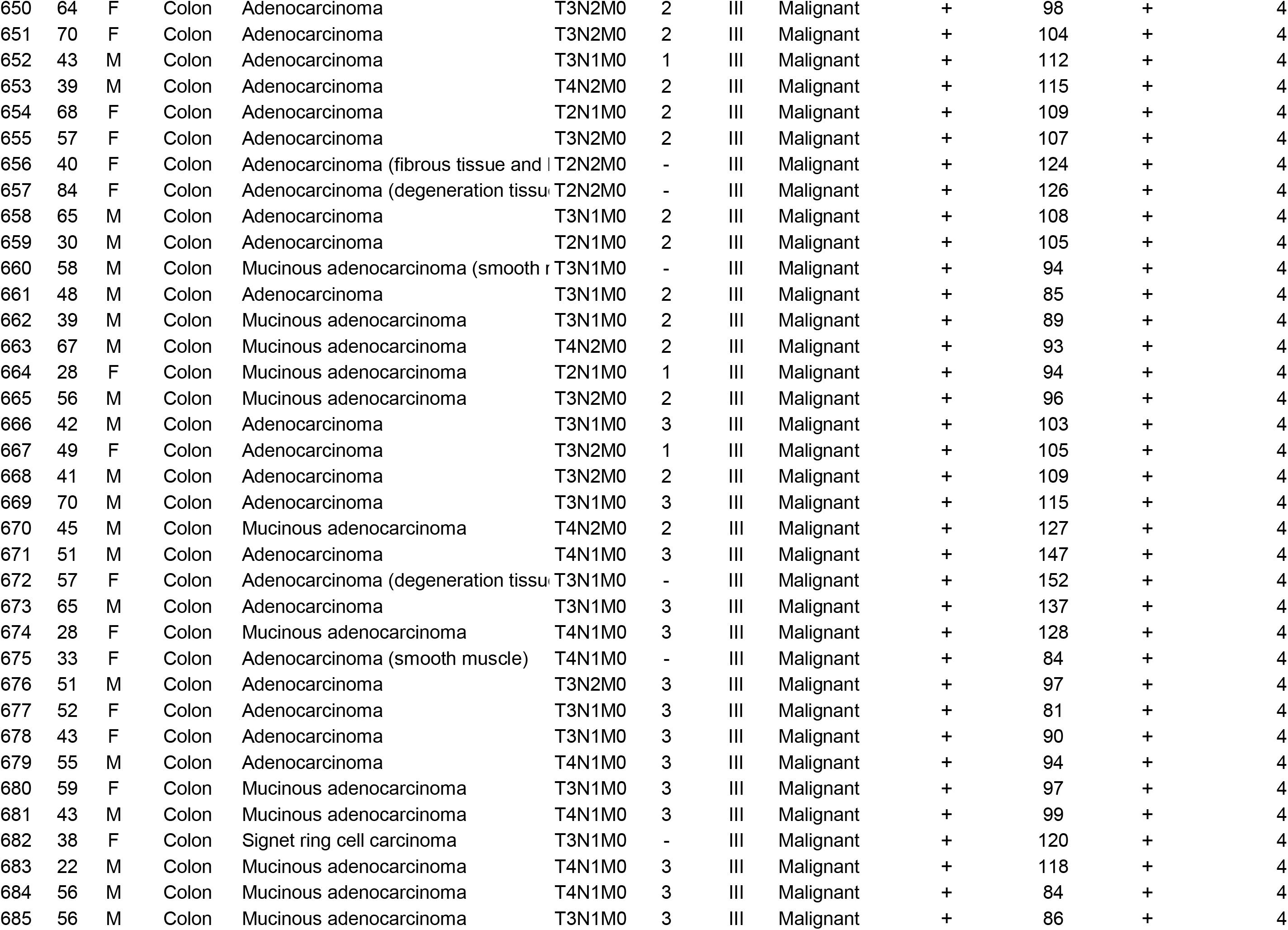
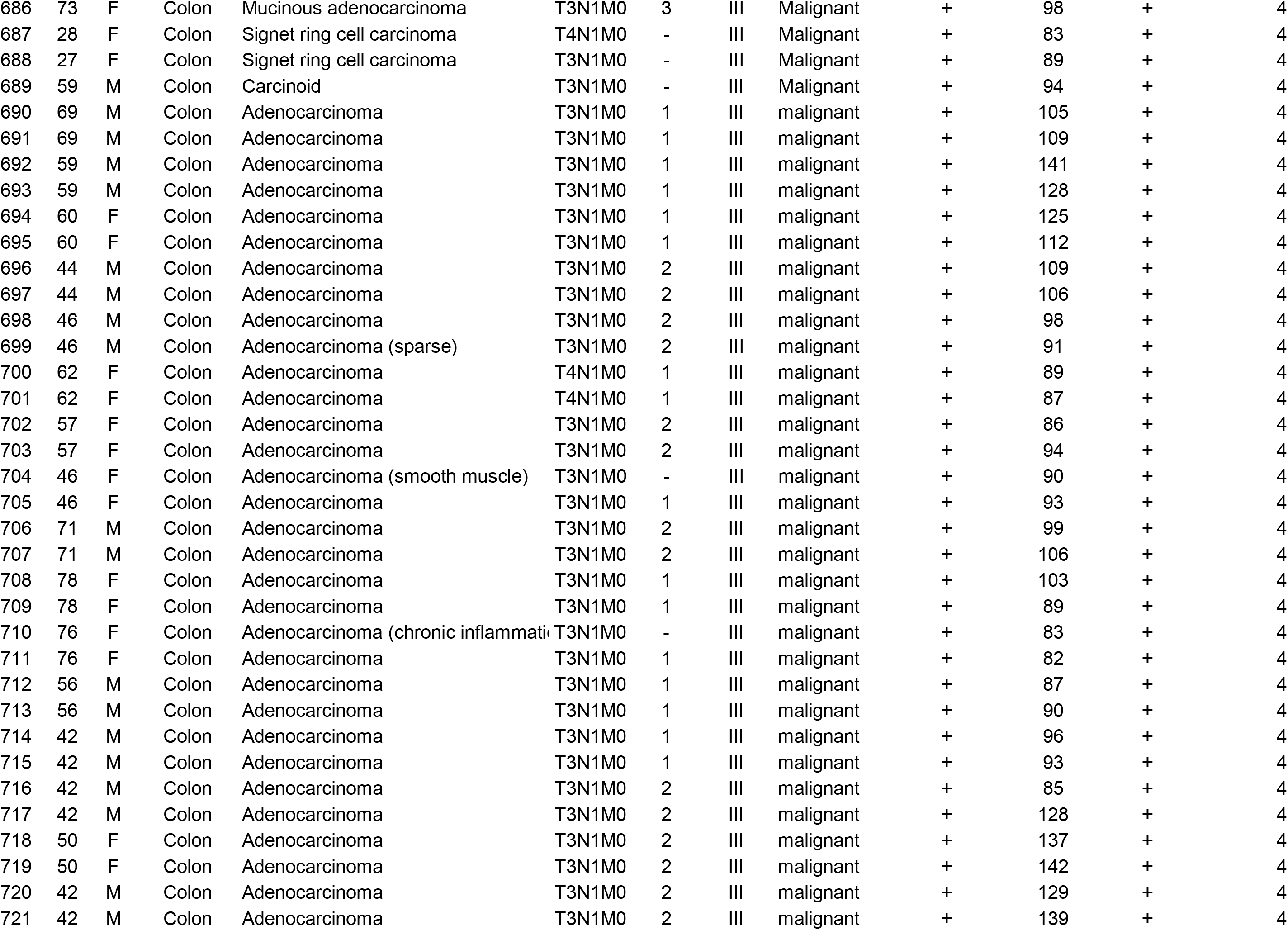
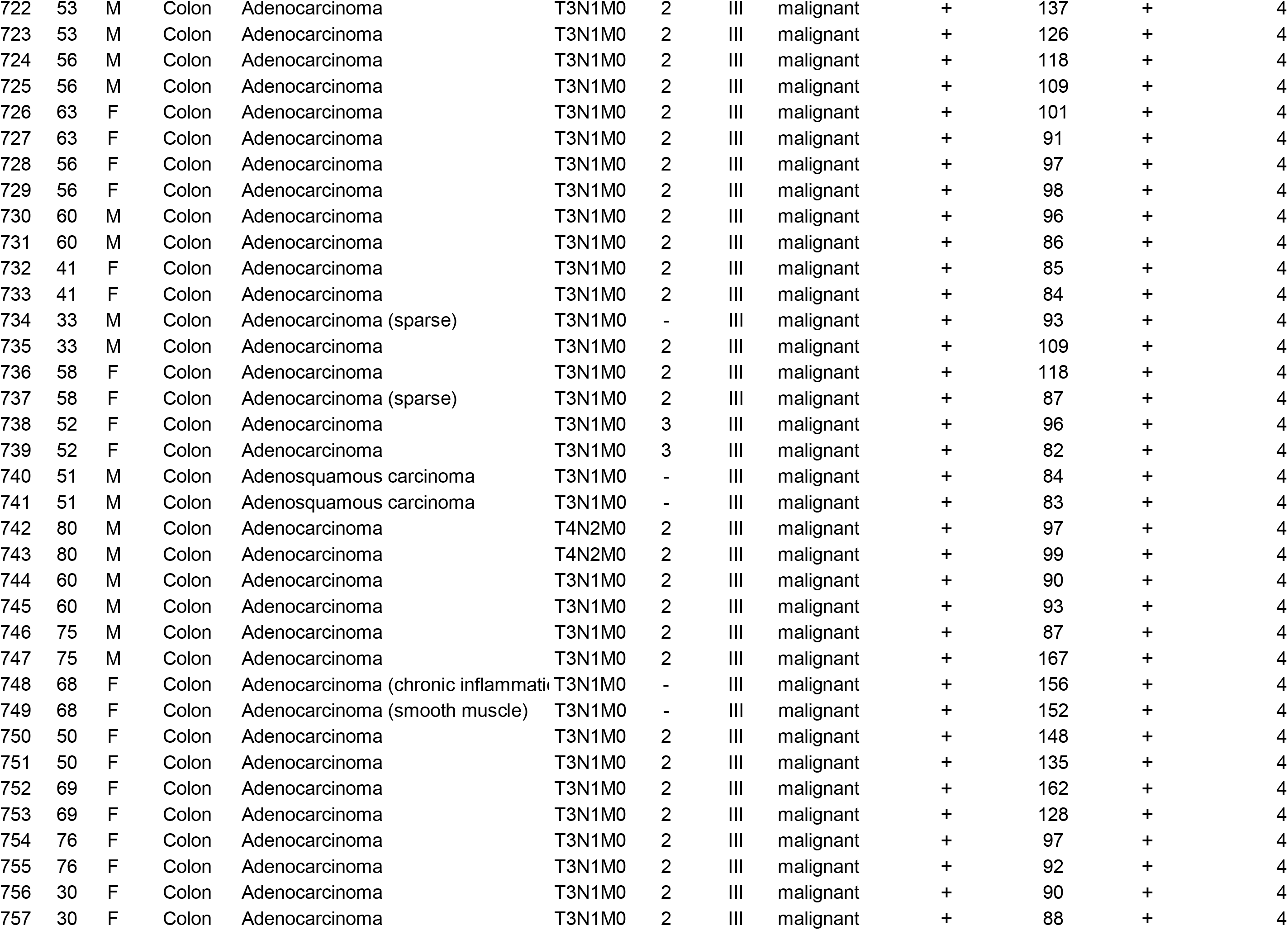
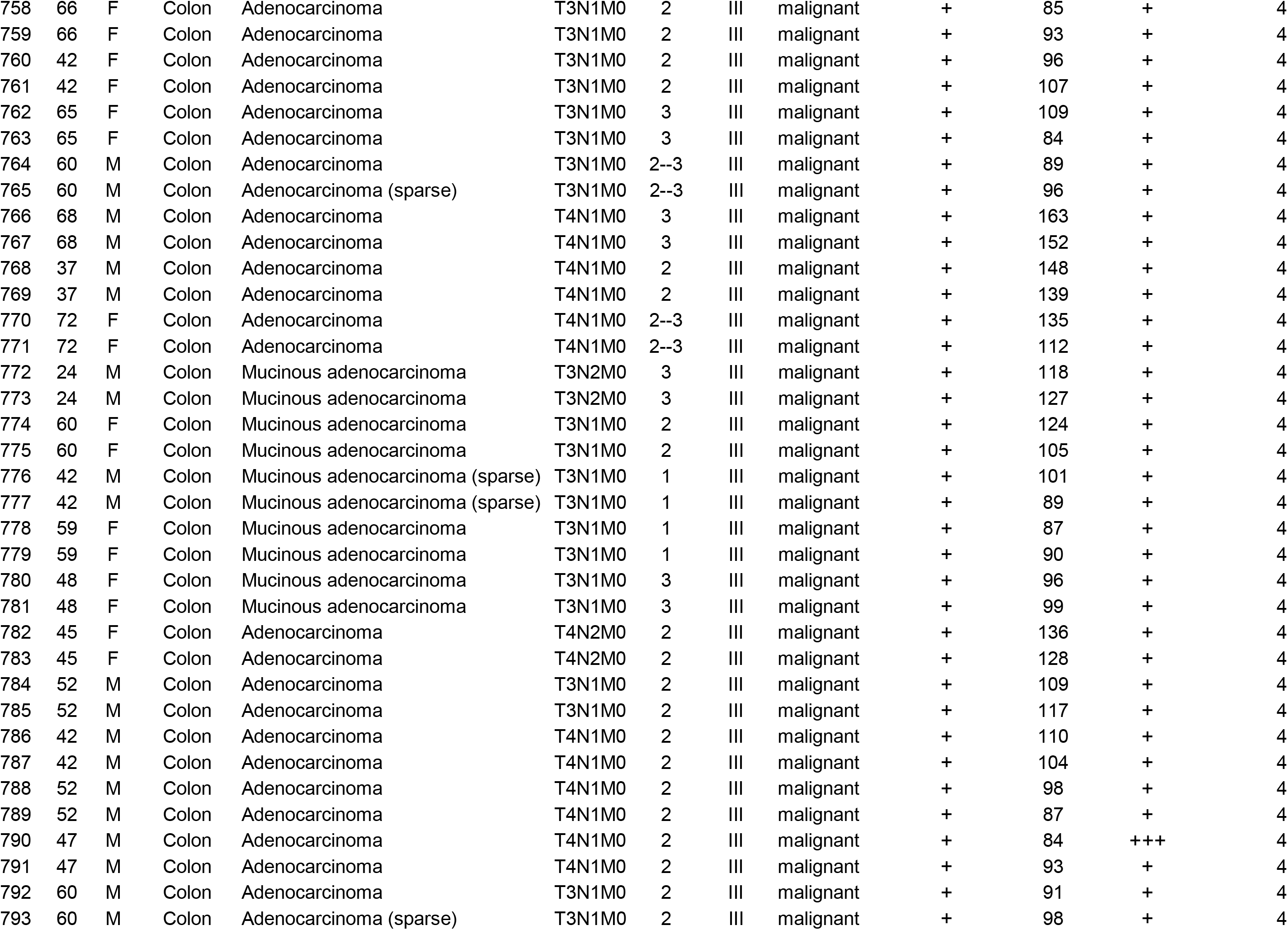
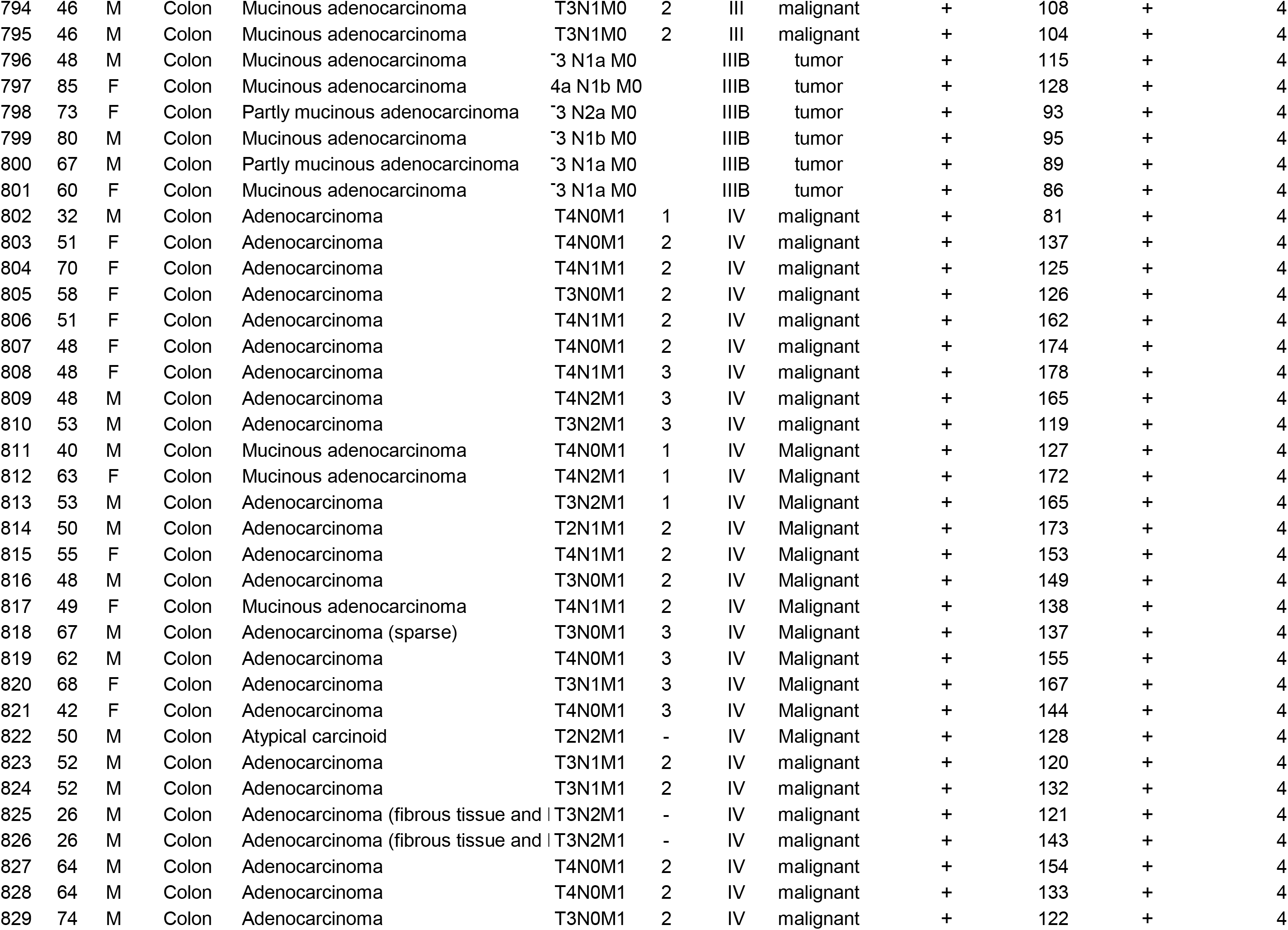
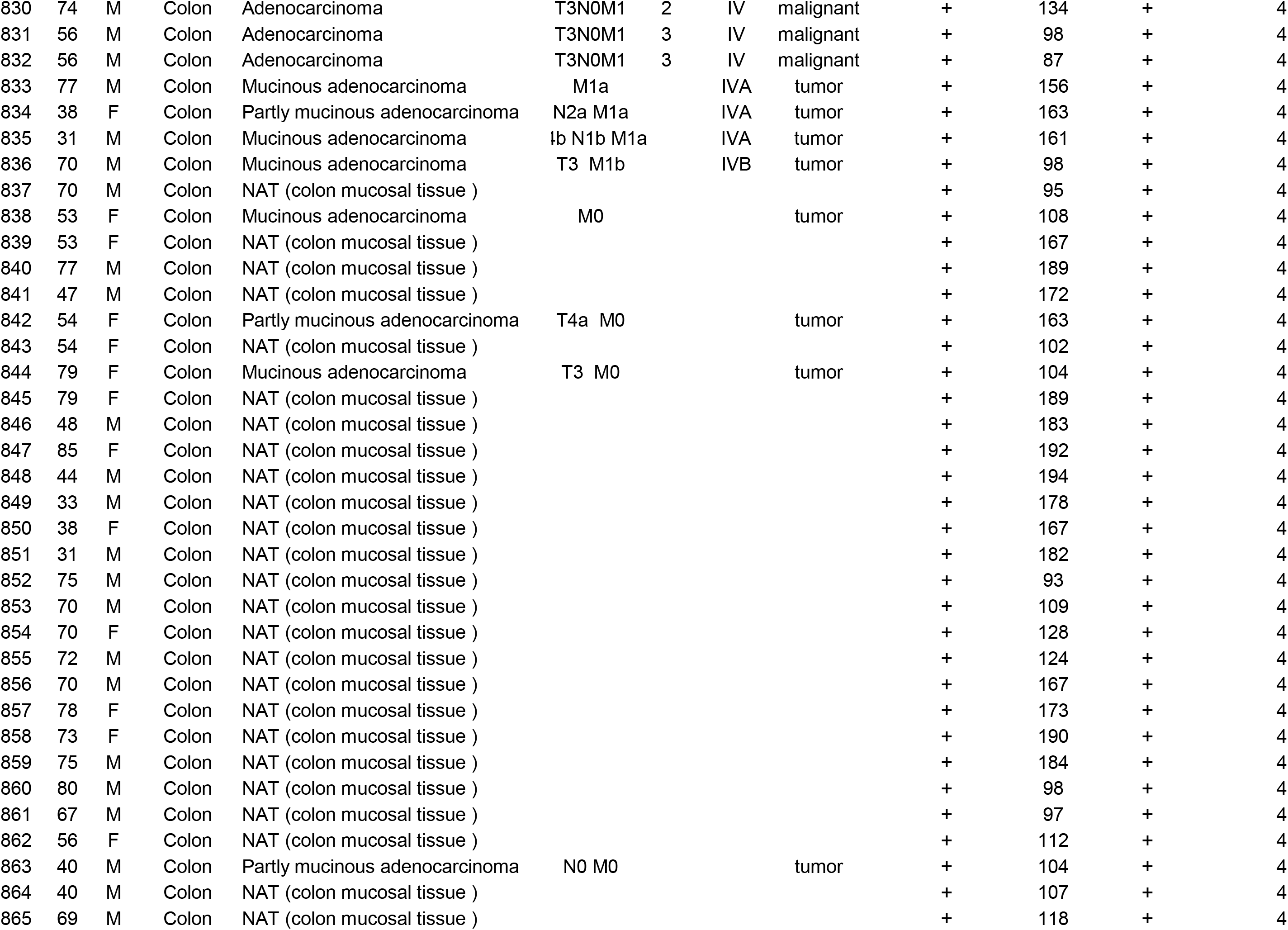
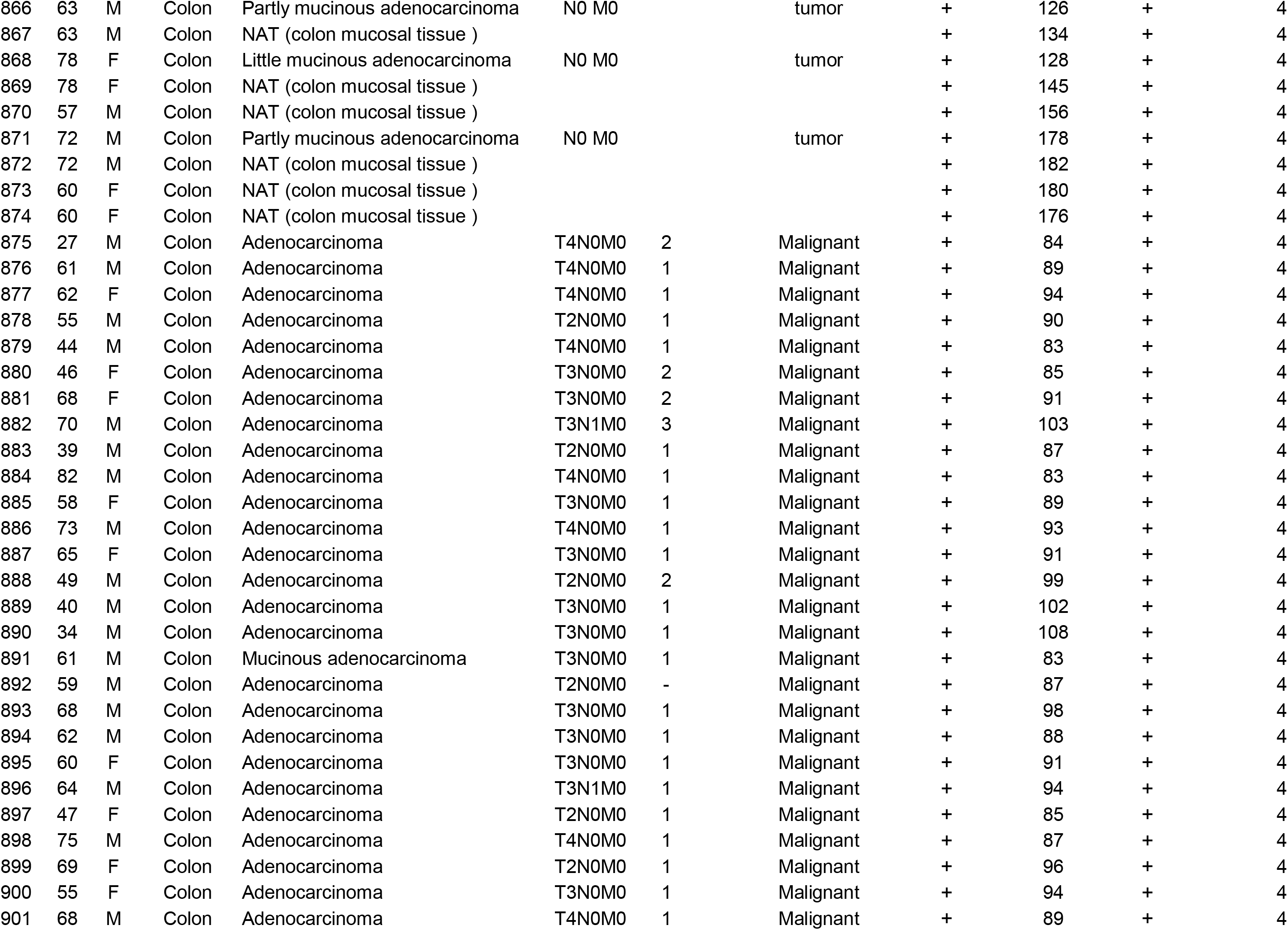
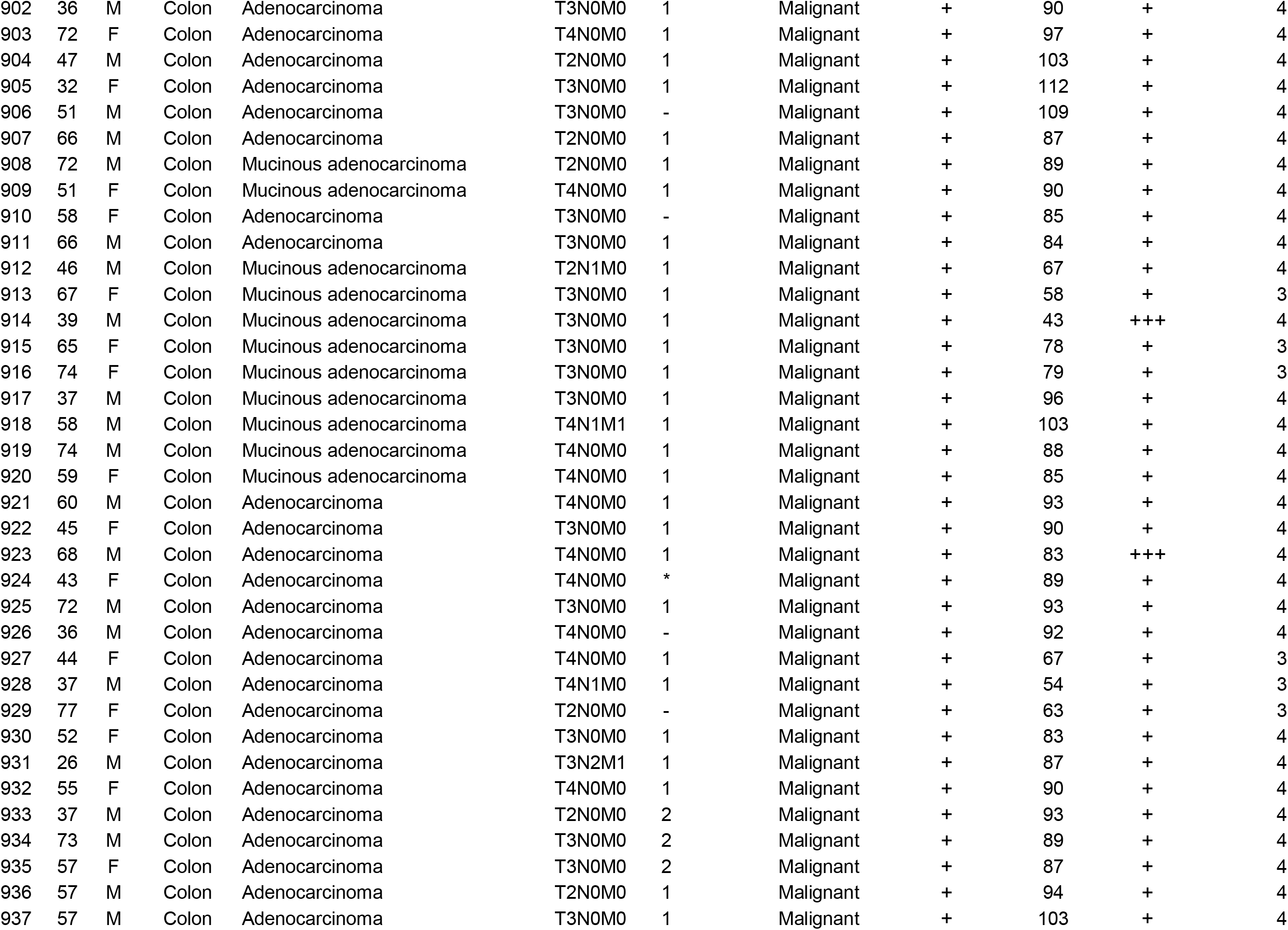
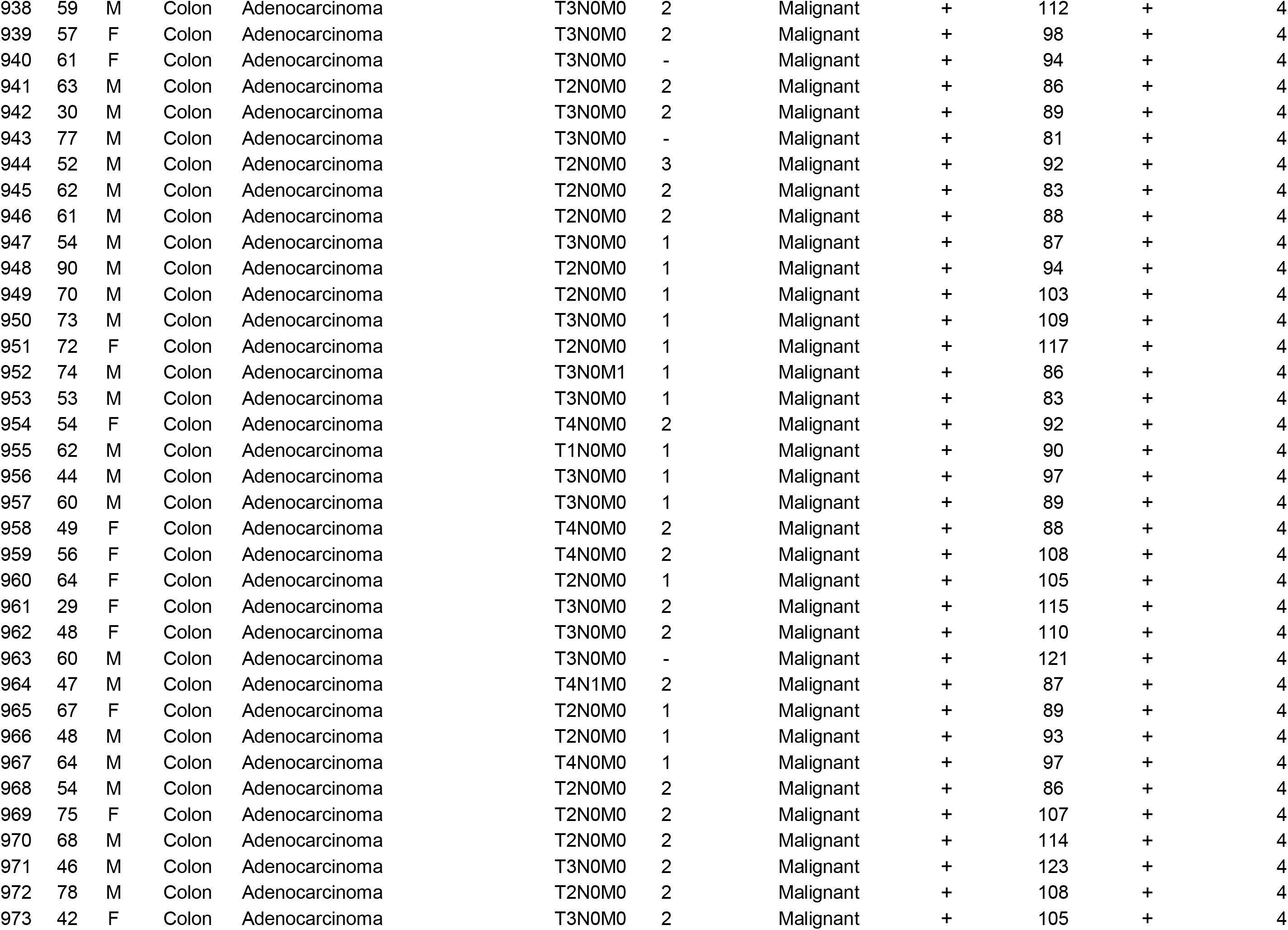
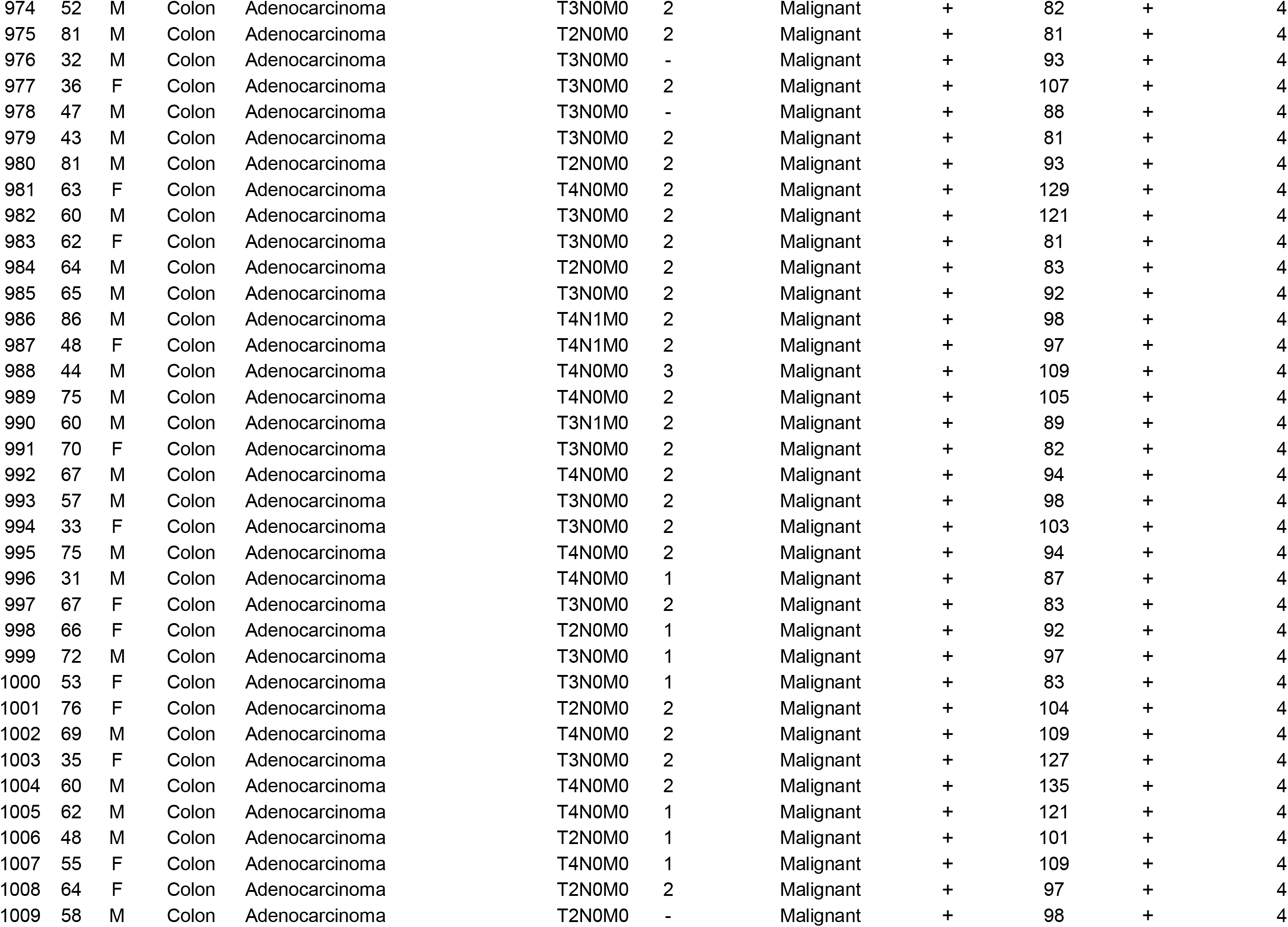
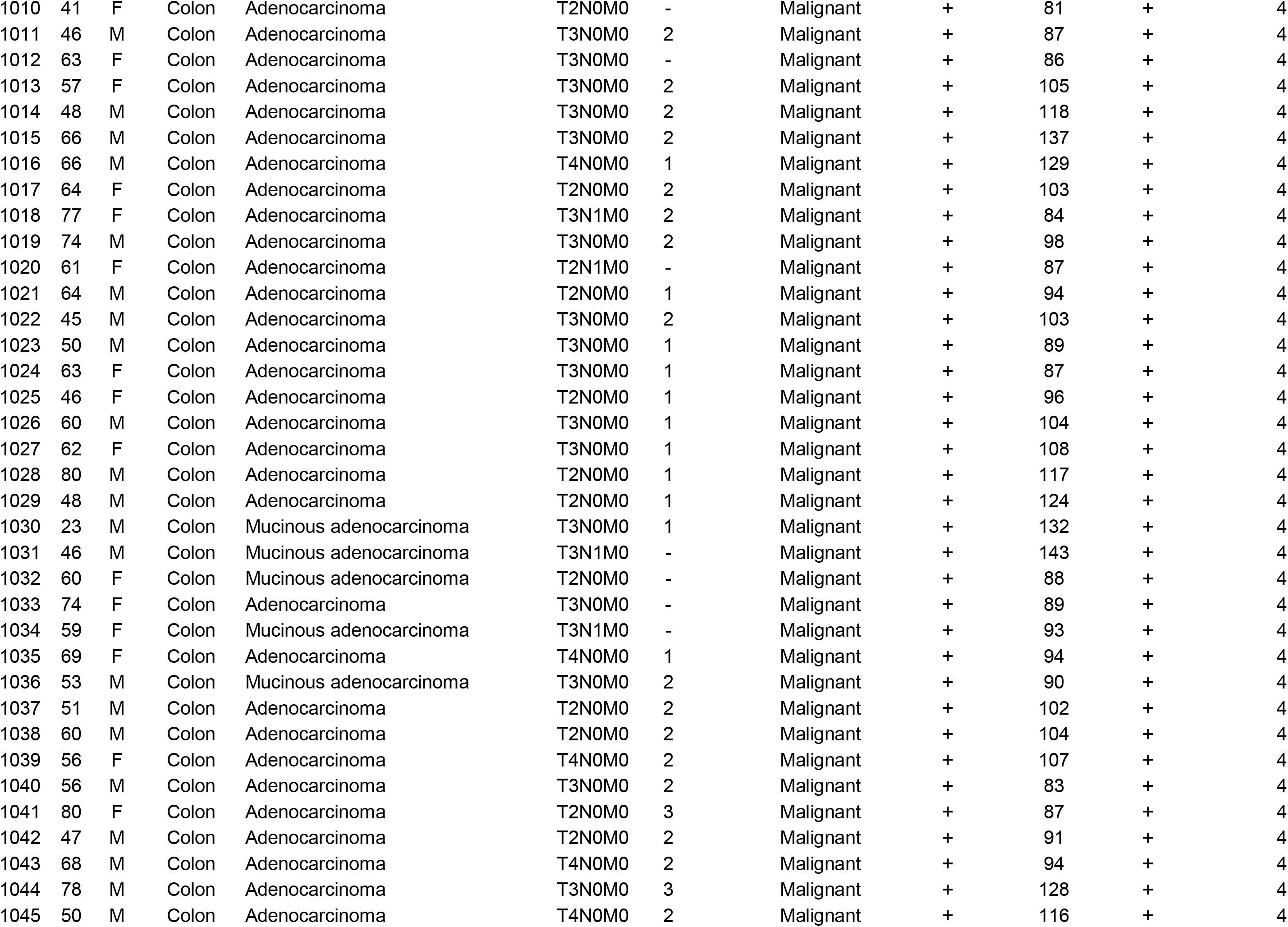
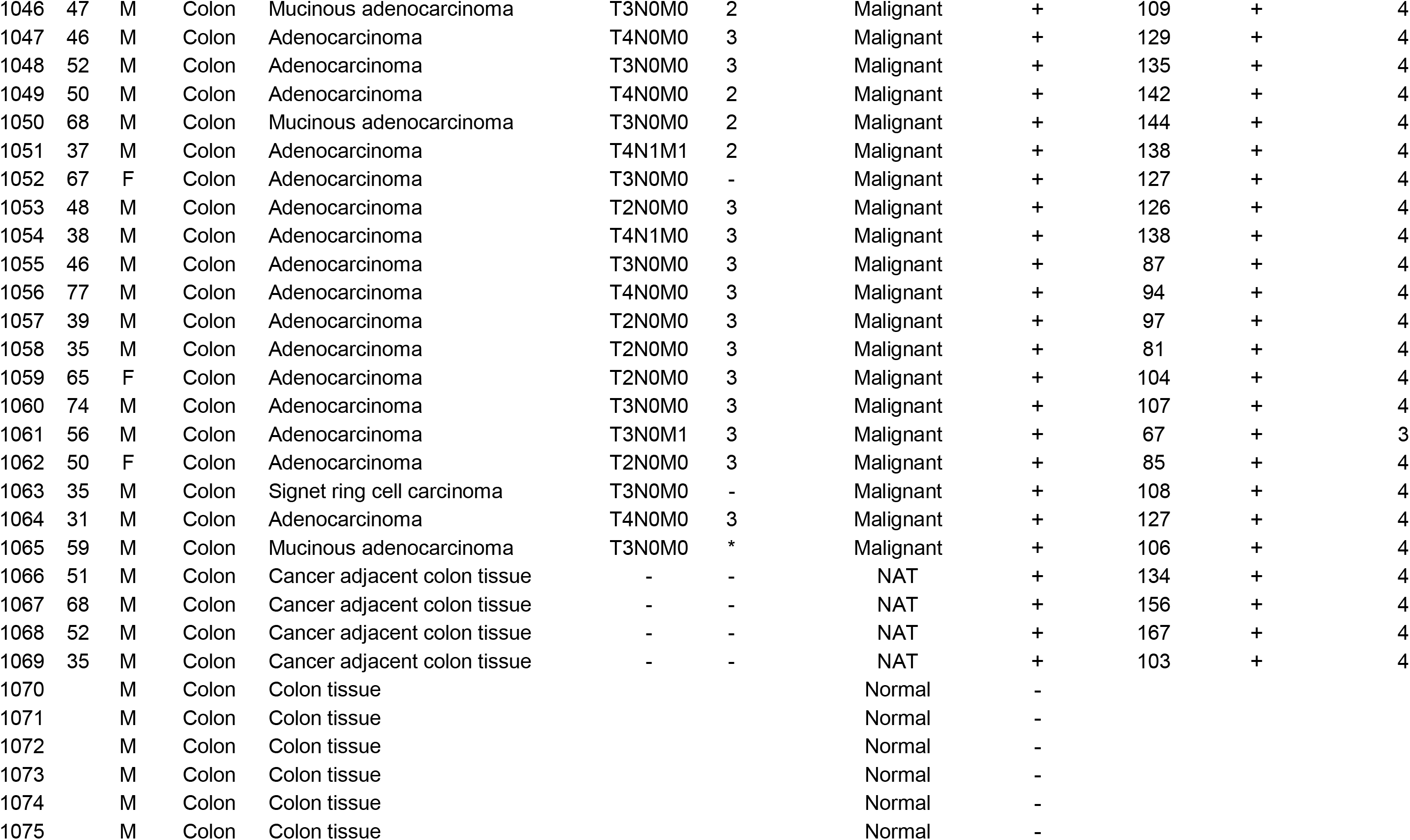

